# Social bonding, gestural complexity and displacement behaviour of wild chimpanzee

**DOI:** 10.1101/678805

**Authors:** Anna Ilona Roberts, Sam George Bradley Roberts

## Abstract

Kinship and demography affect social affiliation in many different contexts such as co-feeding, resting, travel, grooming, visual attention and proximity. Chimpanzees may coordinate these social interactions by using gestural communication to make signaller’s goal transparent to the recipient and also by increasing commitment of the recipient through including rewarding property in communication. The rewards of gesturing can be measured through the rates of displacement behaviour made in response to these gestures by the recipient. We tested hypothesis that gestural communication affects social affiliation after controlling for kinship and demography in wild, adult chimpanzees living in Budongo Forest, Uganda. We found that affiliative but not antagonistic gestures positively predicted social affiliation. Contexts differed in their association with gestures according to complexity and association with displacement behaviour. More complex, less intense gestures predicted mutual grooming, travel, visual attention whereas less complex, more intense gestures predicted unidirectional grooming. Mirroring these patterns, reduced displacement activity occurred in response to gestures associated with unidirectional grooming but not other contexts. We highlight that these tactical decisions that wild chimpanzees make in their use of gestural communication may be driven by complexity of social environment that influences effectiveness with which signalers can influence the recipient.

## Introduction

Social bonds of animals, whereby particular dyads within the group preferentially associate and affiliate with each other have fitness benefits by regulating within-group competition in contexts such as feeding and mating. Such relationships have been found in numerous animal species which include for instance, bottlenose dolphins (Tursiops aduncus) (Möller, Beheregaray, Harcourt, & Krützen, 2001), Bechstein’s bats (*Myotis bechsteinii*) (Kerth, Perony, & Schweitzer, 2011), and sperm whales (*Physeter macrocephalus*) (Ortega-Ortiz, Engelhaupt, Winsor, Mate, & Rus Hoelzel, 2012). By far however, the most dynamic and stable societies are those of primate species, and particularly great apes such as chimpanzees whereby conspecifics regularly form social bonds manifest in many different contexts (F. B. M. De Waal, 1982; Goodall, 1986). Unsurprisingly perhaps, considerable research effort has been directed at explaining the emergence and persistence of social bonds in primates (K. Langergraber, Mitani, & Vigilant, 2009; John C Mitani, Watts, Pepper, & Merriwether, 2002; J. B. Silk, Seyfarth, & Cheney, 2018). At first, inclusive fitness behind social bonding with kin was ubiquitously claimed to drive patterns of social bond formation (Hamilton, 1964). Although primates can and do socially bond with kin (Wrangham, 1980), the principal role of kinship in driving primate sociality has been discounted on the basis that in many socially complex primate species such as gorillas, chimpanzees and the bonobos social bonding within dyads is not determined by kinship (F. B. de Waal, 1986; John C Mitani et al., 2002; Watts, 1992). One another explanation that has been provided for existence of social bonds in primates, is that demographic constraints such as membership of the same age cohort can foster social bonding due to potency of peer influence on one’s behaviour and familiarity inherent in growing up in the same social context (Goldberg & Wrangham, 1997; Van Schaik & Van Hooff, 1994). For those reasons, primates of same reproductive status or sex, may also be found to have affiliative contact with each other at a higher rate than among primates of different reproductive status or sex. For instance, male chimpanzees are more likely to groom with other males than the females (John C Mitani et al., 2002). Thus, formation of social bonds in primates, often demands that some similarity between interactants exists (Massen & Koski, 2014). However, primate social groups are complex and large, and primates often come into contact with conspecifics who are dissimilar and/or unrelated. For these complex social groups of unrelated individuals to form and persist over time, the social cohesion mechanism has to evolve that facilitates these social interactions. We propose that gestural communication defined as voluntary movements of the hands, legs, head, bodily postures or locomotory gaits (Hewes, 1973; Liebal, Call, & Tomasello, 2004) is one important mechanisms that facilitates formation of social bonds between individuals regardless of similarity.

The gestural communication is highly variable across species but within species, there is also a considerable variation that may facilitate choice of a social partner on the basis of several gesture features. For instance, gestural communication can be multimodal and co-occur with facial expressions or vocalisations, or it can be unimodal. Primates may incorporate objects such as branch of a tree in signalling or the gesture may not contain use of objects. Gestures can be combined with other gestural signals in a single utterance, or gesture may be produced singly (A. I. Roberts, Vick, Roberts, Buchanan-Smith, & Zuberbühler, 2012). The variation in gestural communication may be associated with the degree of social bonding, because gestures vary in the efficiency with which signaller can influence behaviour of the recipient (A. I. Roberts, Vick, Roberts, & Menzel, 2014). Signals of greater complexity that deviate from the population mean, may be more effective at influencing the recipient, than the average signals (Dawkins & Guilford, 1997; Zahavi & Zahavi, 1997). There are however many different pathways to achieving a greater efficiency in gestural communication. Studies in humans, showed that during perception of facial expression accompanied by direct gaze and pointing, the amygdala processes emotional content of communication. However, there is a concurrent integration of the directional communication (pointing, gaze) in the premotor cortex that facilitates re-evaluation of prior expectations regarding perceived signaller’s immediate intent from emotional expression alone (Conty, Dezecache, Hugueville, & Grèzes, 2012). Thus, pointing gestures with facial expressions have a greater signal value than facial expressions alone and they can elicit a more accurate response from the recipient. Recently studies documented similar behavioural strategies in primates (S. Roberts & A. I. Roberts, 2018). Chimpanzee females, for example, lower back to make infant climb on it for travel as well as combining this bodily gesture with a manual sweep backwards in direction of the infant, potentially using manual directional gesture to draw the attention of the infant to the bodily gesture even though the infant is only a meter away and can respond to bodily gesture alone. When initiating grooming, male chimpanzee may present the bodily part that he wants to be groomed by the recipient, as well as making almost imperceptible sound to an observer standing only a short distance away, but clearly of fundamental importance to the recipient.

The social relationships of primates, however, may not always be guided by selection for effectiveness of communication (Dawkins & Guilford, 1997). If both interactants have mutual interest in signalling and responding then the recipient may become sensitive to the signaller, whereas signaller may make communication less complex. These scenarios are evolutionarily stable because the communication is effective at achieving its desired goal, whilst reducing the costs of potential harassment by third party or predators in circumstances when the signal is conspicuous and large. An example would be left handed gesture seen in a common goal contexts signifying stronger social bonding (e.g. mutual grooming, joint travel, mutual visual monitoring) versus right handed gestures seen in contexts whereby interactants do not have a common goal signifying weaker social bonding (e.g. unidirectional grooming, lack of mutual visual monitoring) (Noë, 2006). Studies have recently emphasized how low intensity of communication is important for fitness by reducing stress inherent in social interaction (Nakayama, Goto, Kuraoka, & Nakamura, 2005; A. I. Roberts & S. G. B. Roberts, 2016). When individuals who are less strongly bonded interact, there may be a greater conflict of interest and therefore low intensity communication may be ineffective at influencing behavioural change in the recipient (Dawkins & Guilford, 1997). There is a wealth of studies that describe the phenomena whereby recipients respond more strongly to visual communication that is complex than to visual communication that is non-complex. Possibly, the most well-known are the elaborations and repetitions of visual signals seen in mating contexts. For instance, male frogs incorporating a higher rate of action repetition in visual display are more likely to attract female for mating than the males incorporating lower rate of action repetition in the visual display (PAYNE & PAGEL, 1997).

Complex visual signals as effective means of signalling, however, may reach a limit at which further increasing of complexity will not have any bearing on effectiveness of interaction with the recipient. Another way, in which signallers can influence efficiency of their signalling is by stimulating the reward system of the recipient. By making a gesture rewarding, signaller enhances recipient’s commitment to the interaction, thereby making these social relationships possible. Such circumstances, for instance, may be present in complex social settings, whereby presence of other conspecifics in vicinity, provides competitive background against which signaller competes for attention of the recipient. By increasing their rewarding value, signals become more pleasurous to the receivers and to that extent become more capable of evoking a response from the recipient. During grooming, chimpanzees sometimes direct visual gestures to make the recipient present body for grooming, but when recipient is unresponsive they switch to gentle, but more intense tactile gestures to achieve their goal (A. I. Roberts, Vick, & Buchanan-Smith, 2013). Higher intensity signals are more perceivable by the receivers and by virtue of being more intense can exert greater influence on the recipient (Zahavi & Zahavi, 1997). Subordinate male chimpanzees for instance, use visual gestures designed to attract female for mating in absence of visual attention from the dominant male, but these gestures become intense and loud when the female is unresponsive because the dominant male is watching the interaction (A. I. Roberts & Roberts, 2015). High intensity however, can reduce or even reverse the preference for signals that are based on complexity of structure hence high intensity signals can become less complex (Gerhardt & Doherty, 1988). For instance, chimpanzee visual gestures are often accompanied by high complexity such as use of mutual attention, low intensity call or synchronized call in contrast to tactile and auditory gestures that are less often accompanied by these characteristics (S. Roberts & A. I. Roberts, 2018).

Final issue of importance for establishing which communication characteristics can influence preference for social affiliation is the positive versus negative communication style. Maestripieri and colleagues (1999, 2005) were amongst the first to recognize that affiliative as opposed to antagonistic gestural communication was of profound importance to the likelihood of social affiliation. Comparing three species of macaques with different social styles, they were able to show that more tolerant macaque species where the influence of kinship on social relationships was reduced displayed a larger repertoire of affiliative communication when compared to less tolerant macaque species where the influence of kinship was stronger. In more tolerant macaque species social interactions can easily disintegrate in response to exposure to aggression demanding higher investment in repair through affiliative communication. More recently, a study of a wild chimpanzee group showed that higher rates of affiliative gestural communication relative to antagonistic gestural communication predict a longer duration of time spent in proximity (S. G. B. Roberts & A. I. Roberts, 2016).

Chimpanzees, *Pan troglodytes*, are particularly good model species to illustrate how gestural communication affects primate social affiliation. Chimpanzees, live in large, fission-fusion social groups of up to 120 individuals, associating and affiliating in subgroups of varying duration and composition (Goodall, 1986). Due to fission-fusion nature of social system, mother and maternal kin are often unavailable as social partners demanding that chimpanzees form social bonds with unrelated group members (Goldberg & Wrangham, 1997; John C Mitani et al., 2002). The chimpanzees derive fitness benefits of social affiliation with unrelated group members in many different contexts such as grooming, coalitions, sharing meat, and hunting (Boesch, 1996; Foerster et al., 2015; Goldberg & Wrangham, 1997; K. E. Langergraber, Mitani, & Vigilant, 2007). Recent research has generated proofs that gestural communication of chimpanzees can influence preference for social affiliation. For example, recent studies have demonstrated the link between proximity measures (an index of sociality) and a number of different indices of communicative complexity such as intentionality (A. I. Roberts, 2018; A. I. Roberts & S. G. B. Roberts, 2018), repertoire size (A. I. Roberts, Chakrabarti, & Roberts, 2019) and multimodality (S. Roberts & A. I. Roberts, 2018). Whereas the role of gestures in maintaining proximity has been made clear, the role of gestural communication has received only limited research attention with regard to its influence on any other aspect of social affiliation in primates. However, in order to develop a deeper understanding of the factors driving social relationships, several measures of the social complexity are needed. As well as proximity, primates use visual attention to monitor conspecifics and also preferentially associate with others during grooming, feeding, resting and travel (Nishida, Zamma, Matsusaka, Inaba, & McGrew, 2010). All these different modes of interaction contribute to forming social relationships (Hinde, 1976). Studies which only focus on one behavioural aspect may therefore miss important features of social relationships.

Here we provide first systematic test of the hypothesis that gestural communication can influence preference of the recipient to form social relationships with the signaller. First, to test this hypothesis we use social network analysis to determine the link between communication and social bonding behaviours. Social network analysis represents individuals as nodes (e.g. individual A or B) and the social relationships (e.g. duration of time spent travelling, per hour spent within 10 m between AB dyad) as the edge or a ‘tie’. Social network analysis examines how increase in the value of one variable produced by a dyad is associated with the increase or decrease in the value of another variable by a dyad (Croft, James, & Krause, 2010). For instance, if individual A directs tactile gestures and grooming at the individual B, then social network analysis can determine whether the rate of tactile gestures directed by individual A at the individual B is associated with increase or decrease in the duration of time individual A spends grooming individual B.

It has long been established that animal signals evolve to function effectively in a particular social environment. The variation in the conflict of interest between signaler and the recipient is reflected in how effectively different types of gesturing can influence social bonding. When social bonds are weaker, and the conflict of interest between signaler and the recipient is high, chimpanzees use more effective, and intense gestures (e.g. tactile, auditory). In contrast, when the social bonds are stronger and the conflict of interest is low, chimpanzees use less effective and less intense gestures (e.g. visual). However, intensity of gestures affects the specificity with which signaller can convey their goal of the interaction. Low intensity, visual gestures are less specific to context than more intense tactile or auditory gestures. Lack of specificity may affect the recipient’s ability to effectively decode signal meaning and thus respond adaptively. One would therefore predict that communicative complexity would more likely co-occur with less intense gestures (e.g. visual) than with more intense gestures to enable the signaller to increase specificity of the gestures so that the recipient can decode signal meaning. Thus, one may predict that low intensity, complex signals will co-occur with indices of stronger social bonding (e.g. mutual grooming, joint travel), whereas higher intensity, non-complex signals will co-occur with indices of weaker social bonding (e.g. unidirectional grooming). In addition, it seems reasonable to assume that style of the gestural communication will affect the efficiency with which individuals can form social relationships. For instance, more bonded partners may exhibit a higher rate of gestures used in affiliation contexts as compared with antagonistic contexts because signaller gestures affiliatively to increase duration of social bonding. However, to date, this association between social relationships and complexity of gestural communication has not been examined systematically.

Second, we determine whether displacement behaviours such as self-scratching that help primates cope with anxiety (Aureli & Schaik, 1991; Baker & Aureli, 1997; Castles & Whiten, 1998; Diezinger & Anderson, 1986; Schino, Perretta, Taglioni, Monaco, & Troisi, 1996) are associated with gestural communication that facilitates social bonding. Studies of long-tailed macaques (Aureli & Schaik, 1991), olive baboons (Castles & Whiten, 1998) and chimpanzees (Baker & Aureli, 1997) have all used scratching to reliably estimate primate’s level of anxiety. Chimpanzees may experience increased anxiety in situations of social uncertainty linked to presence of unpredictable social partners with whom social interactions are infrequent or dominance relationships have been unresolved (Aureli, 1997; Schino, Scucchi, Maestripieri, & Turillazzi, 1988). In this case, one may predict that communication aimed at these social partners may be associated with higher rate of signaller’s displacement activity. Furthermore, chimpanzees may experience reduced anxiety in response to communication that has a rewarding property. For instance, vocal communication can prompt a surge in the production of social neurohormones in the recipient, such as a surge of oxytocin that relieves anxiety (Feldman, Gordon, & Zagoory-Sharon, 2011). In this case, it would be predicted that communication associated with weaker social bonds would be associated with reduced rate of displacement activity in the recipient.

## Methods

Twelve adult East African chimpanzees (*Pan troglodytes schweinfurthii*) (6 males and 6 females) were observed for a mean (SD) duration of 751.5 (250.9) min per each focal individual (Table 1). The sample included almost all of the uninjured adult males in the community and to match the sample size of the males, a similar number of uninjured females was chosen as focal individuals. The subjects were chosen on the basis that they were lacking physical injury, because the injuries such as deformation of the hand by the snare can influence the type of communication used. Moreover, the sample of males and females was matched as closely as possible in terms of rank categories, including three low ranking and three high ranking males and females).

**Table 1:** Focal ID, sex, year of birth, kinship and reproductive status of the 12 focal subjects included in the study.

| **Focal subject ID** | **Sex** | **Age** | **Female reproductive status** | **Kinship** | **Total observation duration (minutes)** |
| --- | --- | --- | --- | --- | --- |
| KU | Female | 29 | Pregnant | KT (son) | 910 |
| KW | Female | 27 | Nursing |  | 510 |
| ML | Female | 33 | Cycling |  | 1118 |
| NB<sup>b</sup> | Female | 46 | Cycling | MS (son) | 500 |
| RH | Female | 43 | Nursing | NK (son) | 1038 |
| ZM | Female | 40 | Cycling |  | 710 |
| BB | Male | 21 | - |  | 516 |
| HW | Male | 15 | - |  | 1030 |
| KT | Male | 15 | - | KU (mother) | 1026 |
| MS | Male | 17 | - | NB (mother) | 524 |
| NK<sup>a</sup> | Male | 26 | - | RH (mother) | 582 |
| SQ | Male | 17 | - |  | 554 |

Notes. <sup>a</sup> Alpha male <sup>b</sup> Alpha female.

The chimpanzees lived in the tropical forest, Sonso community at the Budongo Conservation Field Station in Uganda, East Africa. The data was collected during 18 minute focal animal follows, consisting of 9 scans at 2 minute intervals, recording the activity of the focal individual (grooming, travel, resting, feeding), the visual attention, activity and proximity of the nearest neighbor and the identity of all individuals present within 10 m (see Table 2 for definitions). Chimpanzees in the wild, especially when present in large parties, can change behavior and activity frequently. Thus, in order to reduce variability within samples, a focal follow of 18 minute duration was chosen. Gestures were recorded continuously using a digital video camera, verbally recording directly onto the footage the identity and the behavior of the signaler and recipient, along with the goal directedness in the behavior and the functional context.

**Table 2.**
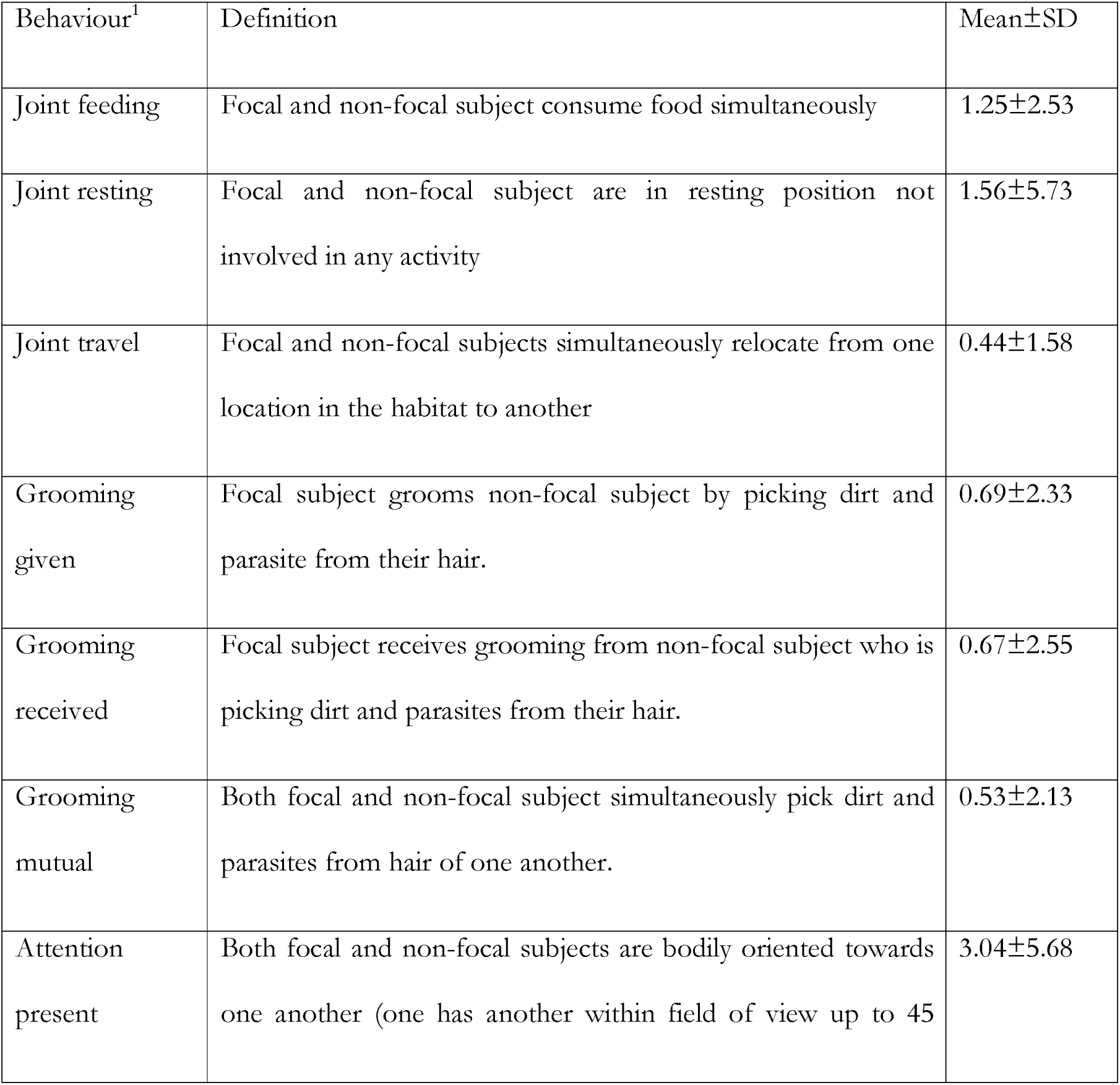
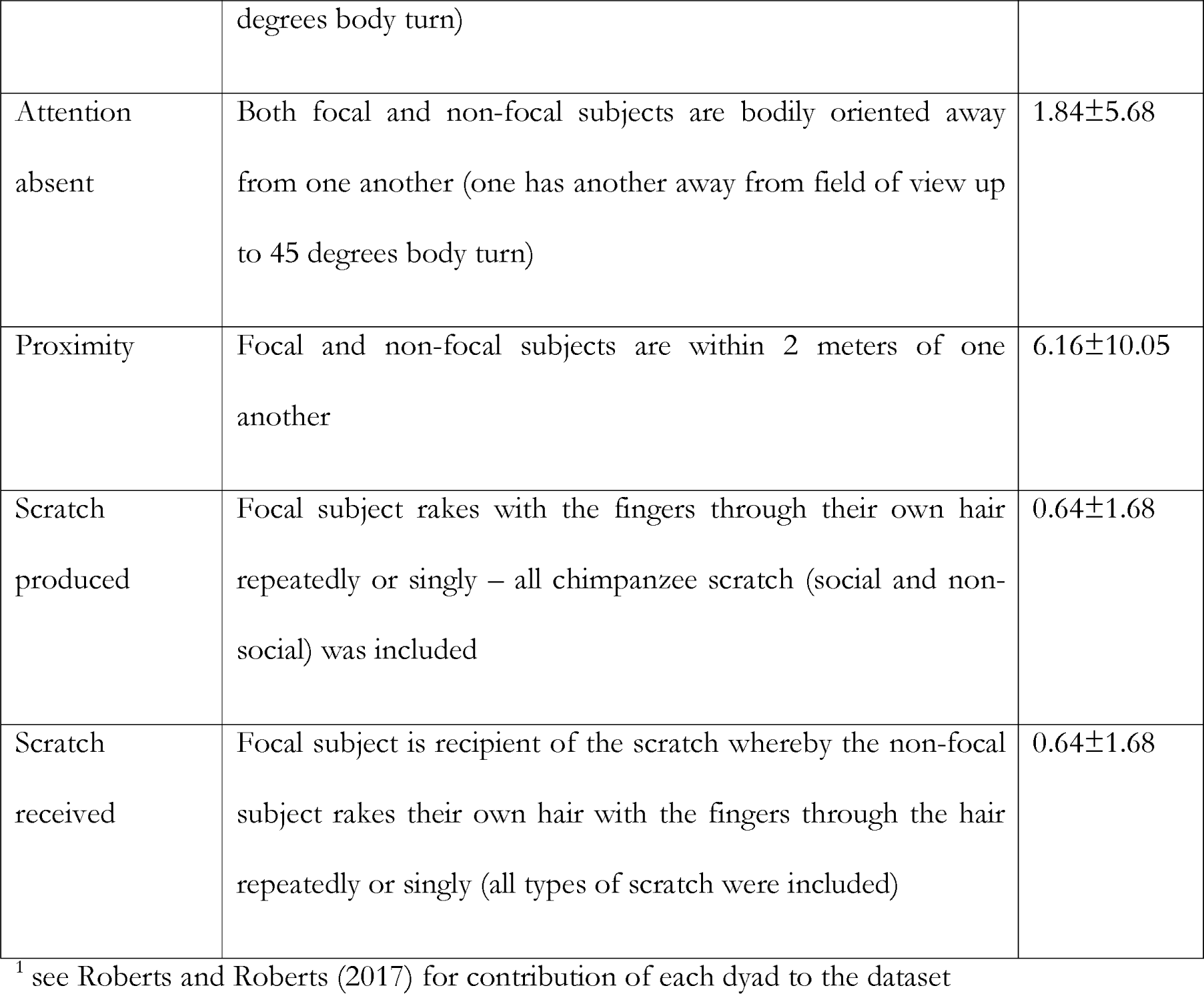
Definitions, means and standard deviations (SD) for social behaviours, based on 132 chimpanzee dyads. All social behaviours measured as durations (mins), per hour dyad spent within 10 m. For joint behaviours (e.g. feeding, resting, travelling) both dyad partners were engaged in the same behaviour. The behaviors are described in depth in (Nishida et al., 2010).

Using the video footage, the inventory identified gestures as acts of non-verbal behavior that were expressive movements of the limbs, head or the body posture that were intentional, communicative and mechanically ineffective. The detailed description of intentionality coding, along with the video clips of each gesture type, has been described previously (A. I. Roberts, Roberts, & Vick, 2014; A. I. Roberts, Vick, & Buchanan-Smith, 2012; A.I. Roberts et al., 2012). Validation of the coding procedure was established by a second coder and reliability (Kappa coefficient, K) was good for function (K = 0.70), modality of gesturing (K = 0.946) and intentionality (K = 0.74) (S. G. B. Roberts & A. I. Roberts, 2016). The behavioural measures (e.g. joint feeding, mutual grooming) were calculated as the duration of time pairs of chimpanzees engaged in these behaviors, per hour they spent within 10 m. The definitions as well as mean ± SD for all variables entered into the models can be found in Tables 2 and 3. Dyads were classified according to maternal kinship, sex, age class and reproductive state. All communication measures were calculated as a rate per hour chimpanzees spent within 10 m (see Supplementary Table 1 for definitions).

**Table 3.**
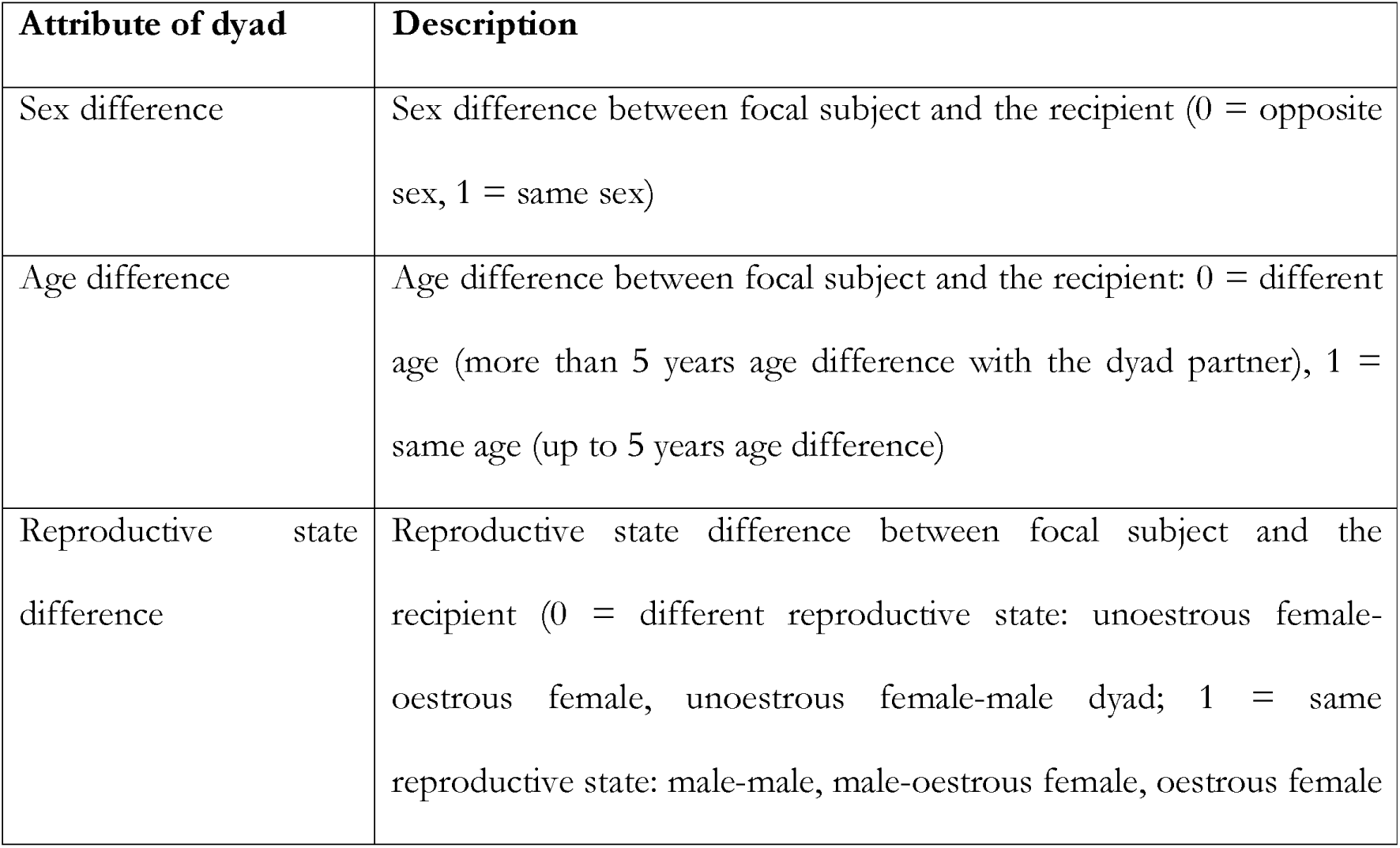
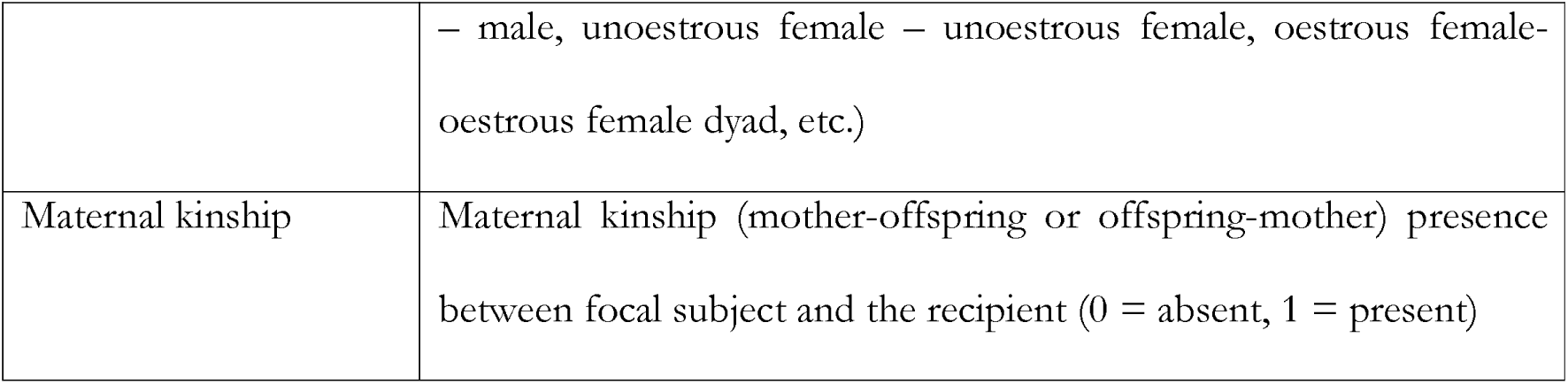
Attributes used to classify dyads

The rates of communication and durations of behavior were used to construct weighted, directed networks. The data was transformed and analyzed using UCINET 6 for Windows (Borgatti, Everett, & Johnson, 2013). The data used in social network matrices is dependent and therefore analysis using randomization (or permutation) have been developed, where the observed value is compared against a distribution of values generated by a large number of random permutations of the networks. The *p* value is calculated by calculating the proportion of random permutations in which a value as large (or as small) as the one observed is present. The relationships between networks were analyzed using Multiple Regression Quadratic Assignment Procedure (MRQAP) (Borgatti et al., 2013). This regression technique resembles a standard regression model as it enables us to examine the association between a number of predictor variables (e.g. rates of gestural communication between dyads, control variables relating to sex and age differences of dyads) and a single dependent variable (e.g. duration of proximity between dyads). We used Double Dekker Semi Partialling MRQAP regression as it is more robust against the effects of network autocorrelation and skewness in the dataset than other forms of network regression (Dekker, Krackhardt, & Snijders, 2007). For this analysis, 2,000 permutations were used following the settings of UCINET.

Running large numbers of models risks inflating significance levels (Field, 2013). To address this issue, following the methods used by Pearce et al. (2017), we examined whether the overall distribution of significant results varied across the different domains of sociality (joint activity, grooming, visual attention, proximity and scratch) and behavioural categories. These data are not subject to potential problems arising from multiple comparisons effects because the p value is used to categorise the results in each category as statistically significant or not, and one chi-square test is then carried out on the overall distribution of results (Pearce et al., 2017). Thus, we used a Chi-square test to examine the overall pattern of association between the behavioral indices and gestural communication. Following the same method as Pearce et al. (2017), this allowed us to examine whether the significant associations between gestural communication and behavior showed a distinct pattern, with certain types of gestural communication associated with specific behavioral indices more commonly than would be expected, as compared to a random distribution. This method is used as an alternative to Bonferroni correction when the multiple comparisons prevent the Bonferroni correction being used (M. J. Silk, Jackson, Croft, Colhoun, & Bearhop, 2015). We created contingency table whereby we included percentage of significant associations (p < 0.05) with each variable within each domain of social complexity (Supplementary Table 2). The results showed that the distribution of significant associations was non-random (χ2 = 364.90, df =60, p < 0.001). This demonstrated that the significant results were not randomly distributed across the different domains of sociality, suggesting that different types of gestural communication are differentially associated with the types of sociality. In all tests, two-tailed probabilities were used, with p set at < 0.05. Supplementary Information 2 gives a summary of these findings, with the full models provided in Supplementary Information 1 (Tables S3 – S19).

## Results

Based on our 12 focal subjects, we studied directed patterns of social behaviour in pairs of chimpanzees, giving 132 unique dyadic relationships (e.g. the duration of grooming given by BB to HW, and the duration of grooming given by HW to BB). There was a great deal of variation in the rate of social behaviours amongst the dyads and across the different behaviours, as indicated by the fact that for all of the behaviours the standard deviation was larger than the mean (Table 2). Per hour spent within 10 m, the mean ± SD number of minutes dyads spent in social behaviours ranged between 0.44 ±1.58 spent in joint travel and 6.16 ± 10.05 spent in close proximity (within 2 m). Dyads scratched at a mean rate ± SD of 0.64 ± 1.68 instances of scratching, per hour spent within 10 m. Similarly for the communication indices, there was a large amount of variation between dyads, with lowest mean ± SD rate of behaviour for gestures accompanied by facial expression (0.09 ± 0.52) and the highest mean ± SD rate of events (3.18 ± 7.82) – Supplementary Table 2.

### Association between the duration of social behaviour and gestural communication categorized according to structure

In the next set of analyses, we used MRQAP network regression analyses to examine how the duration of social behaviours (the dependent variable in all the models) was associated with the rate of different types of gestural communication and demography (see Figures 1 and 2 for visual representation of the networks).

**Figure 1.**
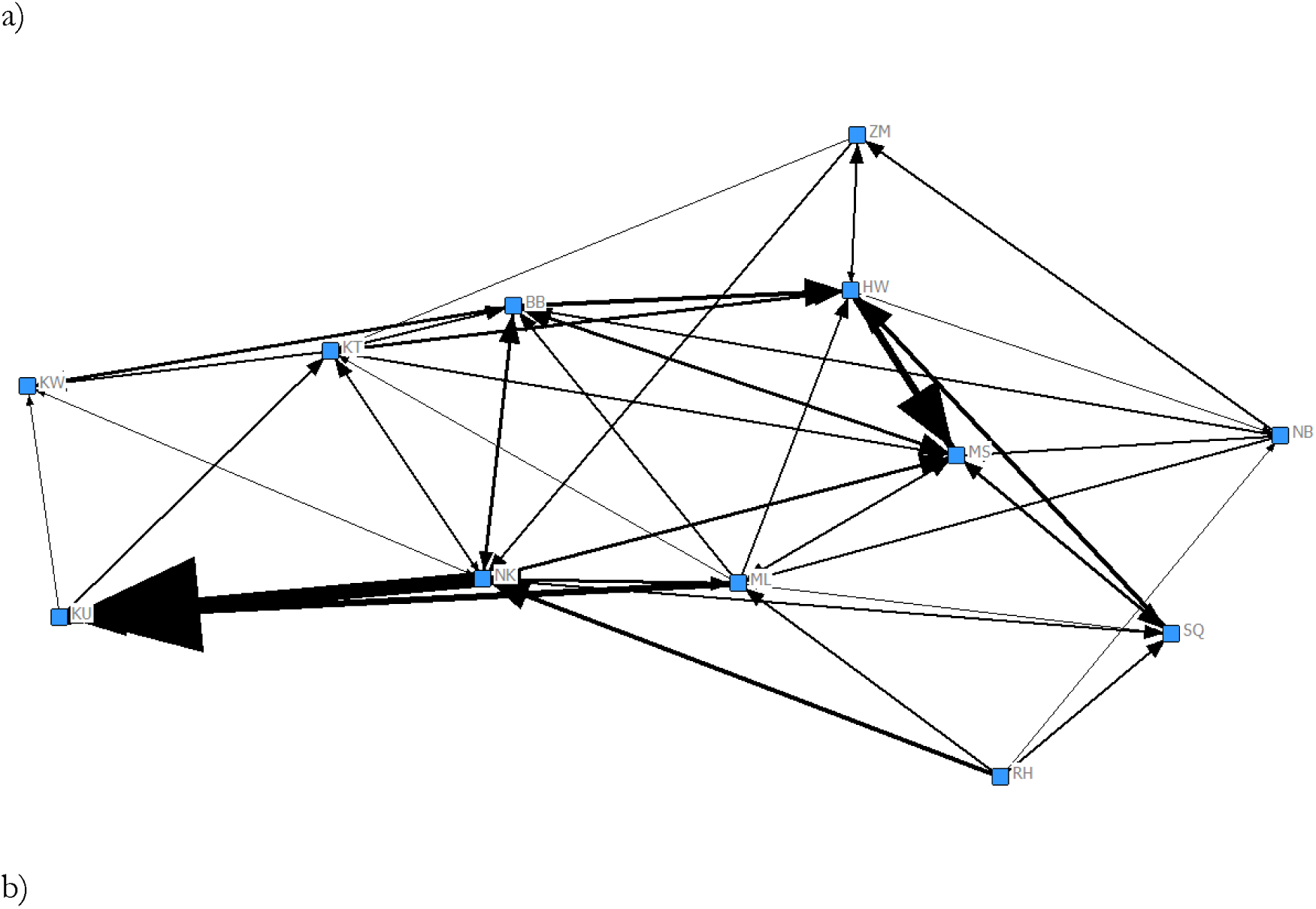
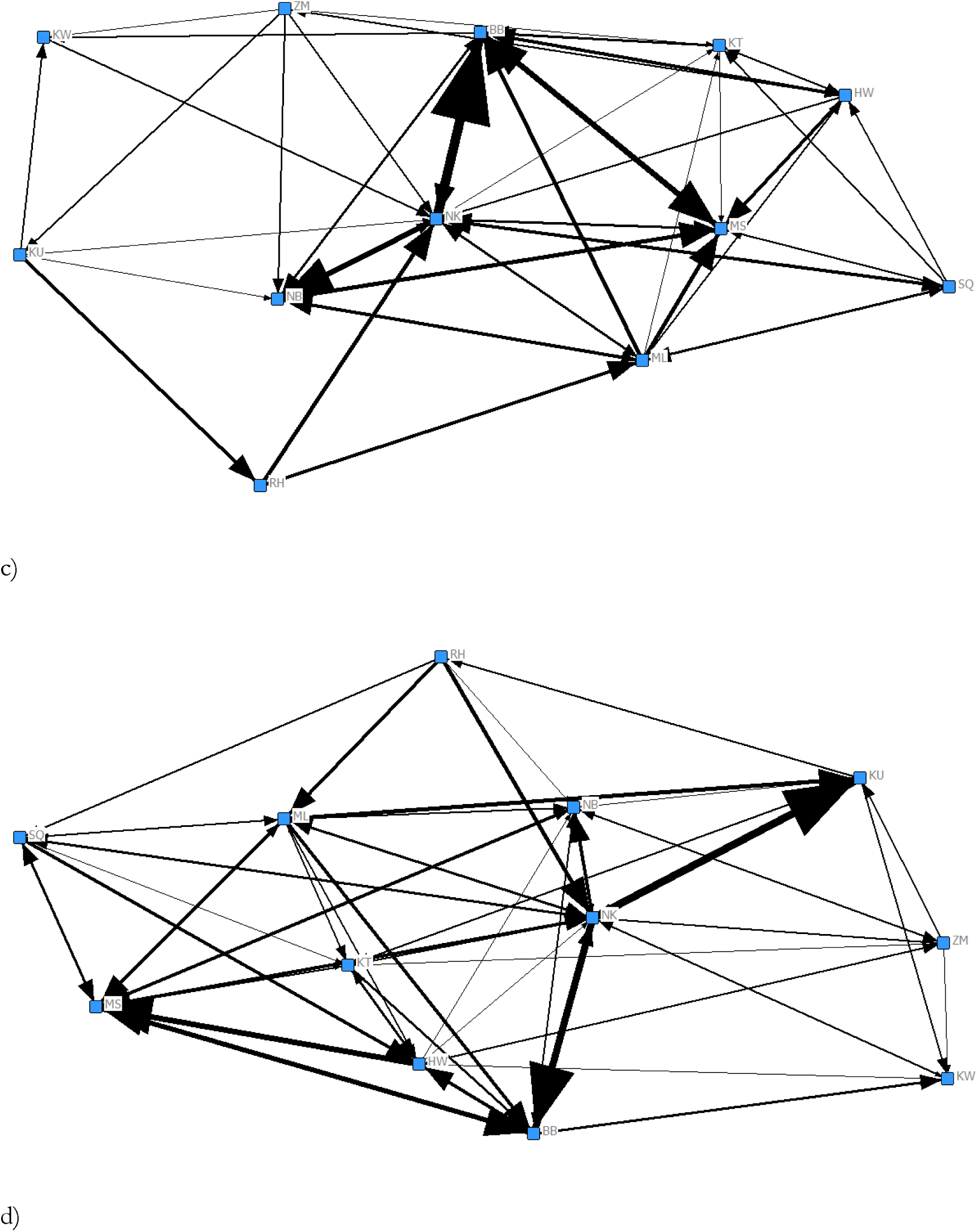
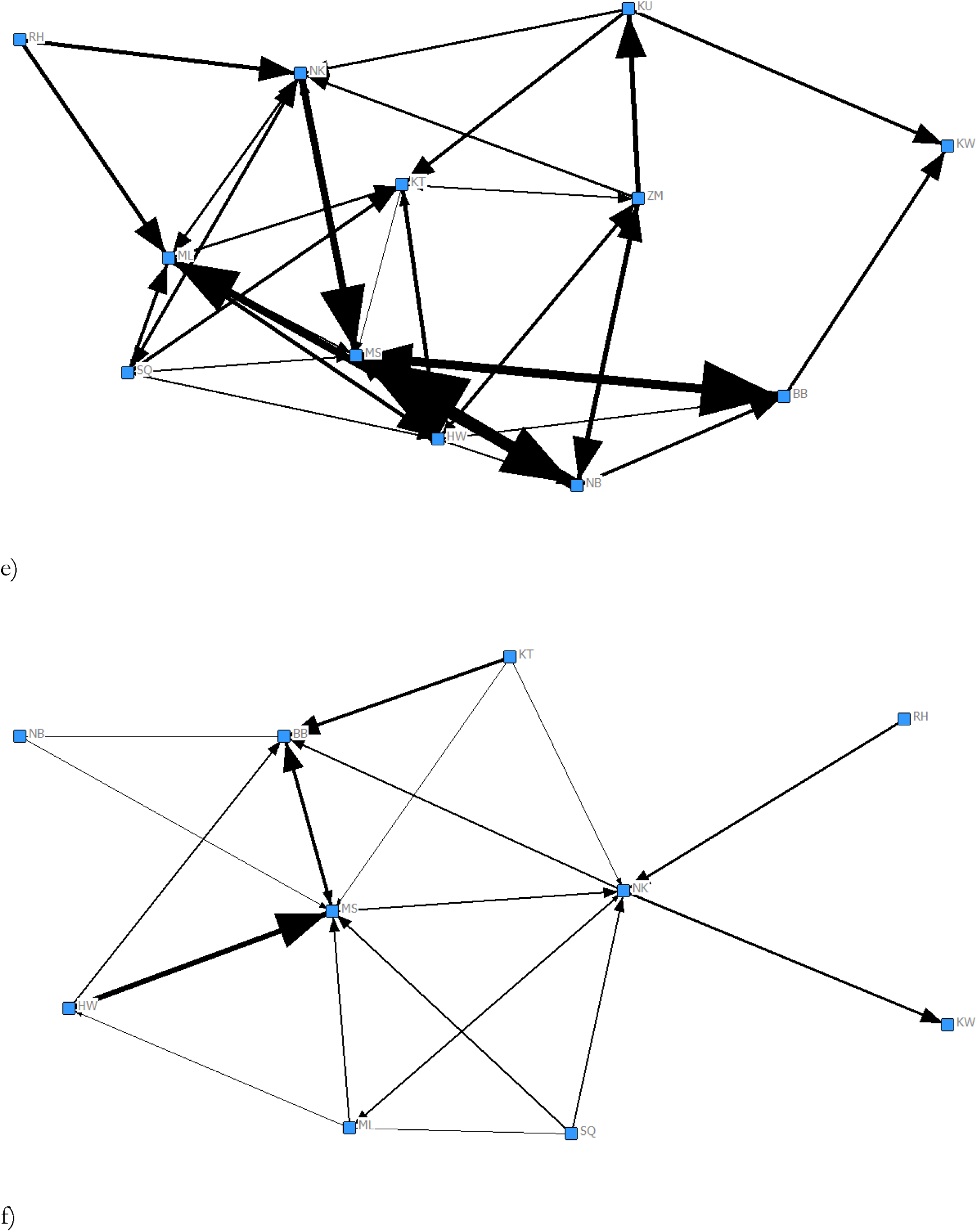
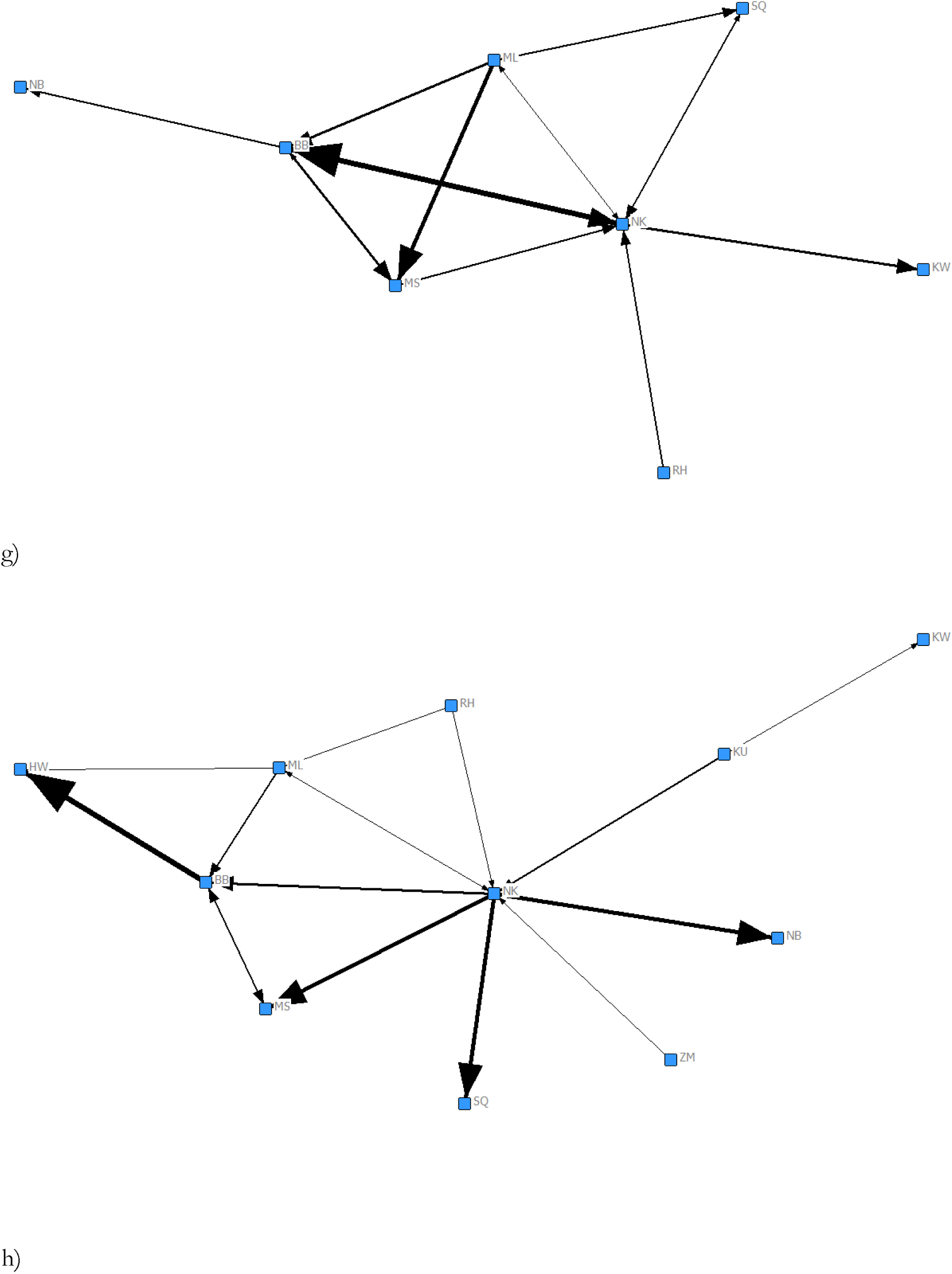
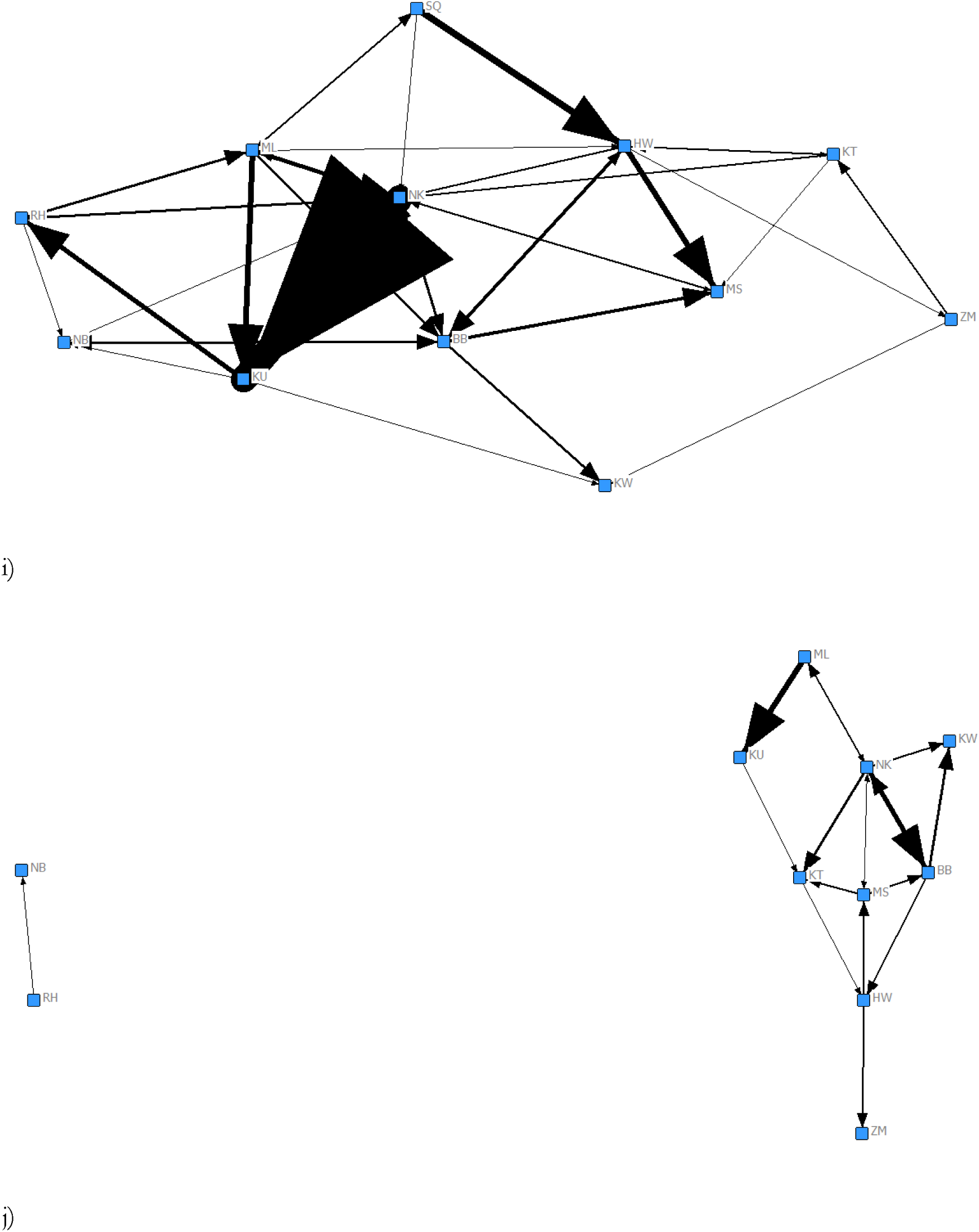
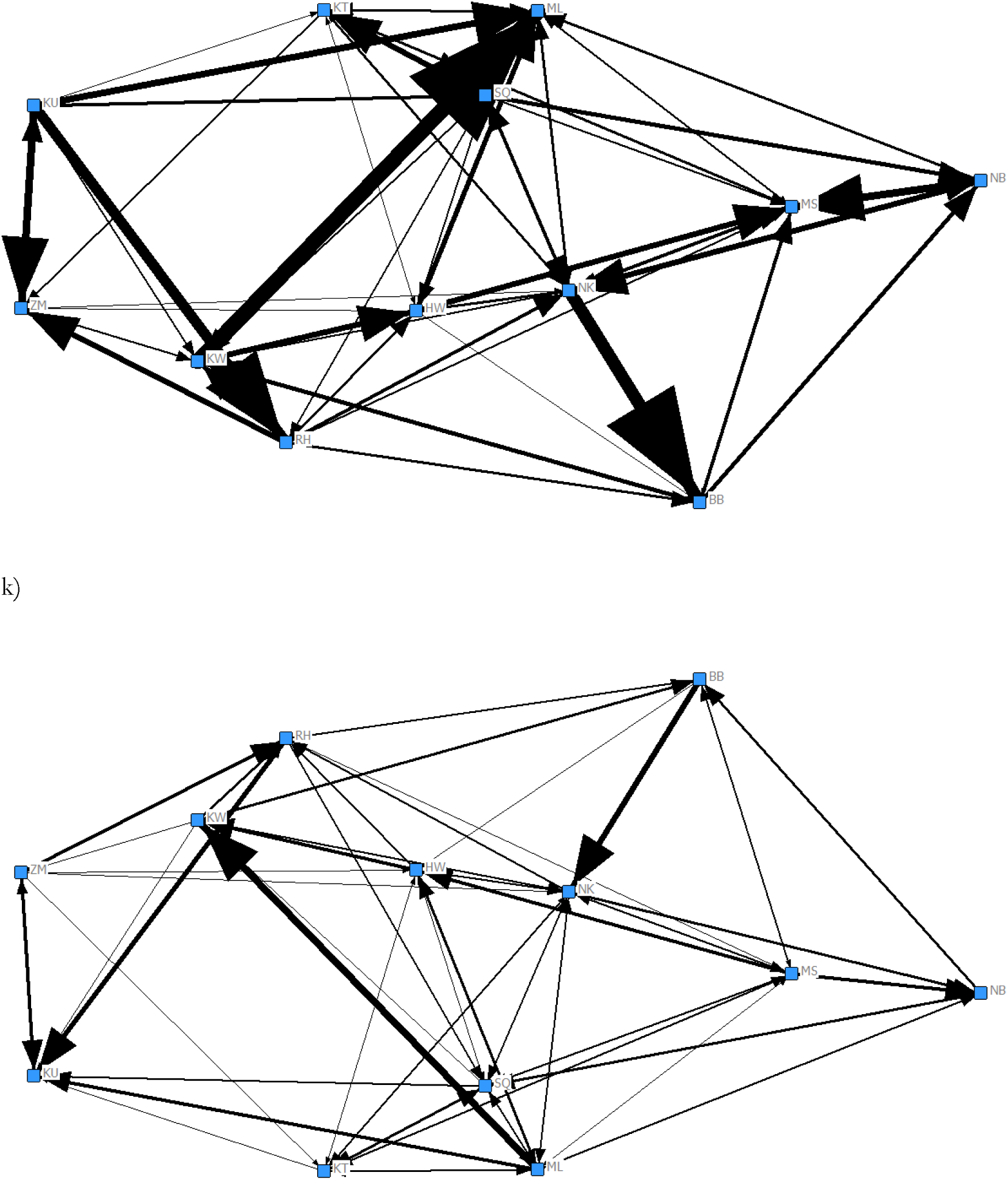
Social networks: a) attention absent, b) attention present, c) proximity, d) joint feeding, e) grooming given, f) grooming mutual, g) grooming received, h) joint resting, i) joint travel, j) scratch produced, k) scratch received

**Figure 2.**
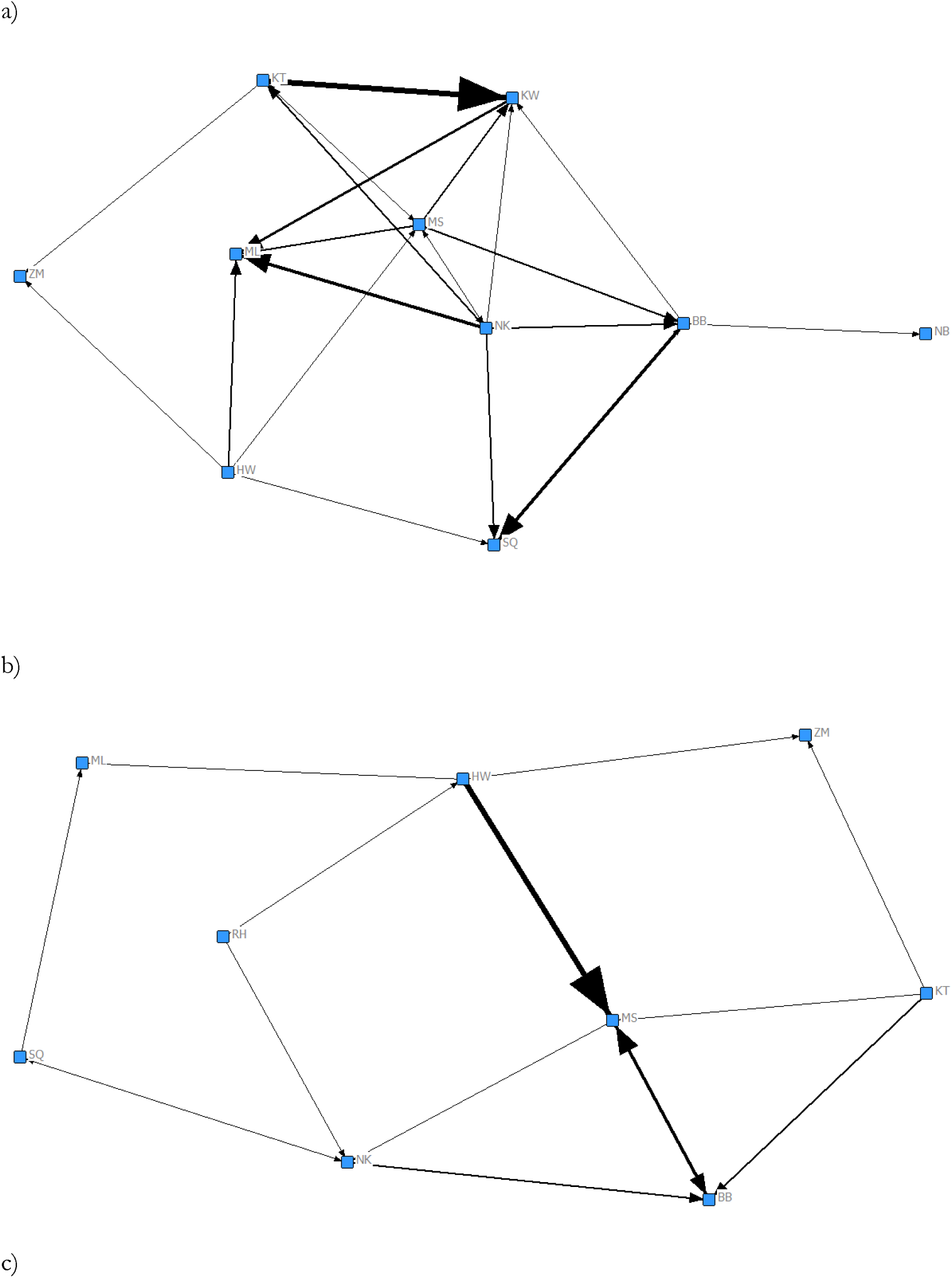
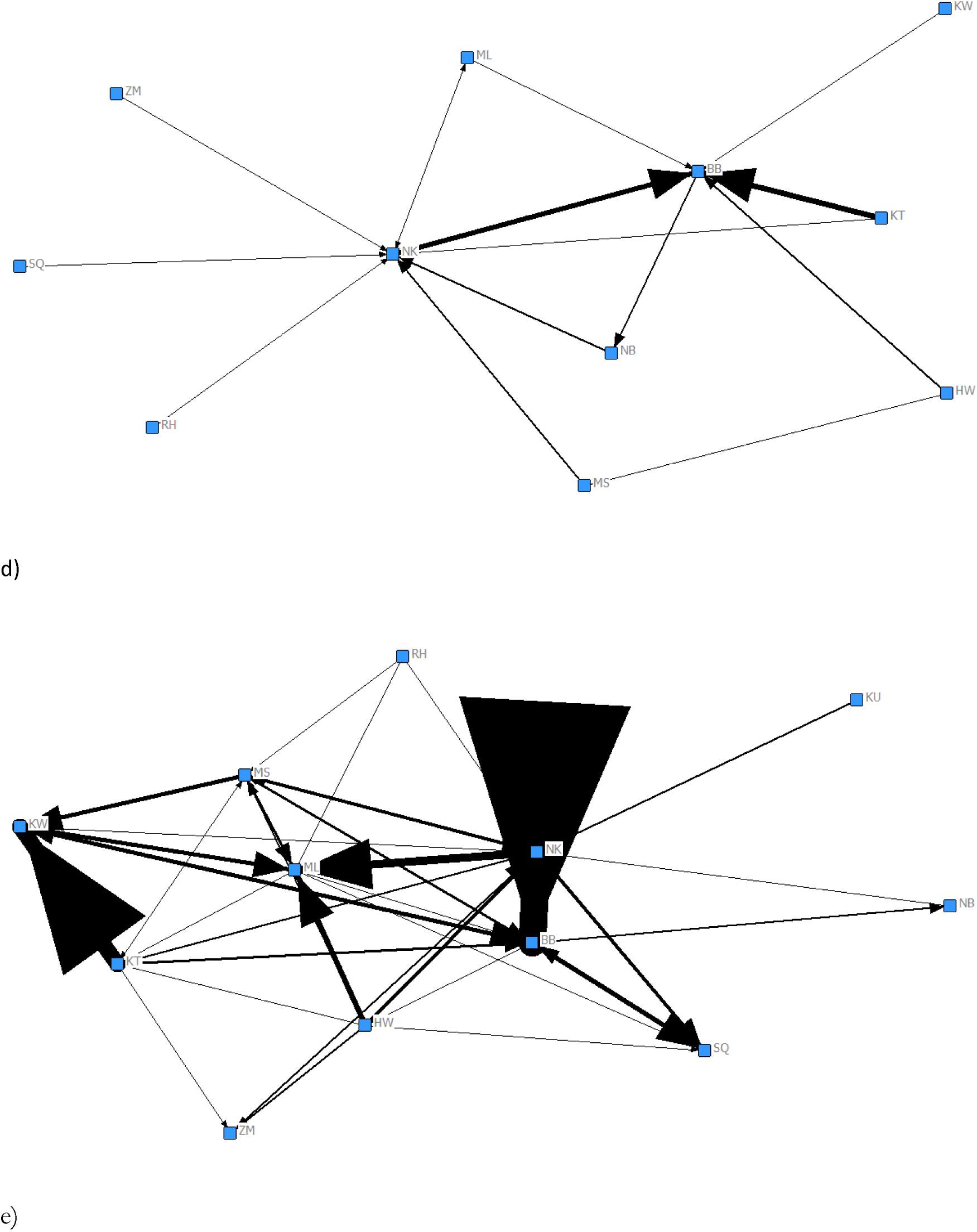
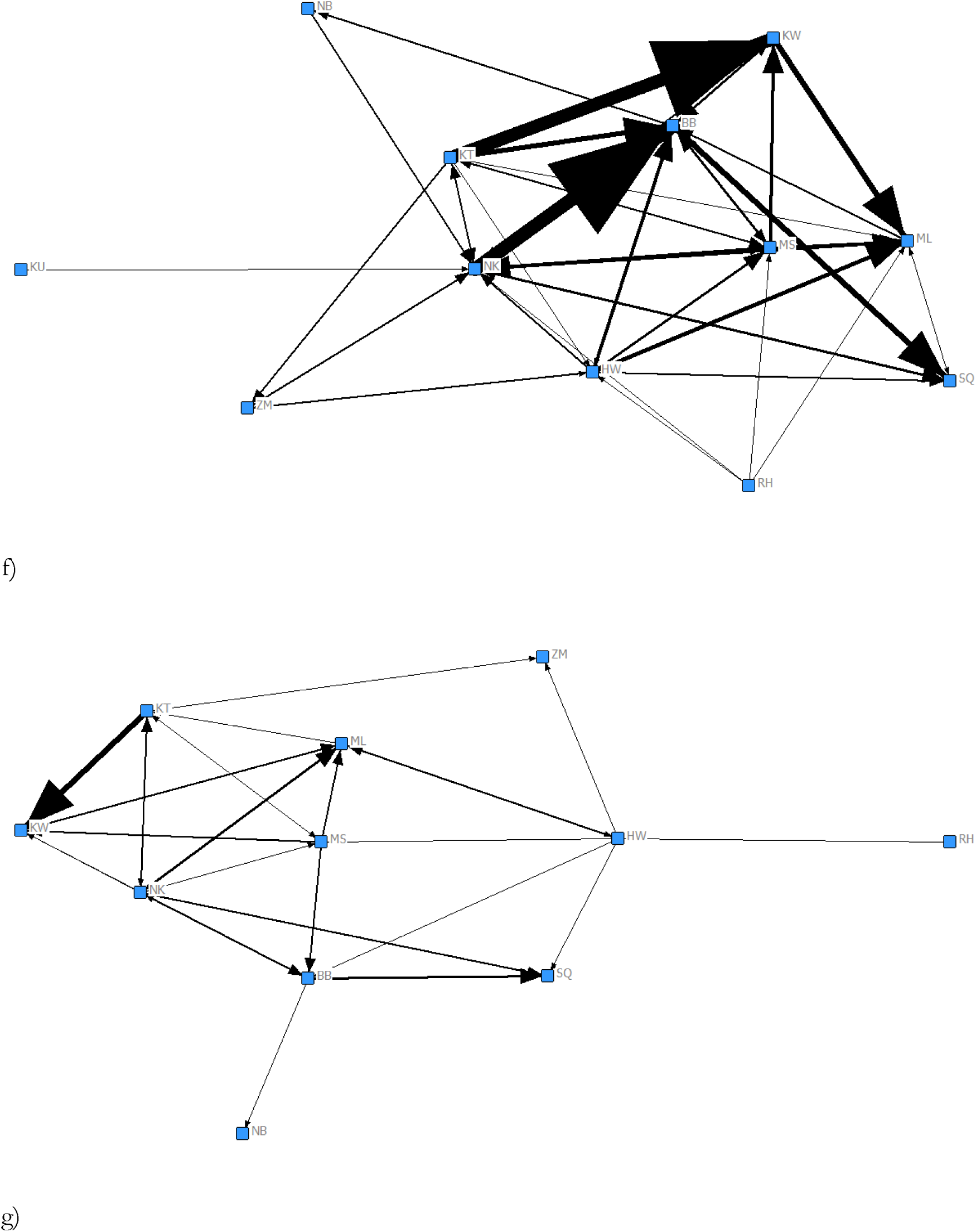
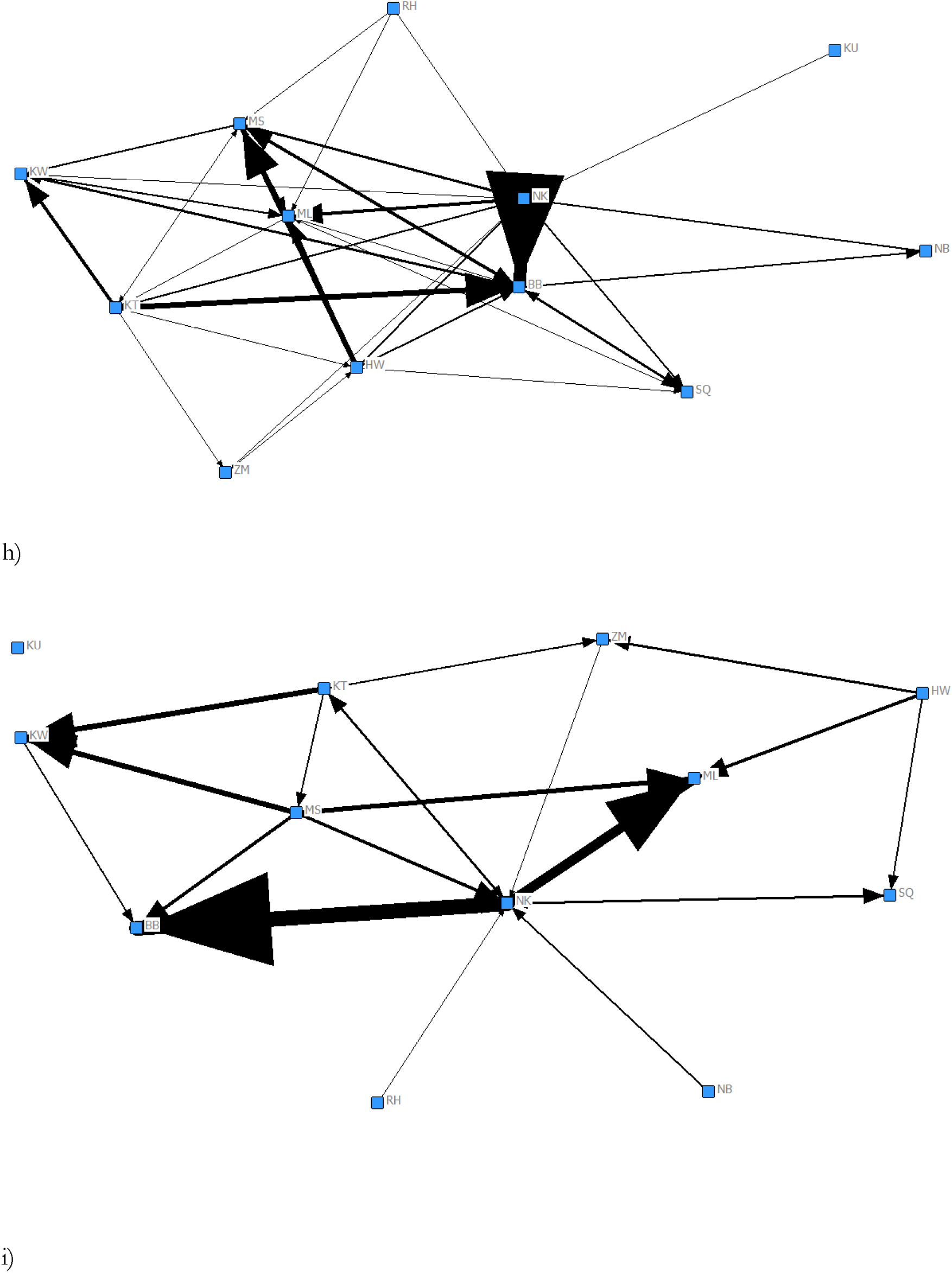
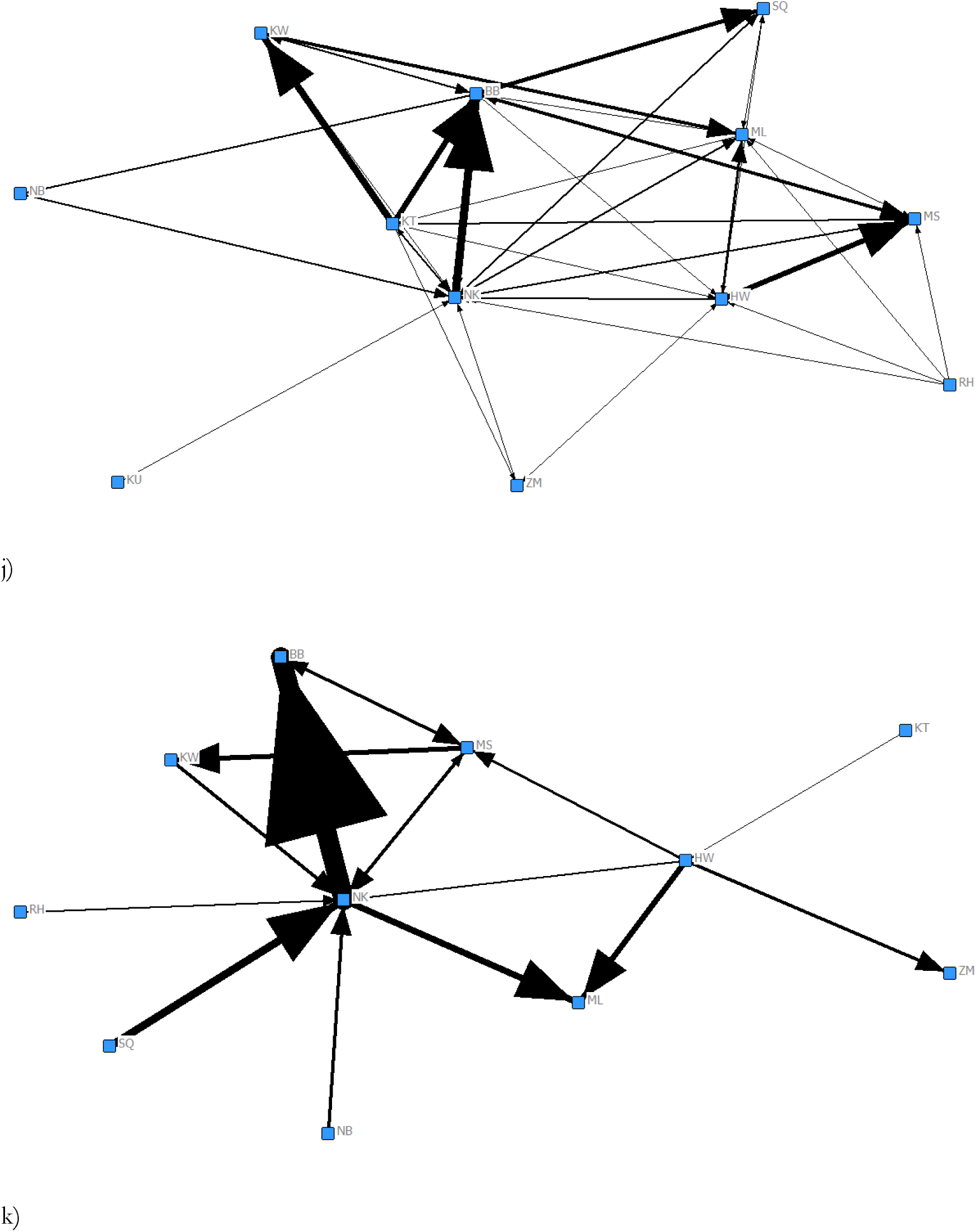
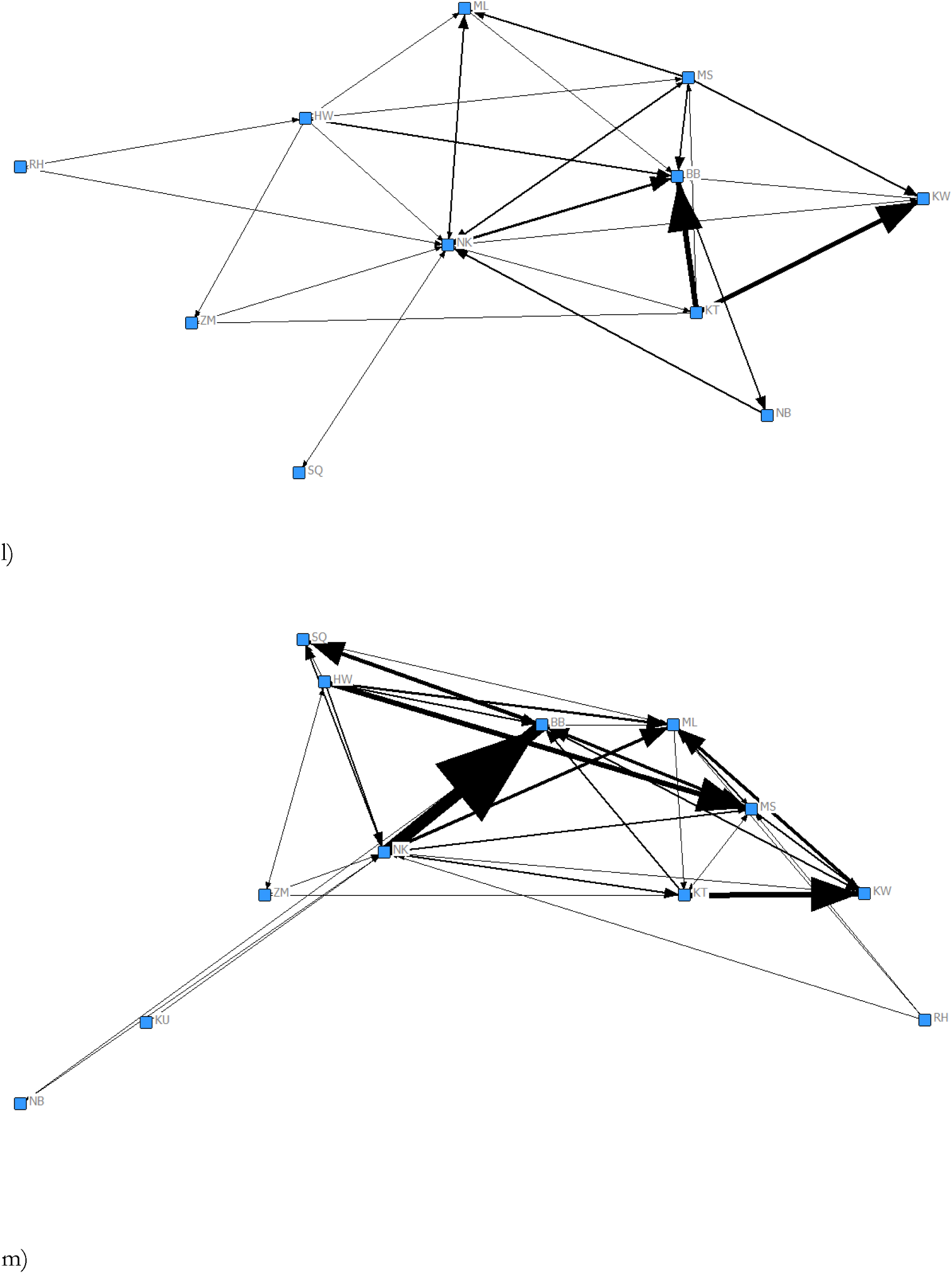
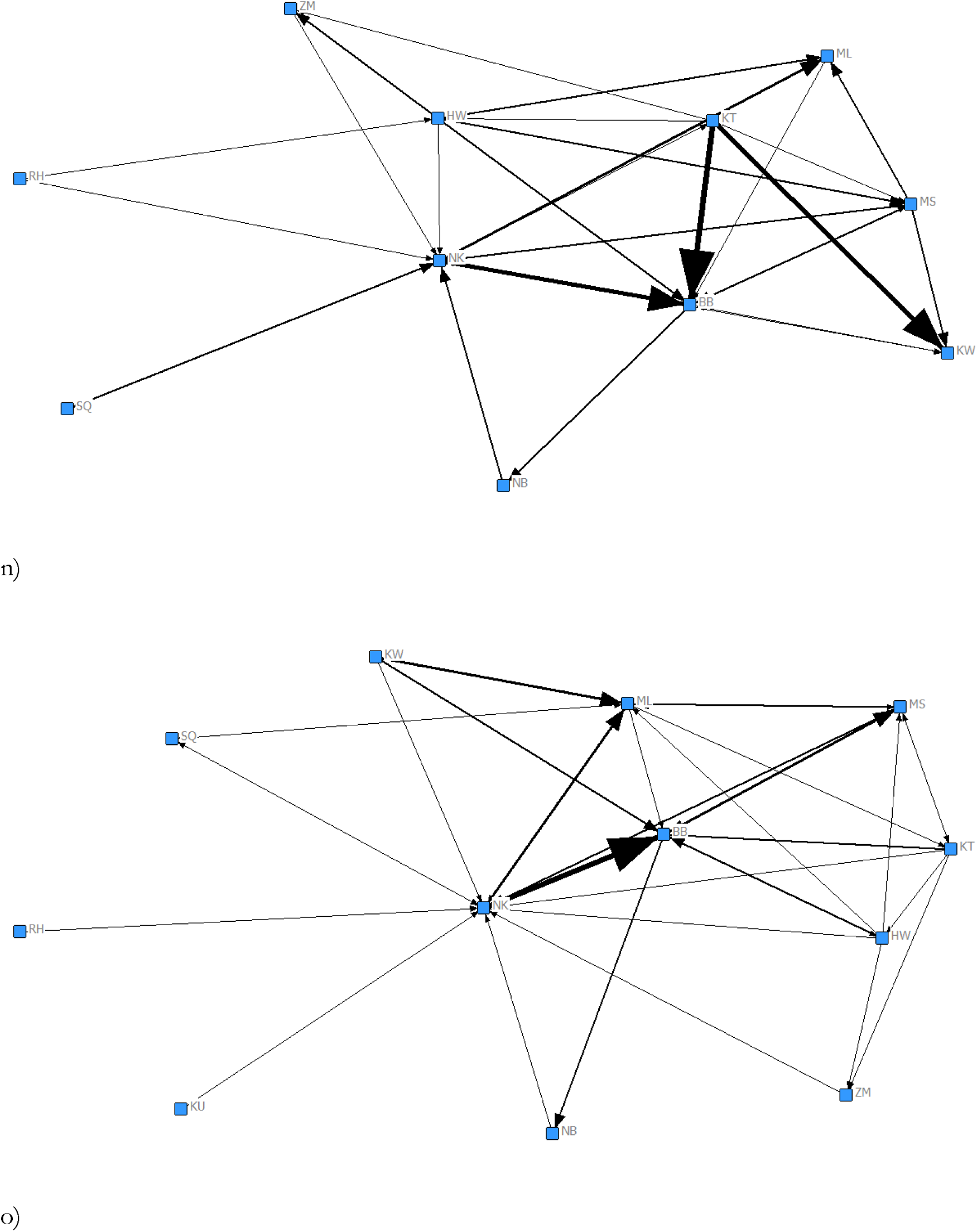
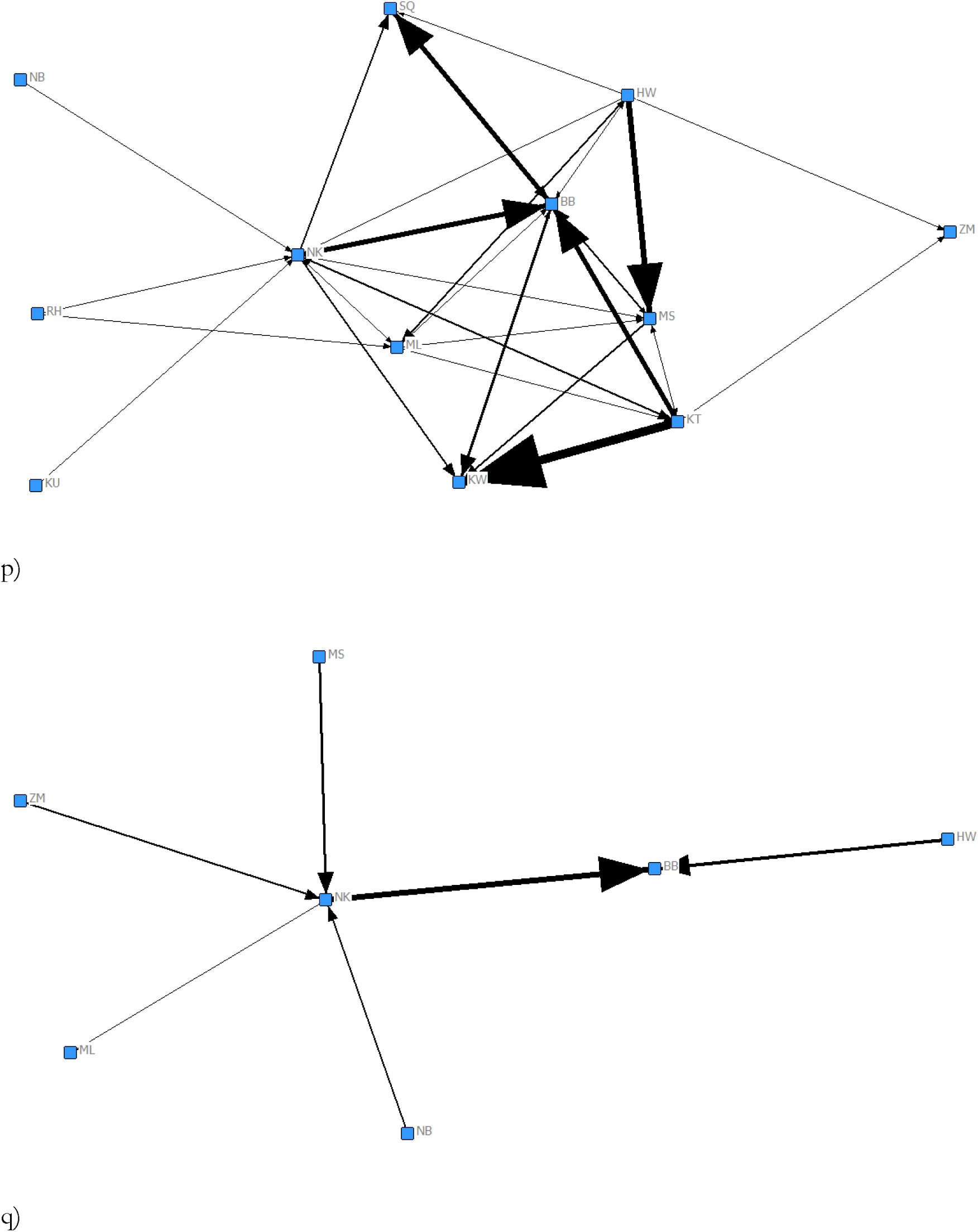
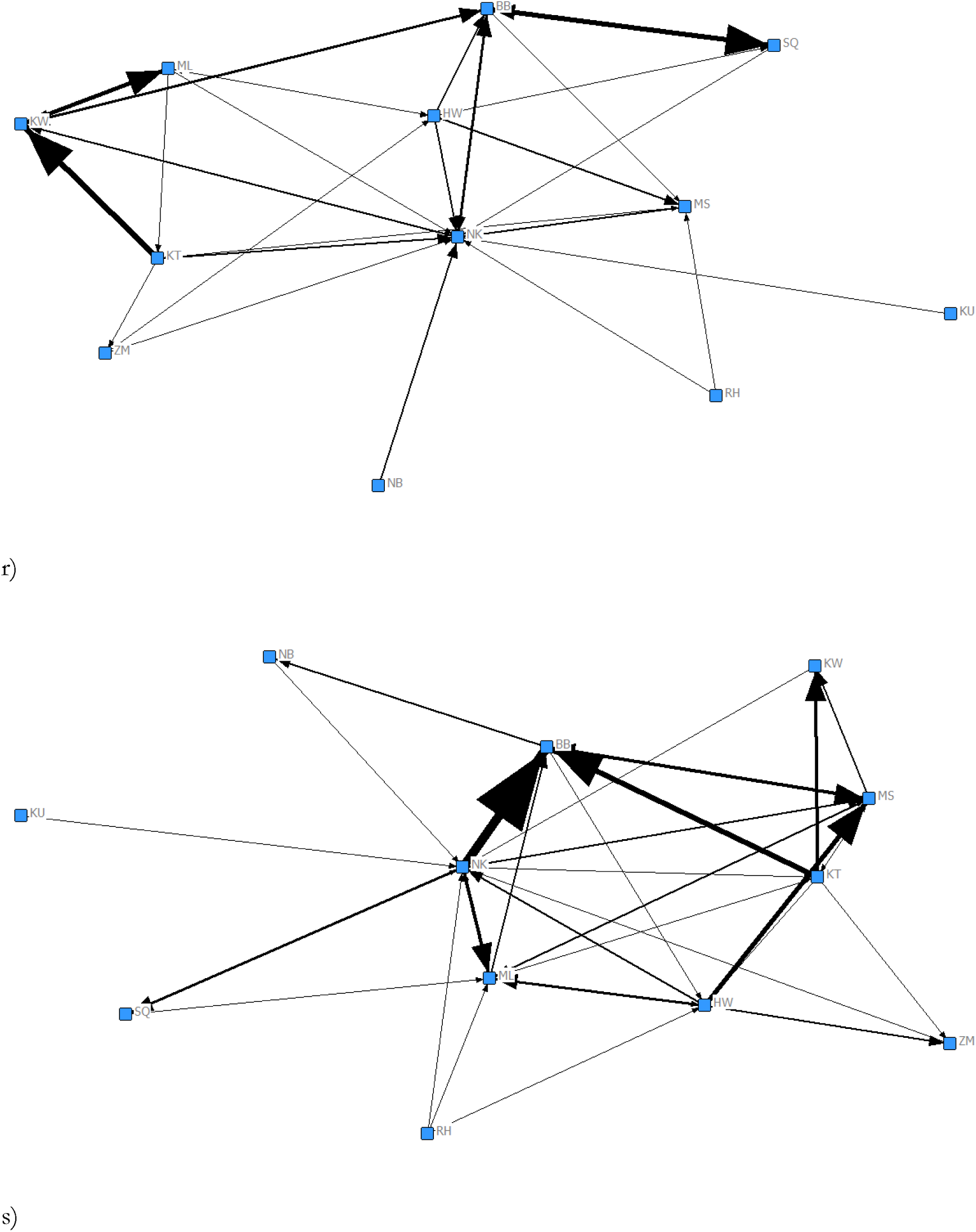
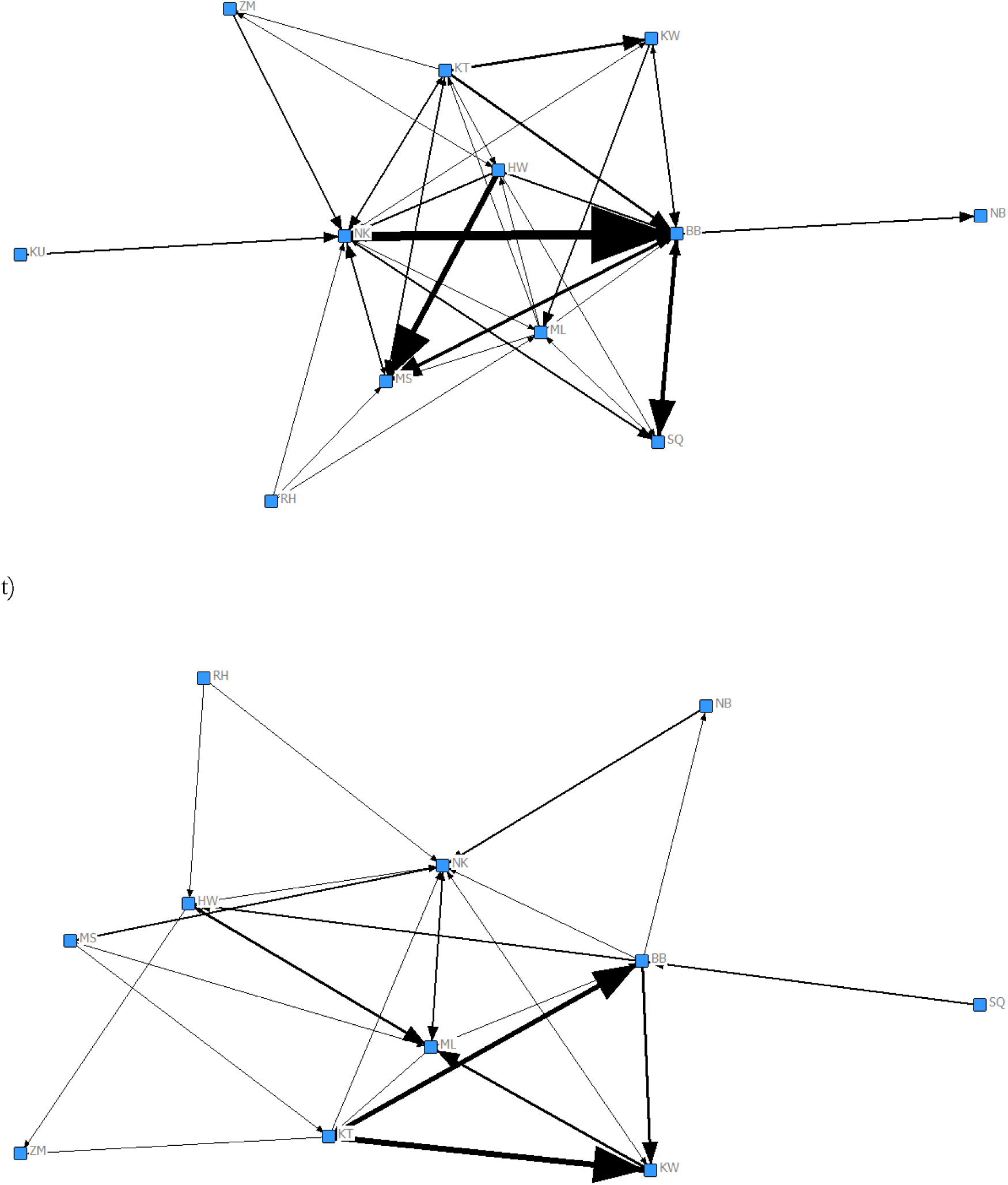
Social networks of gestures categorised according to Modality: a) auditory long range, b) auditory short range, c) tactile, d) visual; e) Repertoire size; Object use: f) objects, g) no objects; Combined gestures: h) combined, i) single (non-combined); Indicative: j) manual indicative, k) manual non-indicative; Bodily and manual: l) bodily, m) manual; Attention: n) mutual attention present, o) mutual attention absent; Multimodal: p) multimodal (facial expression), q) multimodal (vocal), r) unimodal; Homogeneity: s) homogenous, t) heterogeneous

#### Joint feeding

Same sex dyads (β = 0.31, *p* = 0.001), kin (β = 0.29, *p* = 0.010) and same reproductive status (β = 0.22, *p* = 0.040) dyads spent a significantly longer amount of time engaged in joint feeding, as compared to different sex dyads, unrelated dyads or different reproductive status dyads. Dyads who spent a longer amount of time jointly feeding had a significantly higher rate of auditory short-range gestures (β = 0.21, *p* = 0.040) and significantly lower rate of tactile gestural communication (β = −0.22, *p* = 0.004).

#### Joint resting

Same age dyads, as compared to different age dyads, spent a significantly longer amount of time engaged in joint resting (β = 0.29, *p* = 0.003). There was a positive association between the duration of time spent in joint resting and the rate of auditory short-range (β = 0.13, *p* = 0.040) and close proximity (β = 0.12, *p* = 0.046) gestures.

#### Joint travel

Same-age dyads, as compared to different-age dyads, spent a significantly longer amount of time jointly travelling (β = 0.28, *p* = 0.010). Further, the duration of joint travel was positively associated with a higher rate of gestural communication without objects (β = 0.47, *p* = 0.005), visual (β = 0.62, *p* = 0.010), unimodal (β = 0.22, *p* = 0.020) and multimodal with facial expression (β = 0.30, *p* = 0.020), combined (β = 0.41, *p* = 0.004), accompanied by mutual attention present (β = 0.37, *p* = 0.010), bodily (β = 0.50, *p* = 0.003), manual indicative (β = 0.42, *p* = 0.011), close (β = 0.27, *p* = 0.015) and far (β = 0.22, *p* = 0.013), non-repetitive (β = 0.55, *p* = 0.006), homogeneous (β = 0.43, *p* = 0.010), single (no sequence) (β = 0.22, *p* = 0.026), rapid (β = 0.20, *p* = 0.020) and persistence sequence (β = 0.14, *p* = 0.045), and accompanied by piloerection (β = 0.38, *p* = 0.006), The repertoire size of gesture types (β = 0.34, *p* = 0.010) and number of events exchanged by each dyad (β = 0.39, *p* = 0.003) was positively associated with the duration of joint travel. In contrast, a higher rate of auditory long-range gestures was associated with shorter durations of time spent in joint travel (β = −0.28, *p* = 0.010).

#### Grooming given

Same-age dyads spent significantly longer engaged in giving grooming (β = 0.21, *p* = 0.020). The duration of time spent giving grooming was positively associated with a higher rate of production of following categories of gestural communication: no object (β = 0.64, *p* = 0.001), auditory short-range (β = 0.73, *p* = 0.001), auditory long-range (β = 0.19, *p* = 0.010), tactile (β = 0.41, *p* = 0.001), unimodal (β = 0.75, *p* = 0.001), dyadic repertoire size (β = 0.26, *p* = 0.020), single (non-combined) (β = 0.64, *p* = 0.001), mutual attention absent (β = 0.78, *p* = 0.001), bodily (β = 0.22, *p* = 0.023), manual (β = 0.29, *p* = 0.016), events (β = 0.46, *p* = 0.002), manual indicative (β = 0.17, *p* = 0.036), manual non-indicative (β = 0.35, *p* = 0.014), close (β = 0.76, *p* = 0.001), repetitive (β = 0.60, *p* = 0.001), homogenous (β = 0.52, *p* = 0.004) and single (no sequence) (β = 0.66, *p* = 0.001). In contrast, the dyads who spent a shorter duration of time giving grooming communicated at a higher rate through the use of objects (β = −0.20, *p* = 0.010), visual gesture (β = −0.26, *p* = 0.010), multimodal (facial expression) (β = −0.23, *p* = 0.010), combined (β = −0.19, *p* = 0.020), mutual attention present (β = −0.11, *p* = 0.040) and far (β = −0.11, *p* = 0.012).

#### Grooming mutual

Same age dyad partners spent a longer duration of time mutually grooming (β = 0.20, *p* = 0.049). A longer duration of time spent mutually grooming was also associated with a higher rate of the following gesture types: no object (β = 0.64, *p* < 0.001), visual (β = 0.88, *p* < 0.001), unimodal (β = 0.27, *p* = 0.020), multimodal (facial expression) (β = 0.44, *p* = 0.002), dyadic repertoire size (β = 0.45, *p* = 0.003), combined (β = 0.40, *p* = 0.003), mutual attention present (β = 0.63, *p* = 0.001), bodily (β = 0.58, *p* = 0.001), events (β = 0.45, *p* = 0.003), manual indicative (β = 0.56, *p* = 0.001), non-repetitive (β = 0.72, *p* = 0.001), homogeneous (β = 0.60, *p* < 0.001), close (β = 0.38, *p* = 0.004), far (β = 0.22, *p* = 0.018), single (no sequence) (β = 0.33, *p* = 0.012), rapid (β = 0.16, *p* = 0.041) and persistence sequence (β = 0.16, *p* = 0.030) and piloerection (β = 0.42, *p* = 0.004).. In contrast, a higher rate of gestures object (β = −0.15, *p* = 0.020), auditory long range (β = −0.51, *p* < 0.001), repetitive (β = −0.18, *p* = 0.004) and heterogeneous (β = −0.15, *p* = 0.016) predicted a shorter duration of time spent mutually grooming.

#### Grooming received

Same-sex dyads, compared to different-sex dyads, spent a significantly longer amount of time receiving grooming (β = 0.20, *p* = 0.030). Individuals who received a longer duration of grooming from the dyad partners produced gestures with no object (β = 0.22, *p* = 0.040), visual (β = 0.50, *p* = 0.020), unimodal (β = 0.25, *p* = 0.040), bodily (β = 0.25, *p* = 0.036), manual indicative (β = 0.17, *p* = 0.046), non-repetitive (β = 0.20, *p* = 0.035) and homogeneous (β = 0.17, *p* = 0.048) at a higher rate. However, individuals who received a shorter duration of grooming from the dyad partner directed auditory long range (β = −0.27, *p* = 0.020) and tactile (β = −0.20, *p* < 0.001) gestures at them at a higher rate.

#### Attention present

Presence of mutual attention occurred between dyads of same-sex (β = 0.30, *p* = 0.010) and same reproductive status (β = 0.28, *p* = 0.040) at a higher rate. Individuals who spent longer duration of time mutually attending with the dyad partner displayed a significantly higher rate of gestures with no object (β = 0.69, *p* < 0.001), visual (β = 0.75, *p* < 0.001), auditory short-range (β = 0.24, *p* = 0.010), unimodal (β = 0.44, *p* < 0.001), multimodal (facial expression) (β = 0.31, *p* = 0.003), dyadic repertoire size (β = 0.42, *p* = 0.002), single (non-combined) (β = 0.29, *p* = 0.004), combined (β = 0.30, *p* = 0.002), mutual attention present (β = 0.56, *p* < 0.001), bodily (β = 0.54, *p* = 0.001), events (β = 0.47, *p* = 0.001), manual indicative (β = 0.54, *p* = 0.001), close (β = 0.50, *p* = 0.001), far (β = 0.13, *p* = 0.045), non-repetitive (β = 0.56, *p* = 0.004), homogeneous (β = 0.60, *p* = 0.001), single (no sequence) (β = 0.47, *p* = 0.001) and piloerection (β = 0.35, *p* = 0.005). In contrast, individuals who spent a shorter duration of time mutually attending with the dyad partner displayed a significantly higher rate of gestures with object (β = −0.18, *p* = 0.003), auditory long-range (β = −0.44, *p* < 0.001), multimodal (vocal) (β = −0.09, *p* < 0.049) and heterogeneous (β = −0.09, *p* = 0.046).

#### Attention absent

Same-aged dyad partners spent a longer duration of time mutually non-attending towards one another as compared to different age partners (β = 0.36, *p* = 0.001). The duration of time dyad partners spent mutually non-attending was positively associated with the rates of gestures with no object (β = 0.24, *p* = 0.028), auditory short-range (β = 0.29, *p* = 0.011), unimodal (β = 0.27, *p* = 0.024), single (non-combined) (β = 0.18, *p* = 0.045), mutual attention absent (β = 0.28, *p* = 0.015), events (β = 0.15, *p* = 0.049), close (β = 0.28, *p* = 0.021), single (no sequence) (β = 0.23, *p* = 0.026) and homogeneous (β = 0.17, *p* = 0.043).

#### Proximity

In comparison with different-age dyad partners, same-age dyad partners (β = 0.32, *p* < 0.001) spent a significantly longer duration of time in close proximity (within 2 m). There was a significant positive association between the duration of time spent in close proximity and a higher rate of a wide range of communication: no object (β = 0.56, *p* < 0.001), visual (β = 0.45, *p* = 0.010), auditory short-range (β = 0.34, *p* = 0.010), unimodal (β = 0.44, *p* = 0.001), multimodal (facial expression) (β = 0.14, *p* = 0.049), dyadic repertoire size (β = 0.28, *p* = 0.010), combined (β = 0.29, *p* = 0.010), mutual attention absent (β = 0.26, *p* = 0.010), mutual attention present (β = 0.30, *p* = 0.010), bodily (β = 0.40, *p* = 0.003), events (β = 0.37, *p* = 0.003), manual indicative (β = 0.37, *p* = 0.007), close (β = 0.48, *p* = 0.001), non-repetitive (β = 0.33, *p* = 0.012), homogeneous (β = 0.45, *p* < 0.001), single (no sequence) (β = 0.42, *p* < 0.001) and piloerection (β = 0.21, *p* = 0.027). In contrast, there was a significant negative association between the duration of time spent in close proximity and the rate of gestures using object (β = 0.17, *p* = 0.003), auditory long-range (β = −0.27, *p* = 0.010) and multimodal (vocal) (β = −0.11, *p* = 0.040).

#### Scratch produced

We further examined how the rate of scratch produced was related to the rates of communication using MRQAP (Fig. 1). Same-sex dyad partners had a higher rate of scratch produced than different-sex dyad partners (β = 0.20, *p* = 0.039). A higher rates of scratch produced were positively associated with higher rates of the following types of gestures: no object (β = 0.40, *p* = 0.004), visual (β = 0.48, *p* = 0.008), unimodal (β = 0.18, *p* = 0.049), multimodal (facial expression) (β = 0.18, *p* = 0.042), dyadic repertoire size (β = 0.35, *p* = 0.002), single (non-combined) (β = 0.25, *p* = 0.016), mutual attention present (β = 0.52, *p* = 0.001), bodily (β = 0.54, *p* < 0.001), events (β = 0.36, *p* = 0.003), manual indicative (β = 0.40, *p* = 0.003), close (β = 0.21, *p* = 0.023), far (β = 0.24, *p* = 0.001), non-repetitive (β = 0.41, *p* = 0.004), homogeneous (β = 0.37, *p* = 0.002), persistence (β = 0.57, *p* < 0.001) and piloerection (β = 0.23, *p* = 0.022). There was a significant negative association between rates of scratch produced and the rates of auditory long-range (β = −0.16, *p* = 0.046) and manual (β = −0.18, *p* = 0.011) gestures.

#### Scratch received

Same-sex dyad partners had a higher rate of scratch received than different-sex dyad partners (β = 0.20, *p* = 0.043). There was a significant negative association between rate of scratch received and the rate of auditory short-range gestures (β = −0.010, *p* = 0.049), gestures accompanied by mutual attention absent (β = −0.13, *p* = 0.049), close proximity (β = −0.11, *p* = 0.048), repetitive (β = −0.44, *p* = 0.044) and single (no sequence) (β = −0.14, *p* = 0.044) gestures.

### Association between the duration of social behaviour and gestural communication categorized according to function

We examined whether the duration of social behaviours (the dependent variable in all the models) was associated with the rate of different types of gestural communication categorized according to function. Chimpanzee dyads that produced higher rates of gesture threat to dominate spent a shorter duration of time jointly feeding (β = −0.50, *p* = 0.045), mutually grooming (β = −0.65, *p* = 0.006) and visually attending to each other (β = −0.64, *p* = 0.004). Gestures to give groom predicted a longer duration of time spent in jointly feeding (β = 0.17, *p* = 0.039), giving grooming (β = 0.69, *p* < 0.001), attention present (β = 0.25, *p* = 0.011) and absent (β = 0.24, *p* = 0.019), proximity (β = 0.30, *p* = 0.007). Further gestures to give groom elicited significantly lower rates of scratch received (β = −0.15, *p* = 0.020). Chimpanzee dyads that had higher rates of gestures in the context of mutual grooming spent a longer duration of time in mutual grooming (β = 1.14, *p* = 0.010) and visually attending to each other (β = 0.79, *p* = 0.025). Gestures to receive grooming were positively associated with the duration of time spent jointly feeding (β = 0.47, *p* = 0.012), giving grooming (β = 0.20, *p* = 0.036), mutual grooming (β = 0.53, *p* = 0.008), receiving grooming (β = 0.90, *p* = 0.001), attention present (β = 0.51, *p* = 0.001) and absent (β = 0.38, *p* = 0.022) and proximity (β = 0.55, *p* = 0.006). Higher rate of other threat gestures was negatively associated with a longer duration of time spent giving grooming (β = −0.06, *p* = 0.029) and were positively associated with a higher rate of scratch produced (β = 0.48, *p* = 0.002). Synchronized high-intensity panthoot was negatively associated with attention absent (β = −0.09, *p* = 0.049), proximity (β = −0.10, *p* = 0.012) and scratch received (β = −0.13, *p* = 0.019). Synchronized low-intensity panthoot was positively associated with joint feeding (β = 0.14, *p* = 0.043), travel (β = 0.26, *p* = 0.012), attention present (β = 0.10, *p* = 0.048) and absent (β = 0.13, *p* = 0.035) and proximity (β = 0.14, *p* = 0.023). A higher rates of play gestures were positively associated with the duration of give grooming (β = 0.10, *p* = 0.010) and negatively associated with the duration of time spent jointly feeding (β = =0.19, *p* = 0.004), mutual (β = −0.10, *p* = 0.012) and received grooming (β = −0.20, *p* = 0.002), attention present (β = −0.09, *p* = 0.017) and proximity (β = −0.08, *p* = 0.046). Higher rates of greeting gestures were associated with a longer duration of time spent giving grooming (β = 0.11, *p* = 0.025) and a shorter duration of time spent receiving grooming (β = −0.08, *p* = 0.041). Finally, higher rates of travel gestures were positively associated with a longer duration of time spent giving grooming (β = 0.10, *p* = 0.034) and travel (β = 0.10, *p* = 0.048).

## Discussion

Most studies of chimpanzee social relationships in the wild or in captivity focus on one aspect of relationships, such as proximity (A. I. Roberts & S. G. B. Roberts, 2016; A. I. Roberts & Roberts, 2017; S. G. B. Roberts & A. I. Roberts, 2016). Here we provide first systematic evidence that chimpanzees form social relationships across many different contexts more effectively with those individuals with whom they communicate through gestural communication.

Previous research on male chimpanzees suggested that maternal kinship did not play a critical role in social relationships. Thus, chimpanzees who shared the same mother did not affiliate or cooperate more often than expected by chance (Goldberg & Wrangham, 1997; John C Mitani et al., 2002). Although these relationships with maternal brothers appear to be important in infancy, demographic constraints on general availability of maternal brothers as potential coalition partners is believed to limit chimpanzee affiliation with maternal brothers (Goldberg & Wrangham, 1997; John C Mitani et al., 2002). On the basis of findings on affiliation and kinship among chimpanzee brothers we expected to find a weak affiliative relationships within mother and the adult offspring dyads. Against our expectation we found that mother offspring dyads maintained close proximity more often than unrelated dyads in co-feeding context but not other contexts. Although adult chimpanzees and mothers appear not to affiliate, they nonetheless remain in proximity in competitive contexts. This raises an important question as to why adult offspring chimpanzees co-feed with mothers if they do not affiliate with them. Research across chimpanzee populations in Africa shows that both adult male and female chimpanzees experience high levels of competition for plant food suggesting that these costs may be reduced by co-feeding with kin (Muller, 2002). Females who are unable to monopolize feeding site alone may engage in co-feeding with adult sons or daughters, thus enhancing their feeding success. In humans, unrelated individuals cooperate contingent upon strength of a social bond (friendship) and potential for reciprocity, whereas kinship has a direct effect on level of cooperation that is independent of social bond strength or reciprocity - the phenomenon termed ‘kinship premium’ (Curry, Roberts, & Dunbar, 2013). Kinship is more resilient to decay overtime, whereas friendships require more investment into social contact to maintain them at a level of strong social bonding (S. G. Roberts & Dunbar, 2011). The current finding in the chimpanzees that co-feeding with kin occurs in absence of affiliation, suggests that one might expect a similar effect of ‘kinship premium’ in the chimpanzees. The observations that unrelated sex and reproductive cohort dyads maintain proximity during feeding in presence of affiliation supports this suggestion.

Comparably important to kinship are relationships with partners from the same age cohort (Altmann, 1979). These relationships develop in infancy through association between mothers (Murray et al., 2014). Mirroring previous findings across Africa, the results show that these dyads are associated with high rates of affiliation in many different contexts (K. Langergraber et al., 2009; K. E. Langergraber et al., 2007; J.C. Mitani, Watts, & Muller, 2002). By expectation, the positive association between co-feeding and kinship should therefore also have emerged in the same age cohorts. However, these dyads differed in the patterns of co-feeding from other dyads. Chimpanzees from the same age cohort did not spend more time co-feeding in proximity than chimpanzees from different age cohort. Although these dyads share important interactions and behaviours, these relationships appear competitive and costly, possibly due to the fact that same age partners occupy similar niche and this might create greater competition. For instance, studies of mountain gorillas showed that when there is a high degree of co-feeding, individuals deal with competition by using different parts from the same food source (Watts, 1992). In contrast, same age cohort chimpanzees compete for the same food, dominance status, and position in the network. Same age partners share similar rank, but this equitability may create social relationships prone to decay because there is no prior consensus about the direction of potential aggression (F. B. de Waal, 1986; Flack, De Waal, & Waal, 2004). Indeed, partners of the same age class in rhesus macaques have been found to display high rates of both aggression and affiliation (Widdig, Nurnberg, Krawczak, Streich, & Bercovitch, 2002). To uncover the extent to which this lack of co-feeding in same age cohorts is generalizable to other sites it is first important to examine these relationships in small and large parties of chimpanzees, as these differ in the degree of feeding competition. In large parties when the feeding competition is greater, chimpanzees may associate with kin to diffuse competition, but in smaller parties they may well co-feed with same age partners.

Chimpanzees distribute their affiliative and competitive behaviour according to kinship and demography. However, can gestural communication play a role if influence of these factors has been discounted? In other words, are chimpanzees more affiliative than would be expected by chance, if affiliation was dependent on gestural communication alone? Here our findings contribute to prior research by showing that gestural communication exerts important influence on patterns of social affiliation among chimpanzee dyads. After controlling for kinship and demography, the data shows that chimpanzees spend longer duration of time affiliating with the individuals with whom they communicate through gestures. Thus, as there is an increase in the duration of time A spends in social behaviour with B, there is a corresponding increase or decrease in the rate of gestural communication emitted by A to B. These findings support earlier research from vocal modality which showed that sociality is associated with use of vocal communication (McComb & Semple, 2005). Affilitive gestures such as gestures to initiate grooming or play appear to have an especially important influence on affiliation. Affiliative gestural communication positively influenced the duration of time spent in social behaviour, whereas antagonistic gestural communication negatively influenced duration of time spent in social behaviour. For instance, the higher rate of gestures to initiate giving grooming predicts longer grooming, whereas higher rate of gestures to threaten predict shorter grooming. Thus, affiliative gestures are important in maintaining social relationships and function to coordinate these interactions. For instance, affiliative gestural communication is more common in socially complex egalitarian pigtail macaques, than in less socially complex and despotic rhesus macaques (Maestripieri, 1999).

However, affiliation contexts are not uniformly associated with different degrees of gestural complexity but there is a vast variation in how these measures of complexity are distributed across contexts. In contexts signifying stronger social bonding (e.g. mutual grooming, joint travel, mutual visual monitoring) gestures are primarily of low intensity (visual or auditory short range), ranging from 16 to 17 different forms, that are more complex (e.g. use of facial expressions, gestures were combined with other gesture types). In contrast, in contexts signifying weaker social bonding (e.g. unidirectional grooming) (Noë, 2006) gestures are of higher intensity (tactile, auditory), take 13 different forms that tend to be simpler (e.g. unimodal gesture, non-combined with other gesture types). Gestures given in strong social bonding contexts may therefore be qualitatively different from gestures given when social bonds are weaker.

Why is there such a variation in gesture complexity across contexts? Is this variation attributed to the efficiency with which communication can influence the recipient? The most parsimonious explanation of these data is that chimpanzees in the wild are making decisions on how to communicate, by choosing the gestures that can best influence the recipient in the given context. Our data supports previous research suggesting that there are two key pathways to achieving a greater efficiency in gestural communication. Studies in chimpanzees, showed how intensity of communication can influence efficiency of social bonding in relation to its effectiveness of conveying signaller’s goal specifically (A. I. Roberts & S. G. B. Roberts, 2016). For instance, visual bodily gestures accompanied by direct gaze and pointing, have a greater success in coordinating proximity with the recipient than when using visual bodily gestures alone (Conty et al., 2012; S. Roberts & A. I. Roberts, 2018). Thus by making low intensity, inspecific gestures more complex, signallers can increase specificity of their gestures and elicit more accurate response from the recipient. Another way to increase specificity of the gesture is by increasing their intensity and therefore greater complexity of the gesture is not required (A. I. Roberts & S. G. B. Roberts, 2016). This link between intensity of communication and specificity of the response has been shown in studies of nonverbal communication in humans (Zajonc & Sales, 1966). As the intensity of the signal increases, the mapping between structure of the signal and the accompanying context becomes tighter enabling recipient to more accurately make the association between the signal and the signaller’s goal. Finally, adding rewarding property to the signal may increase signal’s effectiveness in conducting successful interaction (A. I. Roberts & S. G. B. Roberts, 2018). These rewarding properties promote greater commitment to the interaction so that the recipient is more likely to respond to the gesture.

The relationships based on behaviours such as mutual grooming or travel appear to be more mutually appealing as shown by a longer duration of time invested in these behaviours between partners who spend more time in proximity. These relationships are more common between same age cohort partners – growing up together provides context whereby dyads share many similarities and are highly familiar with each other. However, a whole range of signals responsible for maintaining these interactions, for example visual gestures, persistence, repertoire, facial expression were associated with higher rate of self-scratch, indicating higher anxiety experienced by the signallers towards recipients of these gestures. One type of answer that can be given to the question of why chimpanzees experienced higher anxiety in response to communicating with social partners with whom they displayed a higher rate of affiliative behaviour is that social relationships with these social partners have been uncertain due to unresolved dominance relationships. What has long puzzled anthropologists is that dominance hierarchy can sometimes lead to greater social cohesion because it enables signaller to more effectively predict outcome of the interaction before they engage in the interaction with the recipient (Flack, Girvan, De Waal, & Krakauer, 2006). Classical ethologists have shown very clearly how lack of linear dominance hierarchy can make animal societies less predictable and more aggressive (Flack, de Waal, & Krakauer, 2005) demanding complex but low intensity communication to resolve ambiguity in social relationships (A. I. Roberts et al., 2019). This communication can increase trust of the recipient, as it may appear more positive, therefore creating a perception of a fitness rewarding intent of the signaler (Roberts & Roberts, 2016a). In social relationships with dominant chimpanzees, the risks of interaction and direction of potential aggression is known in advance and therefore dominance relationships increase certainty by having predictable outcomes (Ay, Flack, & Krakauer, 2007; Flack et al., 2006). In contrast, equitable ranks are less predictable in that both interactants are equally likely to win if engaged in a fight resulting in high levels of uncertainty. The mutual appeal that draws chimpanzees together in these social interactions, has been insufficient and the power of gestural communication has been exploited to facilitate social interactions between these partners. Low intensity signals designed to effectively convey the intentions of a signaller in a way that leads to a reduction in uncertainty of the recipient about the signaller’s goal encompass all common signals seen in these contexts.

In contrast, the relationships based on behaviours such as unidirectional grooming are often directed towards chimpanzees who display higher rank. Gestures made in unidirectional grooming context were associated with reduced rate of self-scratching by the recipient, suggesting that recipients experience reduced anxiety when receiving these gestures. This reveals a possible lack of mutual appeal in these interactions as the signaller actively influences recipient’s positive affect to enhance its willingness to associate. As the anxiety is reduced, the signaller increases recipient’s commitment to the social interaction forging stronger social bonding. Thus, in these contexts, in addition to conveying goals effectively through more intense gestures, use of signals that increase rewards are important to succeed in engaging the recipient who may otherwise not be particularly interested in the interaction.

The differentiated communication strategies of the chimpanzees may have evolved in response to the demands imposed by competition in complex social settings. Both male and female chimpanzees often compete for food and individual’s ability to gain access to food can influence their reproductive success (Muller, 2002). Evidence from chimpanzees indicates that social variables such as party size can affect individual’s relative competitive success in feeding contexts. Chimpanzees in larger parties derive benefits from lower predation pressure but may face a higher feeding competition than the chimpanzees in smaller parties. Competition for food may therefore promote evolution of the strategies the individuals use to diffuse this competition. Influential strategy includes grooming to reduce likelihood of potential aggression during co-feeding. In small social groups, chimpanzees face lower social competition and thus can invest in grooming with the social partners that are insecure but appealing. However, when the size of the group increases, the safety of secure relationships promotes feeding efficiency and therefore grooming interactions focus on these social partners. Tactile gestures that have a rewarding property can compete for recipient’s attention in larger groups more effectively than visual gestures by being more capable of redirecting recipient’s attention from the wider audience onto the signaller. Furthermore, rewarding gestures may have a better coordination value by being able to influence recipient’s behaviour more directly through more intense and specific gestures (A. I. Roberts & S. G. B. Roberts, 2016). Studies showed that when larger audience is present, both rewarding and coordination properties of gestural communication facilitate longer grooming and this in turn is associated with longer co-feeding in proximity (A. I. Roberts, 2018). Given the time budgeting constraints on grooming (Dunbar, 1992), there may be a limit for use of grooming as a tool for diffusing social competition in feeding contexts. Our data reveals that chimpanzees incorporated use of objects and vocalisations when gesturing towards conspecifics with whom they spent short periods of time in proximity. Similarly, use of objects such as a trunk of a tree to make sounds accompanied by use of rhythmic vocalisations (‘synchronized high-intensity pant hoot’) may potentially fulfil such a function as this communication is used most often when joining feeding sites or during travel (Clark & Wrangham, 1994; S. G. B. Roberts & A. I. Roberts, 2016). Supporting these suggestions, our evidence shows that synchronized high-intensity pant hoots were associated with reduced displacement behaviours in the recipient. When social parties become large, the ability to groom with conspecifics may decline and ‘synchronized high-intensity pant hoot’ may facilitate social cohesion by reducing anxiety of the recipients (S. G. B. Roberts & A. I. Roberts, 2016). This property may draw attention of the recipients from the wider audience onto the signaller facilitating longer interactions such as travel or co-feeding. This finding supports research on humans (Jackson et al., 2018) showing that joint, high intensity behaviours have a role in social cohesion by being intensely rewarding to the dyad partners, regardless of the history of prior interaction.

The complexity of a social system depends on the complexity of individual relationships between animals, as the individual-level social interactions scale to the emergent properties found in the social system (Krause, Croft, & James, 2007). In this study we highlight the tactical decisions that wild chimpanzees make in their use of gestural communication to develop and maintain complexity of their social system. The selective pressures arising from maintaining this social complexity is proposed to have played a key role both in the evolution of communicative complexity and in the evolution of larger brains both in primates and in hominins. However, detailed behavioural evidence of an association between communicative complexity and these differentiated social relationships is lacking. Here we address this issue by examining how different types of social behaviour relate to patterns of gestural communication. Overall, our results suggest that differentiated patterns of gestural communication can help chimpanzees maintain a network of differentiated social relationships, and that this may allow individual primates to successfully navigate a complex social world. Future studies exploring the relationship between the complexity of communication skills, sociality and brain size across a range of primate species would allow for a deeper understanding of the association between complex social systems and complex communication.

## Acknowledgement

The data collection was funded by the Economic and Social Research Council 3+ fellowship and the University of Stirling. We thank the Royal Zoological Society of Scotland for providing core funding to the Budongo Conservation Field Station, Uganda, as well as the Ugandan Wildlife Authority and the Uganda National Council for Science and Technology for permission to carry out this research. We are most grateful to Prof. Klaus Zuberbuhler and Geresomu Muhumuza for fantastic fieldwork.

## Supplementary Information 1 Social bonding, gestural complexity and displacement behaviour of wild chimpanzee

**Table S1.**
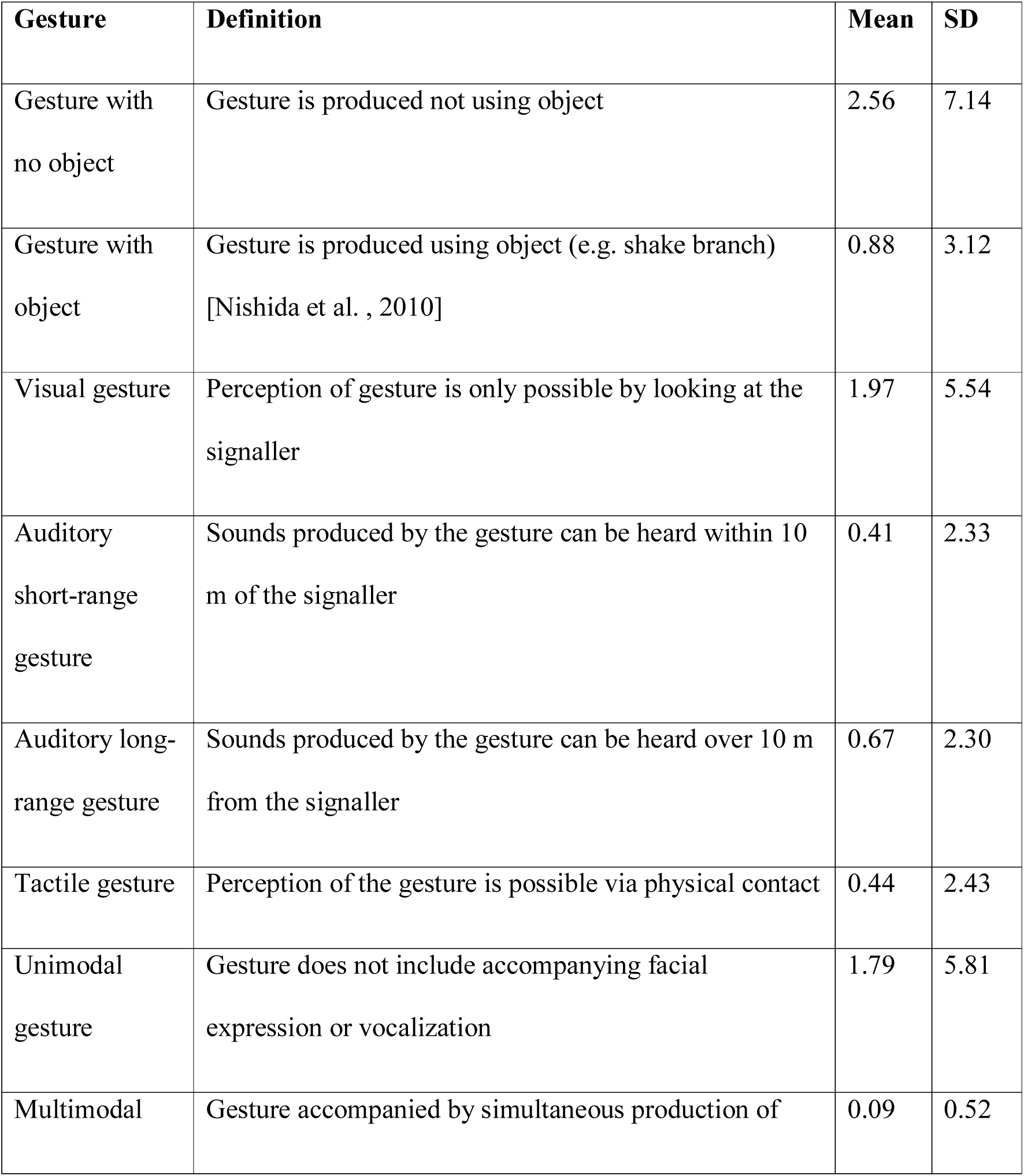
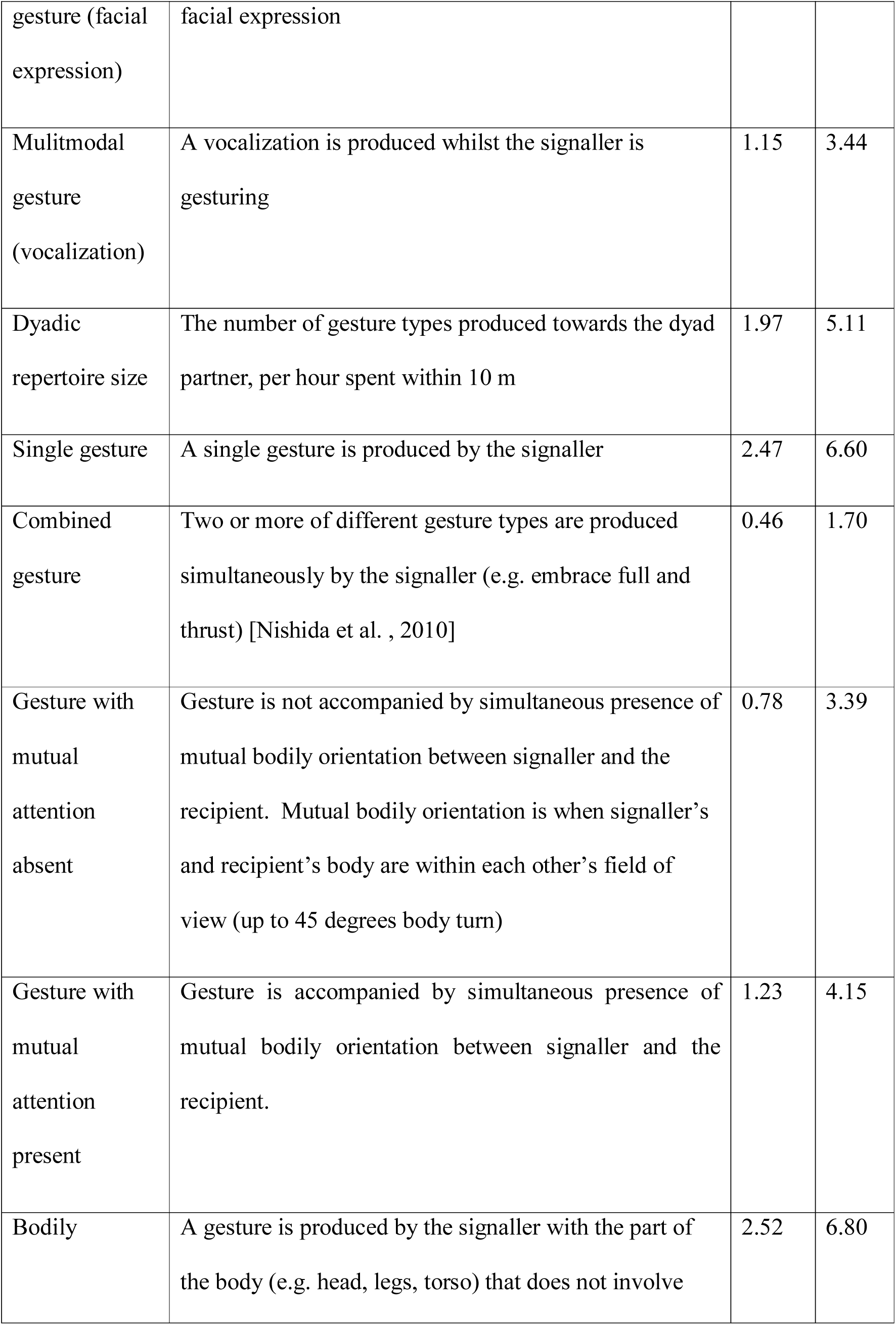
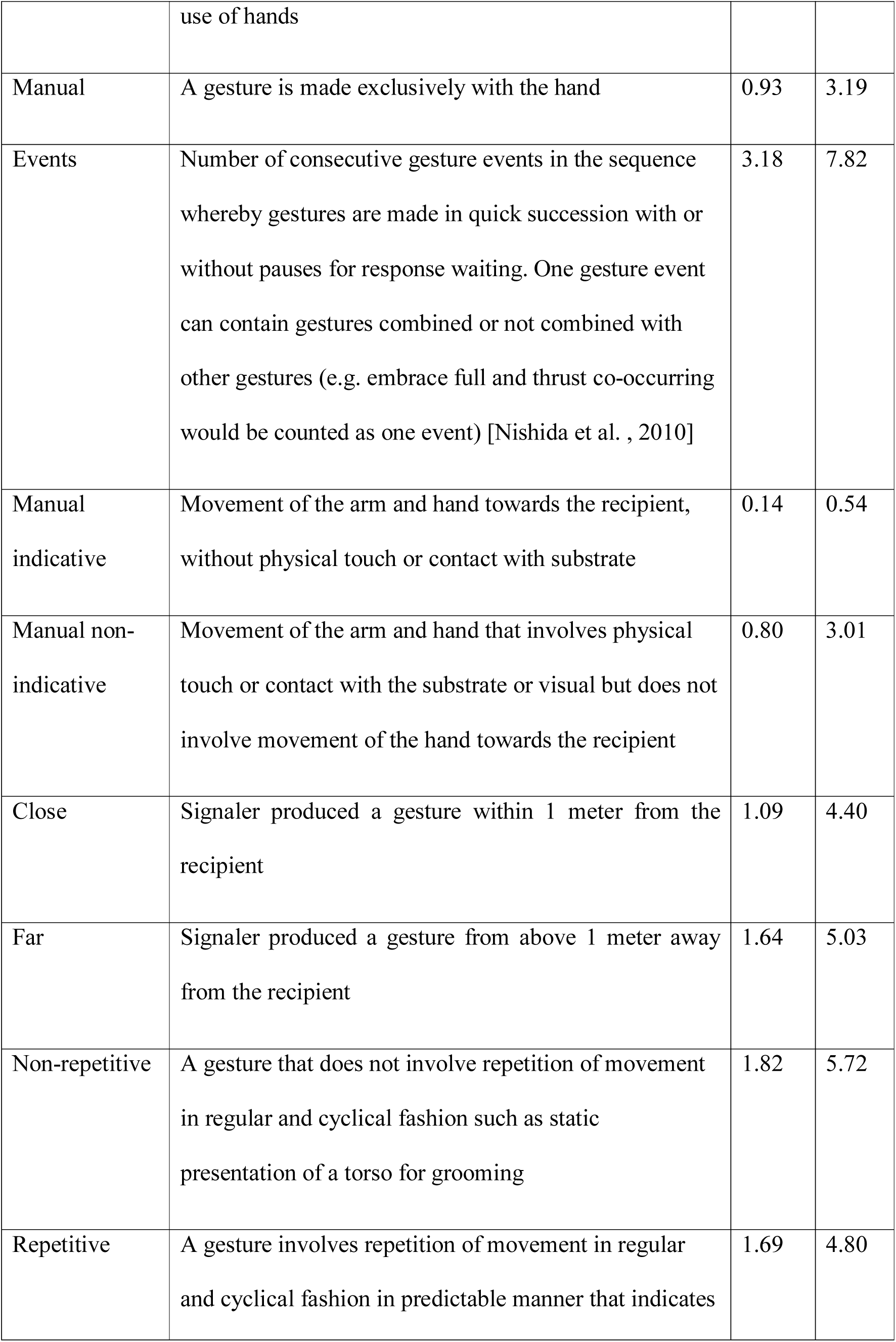
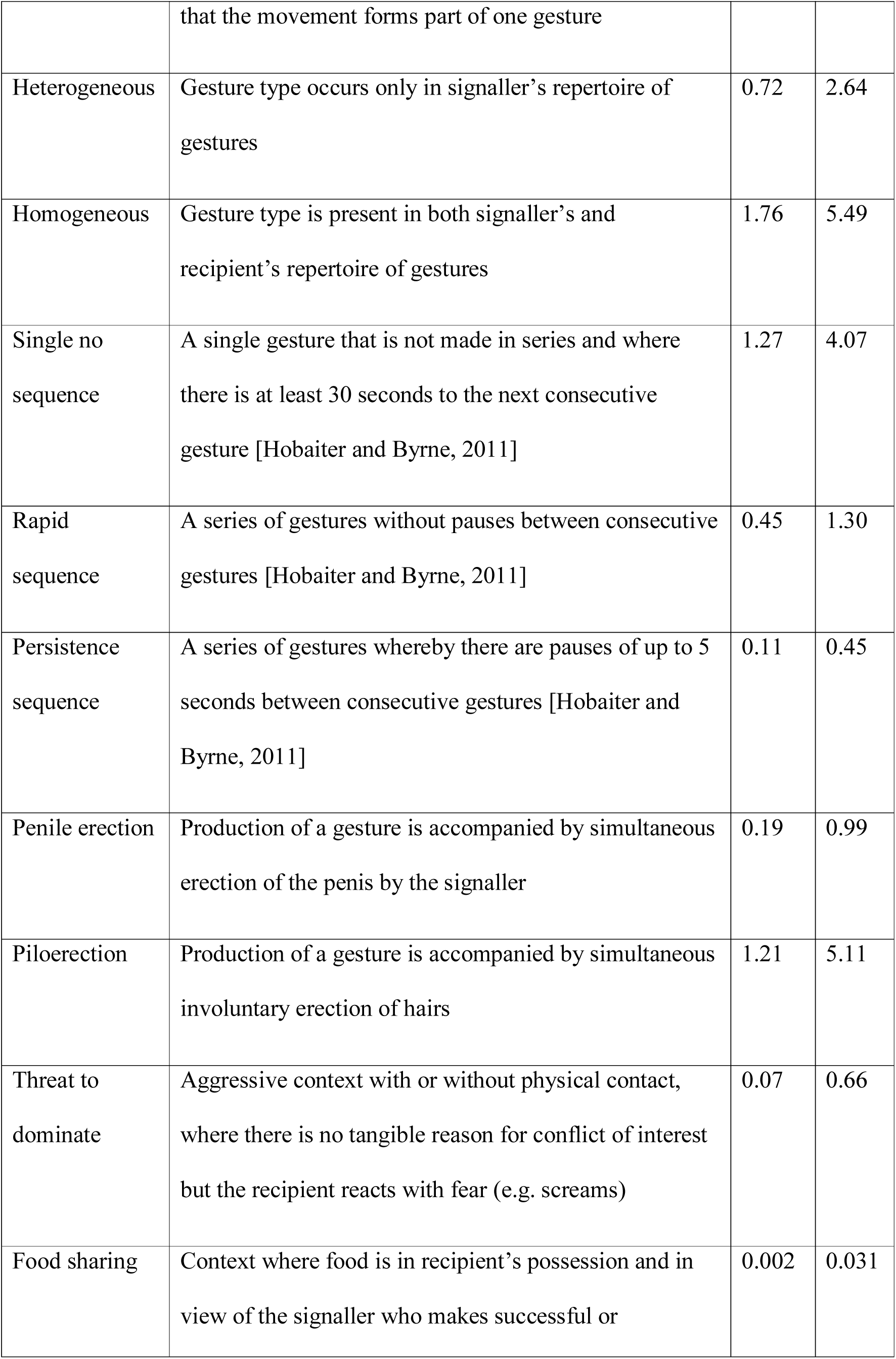
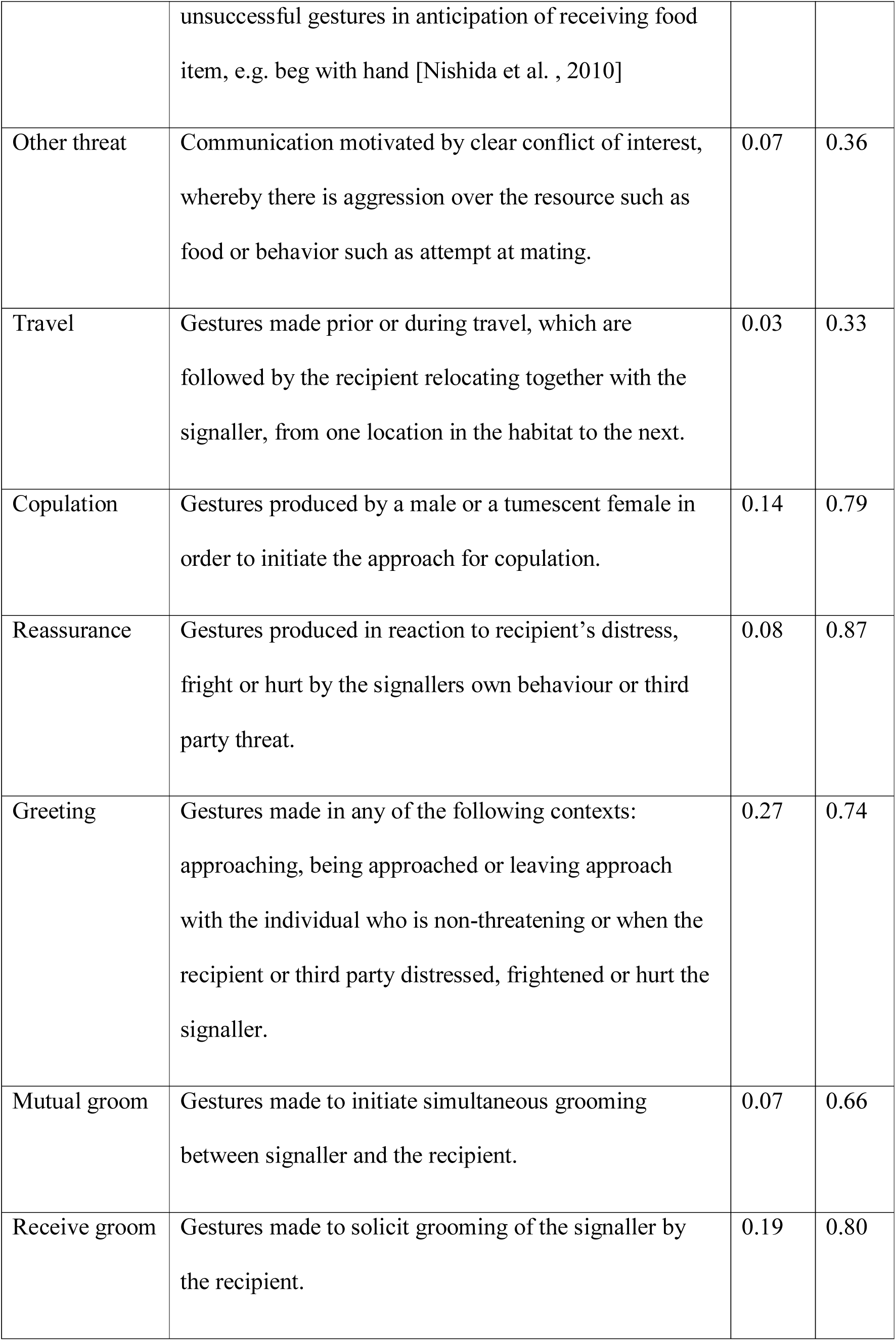
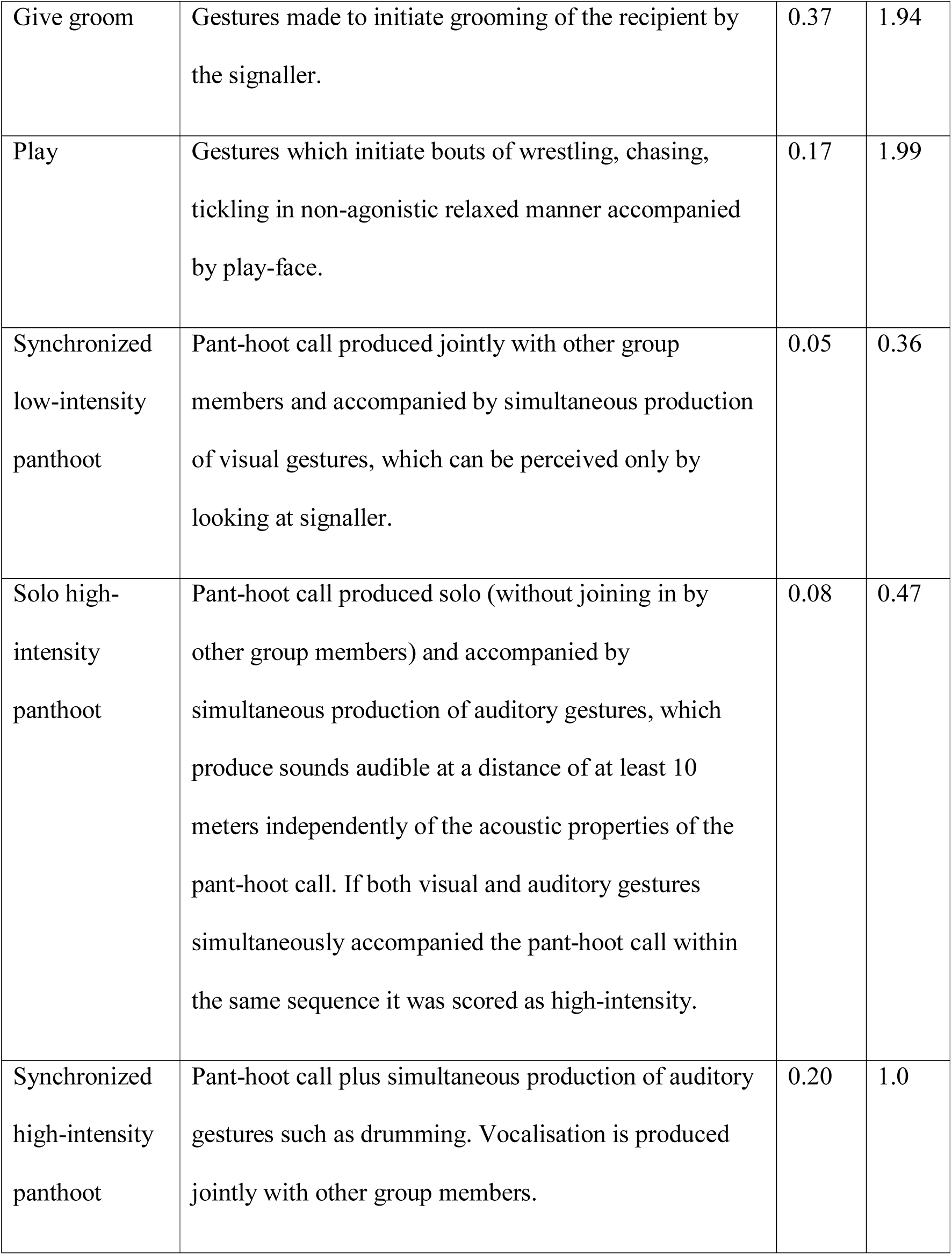
Definitions, means and standard deviations (SD) for different types of gestural communication. Data based on gestural communication between 132 chimpanzee dyads. All gestural communication measured as the rate per hour dyad spent within 10 m. For detailed description of all gesture types and accompanying video footage see [Roberts et al., 2014].

**Table S2.**
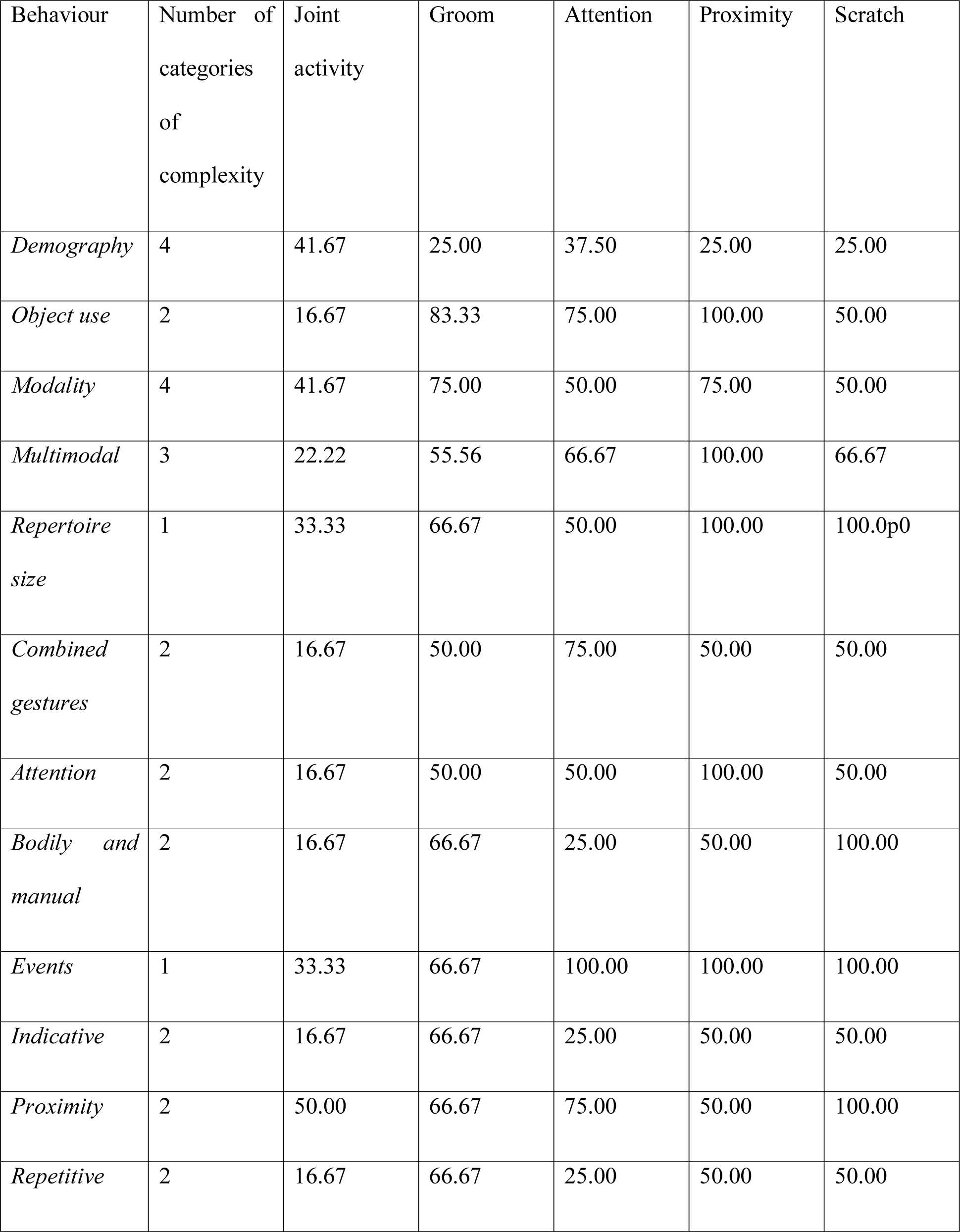
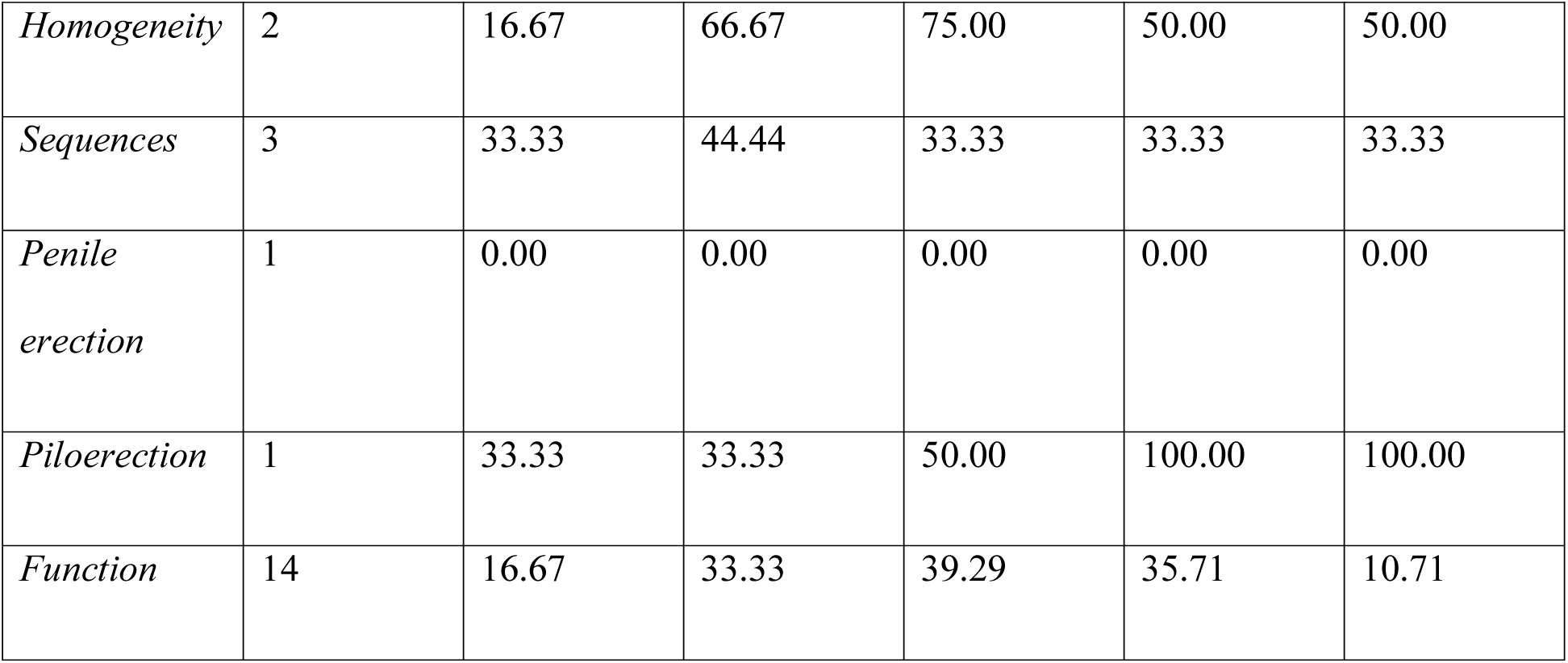
Percentage of indicators for each behavioural categories that is significantly associated with behavioural indices of 5 domains of sociality (joint activity, grooming, visual attention, proximity and scratch).

| Behaviour | Number of categories of complexity | Joint activity | Groom | Attention | Proximity | Scratch |
| --- | --- | --- | --- | --- | --- | --- |
| *Demography* | 4 | 41.67 | 25.00 | 37.50 | 25.00 | 25.00 |
| *Object use* | 2 | 16.67 | 83.33 | 75.00 | 100.00 | 50.00 |
| *Modality* | 4 | 41.67 | 75.00 | 50.00 | 75.00 | 50.00 |
| *Multimodal* | 3 | 22.22 | 55.56 | 66.67 | 100.00 | 66.67 |
| *Repertoire size* | 1 | 33.33 | 66.67 | 50.00 | 100.00 | 100.00 |
| *Combined gestures* | 2 | 16.67 | 50.00 | 75.00 | 50.00 | 50.00 |
| *Attention* | 2 | 16.67 | 50.00 | 50.00 | 100.00 | 50.00 |
| *Bodily and manual* | 2 | 16.67 | 66.67 | 25.00 | 50.00 | 100.00 |
| *Events* | 1 | 33.33 | 66.67 | 100.00 | 100.00 | 100.00 |
| *Indicative* | 2 | 16.67 | 66.67 | 25.00 | 50.00 | 50.00 |
| *Proximity* | 2 | 50.00 | 66.67 | 75.00 | 50.00 | 100.00 |
| *Repetitive* | 2 | 16.67 | 66.67 | 25.00 | 50.00 | 50.00 |
| *Homogeneity* | 2 | 16.67 | 66.67 | 75.00 | 50.00 | 50.00 |
| *Sequences* | 3 | 33.33 | 44.44 | 33.33 | 33.33 | 33.33 |
| *Penile erection* | 1 | 0.00 | 0.00 | 0.00 | 0.00 | 0.00 |
| *Piloerection* | 1 | 33.33 | 33.33 | 50.00 | 100.00 | 100.00 |
| *Function* | 14 | 16.67 | 33.33 | 39.29 | 35.71 | 10.71 |

### Association between the duration of social behavior and gestural communication categorized according to structure

#### Demographic Factors

Supplementary Table S3. MRQAP regression models predicting durations of social behavior, per hour dyad spent within 10m. Predictors were demographic variables. Dyads were classified as same age or different age (within 5 years), same sex or different sex, related by maternal kinship and as the same or different reproductive status (reproductively active, not reproductively active). Based on 132 dyadic relationships of the chimpanzees. Significant *p* values are indicated in bold. R squared (*r^2^)* denotes amount of variance in the dependent variable explained by the regression model.

**Table S3.1.** Duration of joint feeding behaviour (*r^2^* = 0.119)

|  | Standardized coefficient | Standard error | *p* |
| --- | --- | --- | --- |
| Age | -0.01 | 0.65 | 0.49 |
| Sex | 0.31 | 0.69 | **0.001** |
| Kinship | 0.29 | 1.28 | **0.01** |
| Reproductive status | 0.22 | 0.72 | **0.04** |

**Table S3.2.** Duration of joint resting behaviour (*r^2^* = 0.070)

|  | Standardized coefficient | Standard error | *p* |
| --- | --- | --- | --- |
| Age | 0.29 | 1.37 | **0.003** |
| Sex | -0.14 | 1.31 | 0.13 |
| Kinship | -0.02 | 2.25 | 0.47 |
| Reproductive status | -0.04 | 1.46 | 0.36 |

**Table S3.3.** Duration of joint travelling behaviour (*r^2^* = 0.09)

|  | Standardized coefficient | Standard error | *p* |
| --- | --- | --- | --- |
| Age | 0.28 | 0.42 | **0.01** |
| Sex | 0.05 | 0.39 | 0.68 |
| Kinship | 0.01 | 0.72 | 0.86 |
| Reproductive status | 0.01 | 0.43 | 0.93 |

**Table S3.4.** Duration of giving grooming (*r^2^* = 0.07)

|  | Standardized | Standard error | *p* |
| --- | --- | --- | --- |

|  | coefficient |  |  |
| --- | --- | --- | --- |
| Age | 0.21 | 0.54 | **0.02** |
| Sex | 0.11 | 0.53 | 0.18 |
| Kinship | 0.10 | 0.96 | 0.12 |
| Reproductive status | 0.03 | 0.66 | 0.42 |

**Table S3.5.** Duration of mutual grooming (*r^2^* = 0.04)

|  | Standardized coefficient | Standard error | *p* |
| --- | --- | --- | --- |
| Age | 0.20 | 0.70 | **0.050** |
| Sex | 0.02 | 0.72 | 0.49 |
| Kinship | 0.02 | 1.29 | 0.29 |
| Reproductive status | 0.14 | 0.87 | 0.18 |

**Table S3.6.**
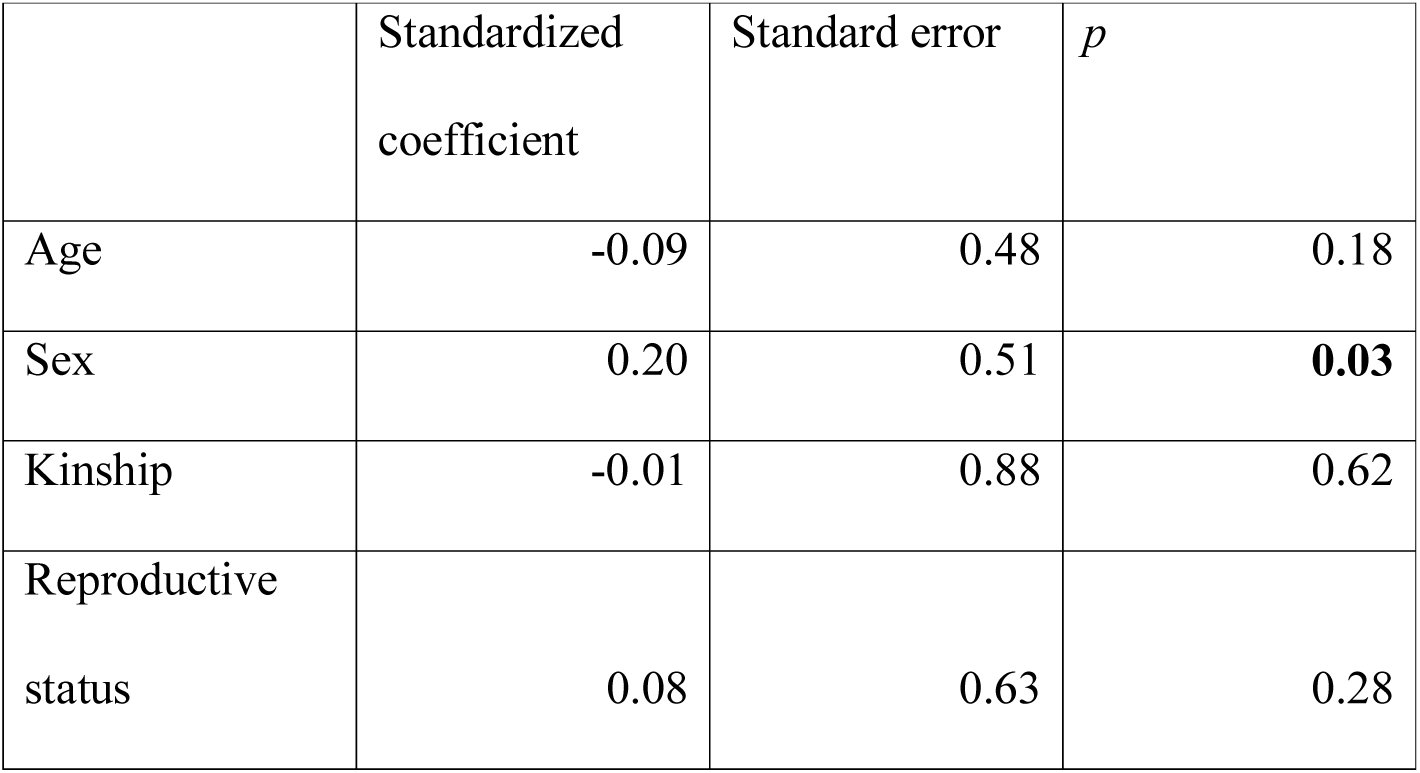
Duration of receiving grooming (*r^2^*= 0.03)

|  | Standardized coefficient | Standard error | *p* |
| --- | --- | --- | --- |
| Age | -0.09 | 0.48 | 0.18 |
| Sex | 0.20 | 0.51 | **0.03** |
| Kinship | -0.01 | 0.88 | 0.62 |
| Reproductive status | 0.08 | 0.63 | 0.28 |

**Table S3.7.**
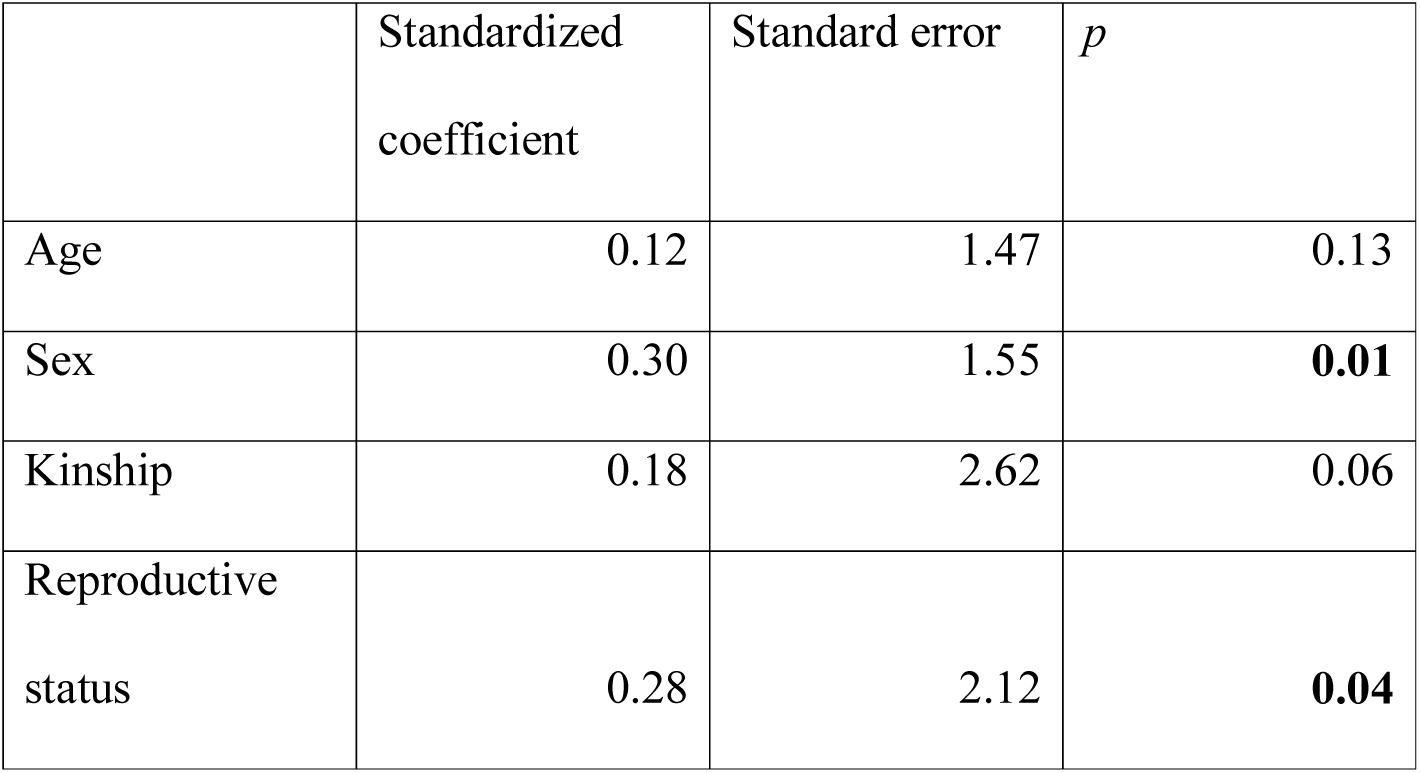
Duration of visual attention towards dyad partner (*r^2^*= 0.11)

**Table S3.8.** Duration of visual attention away dyad partner (*r^2^* = 0.12)

|  | Standardized<br>coefficient | Standard error | *p* |
| --- | --- | --- | --- |
| Age | 0.367 | 1.730 | **0.001** |
| Sex | -0.064 | 1.277 | 0.251 |
| Kinship | 0.071 | 2.783 | 0.145 |
| Reproductive<br>status | 0.041 | 1.395 | 0.370 |

**Table S3.9.**
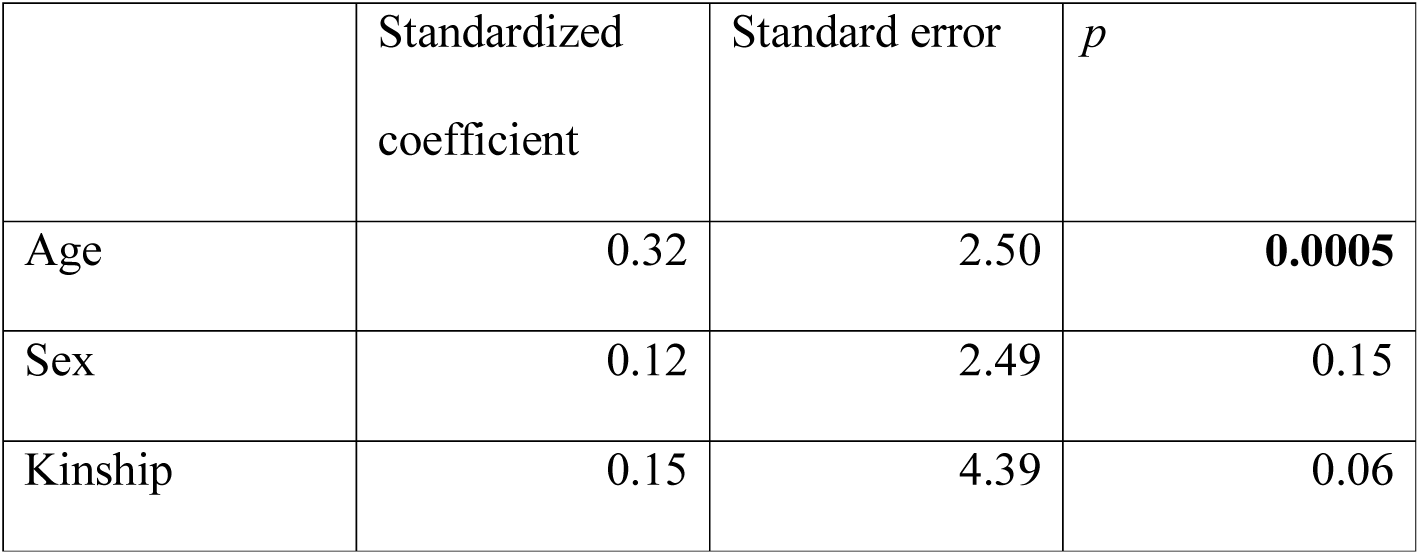
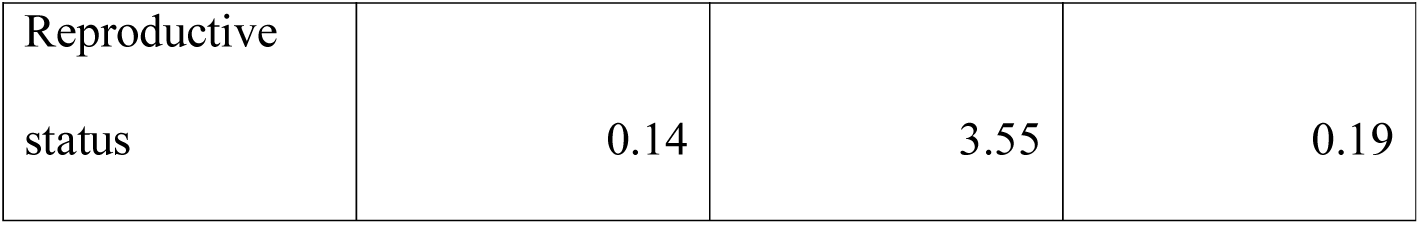
Duration of time in close proximity – within 2 m (*r^2^*= 0.11)

|  | Standardized<br>coefficient | Standard error | *p* |
| --- | --- | --- | --- |
| Age | 0.32 | 2.50 | **0.0005** |
| Sex | 0.12 | 2.49 | 0.15 |
| Kinship | 0.15 | 4.39 | 0.06 |
| Reproductive status | 0.14 | 3.55 | 0.19 |

**Table S3.10.** Rate of scratch produced (*r^2^*= 0.046)

|  | Standardized coefficient | Standard error | *p* |
| --- | --- | --- | --- |
| Age | 0.016 | 0.606 | 0.41829 |
| Sex | 0.201 | 0.560 | **0.03948** |
| Kinship | 0.079 | 1.143 | 0.18341 |
| Reproductive status | 0.030 | 0.423 | 0.38131 |

**Table S3.11.** Rate of scratch received (*r^2^* = 0.046)

|  | Standardized coefficient | Standard error | *p* |
| --- | --- | --- | --- |
| Age | 0.02 | 0.61 | 0.42779 |
| Sex | 0.20 | 0.56 | **0.04348** |
| Kinship | 0.08 | 1.14 | 0.18991 |
| Reproductive status | 0.03 | 0.44 | 0.37481 |

#### Object use in gestural communication

Supplementary Table S4. MRQAP regression models predicting durations of social behavior, per hour dyad spent within 10m. Predictor variables were rates of gestural communication (without objects and with objects) and demographic variables. Based on 132 chimpanzee dyads. Significant *p* values are indicated in bold. R squared (*r^2^)* denotes amount of variance in the dependent variable explained by the regression model.

**Table S4.1.** Duration of joint feeding behaviour (*r^2^*= 0.12)

| | Standardized<br>coefficient | Standard error | $p$ |
| --- | --- | --- | --- |
| Age | -0.01 | 0.68 | 0.47 |
| Sex | 0.31 | 0.70 | **0.003** |
| Kinship | 0.29 | 1.35 | **0.009** |
| Reproductive<br>status | 0.22 | 0.76 | **0.047** |
| Object use | -0.01 | 0.08 | 0.56 |
| No object use | 0.03 | 0.04 | 0.30 |

**Table S4.2.** Duration of joint resting behaviour (*r^2^* = 0.08)

| | Standardized<br>coefficient | Standard error | $p$ |
| --- | --- | --- | --- |
| Age | 0.28 | 1.45 | **0.01** |
| Sex | -0.16 | 1.37 | 0.10 |
| Kinship | -0.02 | 2.41 | 0.42 |
| Reproductive<br>status | -0.05 | 1.45 | 0.35 |
| Object use | -0.08 | 0.18 | 0.07 |
| No object use | 0.10 | 0.08 | 0.07 |

**Table S4.3.**
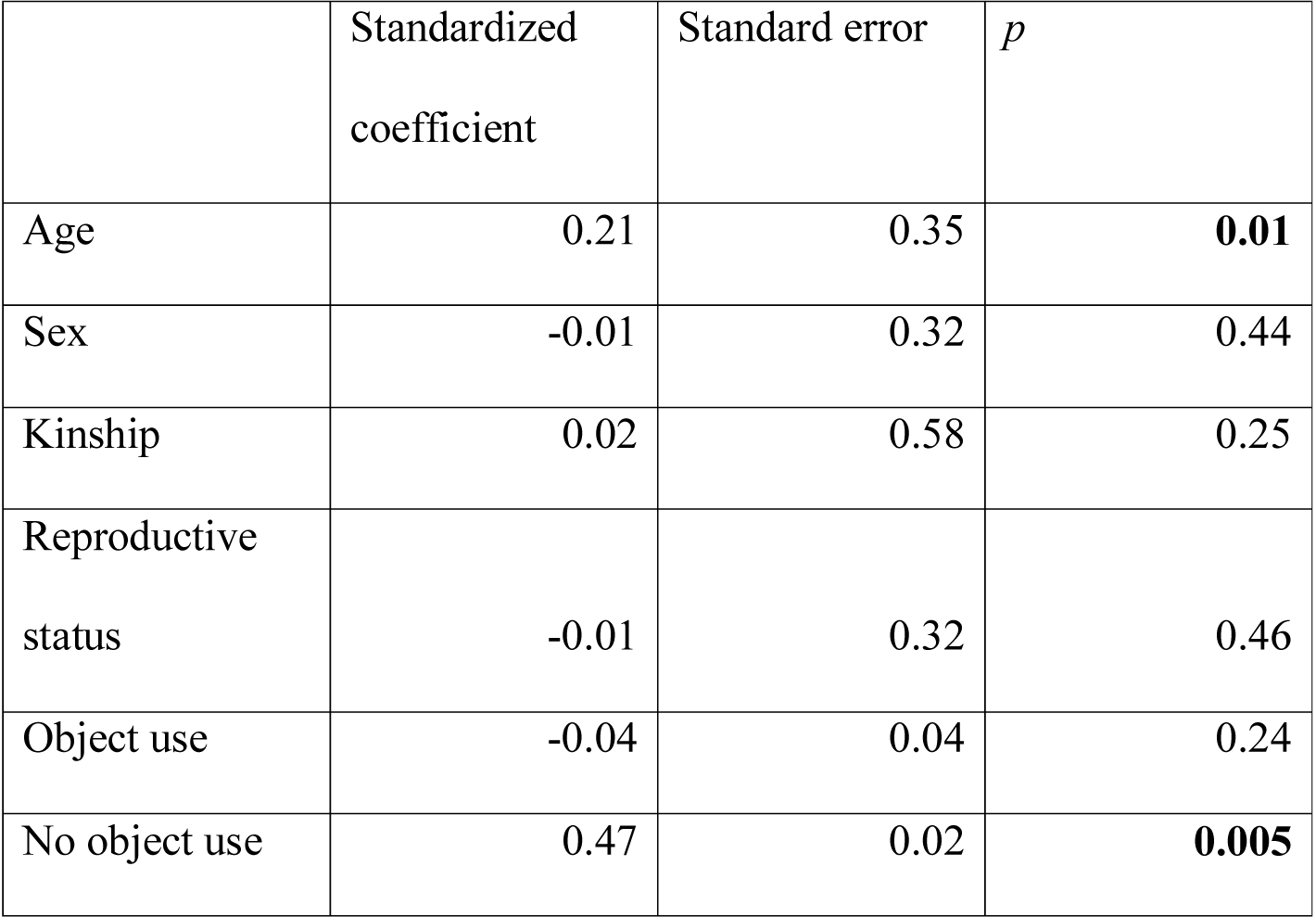
Duration of joint travelling behaviour (*r^2^*= 0.29)

|  | Standardized<br>coefficient | Standard error | *p* |
| --- | --- | --- | --- |
| Age | 0.21 | 0.35 | **0.01** |
| Sex | -0.01 | 0.32 | 0.44 |
| Kinship | 0.02 | 0.58 | 0.25 |
| Reproductive<br>status | -0.01 | 0.32 | 0.46 |
| Object use | -0.04 | 0.04 | 0.24 |
| No object use | 0.47 | 0.02 | **0.005** |

**Table S4.4.** Duration of giving grooming (*r^2^* = 0.39)

|  | Standardized<br>coefficient | Standard error | *p* |
| --- | --- | --- | --- |
| Age | 0.12 | 0.45 | 0.08 |
| Sex | 0.02 | 0.44 | 0.41 |
| Kinship | 0.10 | 0.85 | 0.08 |
| Reproductive<br>status | 0.00 | 0.46 | 0.47 |
| Object use | -0.20 | 0.06 | **0.01** |
| No object use | 0.64 | 0.04 | **0.001** |

**Table S4.5.** Duration of mutual grooming (*r^2^*= 0.38)

|  | Standardized<br>coefficient | Standard error | *p* |
| --- | --- | --- | --- |
| Age | 0.11 | 0.55 | 0.10 |
| Sex | -0.07 | 0.53 | 0.24 |
| Kinship | 0.02 | 1.00 | 0.26 |
| Reproductive<br>status | 0.10 | 0.58 | 0.12 |
| Object use | -0.15 | 0.07 | **0.02** |
| No object use | 0.64 | 0.04 | **0.0005** |

**Table S4.6.** Duration of receiving grooming (*r^2^* = 0.07)

|  | Standardized<br>coefficient | Standard error | *p* |
| --- | --- | --- | --- |
| Age | -0.12 | 0.49 | 0.09 |
| Sex | 0.17 | 0.49 | 0.05 |
| Kinship | 0.00 | 0.91 | 0.66 |
| Reproductive<br>status | 0.07 | 0.56 | 0.27 |
| Object use | -0.04 | 0.06 | 0.34 |
| No object use | 0.22 | 0.03 | **0.04** |

**Table S4.7.** Duration of visual attention towards dyad partner (*r^2^* = 0.49)

|  | Standardized<br>coefficient | Standard error | *p* |
| --- | --- | --- | --- |
| Age | 0.02 | 1.10 | 0.39 |
| Sex | 0.20 | 1.12 | **0.01** |
| Kinship | 0.19 | 2.11 | **0.01** |
| Reproductive status | 0.24 | 1.42 | **0.01** |
| Object use | -0.18 | 0.14 | **0.003** |
| No object use | 0.69 | 0.10 | **0.0005** |

**Table S4.8.**
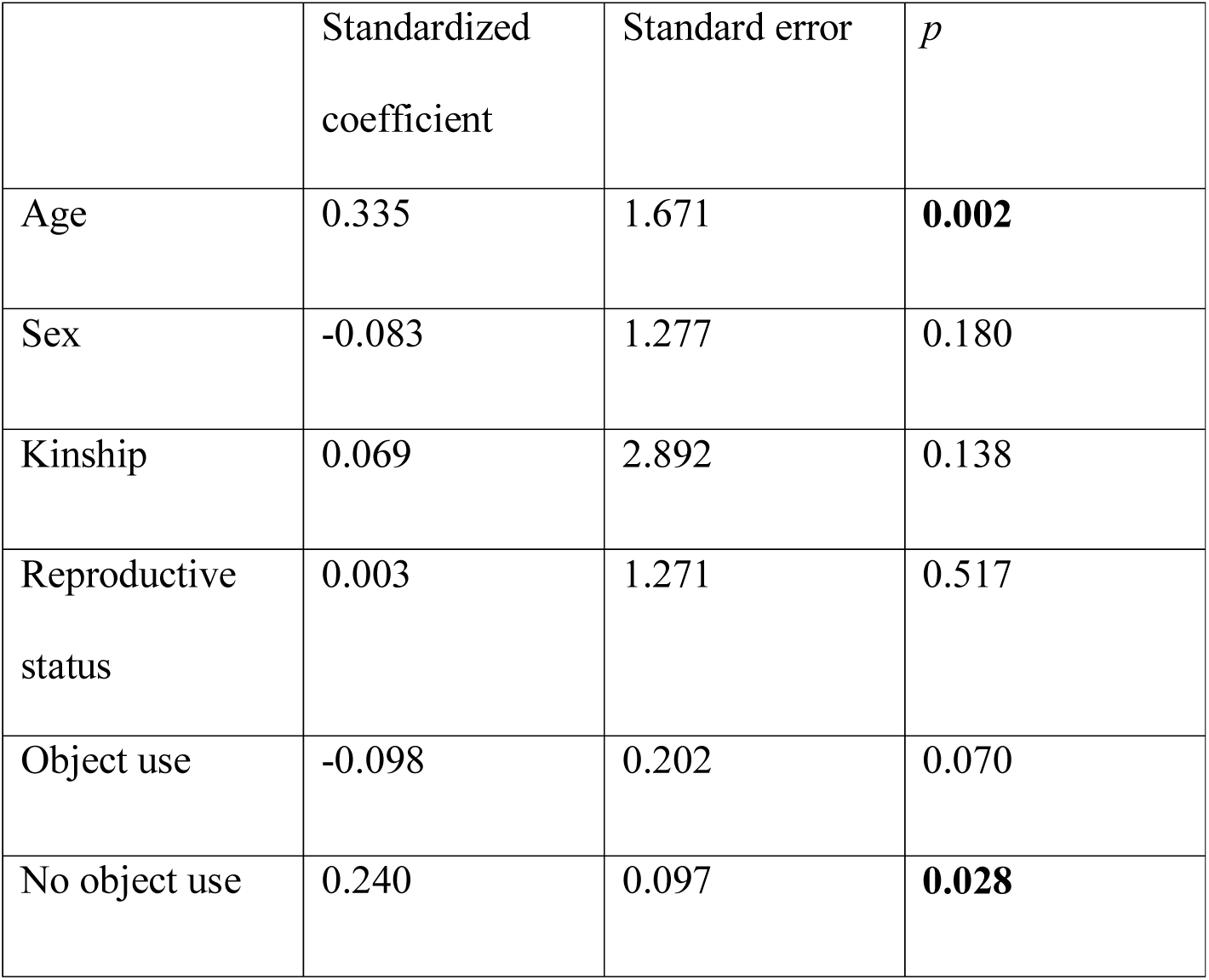
Duration of visual attention away from dyad partner (*r^2^*= 0.16)

|  | Standardized coefficient | Standard error | *p* |
| --- | --- | --- | --- |
| Age | 0.335 | 1.671 | **0.002** |
| Sex | -0.083 | 1.277 | 0.180 |
| Kinship | 0.069 | 2.892 | 0.138 |
| Reproductive status | 0.003 | 1.271 | 0.517 |
| Object use | -0.098 | 0.202 | 0.070 |
| No object use | 0.240 | 0.097 | **0.028** |

**Table S4.9.**
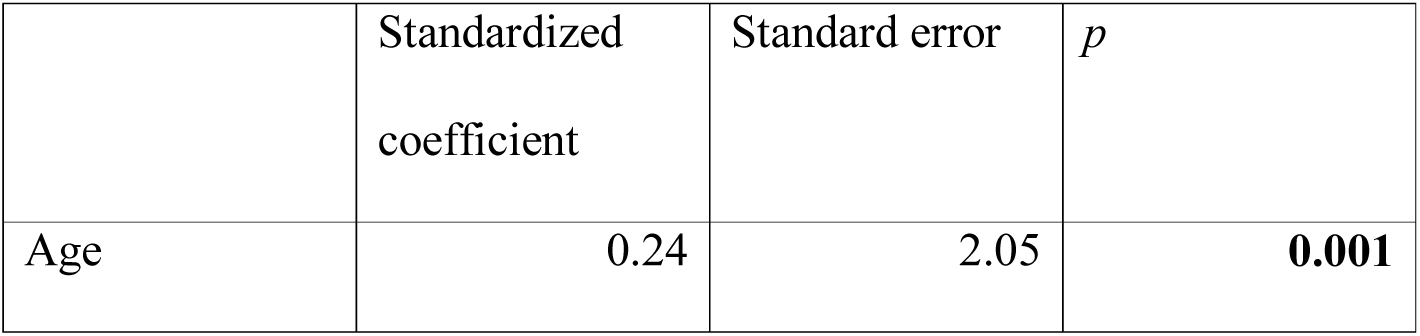
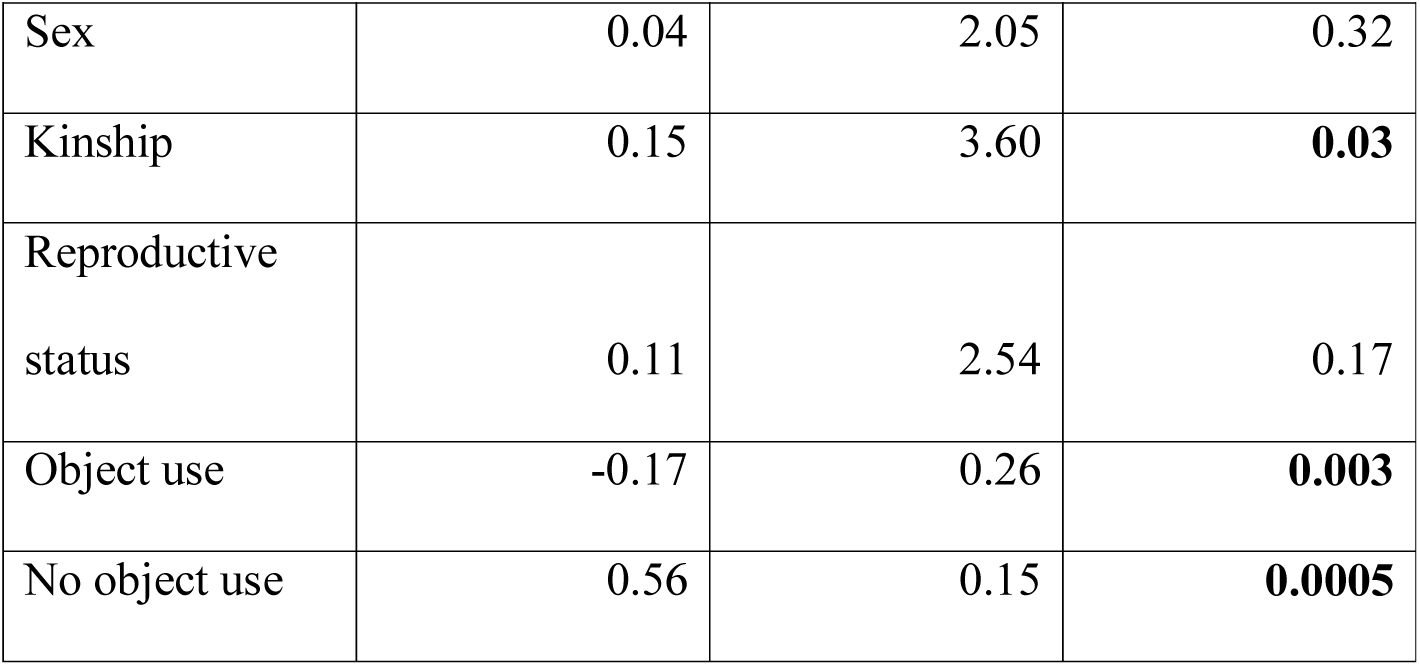
Duration of time in close proximity – within 2 m (*r^2^* = 0.37)

**Table S4.10.**
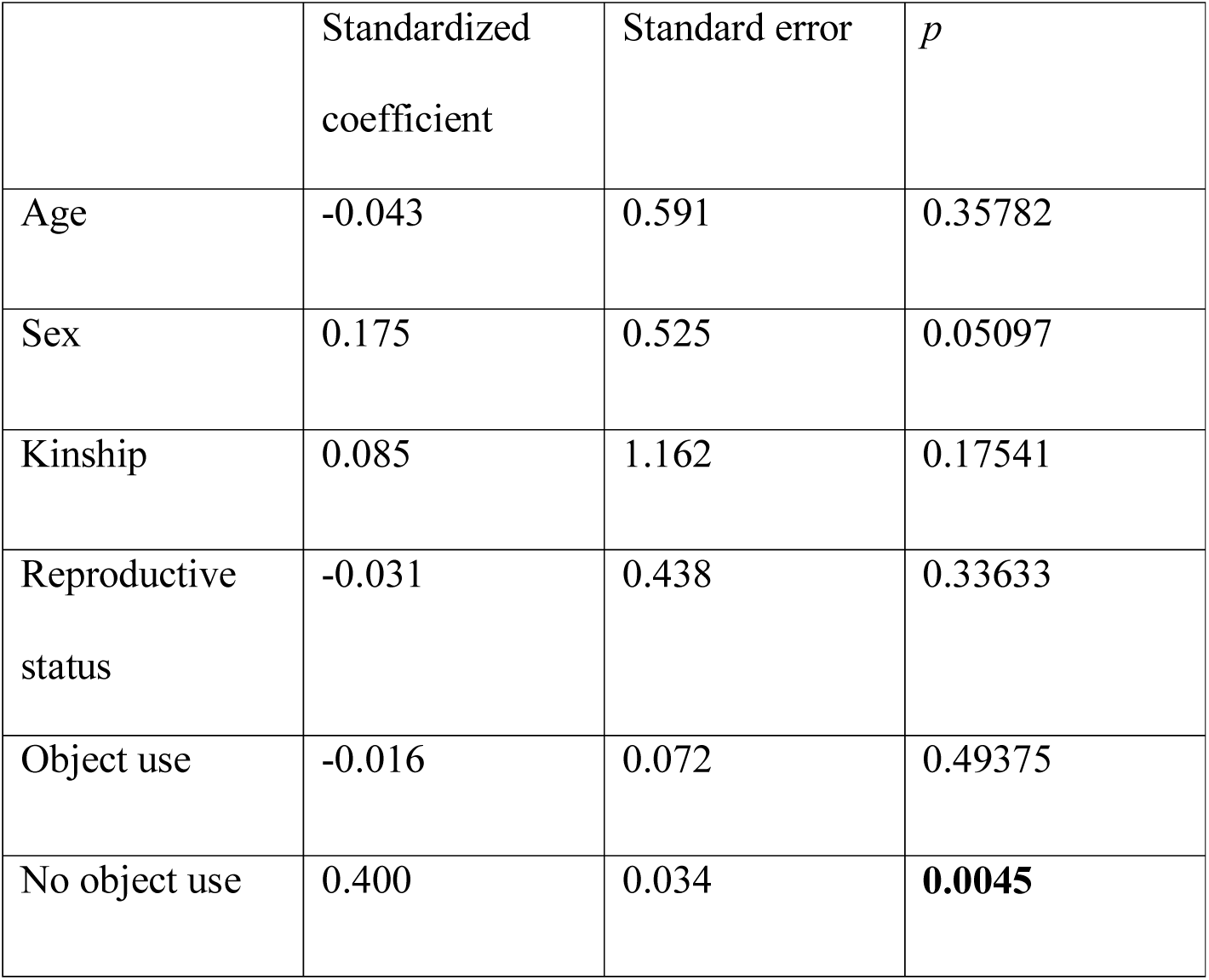
Rate of scratch produced (*r^2^* = 0.190)

**Table S4.11.** Rate of scratch received (*r^2^*= 0.050)

|  | Standardized coefficient | Standard error | *p* |
| --- | --- | --- | --- |
| Age | 0.03 | 0.63 | 0.38331 |
| Sex | 0.20 | 0.57 | **0.03998** |
| Kinship | 0.08 | 1.12 | 0.17841 |
| Reproductive status | 0.04 | 0.45 | 0.30585 |
| Object use | -0.01 | 0.08 | 0.56822 |
| No object use | -0.06 | 0.03 | 0.26087 |

#### Modality of gestural communication

Supplementary Table S5. MRQAP regression models predicting durations of social behavior, per hour dyad spent within 10m. Predictor variables were rates of gestural communication of different modalities (visual, tactile, auditory short-range, auditory long-range) and demographic variables. Based on 132 chimpanzee dyads. Significant *p* values are indicated in bold. R squared (*r^2^)* denotes amount of variance in the dependent variable explained by the regression model.

**Table S5.1.** Duration of joint feeding behaviour (*r^2^* = 0.17)

| | Standardized coefficient | Standard error | $p$ |
| --- | --- | --- | --- |
| Age | -0.07 | 0.68 | 0.27 |
| Sex | 0.33 | 0.70 | **0.002** |
| Kinship | 0.29 | 1.30 | **0.01** |
| Reproductive status | 0.21 | 0.71 | **0.03** |
| Visual gestures | 0.15 | 0.07 | 0.12 |
| Auditory long-range gestures | -0.08 | 0.13 | 0.21 |
| Auditory short-range gestures | 0.21 | 0.11 | **0.04** |
| Tactile gestures | -0.22 | 0.13 | **0.004** |

**Table S5.2.**
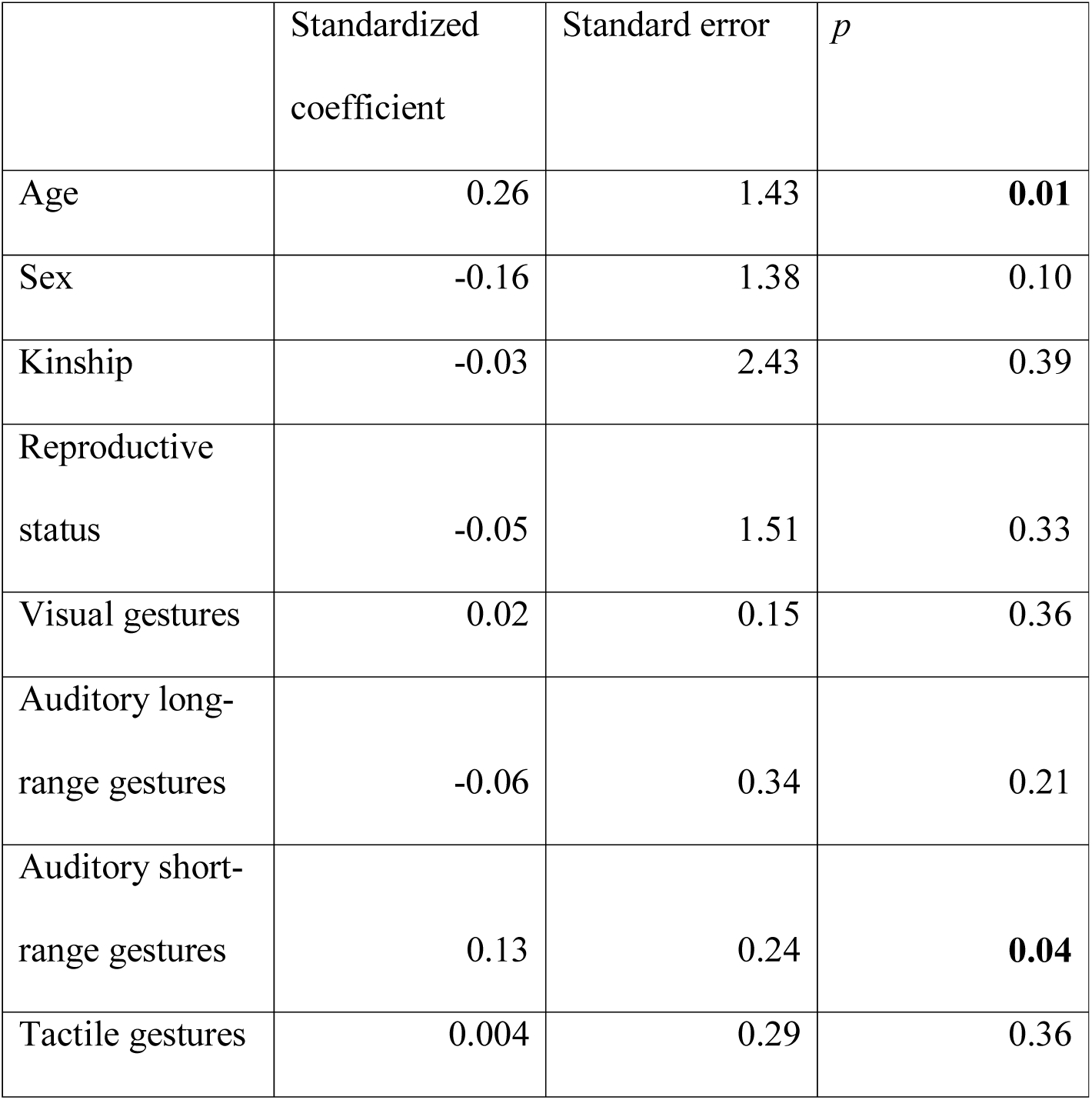
Duration of joint resting behaviour (*r^2^*= 0.09)

|  | Standardized coefficient | Standard error | *p* |
| --- | --- | --- | --- |
| Age | 0.26 | 1.43 | **0.01** |
| Sex | -0.16 | 1.38 | 0.10 |
| Kinship | -0.03 | 2.43 | 0.39 |
| Reproductive status | -0.05 | 1.51 | 0.33 |
| Visual gestures | 0.02 | 0.15 | 0.36 |
| Auditory long-range gestures | -0.06 | 0.34 | 0.21 |
| Auditory short-range gestures | 0.13 | 0.24 | **0.04** |
| Tactile gestures | 0.004 | 0.29 | 0.36 |

**Table S5.3.** Duration of joint travelling behaviour (*r^2^* = 0.32)

|  | Standardized coefficient | Standard error | *p* |
| --- | --- | --- | --- |
| Age | 0.21 | 0.33 | **0.01** |
| Sex | 0.04 | 0.30 | 0.36 |
| Kinship | 0.02 | 0.58 | 0.25 |
| Reproductive status | 0.00 | 0.30 | 0.50 |
| Visual gestures | 0.62 | 0.04 | **0.01** |
| Auditory long-range gestures | -0.28 | 0.08 | **0.01** |
| Auditory short-range gestures | 0.04 | 0.05 | 0.11 |
| Tactile gestures | -0.02 | 0.07 | 0.43 |

**Table S5.4.**
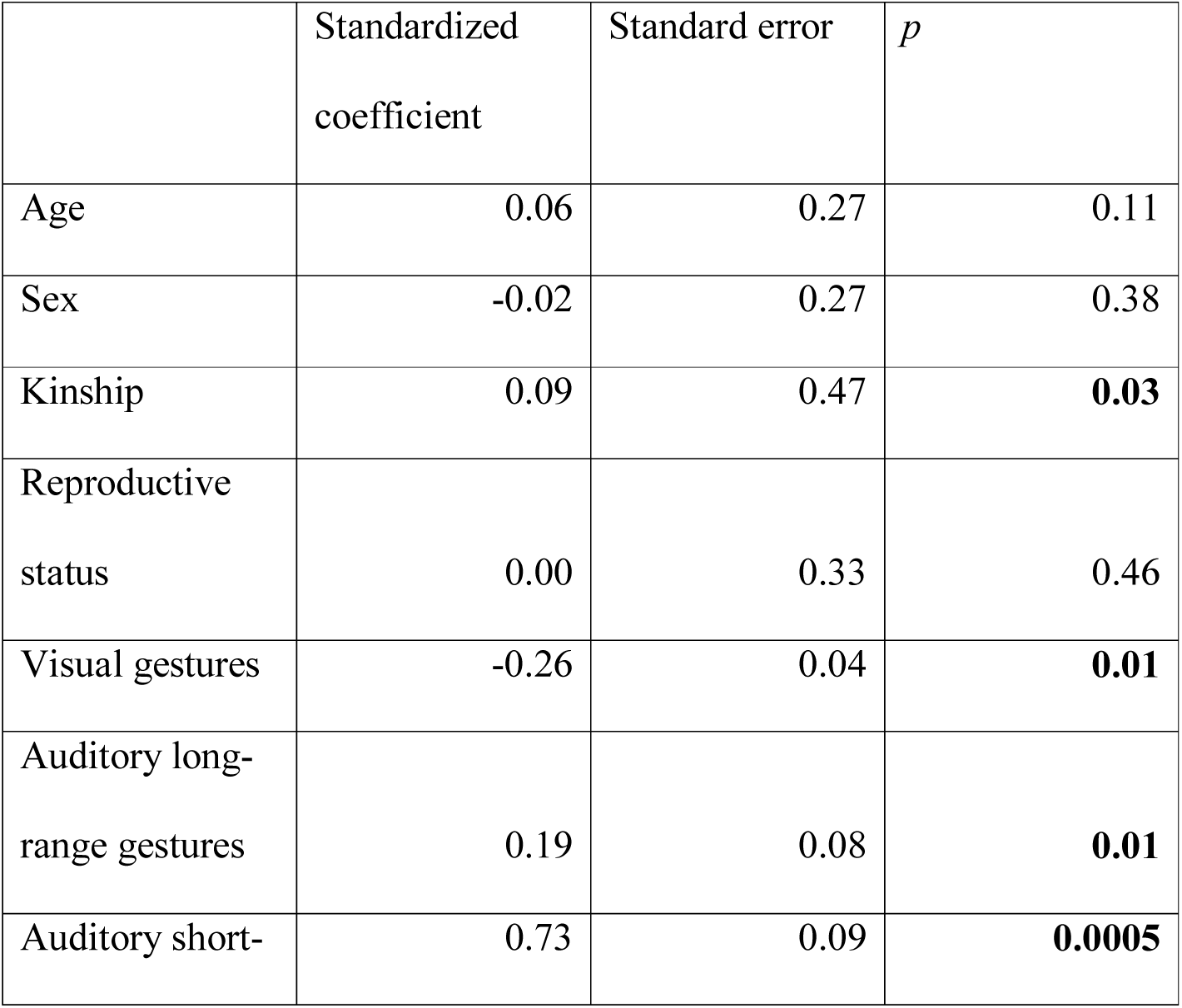
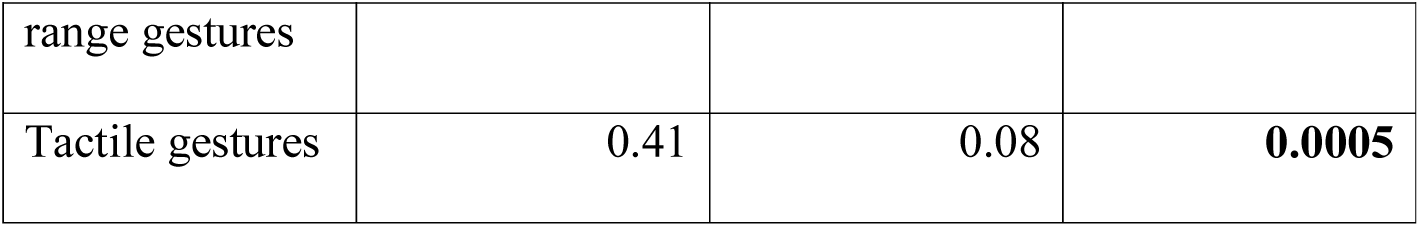
Duration of giving grooming (*r^2^*= 0.74)

**Table S5.5.** Duration of mutual grooming (*r^2^* = 0.48)

|  | Standardized<br>coefficient | Standard error | *p* |
| --- | --- | --- | --- |
| Age | 0.12 | 0.49 | 0.07 |
| Sex | -0.01 | 0.49 | 0.47 |
| Kinship | 0.03 | 0.87 | 0.27 |
| Reproductive<br>status | 0.12 | 0.53 | 0.08 |
| Visual gestures | 0.88 | 0.07 | **0.0005** |
| Auditory long-range gestures | -0.51 | 0.12 | **0.0005** |
| Auditory short-range gestures | 0.01 | 0.08 | 0.25 |
| Tactile gestures | -0.03 | 0.10 | 0.37 |

**Table S5.6.** Duration of receiving grooming (*r^2^*= 0.11)

|  | Standardized<br>coefficient | Standard error | *p* |
| --- | --- | --- | --- |
| Age | -0.15 | 0.50 | **0.04** |
| Sex | 0.21 | 0.49 | **0.02** |
| Kinship | 0.00 | 0.91 | 0.32 |
| Reproductive status | 0.07 | 0.56 | 0.24 |
| Visual gestures | 0.50 | 0.06 | **0.02** |
| Auditory long-range gestures | -0.27 | 0.13 | **0.02** |
| Auditory short-range gestures | 0.04 | 0.08 | 0.11 |
| Tactile gestures | -0.20 | 0.12 | **0.0005** |

**Table S5.7.** Duration of visual attention towards dyad partner (*r^2^*= 0.21)

|  | Standardized coefficient | Standard error | *p* |
| --- | --- | --- | --- |
| Age | -0.01 | 1.06 | 0.47 |
| Sex | 0.25 | 1.13 | **0.004** |
| Kinship | 0.19 | 1.97 | **0.007** |
| Reproductive status | 0.26 | 1.40 | **0.007** |
| Visual gestures | 0.75 | 0.15 | **0.0005** |
| Auditory long-range gestures | -0.44 | 0.25 | **0.0005** |
| Auditory short-range gestures | 0.24 | 0.19 | **0.01** |
| Tactile gestures | -0.02 | 0.22 | 0.50 |

**Table S5.8.** Duration of visual attention away from dyad partner (*r^2^* = 0.20)

|  | Standardized<br>coefficient | Standard error | *p* |
| --- | --- | --- | --- |
| Age | 0.296 | 1.683 | **0.004** |
| Sex | -0.083 | 1.347 | 0.178 |
| Kinship | 0.064 | 2.799 | 0.129 |
| Reproductive<br>status | 0.001 | 1.338 | 0.496 |
| Auditory long-range gestures | -0.032 | 0.368 | 0.412 |
| Auditory short-range gestures | 0.290 | 0.285 | **0.011** |
| Tactile gestures | 0.031 | 0.366 | 0.247 |
| Visual gestures | 0.033 | 0.179 | 0.327 |

**Table S5.9.** Duration of time in close proximity – within 2 m (*r^2^*= 0.40)

|  | Standardized<br>coefficient | Standard error | *p* |
| --- | --- | --- | --- |
| Age | 0.20 | 2.05 | **0.01** |
| Sex | 0.07 | 2.05 | 0.22 |
| Kinship | 0.15 | 3.65 | **0.03** |
| Reproductive<br>status | 0.12 | 2.59 | 0.15 |
| Visual gestures | 0.45 | 0.28 | **0.01** |
| Auditory long-range gestures | -0.27 | 0.48 | **0.01** |
| Auditory short-range gestures | 0.34 | 0.39 | **0.01** |
| Tactile gestures | 0.01 | 0.42 | 0.33 |

**Table S5.10.** Rate of scratch produced (*r^2^*= 0.201)

|  | Standardized coefficient | Standard error | *p* |
| --- | --- | --- | --- |
| Age | -0.064 | 0.601 | 0.27286 |
| Sex | 0.208 | 0.529 | **0.02699** |
| Kinship | 0.090 | 1.128 | 0.14893 |
| Reproductive status | -0.019 | 0.430 | 0.41979 |
| Visual gestures | 0.484 | 0.071 | **0.0085** |
| Auditory long-range gestures | -0.163 | 0.126 | **0.04698** |
| Auditory short-range gestures | 0.127 | 0.100 | 0.07496 |
| Tactile gestures | -0.072 | 0.120 | 0.2014 |

**Table S5.11.** Rate of scratch received (*r^2^* = 0.060)

|  | Standardized<br>coefficient | Standard error | *p* |
| --- | --- | --- | --- |
| Age | 0.03 | 0.65 | 0.34983 |
| Sex | 0.21 | 0.60 | **0.03448** |
| Kinship | 0.08 | 1.23 | 0.18041 |
| Reproductive<br>status | 0.04 | 0.45 | 0.29135 |
| Visual gestures | 0.08 | 0.07 | 0.22539 |
| Auditory long-range gestures | -0.08 | 0.14 | 0.23138 |
| Auditory short-range gestures | -0.10 | 0.10 | **0.04948** |
| Tactile gestures | -0.07 | 0.13 | 0.27186 |

#### Unimodal and multimodal gestural communication

Supplementary Table S6. MRQAP regression models predicting durations of social behavior, per hour dyad spent within 10m. Predictor variables were rates of unimodal and multi-modal gestural communication and demographic variables. Based on 132 chimpanzee dyads. Significant *p* values are indicated in bold. R squared (*r^2^)* denotes amount of variance in the dependent variable explained by the regression model.

**Table S6.1.** Duration of joint feeding behaviour (*r^2^*= 0.14)

|  | Standardized<br>coefficient | Standard error | *p* |
| --- | --- | --- | --- |
| Age | -0.01 | 0.69 | 0.47 |
| Sex | 0.31 | 0.68 | **0.002** |
| Kinship | 0.28 | 1.29 | **0.01** |
| Reproductive status | 0.20 | 0.73 | **0.04** |
| Unimodal gestures | 0.16 | 0.05 | 0.07 |
| Multimodal (facial<br>expressions) | -0.13 | 0.53 | 0.09 |
| Multimodal<br>(vocalisations) | -0.07 | 0.07 | 0.23 |

**Table S6.2.**
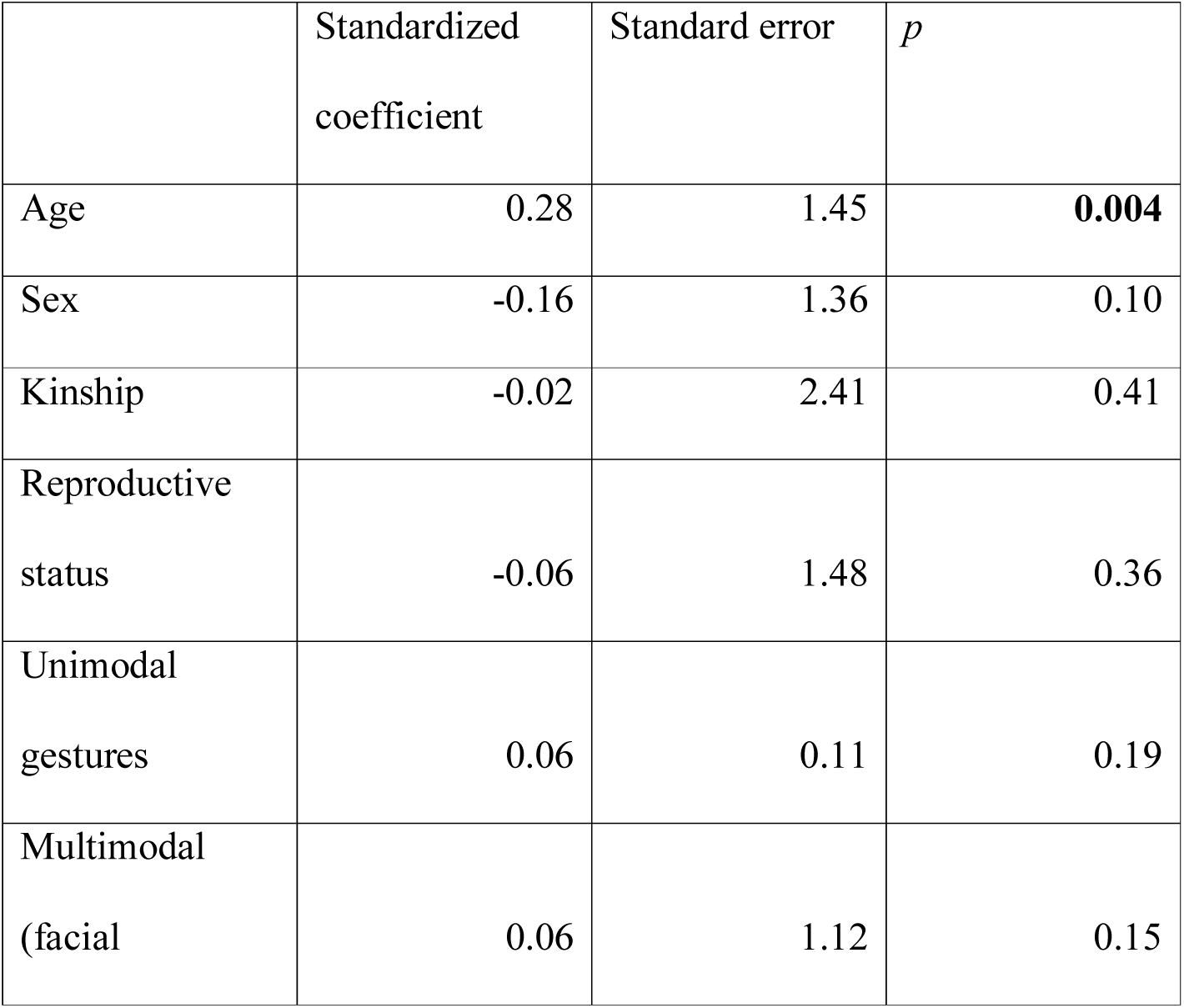
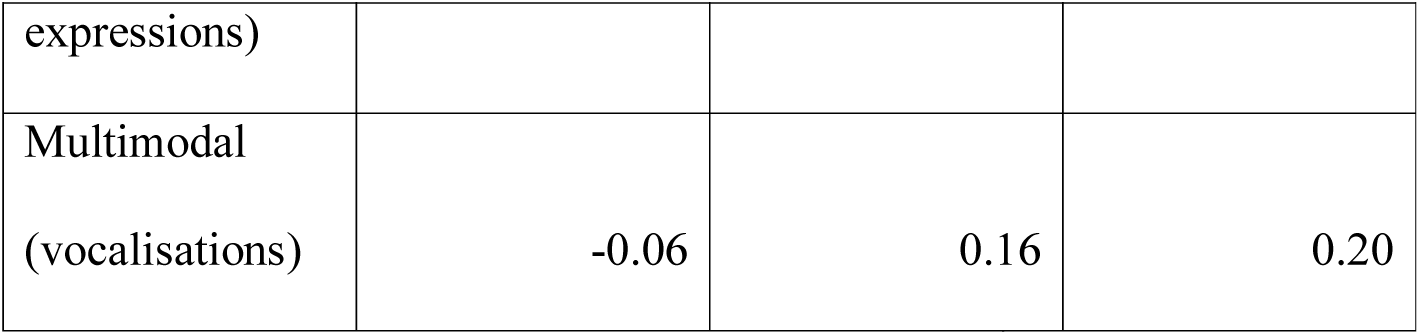
Duration of joint resting behaviour (*r^2^* = 0.08)

**Table S6.3.** Duration of joint travelling behaviour (*r^2^*= 0.31)

|  | Standardized<br>coefficient | Standard error | *p* |
| --- | --- | --- | --- |
| Age | 0.24 | 0.34 | **0.01** |
| Sex | -0.01 | 0.31 | 0.41 |
| Kinship | 0.02 | 0.62 | 0.25 |
| Reproductive<br>status | -0.02 | 0.32 | 0.43 |
| Unimodal<br>gestures | 0.22 | 0.03 | **0.02** |
| Multimodal<br>(facial<br>expressions) | 0.30 | 0.31 | **0.02** |
| Multimodal<br>(vocalisations) | 0.01 | 0.04 | 0.30 |

**Table S6.4.**
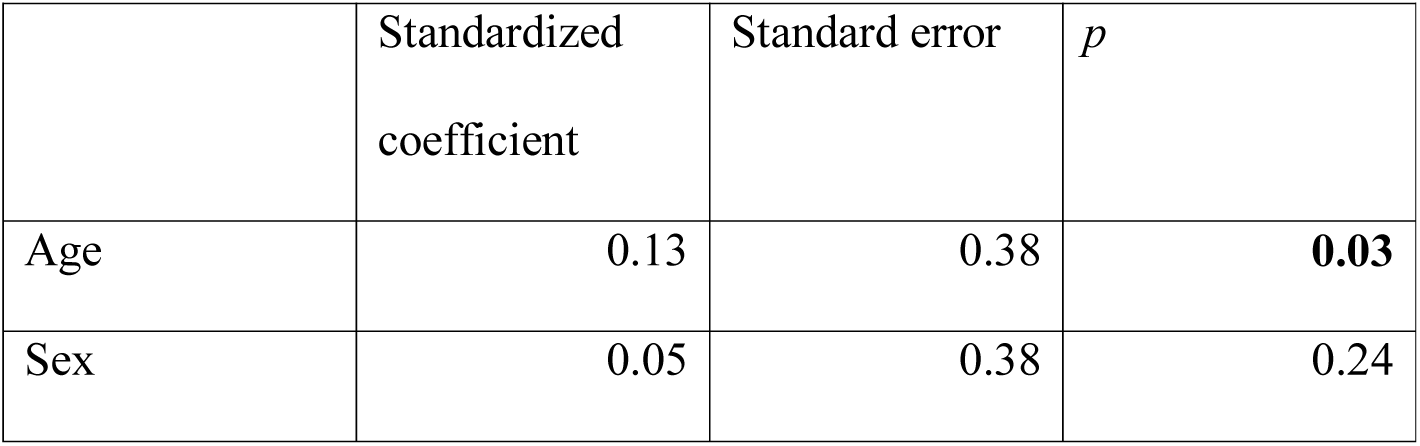
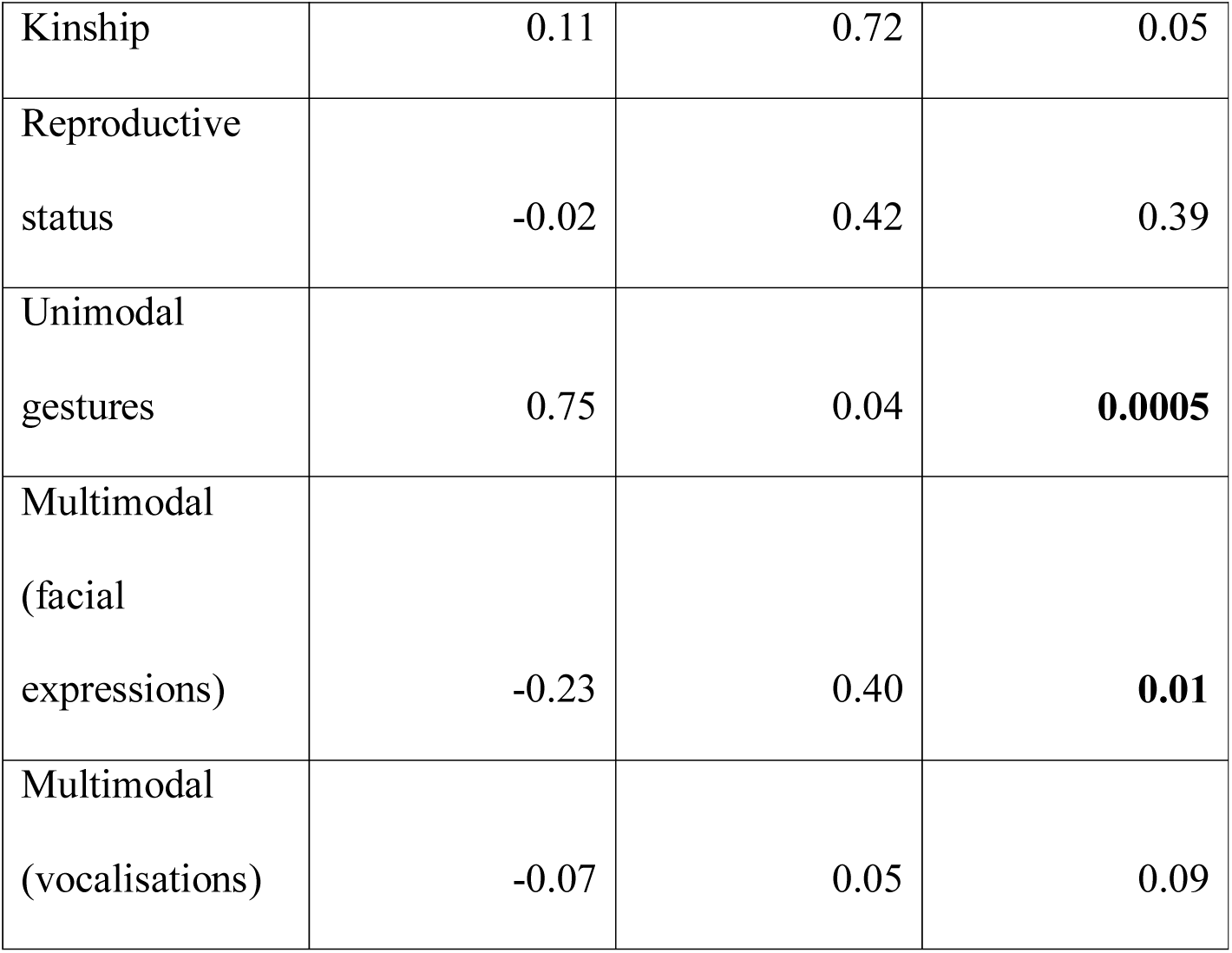
Duration of giving grooming (*r^2^* = 0.46)

|  | Standardized<br>coefficient | Standard error | *p* |
| --- | --- | --- | --- |
| Age | 0.13 | 0.38 | **0.03** |
| Sex | 0.05 | 0.38 | 0.24 |
| Kinship | 0.11 | 0.72 | 0.05 |
| Reproductive status | -0.02 | 0.42 | 0.39 |
| Unimodal gestures | 0.75 | 0.04 | **0.0005** |
| Multimodal (facial expressions) | -0.23 | 0.40 | **0.01** |
| Multimodal (vocalisations) | -0.07 | 0.05 | 0.09 |

**Table S6.5.**
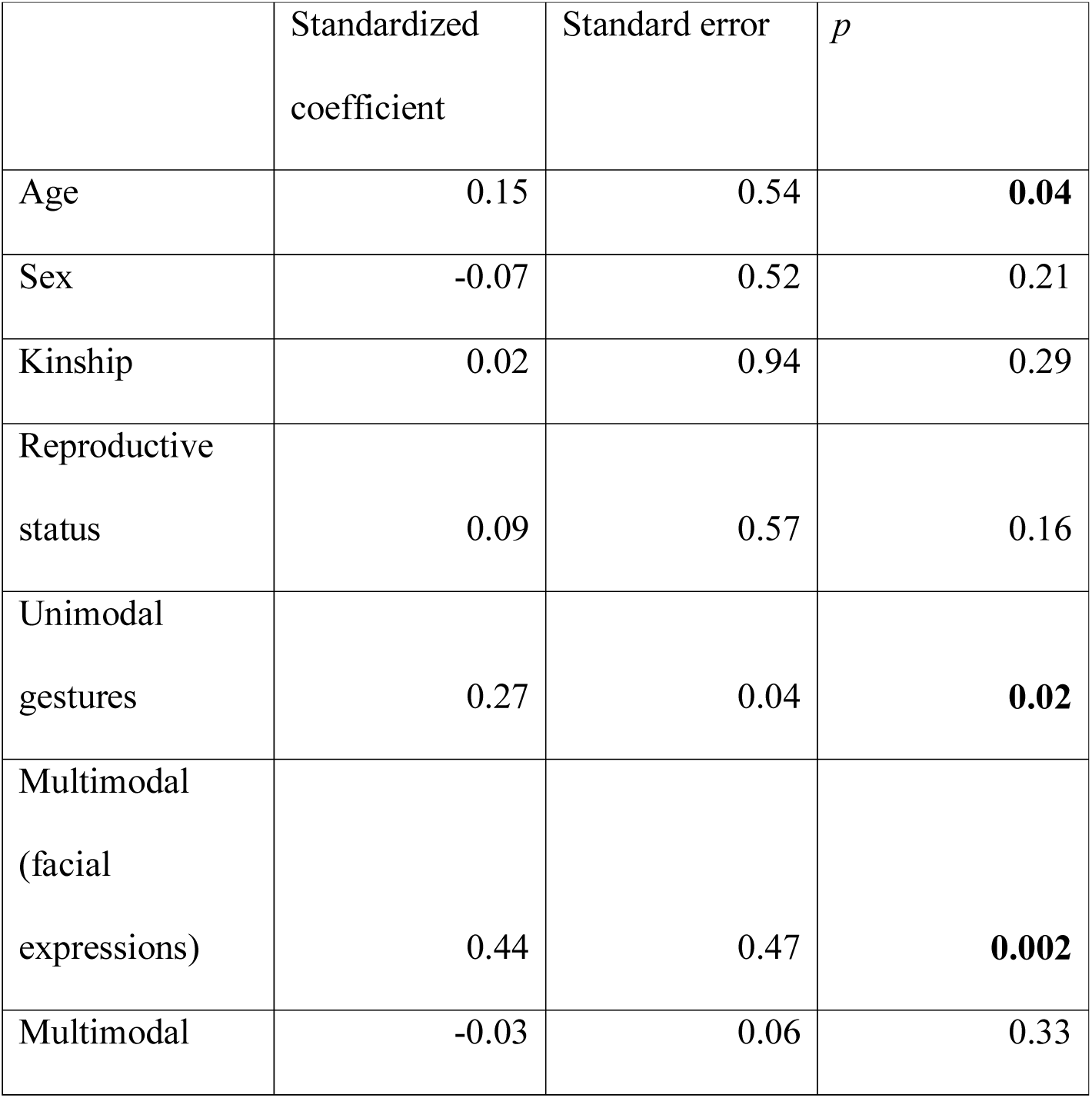
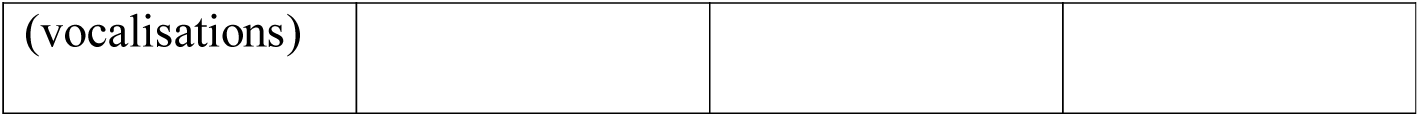
Duration of mutual grooming (*r^2^*= 0.42)

**Table S6.6.** Duration of receiving grooming (*r^2^* = 0.08)

|  | Standardized<br>coefficient | Standard error | *p* |
| --- | --- | --- | --- |
| Age | -0.11 | 0.52 | 0.12 |
| Sex | 0.17 | 0.52 | 0.06 |
| Kinship | 0.00 | 0.93 | 0.65 |
| Reproductive<br>status | 0.04 | 0.61 | 0.36 |
| Unimodal<br>gestures | 0.25 | 0.04 | **0.04** |
| Multimodal<br>(facial<br>expressions) | 0.001 | 0.44 | 0.43 |
| Multimodal<br>(vocalisations) | -0.09 | 0.05 | 0.06 |

**Table S6.7.**
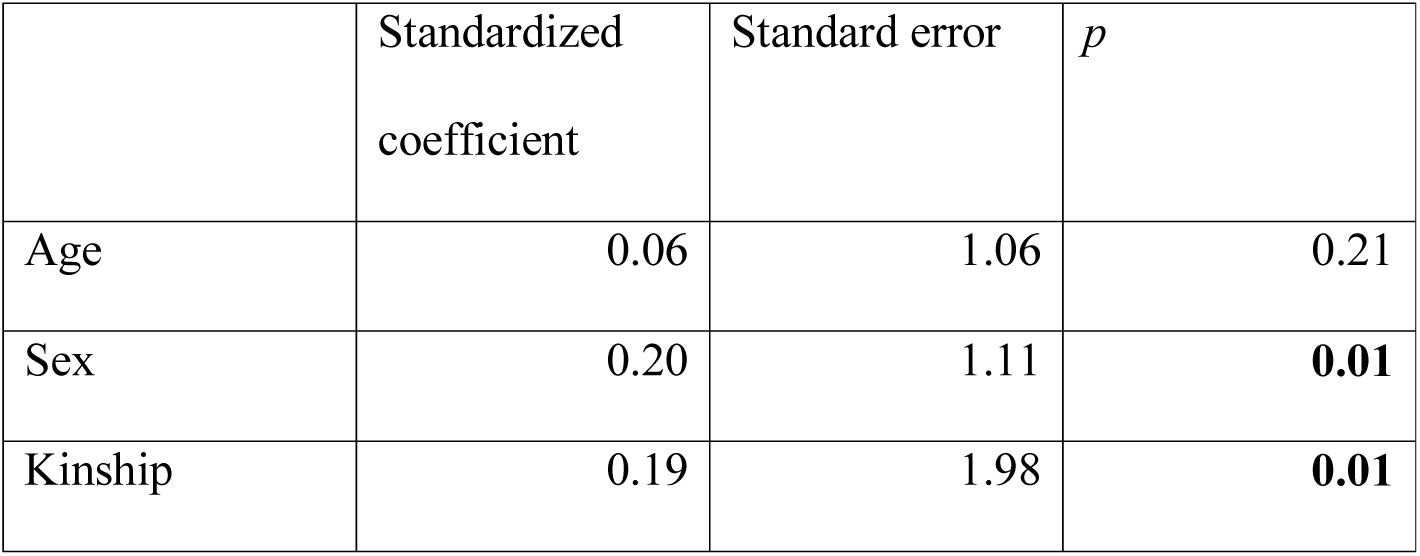
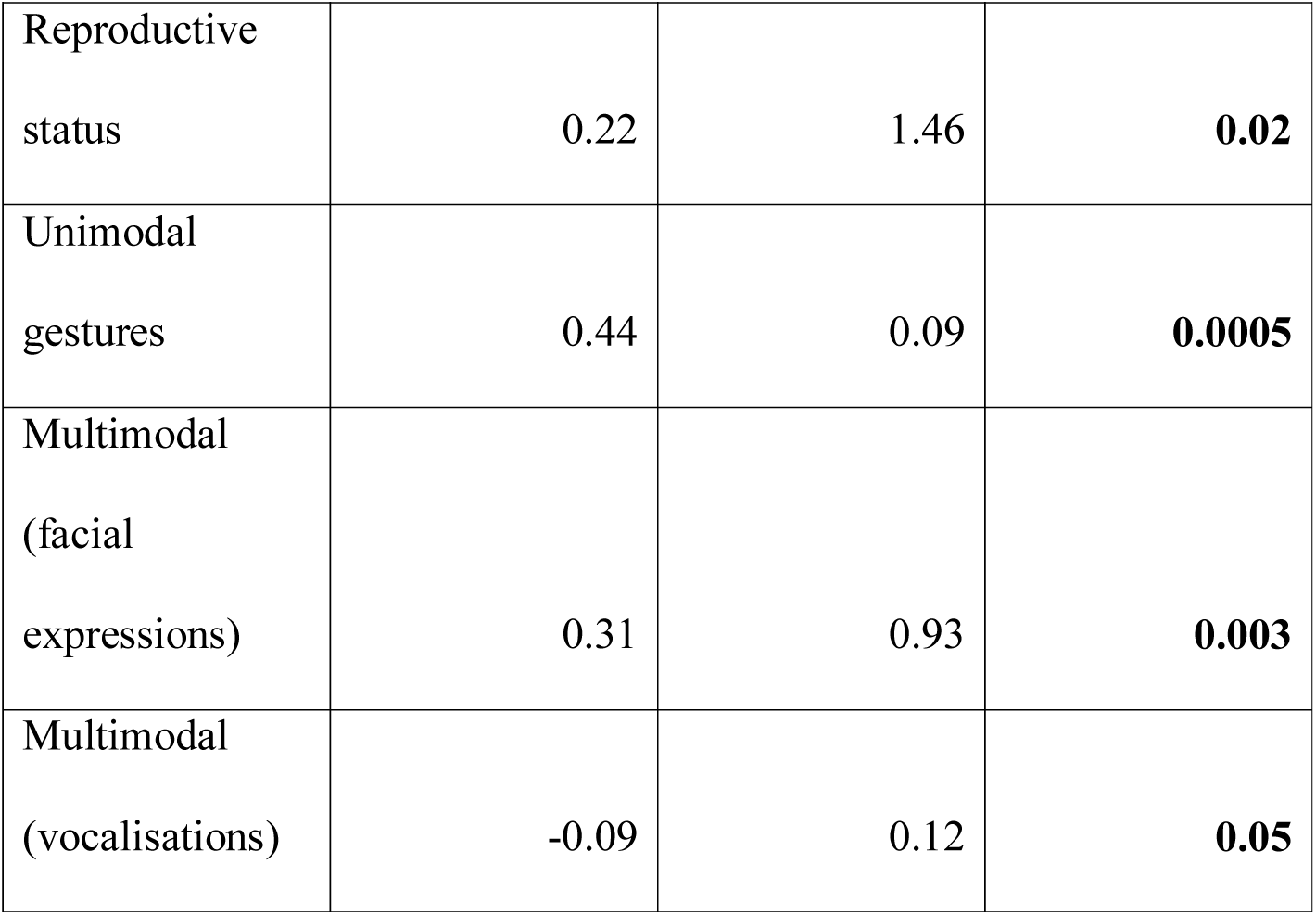
Duration of visual attention towards dyad partner (*r^2^* = 0.50)

|  | Standardized<br>coefficient | Standard error | *p* |
| --- | --- | --- | --- |
| Age | 0.06 | 1.06 | 0.21 |
| Sex | 0.20 | 1.11 | **0.01** |
| Kinship | 0.19 | 1.98 | **0.01** |
| Reproductive status | 0.22 | 1.46 | **0.02** |
| Unimodal gestures | 0.44 | 0.09 | **0.0005** |
| Multimodal (facial expressions) | 0.31 | 0.93 | **0.003** |
| Multimodal (vocalisations) | -0.09 | 0.12 | **0.05** |

**Table S6.8.**
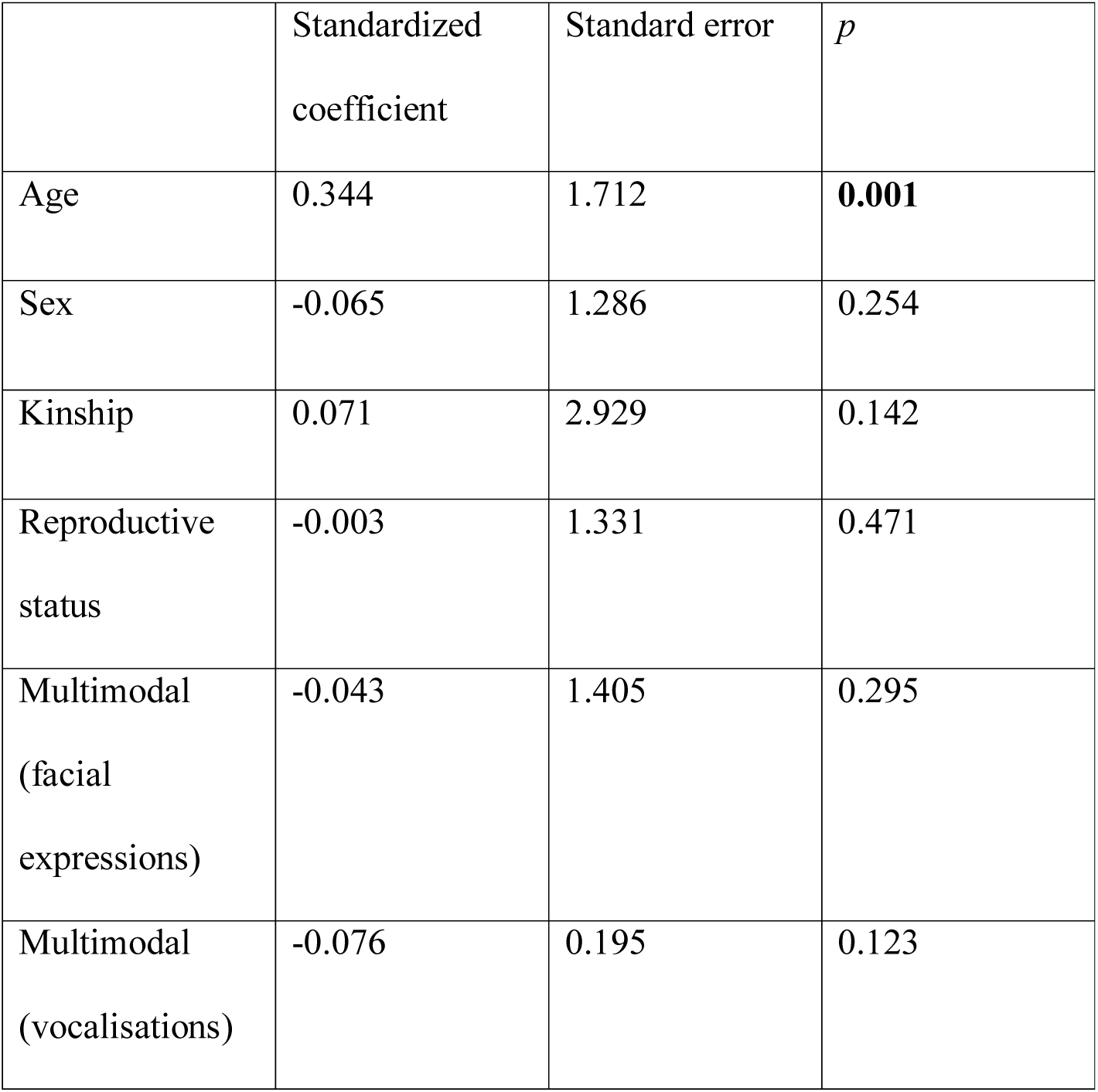
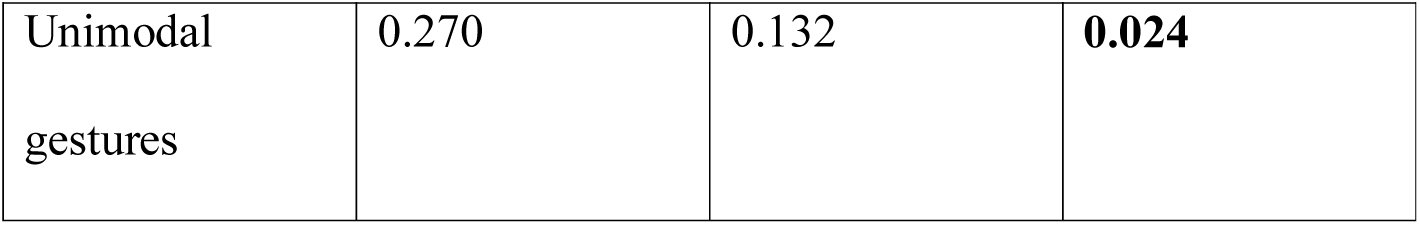
Duration of visual attention away from dyad partner (*r^2^*= 0.17)

|  | Standardized coefficient | Standard error | *p* |
| --- | --- | --- | --- |
| Age | 0.344 | 1.712 | **0.001** |
| Sex | -0.065 | 1.286 | 0.254 |
| Kinship | 0.071 | 2.929 | 0.142 |
| Reproductive status | -0.003 | 1.331 | 0.471 |
| Multimodal (facial expressions) | -0.043 | 1.405 | 0.295 |
| Multimodal (vocalisations) | -0.076 | 0.195 | 0.123 |
| Unimodal gestures | 0.270 | 0.132 | **0.024** |

**Table S6.9.** Duration of time in close proximity – within 2 m (*r^2^* = 0.37)

|  | Standardized coefficient | Standard error | *p* |
| --- | --- | --- | --- |
| Age | 0.27 | 2.07 | **0.002** |
| Sex | 0.05 | 2.01 | 0.31 |
| Kinship | 0.15 | 3.55 | **0.03** |
| Reproductive status | 0.09 | 2.54 | 0.22 |
| Unimodal gestures | 0.44 | 0.17 | **0.001** |
| Multimodal (facial expressions) | 0.14 | 1.69 | **0.05** |
| Multimodal (vocalisations) | -0.11 | 0.23 | **0.04** |

**Table S6.10.** Rate of scratch produced (*r^2^* = 0.191)

|  | Standardized coefficient | Standard error | *p* |
| --- | --- | --- | --- |
| Age | -0.029 | 0.593 | 0.37781 |
| Sex | 0.171 | 0.536 | 0.05847 |
| Kinship | 0.089 | 1.098 | 0.14393 |
| Reproductive<br>status | -0.018 | 0.448 | 0.43278 |
| Unimodal<br>gestures | 0.180 | 0.045 | **0.04948** |
| Multimodal<br>(facial<br>expressions) | 0.184 | 0.489 | **0.04298** |
| Multimodal<br>(vocalisations) | 0.138 | 0.065 | 0.06597 |

**Table S6.11.**
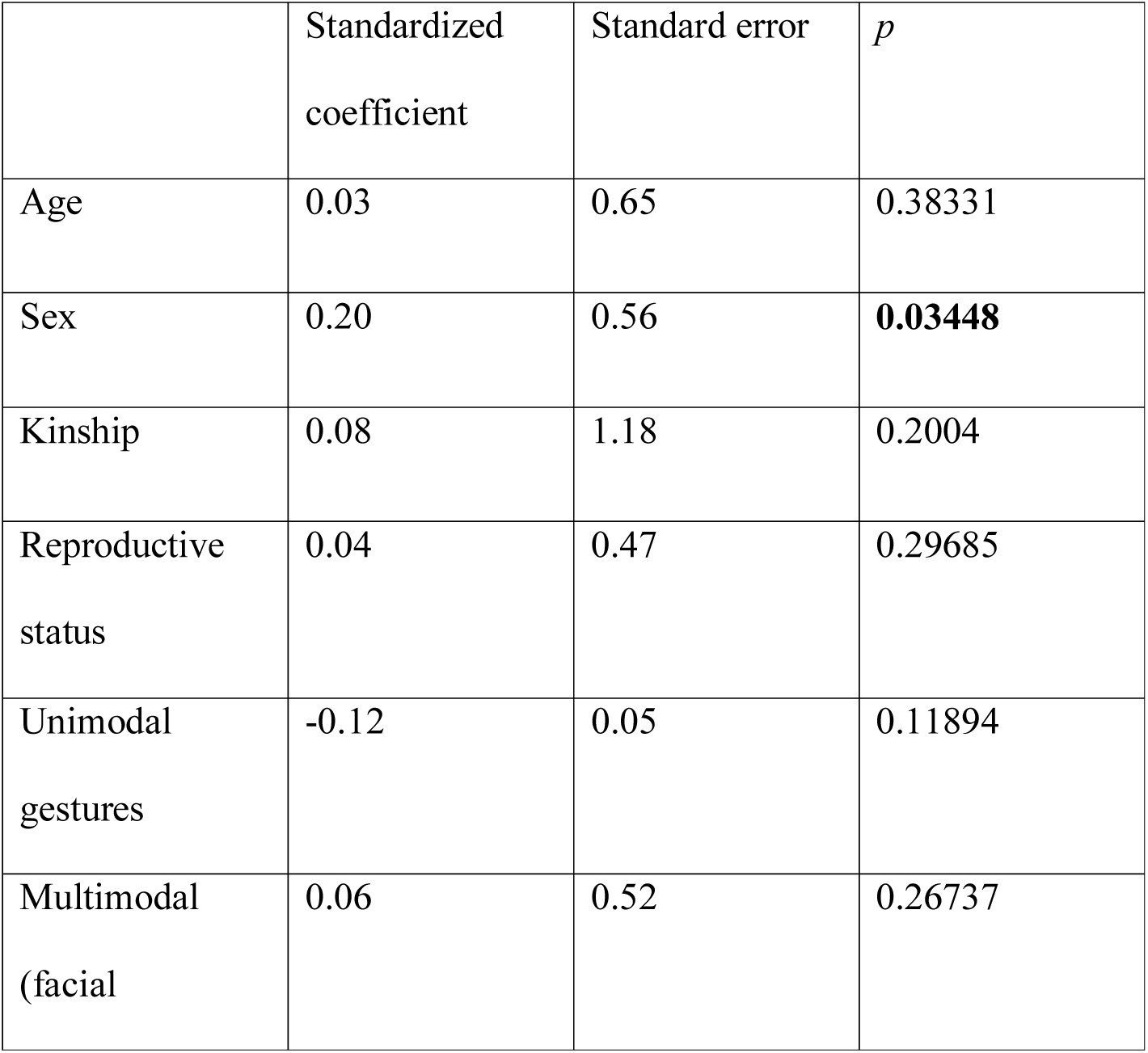
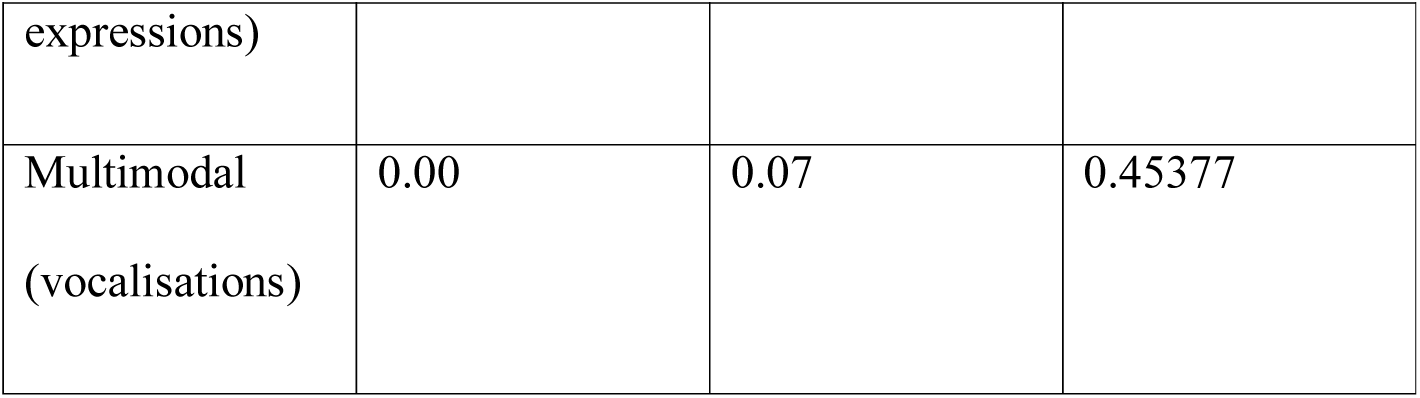
Rate of scratch received (*r^2^*= 0.056)

#### Dyadic repertoire size

Supplementary Table S7. MRQAP regression models predicting durations of social behavior, per hour dyad spent within 10m. Predictor variables were dyadic repertoire size, per hour dyad spent within 10m and demographic variables. Based on 132 chimpanzee dyads. Significant *p* values are indicated in bold. R squared (*r^2^)* denotes amount of variance in the dependent variable explained by the regression model.

**Table S7.1.** Duration of joint feeding behaviour (*r^2^* = 0.12)

|  | Standardized<br>coefficient | Standard error | *p* |
| --- | --- | --- | --- |
| Age | -0.01 | 0.65 | 0.49 |
| Sex | 0.31 | 0.68 | **0.0005** |
| Kinship | 0.28 | 1.32 | **0.01** |
| Reproductive status | 0.22 | 0.75 | **0.04** |
| Dyadic repertoire<br>size | -0.02 | 0.04 | 0.45 |

**Table S7.2.**
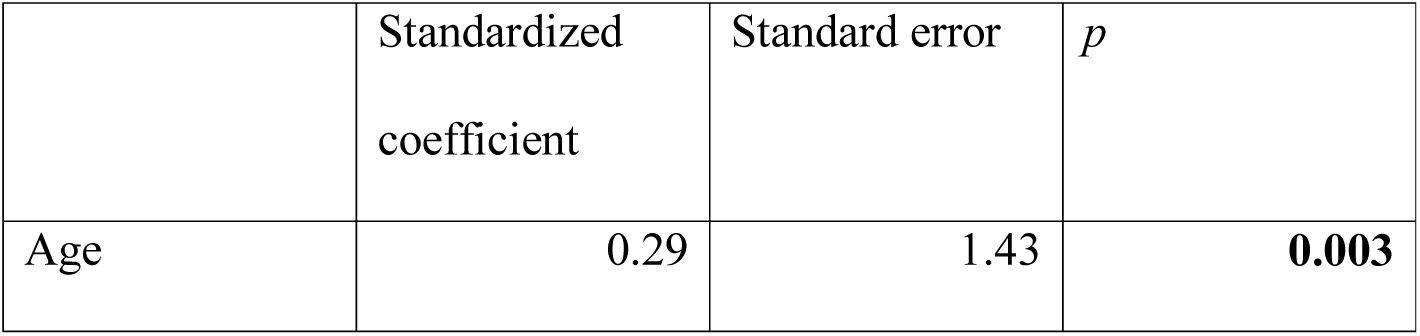
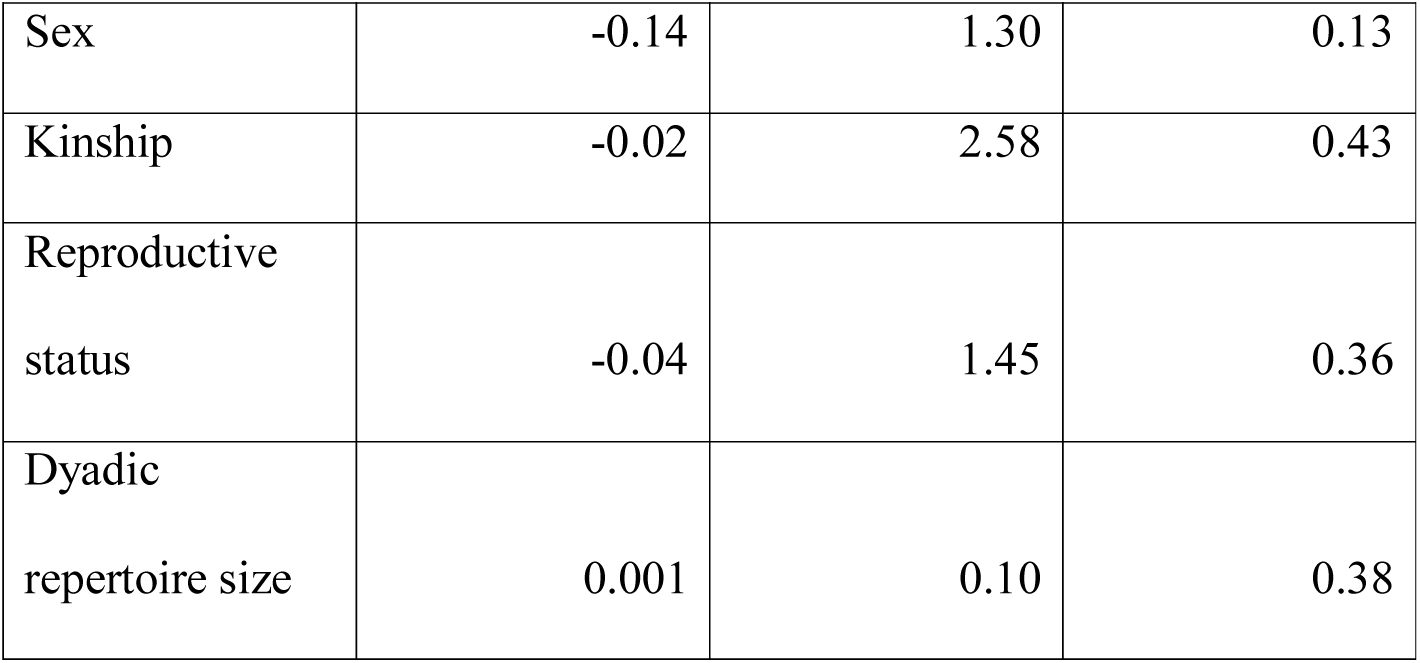
Duration of joint resting behaviour (*r^2^*= 0.07)

**Table S7.3.** Duration of joint travelling behaviour (*r^2^* = 0.21)

|  | Standardized coefficient | Standard error | *p* |
| --- | --- | --- | --- |
| Age | 0.25 | 0.38 | **0.01** |
| Sex | 0.05 | 0.34 | 0.36 |
| Kinship | 0.03 | 0.66 | 0.24 |
| Reproductive status | 0.02 | 0.36 | 0.41 |
| Dyadic repertoire size | 0.34 | 0.03 | **0.01** |

**Table S7.4.**
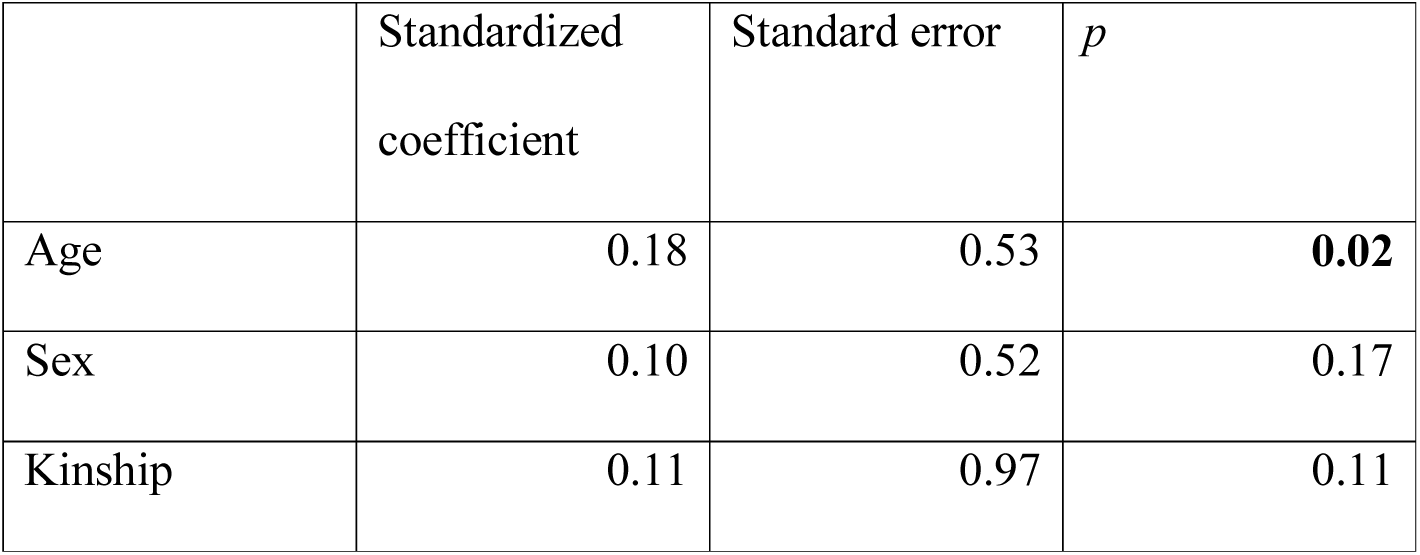
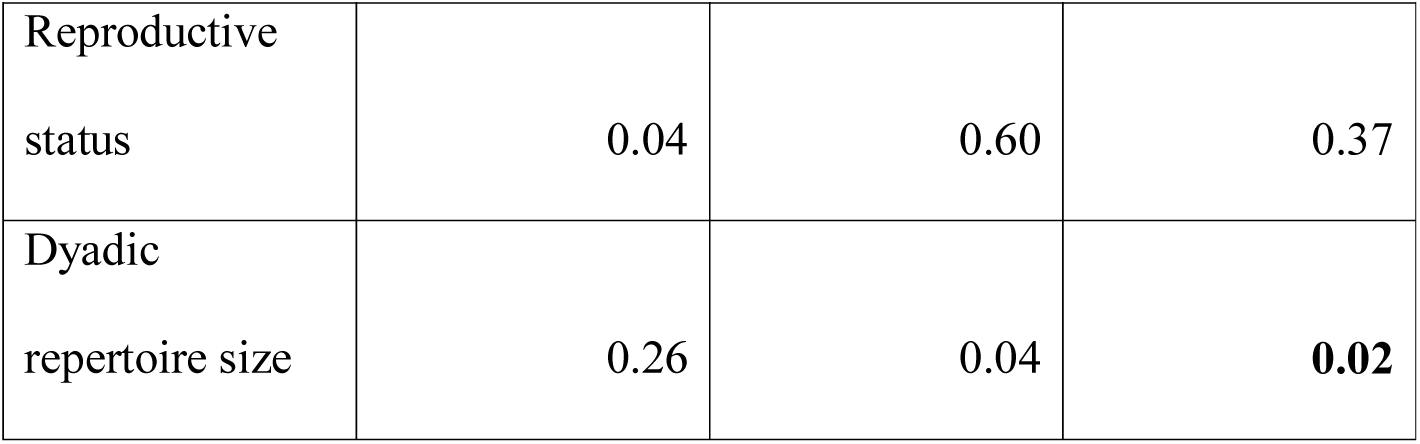
Duration of giving grooming (*r^2^*= 0.13)

**Table S7.5.** Duration of mutual grooming (*r^2^* = 0.24)

|  | Standardized coefficient | Standard error | *p* |
| --- | --- | --- | --- |
| Age | 0.16 | 0.60 | **0.05** |
| Sex | 0.01 | 0.59 | 0.51 |
| Kinship | 0.04 | 1.11 | 0.22 |
| Reproductive status | 0.15 | 0.68 | 0.11 |
| Dyadic repertoire size | 0.45 | 0.05 | **0.003** |

**Table S7.6.** Duration of receiving grooming (*r^2^*= 0.04)

|  | Standardized coefficient | Standard error | *p* |
| --- | --- | --- | --- |
| Age | -0.10 | 0.49 | 0.13 |
| Sex | 0.20 | 0.50 | **0.03** |
| Kinship | 0.00 | 0.87 | 0.33 |
| Reproductive status | 0.08 | 0.60 | 0.24 |
| Dyadic repertoire size | 0.14 | 0.04 | 0.08 |

**Table S7.7.** Duration of visual attention towards dyad partner (*r^2^*= 0.28)

|  | Standardized coefficient | Standard error | *p* |
| --- | --- | --- | --- |
| Age | 0.08 | 1.24 | 0.16 |
| Sex | 0.29 | 1.38 | **0.002** |
| Kinship | 0.21 | 2.50 | **0.02** |
| Reproductive status | 0.29 | 1.87 | **0.02** |
| Dyadic repertoire size | 0.42 | 0.11 | **0.002** |

**Table S7.8.** Duration of visual attention away from dyad partner (*r^2^* = 0.12)

|  | Standardized coefficient | Standard error | *p* |
| --- | --- | --- | --- |
| Age | 0.361 | 1.687 | **0.001** |
| Sex | -0.065 | 1.276 | 0.255 |
| Kinship | 0.074 | 2.826 | 0.129 |
| Reproductive status | 0.036 | 1.371 | 0.370 |
| Dyadic repertoire size | 0.063 | 0.119 | 0.164 |

**Table S7.9.** Duration of time in close proximity – within 2 m (*r^2^*= 0.21)

|  | Standardized<br>coefficient | Standard error | *p* |
| --- | --- | --- | --- |
| Age | 0.29 | 2.38 | **0.001** |
| Sex | 0.11 | 2.27 | 0.15 |
| Kinship | 0.16 | 4.12 | **0.03** |
| Reproductive<br>status | 0.15 | 3.25 | 0.14 |
| Dyadic<br>repertoire size | 0.28 | 0.20 | **0.01** |

**Table S7.10.** Rate of scratch produced (*r^2^*= 0.167)

|  | Standardized<br>coefficient | Standard error | *p* |
| --- | --- | --- | --- |
| Age | -0.016 | 0.583 | 0.43878 |
| Sex | 0.193 | 0.529 | **0.03398** |
| Kinship | 0.096 | 1.076 | 0.13343 |
| Reproductive<br>status | 0.008 | 0.417 | 0.45527 |
| Dyadic<br>repertoire size | 0.352 | 0.042 | **0.002** |

**Table S7.11.** Rate of scratch received (*r^2^* = 0.048)

|  | Standardized<br>coefficient | Standard error | *p* |
| --- | --- | --- | --- |
| Age | 0.02 | 0.62 | 0.42829 |
| Sex | 0.20 | 0.55 | **0.03698** |
| Kinship | 0.08 | 1.21 | 0.21039 |
| Reproductive<br>status | 0.03 | 0.44 | 0.38131 |
| Dyadic<br>repertoire size | -0.04 | 0.04 | 0.33483 |

#### Single and combined gestures

Supplementary Table S8. MRQAP regression models predicting durations of social behavior, per hour dyad spent within 10m. Predictor variables were rates of single and combined gestures, per hour dyad spent within 10m and demographic variables. Based on 132 chimpanzee dyads. Significant *p* values are indicated in bold. R squared (*r^2^)* denotes amount of variance in the dependent variable explained by the regression model.

**Table S8.1.** Duration of joint feeding behaviour (*r^2^*= 0.12)

|  | Standardized<br>coefficient | Standard error | *p* |
| --- | --- | --- | --- |
| Age | -0.01 | 0.71 | 0.47 |
| Sex | 0.31 | 0.68 | **0.001** |
| Kinship | 0.29 | 1.34 | **0.01** |
| Reproductive status | 0.22 | 0.77 | **0.048** |
| Combined gesture | 0.01 | 0.16 | 0.43 |
| Single gesture | 0.02 | 0.04 | 0.42 |

**Table S8.2.** Duration of joint resting behaviour (*r^2^* = 0.07)

|  | Standardized<br>coefficient | Standard error | *p* |
| --- | --- | --- | --- |
| Age | 0.28 | 1.38 | **0.002** |
| Sex | -0.15 | 1.32 | 0.11 |
| Kinship | -0.02 | 2.50 | 0.46 |
| Reproductive<br>status | -0.04 | 1.47 | 0.37 |
| Combined<br>gesture | -0.03 | 0.34 | 0.37 |
| Single gesture | 0.06 | 0.10 | 0.17 |

**Table S8.3.** Duration of joint travelling behaviour (*r^2^*= 0.31)

|  | Standardized<br>coefficient | Standard error | *p* |
| --- | --- | --- | --- |
| Age | 0.23 | 0.35 | **0.01** |
| Sex | 0.05 | 0.32 | 0.31 |
| Kinship | 0.03 | 0.64 | 0.24 |
| Reproductive<br>status | -0.01 | 0.32 | 0.44 |
| Combined<br>gesture | 0.41 | 0.09 | **0.004** |
| Single gesture | 0.09 | 0.02 | 0.12 |

**Table S8.4.**
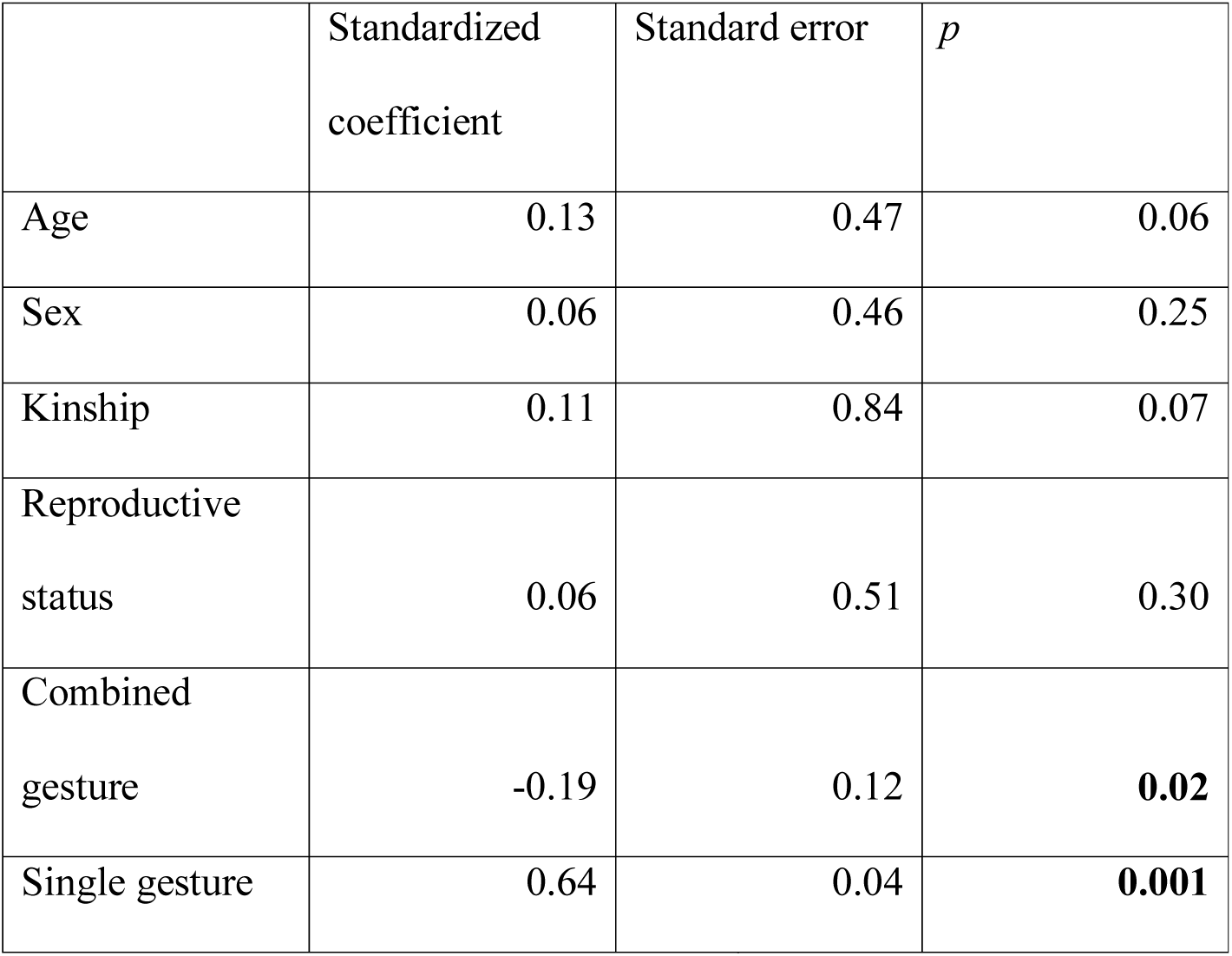
Duration of giving grooming (*r^2^* = 0.35)

**Table S8.5.** Duration of mutual grooming (*r^2^*= 0.32)

|  | Standardized<br>coefficient | Standard error | *p* |
| --- | --- | --- | --- |
| Age | 0.14 | 0.57 | 0.07 |
| Sex | 0.00 | 0.53 | 0.52 |
| Kinship | 0.04 | 1.04 | 0.23 |
| Reproductive<br>status | 0.11 | 0.61 | 0.13 |
| Combined<br>gesture | 0.40 | 0.16 | **0.003** |
| Single gesture | 0.19 | 0.04 | 0.05 |

**Table S8.6.**
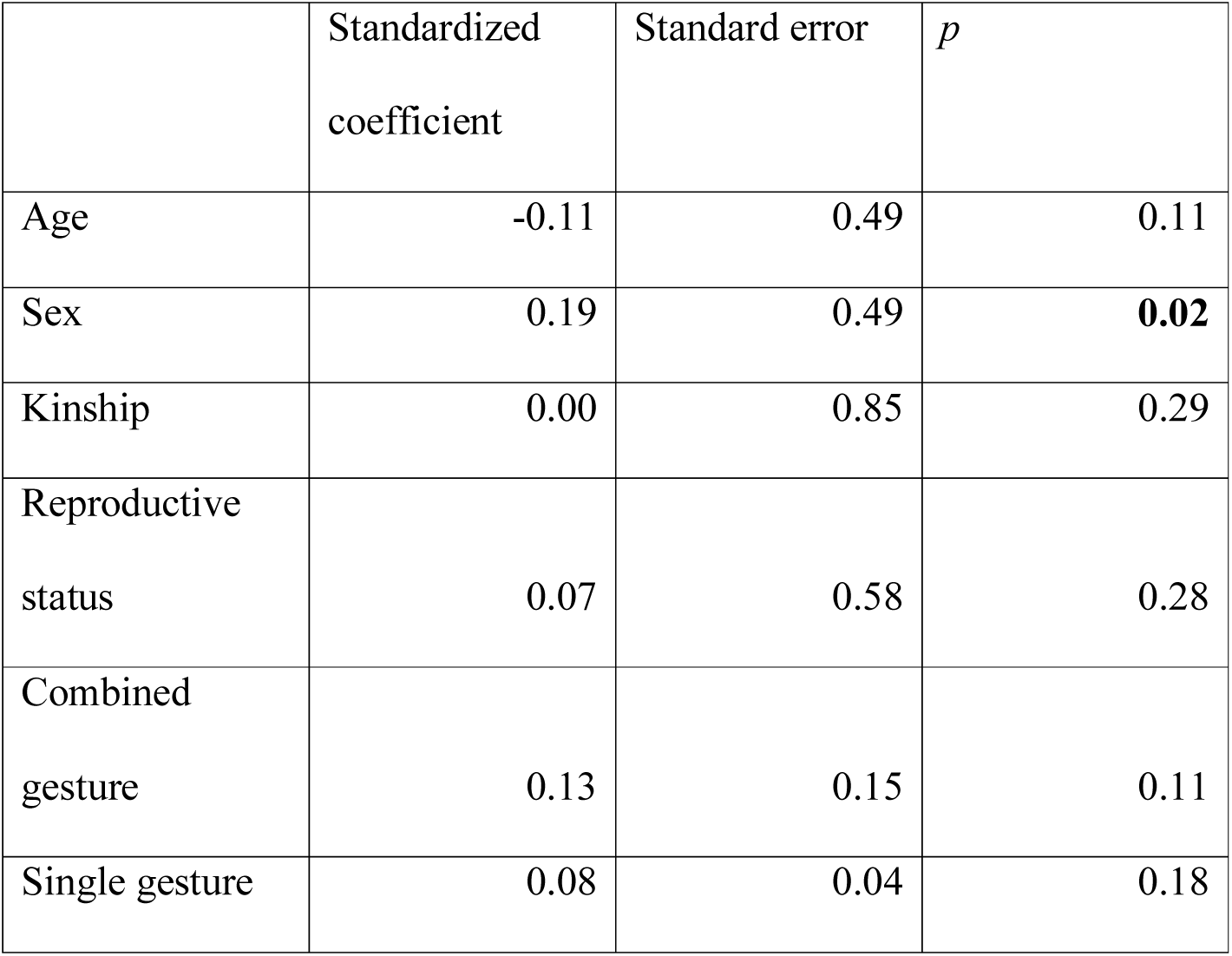
Duration of receiving grooming (*r^2^* = 0.06)

|  | Standardized<br>coefficient | Standard error | *p* |
| --- | --- | --- | --- |
| Age | -0.11 | 0.49 | 0.11 |
| Sex | 0.19 | 0.49 | **0.02** |
| Kinship | 0.00 | 0.85 | 0.29 |
| Reproductive<br>status | 0.07 | 0.58 | 0.28 |
| Combined<br>gesture | 0.13 | 0.15 | 0.11 |
| Single gesture | 0.08 | 0.04 | 0.18 |

**Table S8.7.**
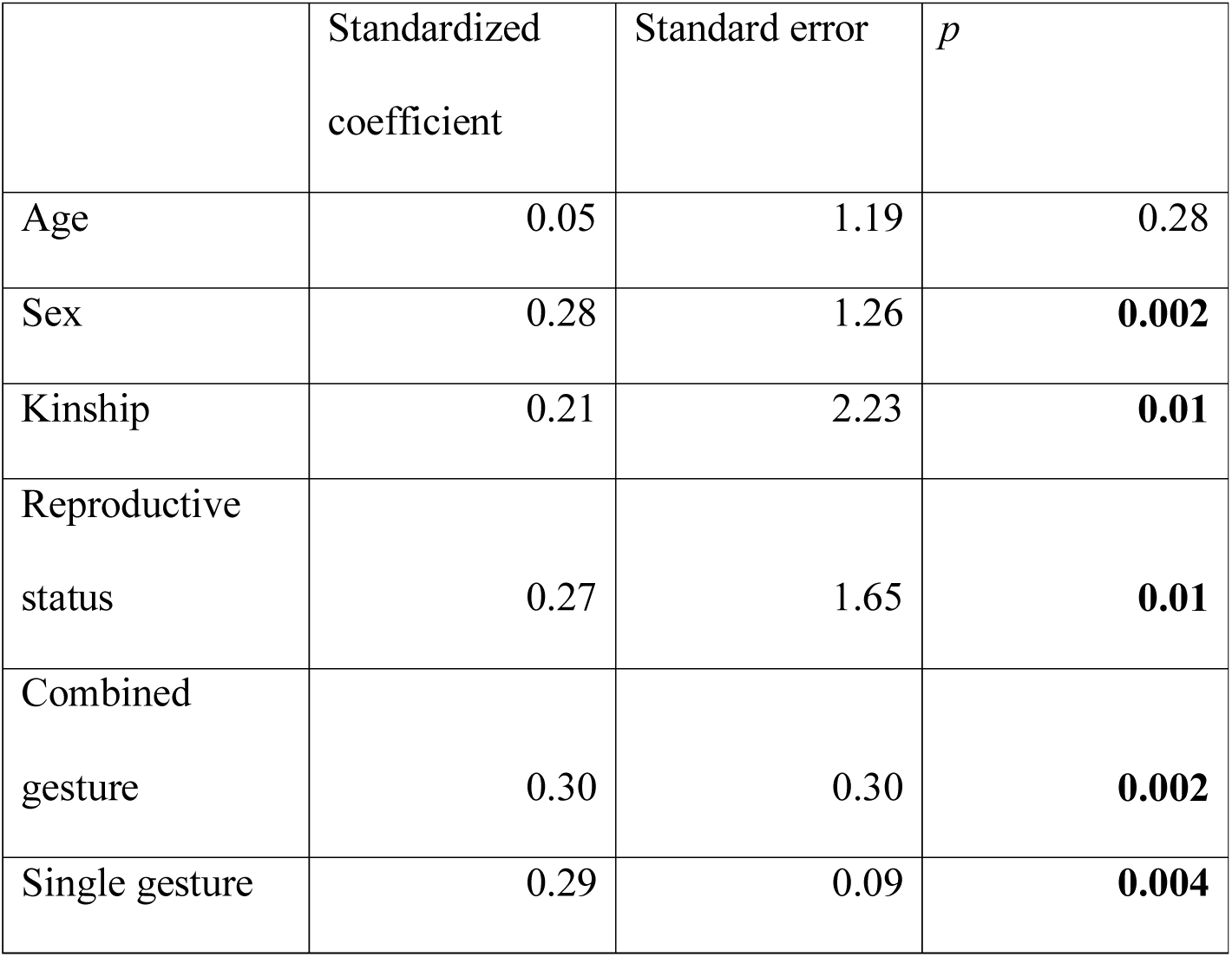
Duration of visual attention towards dyad partner (*r^2^* = 0.38)

|  | Standardized<br>coefficient | Standard error | *p* |
| --- | --- | --- | --- |
| Age | 0.05 | 1.19 | 0.28 |
| Sex | 0.28 | 1.26 | **0.002** |
| Kinship | 0.21 | 2.23 | **0.01** |
| Reproductive<br>status | 0.27 | 1.65 | **0.01** |
| Combined<br>gesture | 0.30 | 0.30 | **0.002** |
| Single gesture | 0.29 | 0.09 | **0.004** |

**Table S8.8.** Duration of visual attention away from dyad partner (*r^2^* = 0.14)

|  | Standardized<br>coefficient | Standard error | *p* |
| --- | --- | --- | --- |
| Age | 0.341 | 1.724 | **0.001** |
| Sex | -0.076 | 1.319 | 0.215 |
| Kinship | 0.075 | 2.879 | 0.123 |
| Reproductive<br>status | 0.026 | 1.379 | 0.419 |
| Combined<br>gesture | -0.029 | 0.422 | 0.385 |
| Single gesture | 0.183 | 0.112 | **0.045** |

**Table S8.9.**
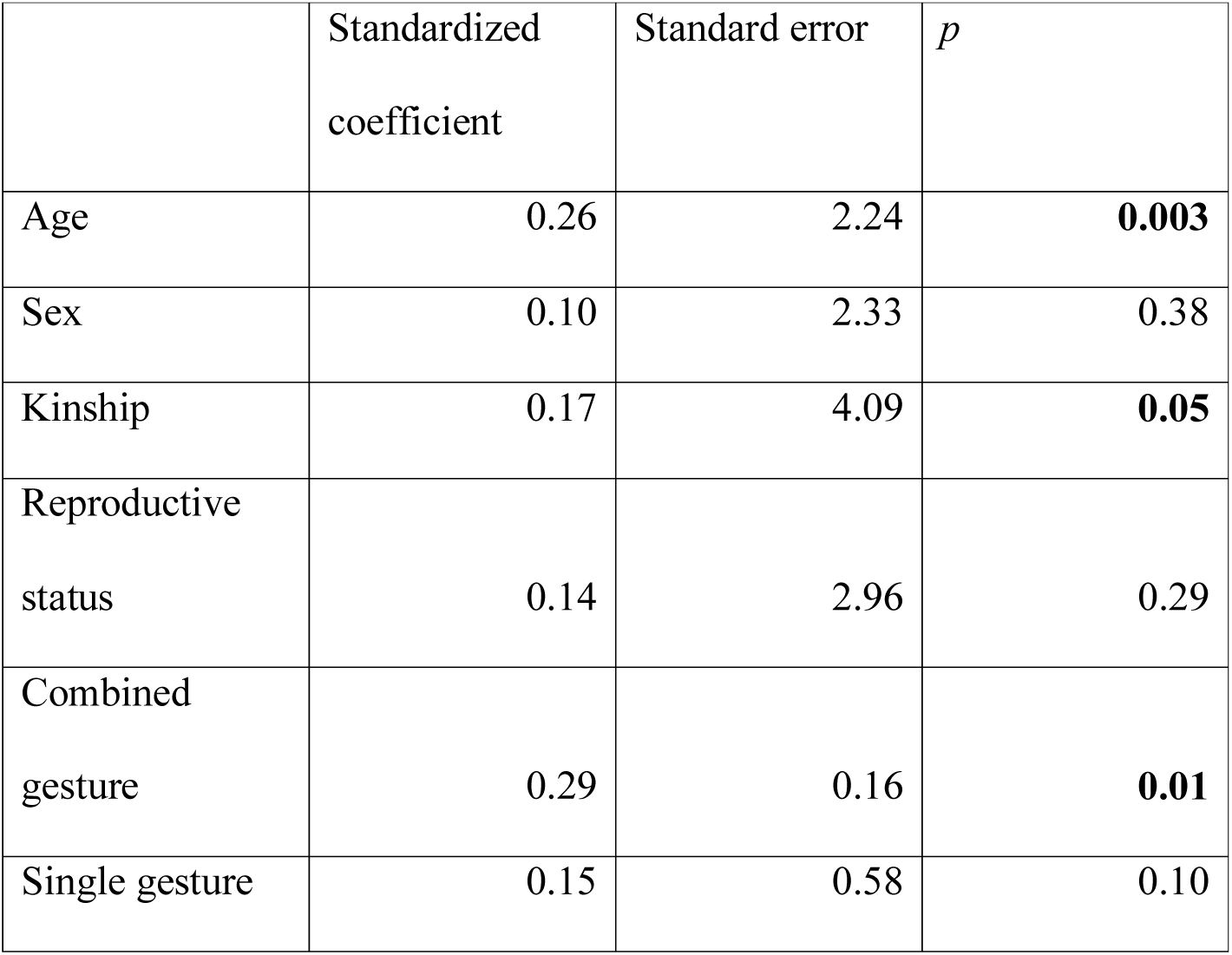
Duration of time in close proximity – within 2 m (*r^2^* = 0.28)

**Table S8.10.** Rate of scratch produced (*r^2^* = 0.179)

|  | Standardized<br>coefficient | Standard error | *p* |
| --- | --- | --- | --- |
| Age | -0.037 | 0.596 | 0.33833 |
| Sex | 0.194 | 0.530 | **0.03348** |
| Kinship | 0.093 | 1.110 | 0.14993 |
| Reproductive<br>status | -0.014 | 0.420 | 0.45027 |
| Combined<br>gesture | 0.155 | 0.154 | 0.06747 |
| Single gesture | 0.259 | 0.039 | **0.01649** |

**Table S8.11.** Rate of scratch received (*r^2^*= 0.051)

|  | Standardized<br>coefficient | Standard error | *p* |
| --- | --- | --- | --- |
| Age | 0.03 | 0.64 | 0.38431 |
| Sex | 0.21 | 0.58 | **0.03248** |
| Kinship | 0.08 | 1.19 | 0.18541 |
| Reproductive<br>status | 0.03 | 0.44 | 0.33133 |
| Combined<br>gesture | 0.09 | 0.16 | 0.18041 |
| Single gesture | -0.14 | 0.04 | 0.05497 |

#### Mutual attention accompanying gestures

Supplementary Table S9. MRQAP regression models predicting durations of social behavior, per hour dyad spent within 10m. Predictor variables were rates of gestures with and without mutual attention between the recipient and the signaller, per hour dyad spent within 10m and demographic variables. Based on 132 chimpanzee dyads. Significant *p* values are indicated in bold. R squared (*r^2^)* denotes amount of variance in the dependent variable explained by the regression model.

**Table S9.1.**
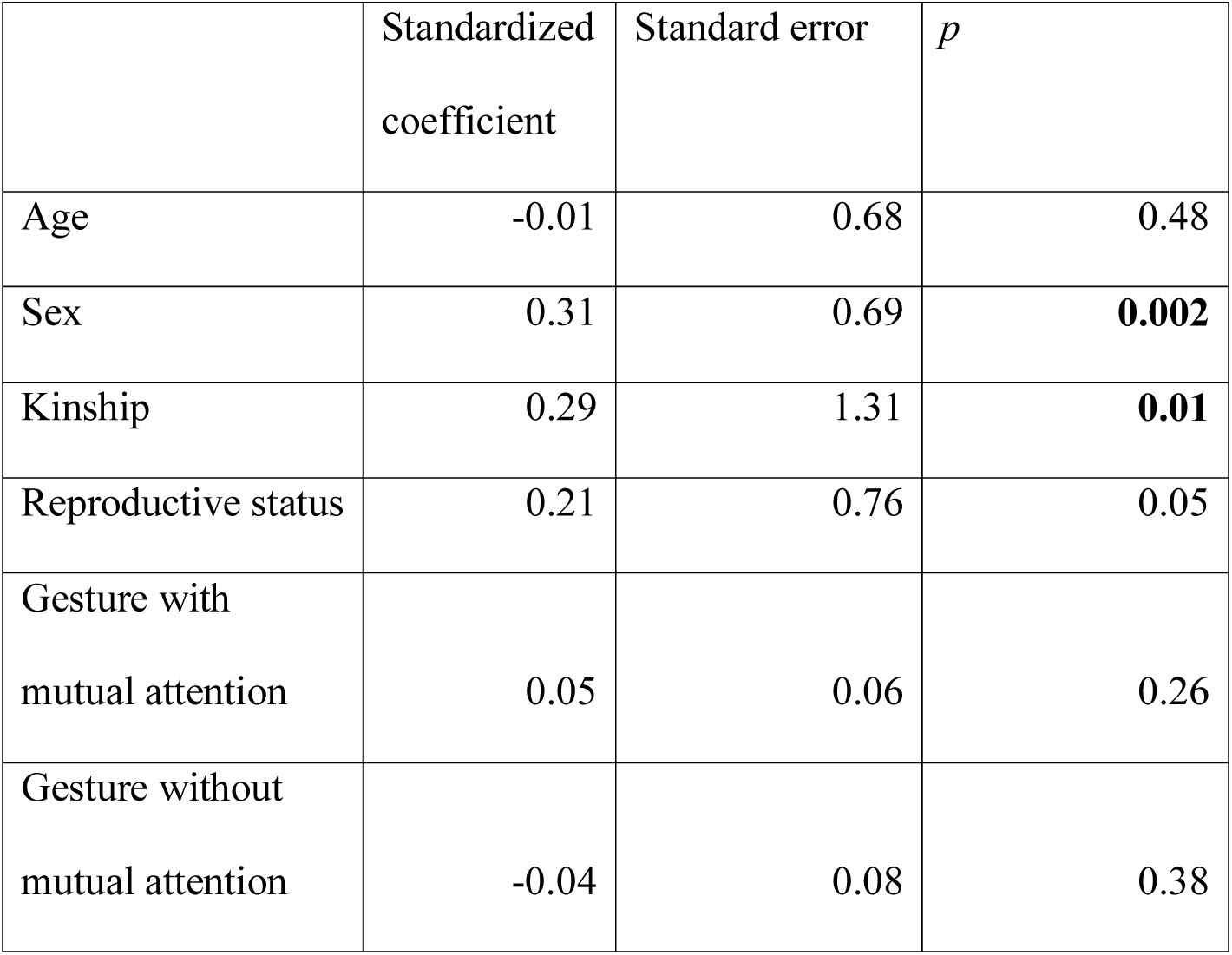
Duration of joint feeding behaviour (*r^2^* = 0.12)

**Table S9.2.** Duration of joint resting behaviour (*r^2^*= 0.08)

| | Standardized coefficient | Standard error | $p$ |
| --- | --- | --- | --- |
| Age | 0.28 | 1.41 | **0.01** |
| Sex | -0.16 | 1.32 | 0.10 |
| Kinship | -0.02 | 2.41 | 0.44 |
| Reproductive status | -0.05 | 1.50 | 0.35 |
| Gesture with mutual attention | 0.004 | 0.16 | 0.37 |
| Gesture without mutual attention | 0.10 | 0.19 | 0.08 |

**Table S9.3.** Duration of joint travelling behaviour (*r^2^* = 0.28)

|  | Standardized coefficient | Standard error | *p* |
| --- | --- | --- | --- |
| Age | 0.23 | 0.36 | **0.01** |
| Sex | -0.02 | 0.32 | 0.40 |
| Kinship | 0.02 | 0.60 | 0.29 |
| Reproductive status | -0.03 | 0.31 | 0.39 |
| Gesture with mutual attention | 0.37 | 0.04 | **0.01** |
| Gesture without mutual attention | 0.12 | 0.04 | 0.07 |

**Table S9.4.**
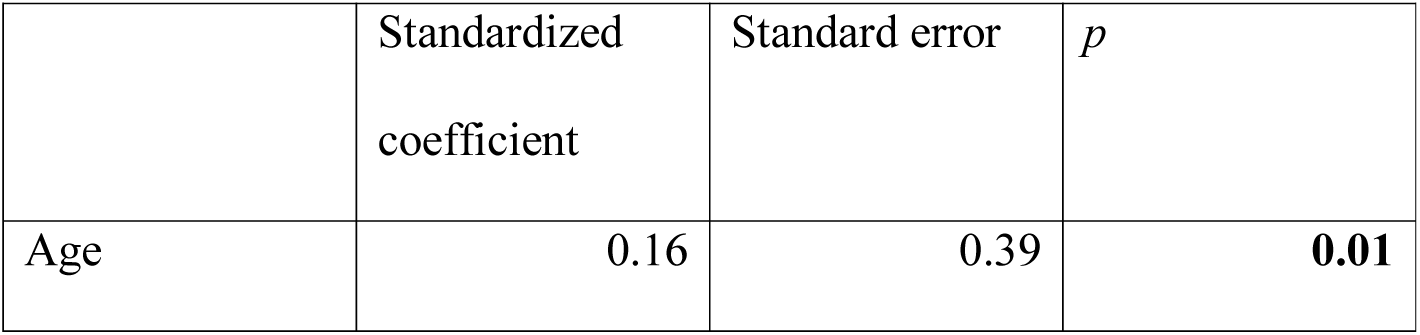
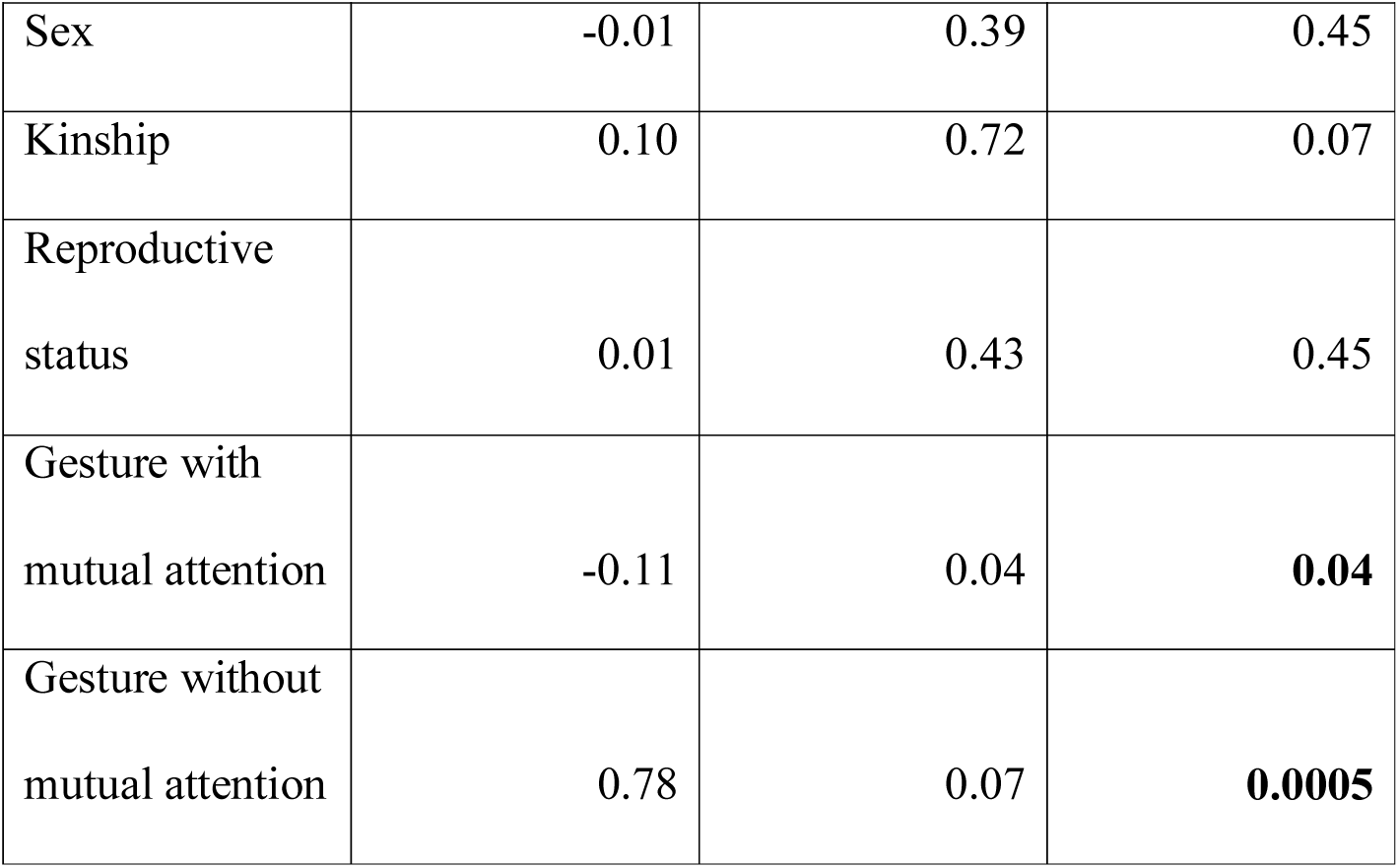
Duration of giving grooming (*r^2^*= 0.57)

**Table S9.5.** Duration of mutual grooming (*r^2^* = 0.42)

| | Standardized coefficient | Standard error | $p$ |
| --- | --- | --- | --- |
| Age | 0.14 | 0.49 | 0.06 |
| Sex | -0.07 | 0.49 | 0.41 |
| Kinship | 0.03 | 0.94 | 0.68 |
| Reproductive status | 0.08 | 0.49 | 0.32 |
| Gesture with mutual attention | 0.63 | 0.07 | **0.001** |
| Gesture without mutual attention | -0.01 | 0.06 | 0.78 |

**Table S9.6.** Duration of receiving grooming (*r^2^*= 0.07)

| | Standardized | Standard error | $p$ |
| --- | --- | --- | --- |

|  | coefficient |  |  |
| --- | --- | --- | --- |
| Age | -0.11 | 0.49 | 0.10 |
| Sex | 0.17 | 0.48 | 0.051 |
| Kinship | 0.00 | 0.86 | 0.67 |
| Reproductive status | 0.06 | 0.55 | 0.30 |
| Gesture with mutual attention | 0.21 | 0.06 | 0.054 |
| Gesture without mutual attention | 0.02 | 0.06 | 0.24 |

**Table S9.7.** Duration of visual attention towards dyad partner (*r^2^*= 0.49)

|  | Standardized coefficient | Standard error | *p* |
| --- | --- | --- | --- |
| Age | 0.06 | 1.03 | 0.23 |
| Sex | 0.20 | 1.07 | **0.01** |
| Kinship | 0.19 | 1.99 | **0.01** |
| Reproductive status | 0.22 | 1.33 | **0.02** |
| Gesture with mutual attention | 0.56 | 0.14 | **0.0005** |
| Gesture without mutual attention | 0.12 | 0.13 | 0.07 |

**Table S9.8.** Duration of visual attention away from dyad partner (*r^2^* = 0.19)

|  | Standardized<br>coefficient | Standard error | *p* |
| --- | --- | --- | --- |
| Age | 0.348 | 1.662 | **0.001** |
| Sex | -0.094 | 1.299 | 0.142 |
| Kinship | 0.068 | 2.725 | 0.130 |
| Reproductive<br>status | 0.003 | 1.329 | 0.494 |
| Gesture with<br>mutual attention | -0.032 | 0.167 | 0.384 |
| Gesture without<br>mutual attention | 0.288 | 0.201 | **0.015** |

**Table S9.9.**
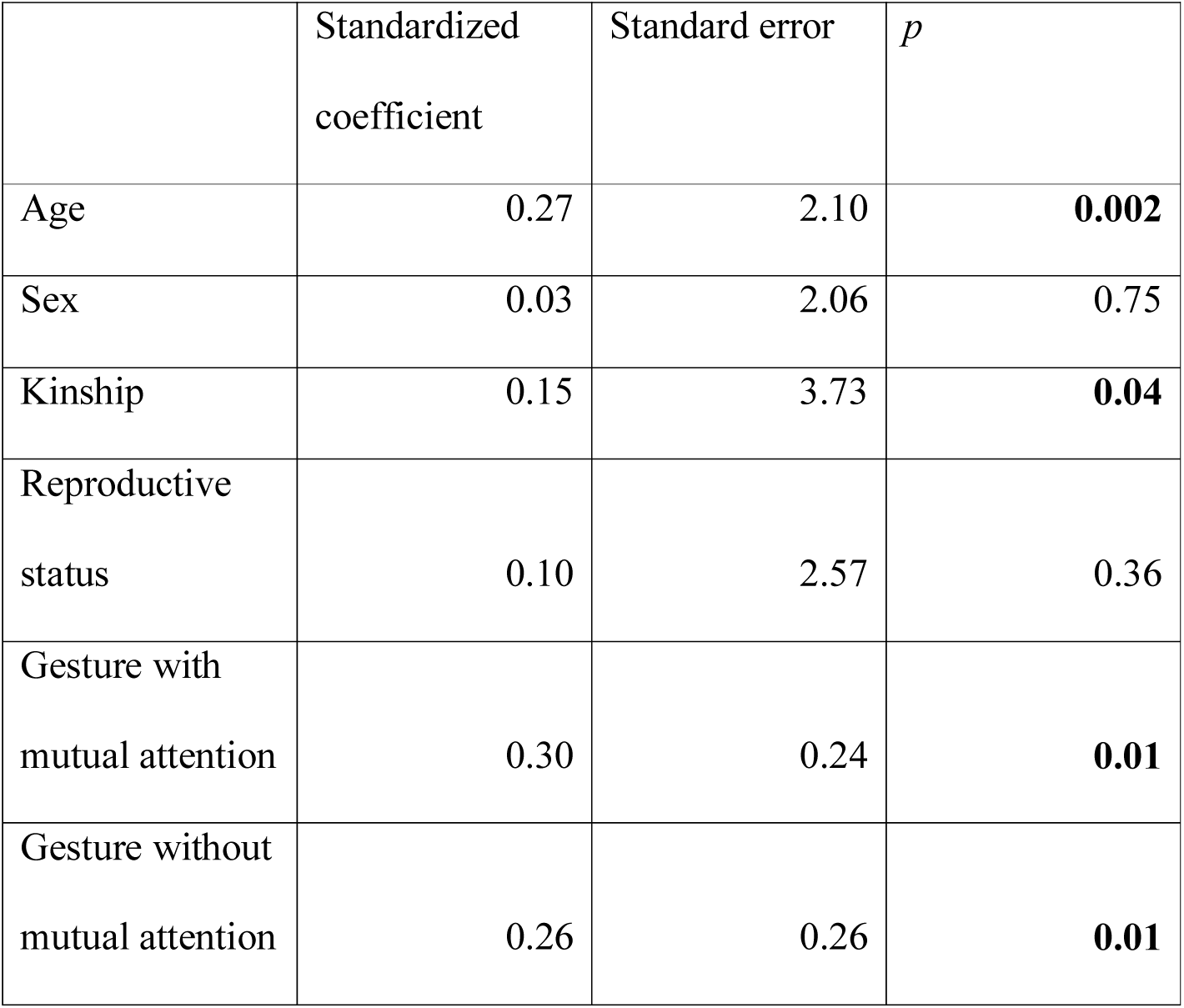
Duration of time in close proximity – within 2 m (*r^2^*= 0.36)

|  | Standardized<br>coefficient | Standard error | *p* |
| --- | --- | --- | --- |
| Age | 0.27 | 2.10 | **0.002** |
| Sex | 0.03 | 2.06 | 0.75 |
| Kinship | 0.15 | 3.73 | **0.04** |
| Reproductive<br>status | 0.10 | 2.57 | 0.36 |
| Gesture with<br>mutual attention | 0.30 | 0.24 | **0.01** |
| Gesture without<br>mutual attention | 0.26 | 0.26 | **0.01** |

**Table S9.10.** Rate of scratch produced (*r^2^*= 0.299)

|  | Standardized<br>coefficient | Standard error | *p* |
| --- | --- | --- | --- |
| Age | -0.031 | 0.540 | 0.36832 |
| Sex | 0.165 | 0.488 | 0.05097 |
| Kinship | 0.086 | 1.004 | 0.15242 |
| Reproductive<br>status | -0.033 | 0.402 | 0.33983 |
| Gesture with<br>mutual attention | 0.527 | 0.061 | **0.001** |
| Gesture without<br>mutual attention | -0.026 | 0.068 | 0.43028 |

**Table S9.11.** Rate of scratch received (*r^2^* = 0.058)

|  | Standardized<br>coefficient | Standard error | *p* |
| --- | --- | --- | --- |
| Age | 0.02 | 0.64 | 0.4018 |
| Sex | 0.21 | 0.56 | **0.03298** |
| Kinship | 0.08 | 1.15 | 0.17841 |
| Reproductive<br>status | 0.04 | 0.44 | 0.32184 |
| Gesture with mutual attention | 0.05 | 0.06 | 0.24138 |
| Gesture without mutual attention | -0.13 | 0.08 | **0.04948** |

#### Bodily and manual gestures

Supplementary Table S10. MRQAP regression models predicting durations of social behavior, per hour dyad spent within 10m. Predictor variables were rates of bodily and manual gestures between the recipient and the signaller, per hour dyad spent within 10m and demographic variables. Based on 132 chimpanzee dyads. Significant *p* values are indicated in bold. R squared (*r^2^)* denotes amount of variance in the dependent variable explained by the regression model.

**Table S10.1.** Duration of joint feeding behaviour (*r^2^*= 0.156)

| | Standardized coefficient | Standard error | $p$ |
| --- | --- | --- | --- |
| Age | -0.043 | 0.674 | 0.365 |
| Sex | 0.150 | 0.581 | 0.083 |
| Kinship | 0.285 | 1.291 | **0.010** |
| Reproductive status | 0.259 | 0.538 | **0.001** |
| Bodily | 0.066 | 0.041 | 0.237 |
| Manual | -0.082 | 0.088 | 0.201 |

**Table S10.2.** Duration of joint resting behaviour (*r^2^* = 0.072)

|  | Standardized<br>coefficient | Standard error | *p* |
| --- | --- | --- | --- |
| Age | 0.280 | 1.405 | **0.004** |
| Sex | -0.116 | 1.102 | 0.081 |
| Kinship | -0.018 | 2.383 | 0.463 |
| Reproductive<br>status | -0.027 | 1.113 | 0.405 |
| Bodily | 0.060 | 0.091 | 0.169 |
| Manual | -0.014 | 0.192 | 0.486 |

**Table S10.3.** Duration of joint travelling behaviour (*r^2^*= 0.294)

|  | Standardized<br>coefficient | Standard error | *p* |
| --- | --- | --- | --- |
| Age | 0.177 | 0.345 | **0.027** |
| Sex | 0.034 | 0.279 | 0.345 |
| Kinship | 0.021 | 0.619 | 0.286 |
| Reproductive<br>status | -0.066 | 0.252 | 0.191 |
| Bodily | 0.503 | 0.026 | **0.003** |
| Manual | -0.062 | 0.049 | 0.194 |

**Table S10.4.** Duration of giving grooming (*r^2^* = 0.291)

|  | Standardized<br>coefficient | Standard error | *p* |
| --- | --- | --- | --- |
| Age | 0.171 | 0.498 | **0.025** |
| Sex | 0.045 | 0.410 | 0.290 |
| Kinship | 0.124 | 0.919 | 0.078 |
| Reproductive<br>status | 0.092 | 0.405 | 0.124 |
| Bodily | 0.220 | 0.034 | **0.024** |
| Manual | 0.293 | 0.072 | **0.016** |

**Table S10.5.** Duration of mutual grooming (*r^2^*= 0.327)

|  | Standardized<br>coefficient | Standard error | *p* |
| --- | --- | --- | --- |
| Age | 0.073 | 0.589 | 0.219 |
| Sex | -0.102 | 0.494 | 0.132 |
| Kinship | 0.031 | 1.081 | 0.270 |
| Reproductive<br>status | 0.097 | 0.456 | 0.115 |
| Bodily | 0.585 | 0.043 | **0.001** |
| Manual | -0.075 | 0.077 | 0.149 |

**Table S10.6.** Duration of receiving grooming (*r^2^* = 0.087)

|  | Standardized<br>coefficient | Standard error | *p* |
| --- | --- | --- | --- |
| Age | -0.148 | 0.516 | **0.046** |
| Sex | 0.123 | 0.426 | 0.099 |
| Kinship | 0.004 | 0.886 | 0.350 |
| Reproductive<br>status | 0.136 | 0.423 | 0.054 |
| Bodily | 0.255 | 0.037 | **0.036** |
| Manual | -0.092 | 0.080 | 0.101 |

**Table S10.7.** Duration of visual attention towards dyad partner (*r^2^* = 0.421)

|  | Standardized<br>coefficient | Standard error | *p* |
| --- | --- | --- | --- |
| Age | -0.013 | 1.188 | 0.452 |
| Sex | 0.070 | 0.977 | 0.202 |
| Kinship | 0.198 | 2.219 | **0.011** |
| Reproductive<br>status | 0.261 | 1.183 | **0.005** |
| Bodily | 0.549 | 0.092 | **0.001** |
| Manual | -0.024 | 0.153 | 0.401 |

**Table S10.8.** Duration of visual attention away from dyad partner (*r^2^* = 0.144)

|  | Standardized<br>coefficient | Standard error | *p* |
| --- | --- | --- | --- |
| Age | 0.340 | 1.771 | **0.002** |
| Sex | -0.069 | 1.294 | 0.219 |
| Kinship | 0.076 | 2.905 | 0.126 |
| Reproductive<br>status | 0.021 | 1.307 | 0.431 |
| Bodily | 0.139 | 0.122 | 0.083 |
| Manual | 0.028 | 0.226 | 0.291 |

**Table S10.9.**
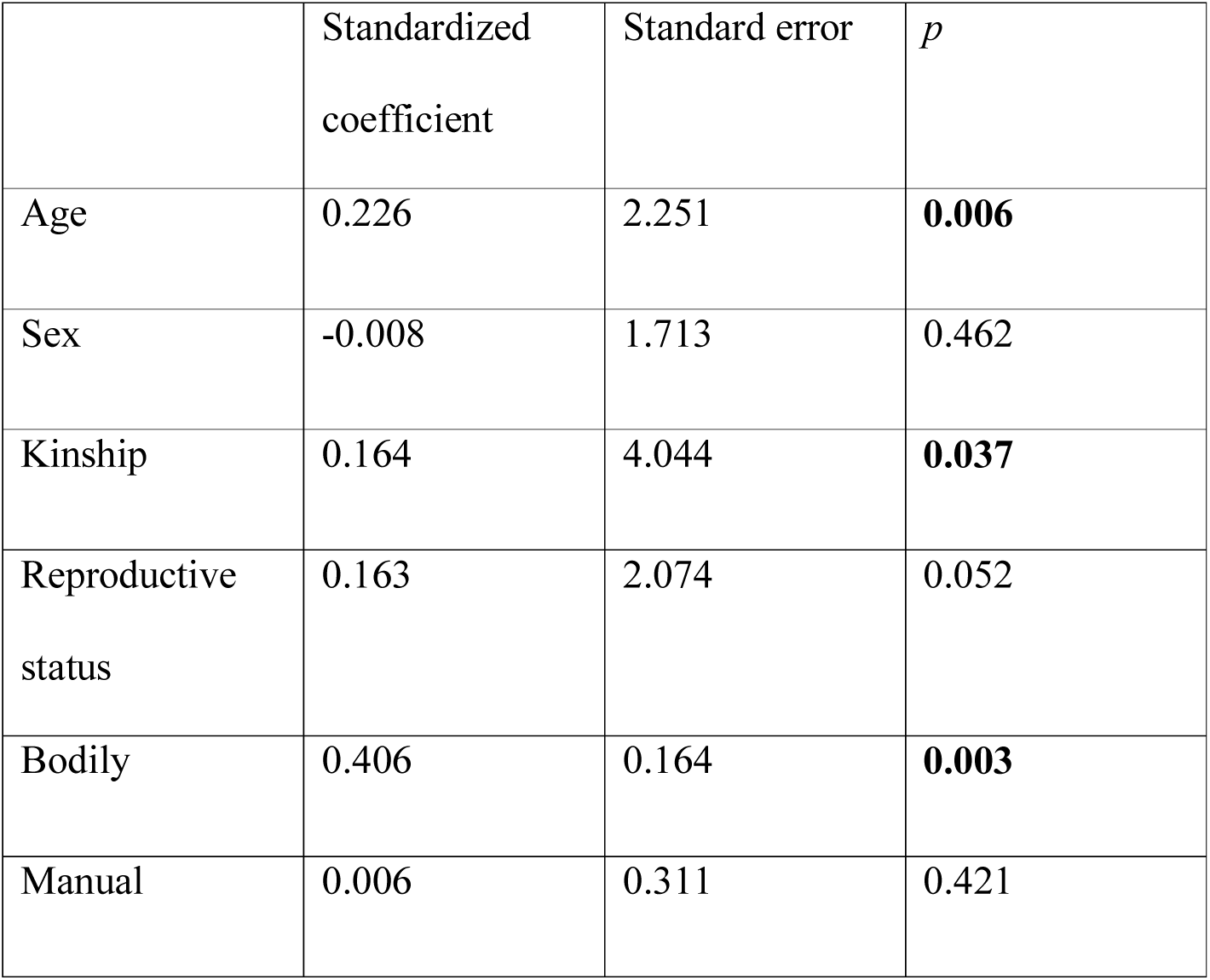
Duration of time in close proximity – within 2 m (*r^2^* = 0.312)

|  | Standardized<br>coefficient | Standard error | *p* |
| --- | --- | --- | --- |
| Age | 0.226 | 2.251 | **0.006** |
| Sex | -0.008 | 1.713 | 0.462 |
| Kinship | 0.164 | 4.044 | **0.037** |
| Reproductive<br>status | 0.163 | 2.074 | 0.052 |
| Bodily | 0.406 | 0.164 | **0.003** |
| Manual | 0.006 | 0.311 | 0.421 |

**Table S10.10.** Rate of scratch produced (*r^2^* = 0.241)

|  | Standardized<br>coefficient | Standard error | *p* |
| --- | --- | --- | --- |
| Age | -0.098 | 0.599 | 0.14443 |
| Sex | 0.189 | 0.522 | **0.03448** |
| Kinship | 0.088 | 1.063 | 0.15442 |
| Reproductive<br>status | -0.018 | 0.394 | 0.4038 |
| Bodily | 0.546 | 0.041 | **0.0005** |
| Manual | -0.184 | 0.079 | **0.01149** |

**Table S10.11.** Rate of scratch received (*r^2^* = 0.051)

|  | Standardized<br>coefficient | Standard error | *p* |
| --- | --- | --- | --- |
| Age | 0.02 | 0.65 | 0.42129 |
| Sex | 0.20 | 0.58 | **0.04248** |
| Kinship | 0.08 | 1.19 | 0.1994 |
| Reproductive<br>status | 0.04 | 0.45 | 0.33183 |
| Bodily | -0.02 | 0.04 | 0.47776 |
| Manual | -0.06 | 0.08 | 0.29685 |

#### Gesture events

Supplementary Table S11. MRQAP regression models predicting durations of social behavior, per hour dyad spent within 10m. Predictor variable was rates of gesture events between the recipient and the signaller, per hour dyad spent within 10m and demographic variables. Based on 132 chimpanzee dyads. Significant *p* values are indicated in bold. R squared (*r^2^)* denotes amount of variance in the dependent variable explained by the regression model.

**Table S11.1.** Duration of joint feeding behaviour (*r^2^*= 0.151)

| | Standardized coefficient | Standard error | $p$ |
| --- | --- | --- | --- |
| Age | -0.026 | 0.656 | 0.417 |
| Sex | 0.149 | 0.567 | 0.076 |
| Kinship | 0.287 | 1.236 | **0.008** |
| Reproductive status | 0.260 | 0.563 | **0.003** |
| Events | -0.008 | 0.027 | 0.509 |

**Table S11.2.**
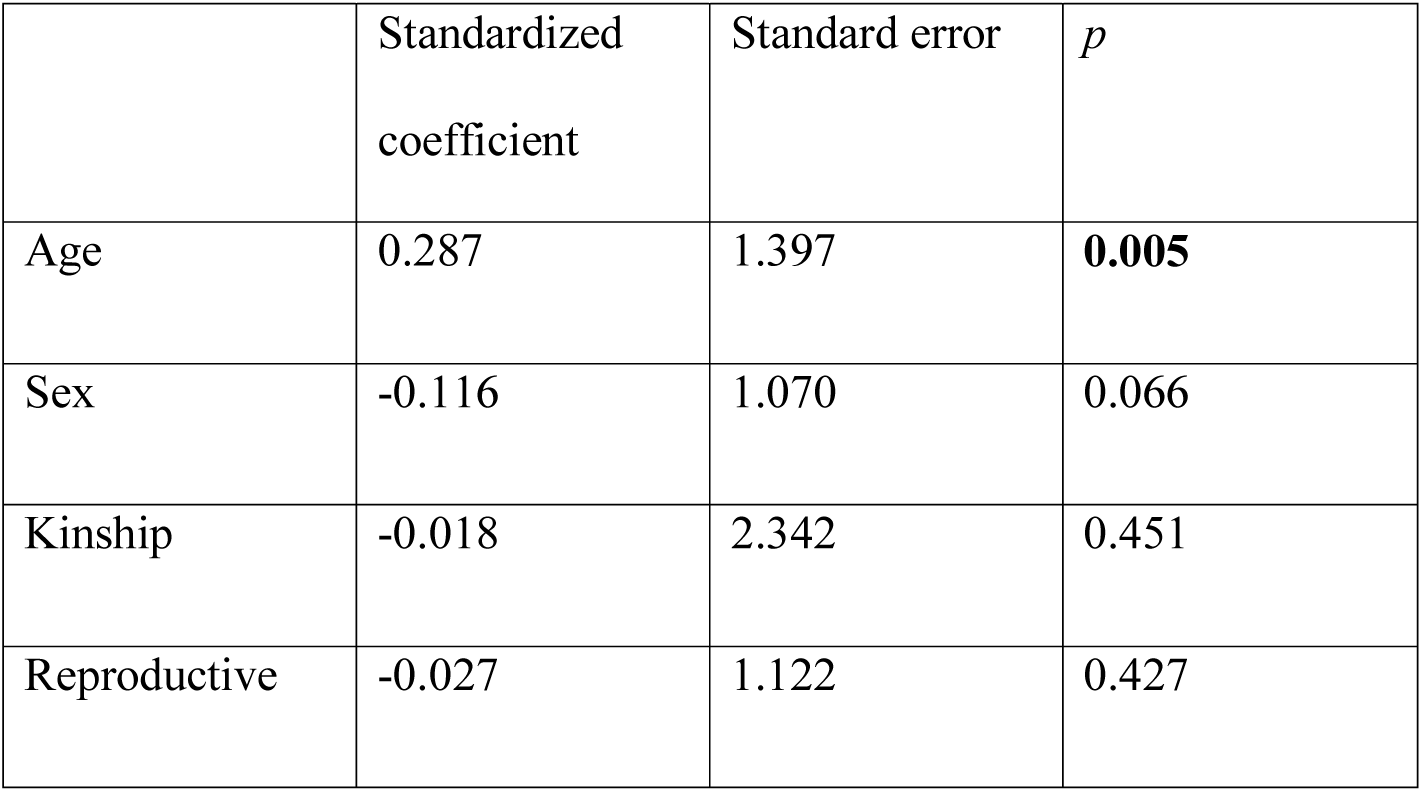
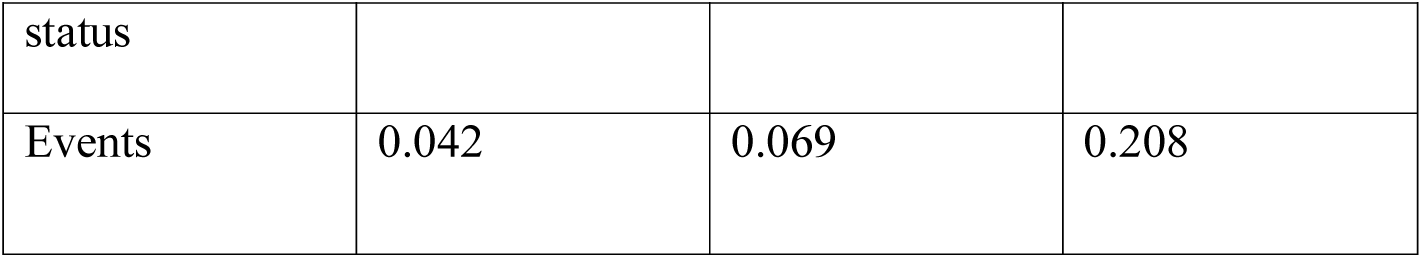
Duration of joint resting behaviour (*r^2^* = 0.071)

**Table S11.3.** Duration of joint travelling behaviour (*r^2^*= 0.240)

|  | Standardized<br>coefficient | Standard error | *p* |
| --- | --- | --- | --- |
| Age | 0.225 | 0.366 | **0.017** |
| Sex | 0.035 | 0.302 | 0.348 |
| Kinship | 0.026 | 0.622 | 0.252 |
| Reproductive<br>status | -0.062 | 0.268 | 0.219 |
| Events | 0.396 | 0.018 | **0.003** |

**Table S11.4.** Duration of giving grooming (*r^2^* = 0.293)

|  | Standardized<br>coefficient | Standard error | *p* |
| --- | --- | --- | --- |
| Age | 0.141 | 0.484 | **0.047** |
| Sex | 0.046 | 0.414 | 0.290 |
| Kinship | 0.125 | 0.886 | 0.064 |
| Reproductive<br>status | 0.086 | 0.389 | 0.128 |
| Events | 0.467 | 0.027 | **0.002** |

**Table S11.5.** Duration of mutual grooming (*r^2^* = 0.248)

|  | Standardized<br>coefficient | Standard error | *p* |
| --- | --- | --- | --- |
| Age | 0.130 | 0.618 | 0.105 |
| Sex | -0.101 | 0.524 | 0.150 |
| Kinship | 0.035 | 1.110 | 0.241 |
| Reproductive<br>status | 0.103 | 0.508 | 0.139 |
| Events | 0.451 | 0.031 | **0.003** |

**Table S11.6.** Duration of receiving grooming (*r^2^* = 0.071)

|  | Standardized<br>coefficient | Standard error | *p* |
| --- | --- | --- | --- |
| Age | -0.117 | 0.483 | 0.086 |
| Sex | 0.123 | 0.404 | 0.092 |
| Kinship | 0.007 | 0.876 | 0.315 |
| Reproductive<br>status | 0.137 | 0.450 | 0.056 |
| Events | 0.164 | 0.025 | 0.065 |

**Table S11.7.**
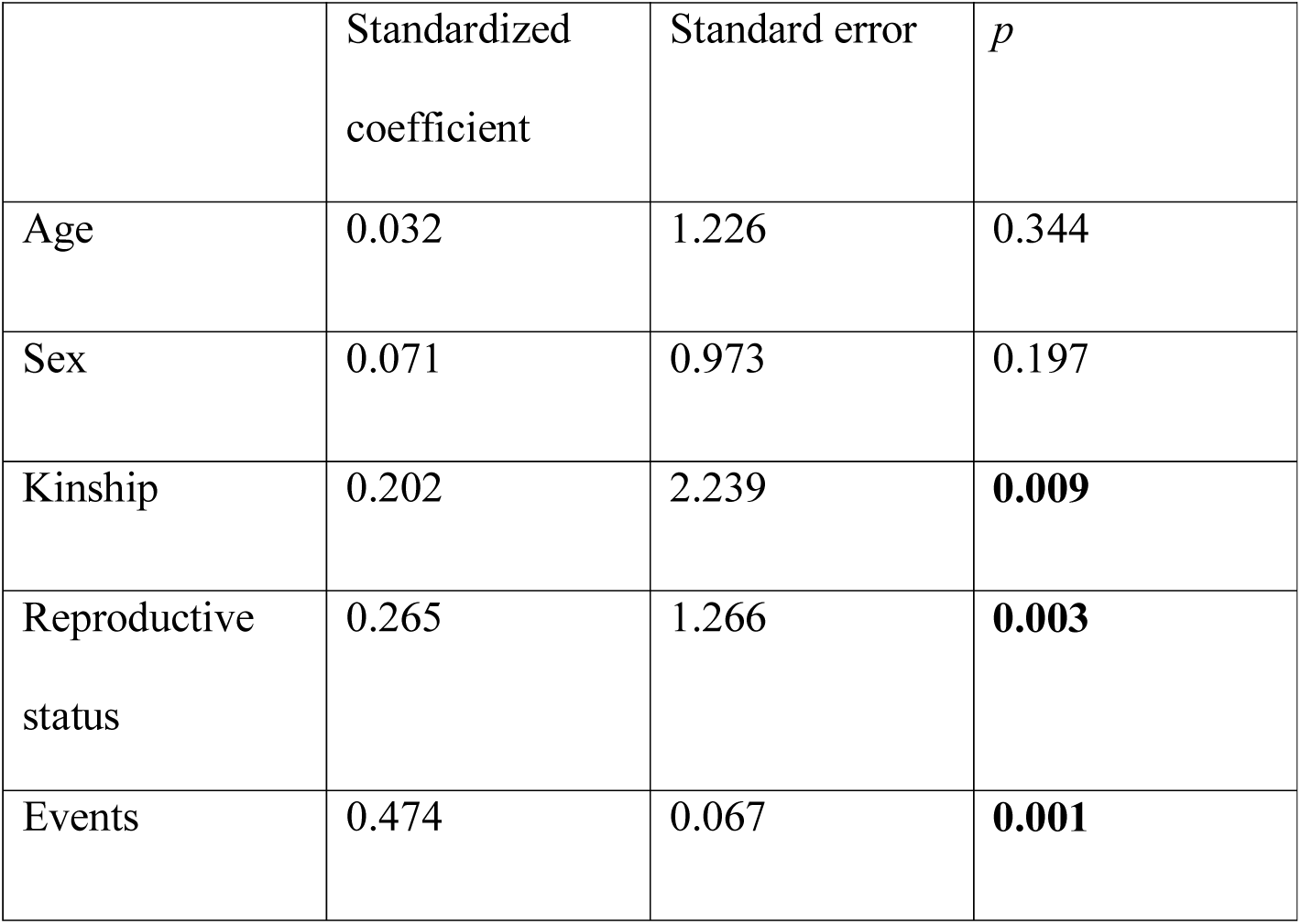
Duration of visual attention towards dyad partner (*r^2^* = 0.369)

|  | Standardized<br>coefficient | Standard error | *p* |
| --- | --- | --- | --- |
| Age | 0.032 | 1.226 | 0.344 |
| Sex | 0.071 | 0.973 | 0.197 |
| Kinship | 0.202 | 2.239 | **0.009** |
| Reproductive<br>status | 0.265 | 1.266 | **0.003** |
| Events | 0.474 | 0.067 | **0.001** |

**Table S11.8.**
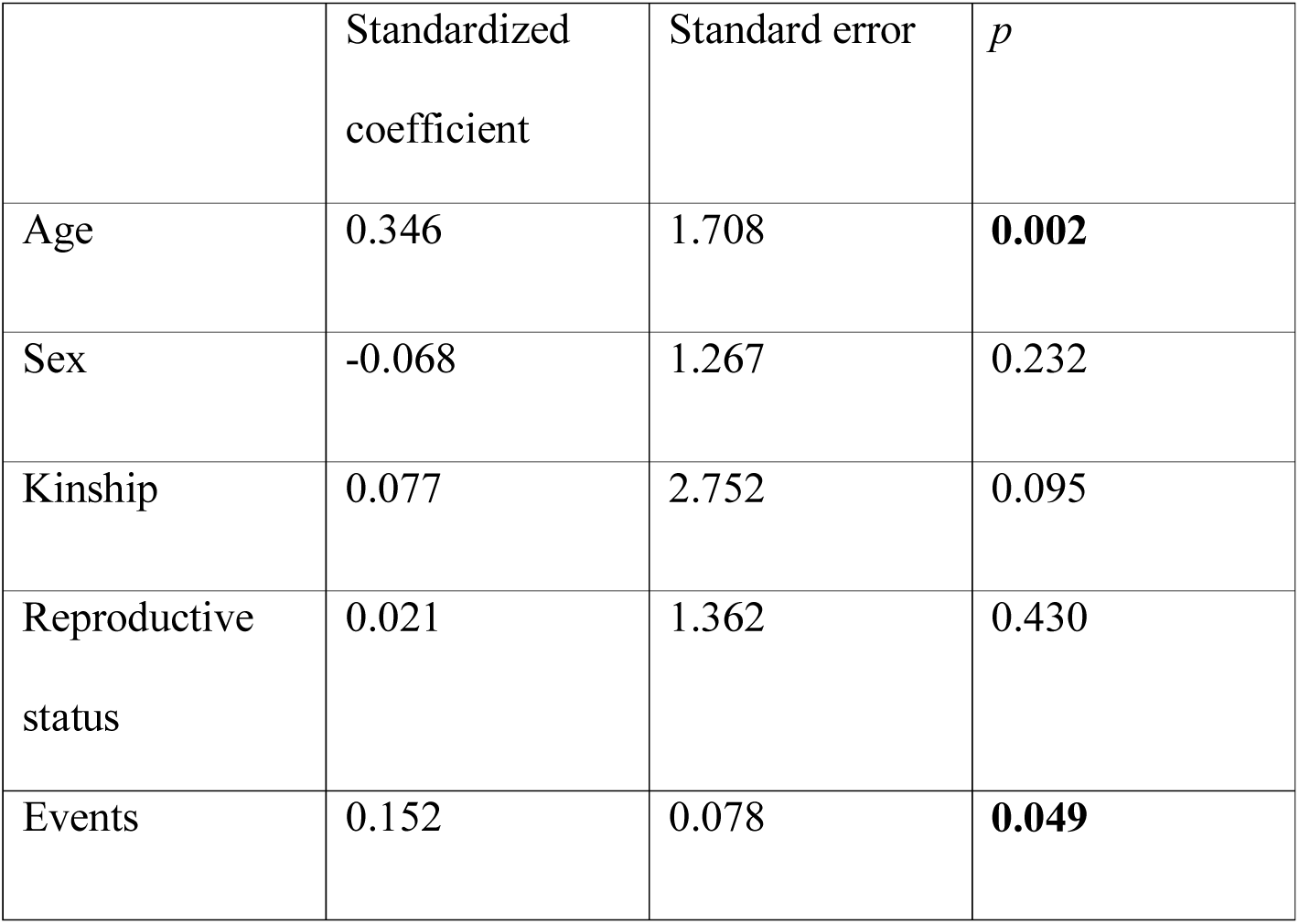
Duration of visual attention away from dyad partner (*r^2^* = 0.143)

**Table S11.9.** Duration of time in close proximity – within 2 m (*r^2^* = 0.288)

|  | Standardized<br>coefficient | Standard error | *p* |
| --- | --- | --- | --- |
| Age | 0.256 | 2.271 | **0.005** |
| Sex | -0.007 | 1.668 | 0.472 |
| Kinship | 0.168 | 3.990 | **0.027** |
| Reproductive status | 0.165 | 2.197 | 0.057 |
| Events | 0.373 | 0.122 | **0.003** |

**Table S11.10.** Rate of scratch produced (*r^2^* = 0.172)

|  | Standardized coefficient | Standard error | *p* |
| --- | --- | --- | --- |
| Age | -0.033 | 0.585 | 0.37281 |
| Sex | 0.189 | 0.528 | **0.03598** |
| Kinship | 0.094 | 1.106 | 0.15692 |
| Reproductive status | -0.016 | 0.432 | 0.43478 |
| Events | 0.364 | 0.027 | **0.003** |

**Table S11.11.** Rate of scratch received (*r^2^* = 0.051)

|  | Standardized coefficient | Standard error | *p* |
| --- | --- | --- | --- |
| Age | 0.03 | 0.62 | 0.38731 |
| Sex | 0.20 | 0.55 | **0.03998** |
| Kinship | 0.08 | 1.14 | 0.1929 |
| Reproductive status | 0.04 | 0.46 | 0.33183 |
| Events | -0.07 | 0.03 | 0.1909 |

#### Non-indicative and indicative gestures

Supplementary Table S12. MRQAP regression models predicting durations of social behavior, per hour dyad spent within 10m. Predictor variables were rates of non-indicative and indicative gestures between the recipient and the signaller, per hour dyad spent within 10m and demographic variables. Based on 132 chimpanzee dyads. Significant *p* values are indicated in bold. R squared (*r^2^)* denotes amount of variance in the dependent variable explained by the regression model.

**Table S12.1.**
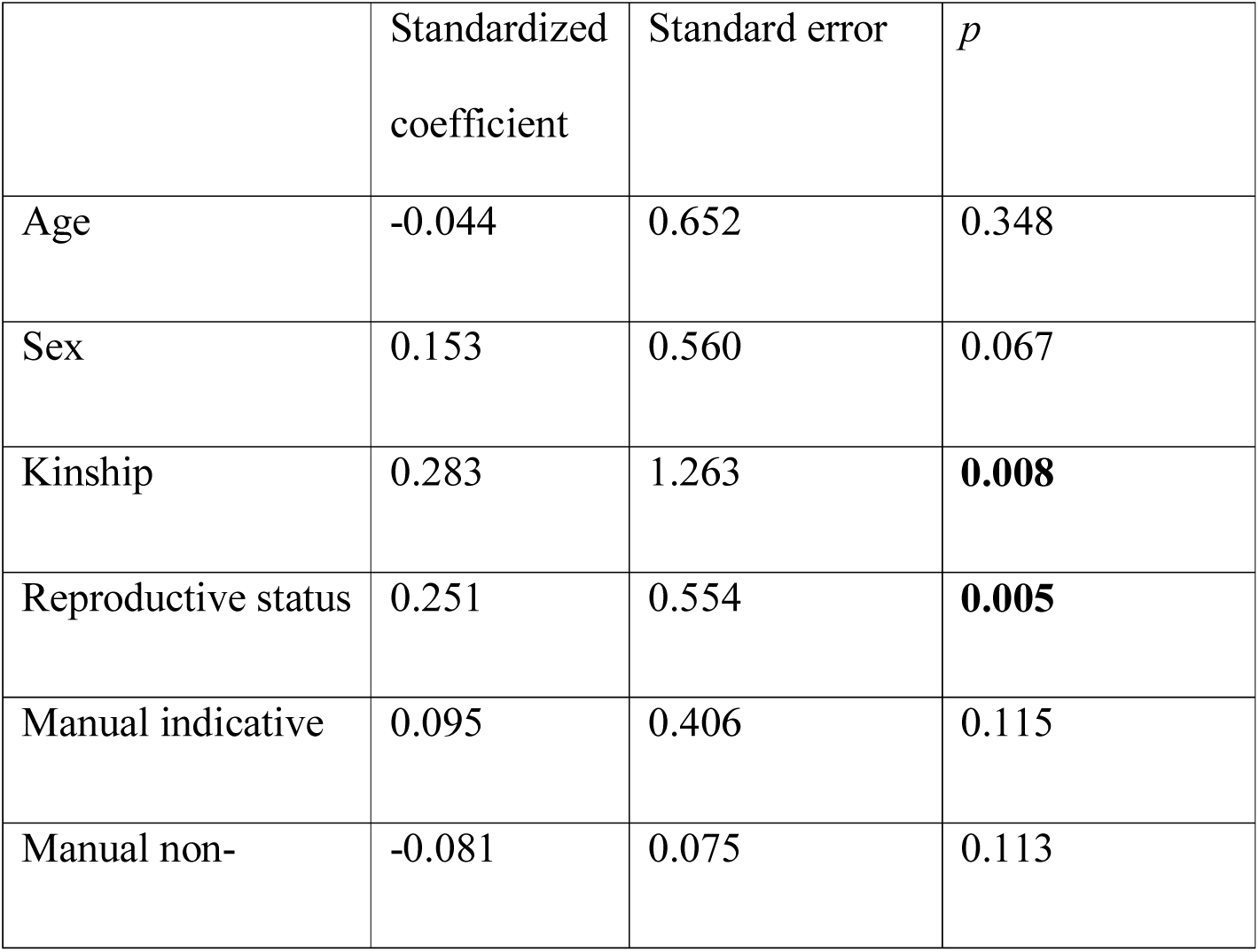
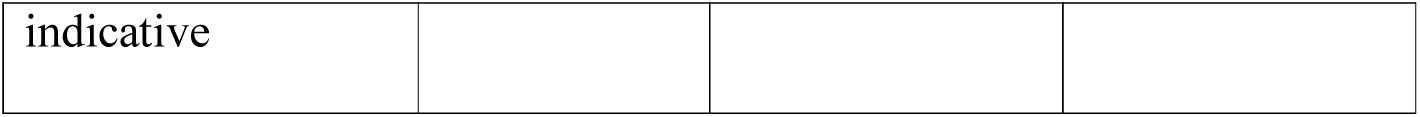
Duration of joint feeding behaviour (*r^2^* = 0.162)

**Table S12.2.** Duration of joint resting behaviour (*r^2^*= 0.072)

|  | Standardized<br>coefficient | Standard error | *p* |
| --- | --- | --- | --- |
| Age | 0.287 | 1.414 | **0.005** |
| Sex | -0.114 | 1.123 | 0.086 |
| Kinship | -0.020 | 2.471 | 0.445 |
| Reproductive<br>status | -0.028 | 1.123 | 0.427 |
| Manual<br>indicative | 0.046 | 1.028 | 0.161 |
| Manual non-indicative | 0.000 | 0.182 | 0.622 |

**Table S12.3.**
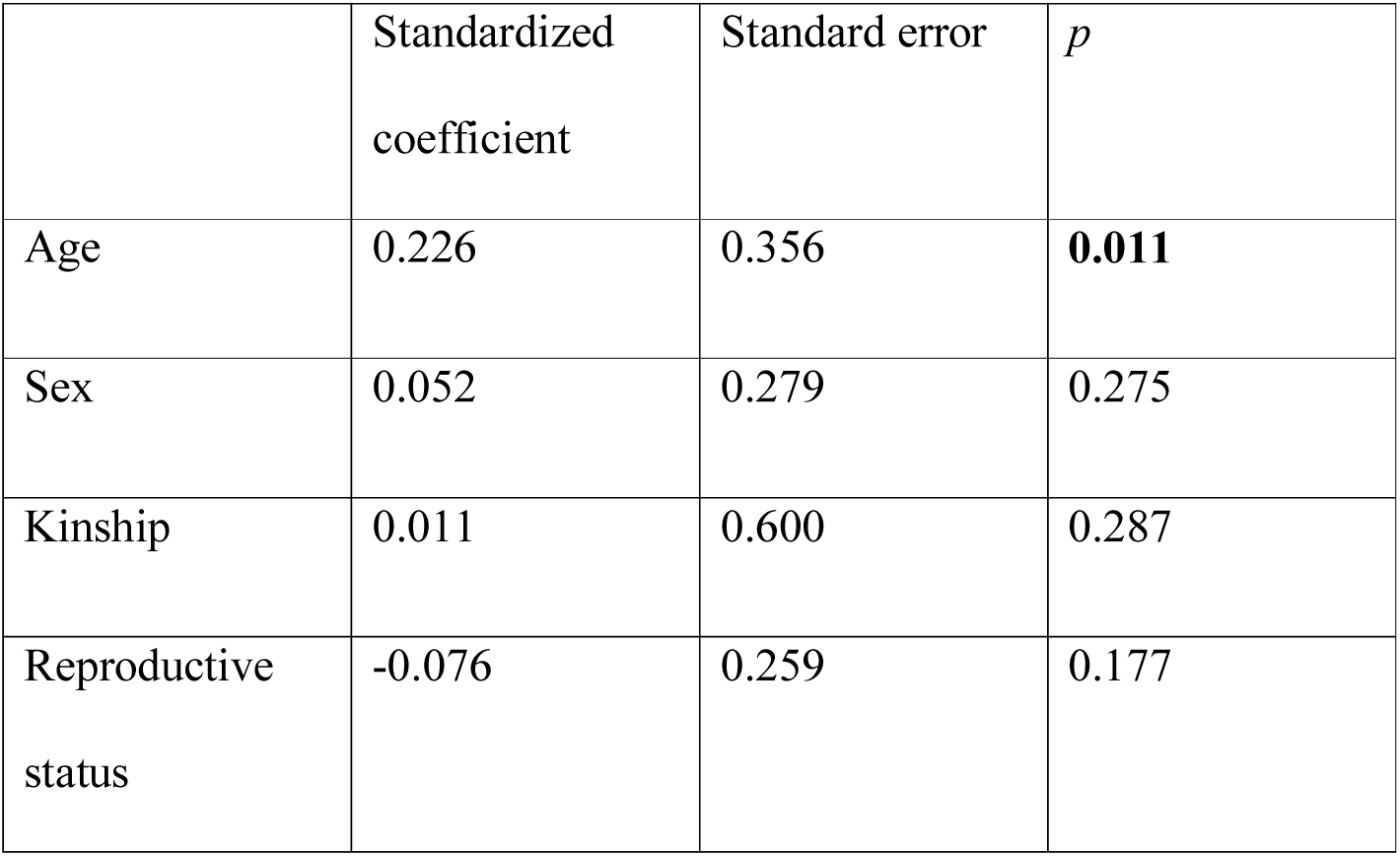
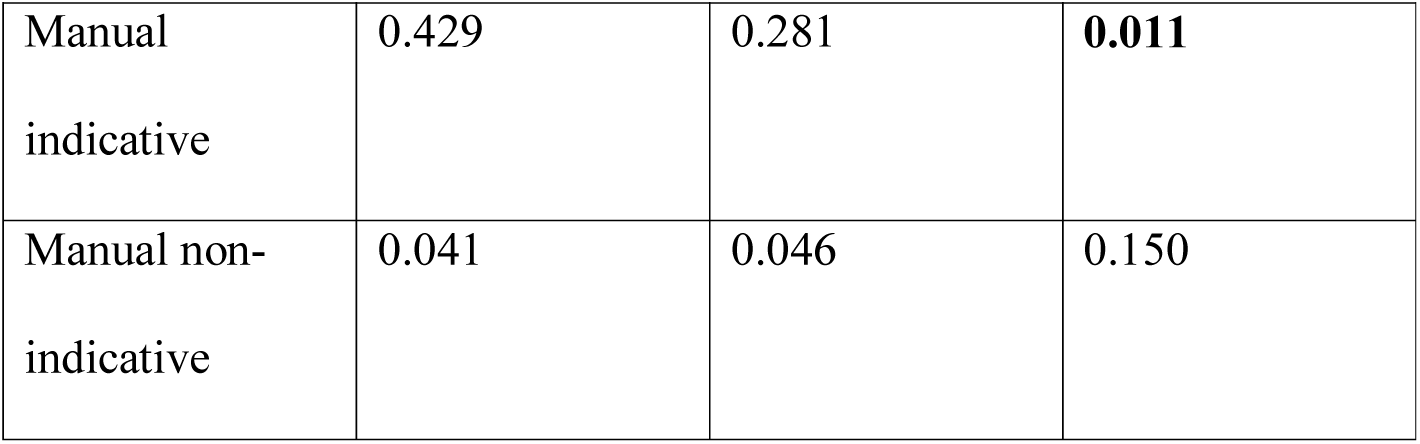
Duration of joint travelling behaviour (*r^2^* = 0.280)

**Table S12.4.** Duration of giving grooming (*r^2^*= 0.275)

|  | Standardized<br>coefficient | Standard error | *p* |
| --- | --- | --- | --- |
| Age | 0.203 | 0.496 | **0.011** |
| Sex | 0.051 | 0.405 | 0.269 |
| Kinship | 0.122 | 0.938 | 0.080 |
| Reproductive<br>status | 0.096 | 0.405 | 0.129 |
| Manual<br>indicative | 0.173 | 0.365 | **0.036** |
| Manual non-indicative | 0.354 | 0.066 | **0.014** |

**Table S12.5.** Duration of mutual grooming (*r^2^* = 0.362)

|  | Standardized<br>coefficient | Standard error | *p* |
| --- | --- | --- | --- |
| Age | 0.120 | 0.556 | 0.094 |
| Sex | -0.079 | 0.470 | 0.186 |
| Kinship | 0.018 | 1.031 | 0.260 |
| Reproductive status | 0.079 | 0.452 | 0.179 |
| Manual indicative | 0.561 | 0.459 | **0.001** |
| Manual non-indicative | 0.016 | 0.068 | 0.246 |

**Table S12.6.** Duration of receiving grooming (*r^2^*= 0.071)

|  | Standardized coefficient | Standard error | *p* |
| --- | --- | --- | --- |
| Age | -0.118 | 0.483 | 0.089 |
| Sex | 0.132 | 0.414 | 0.084 |
| Kinship | 0.000 | 0.807 | 0.685 |
| Reproductive status | 0.135 | 0.436 | 0.060 |
| Manual indicative | 0.170 | 0.382 | **0.046** |
| Manual non-indicative | -0.019 | 0.070 | 0.541 |

**Table S12.7.**
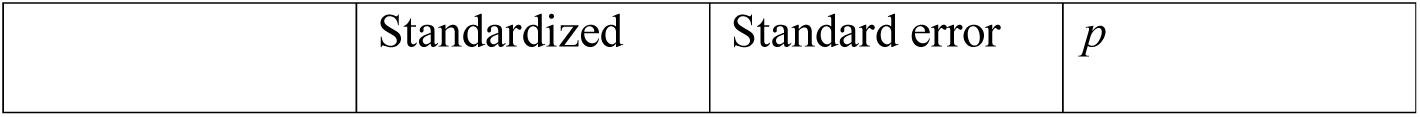
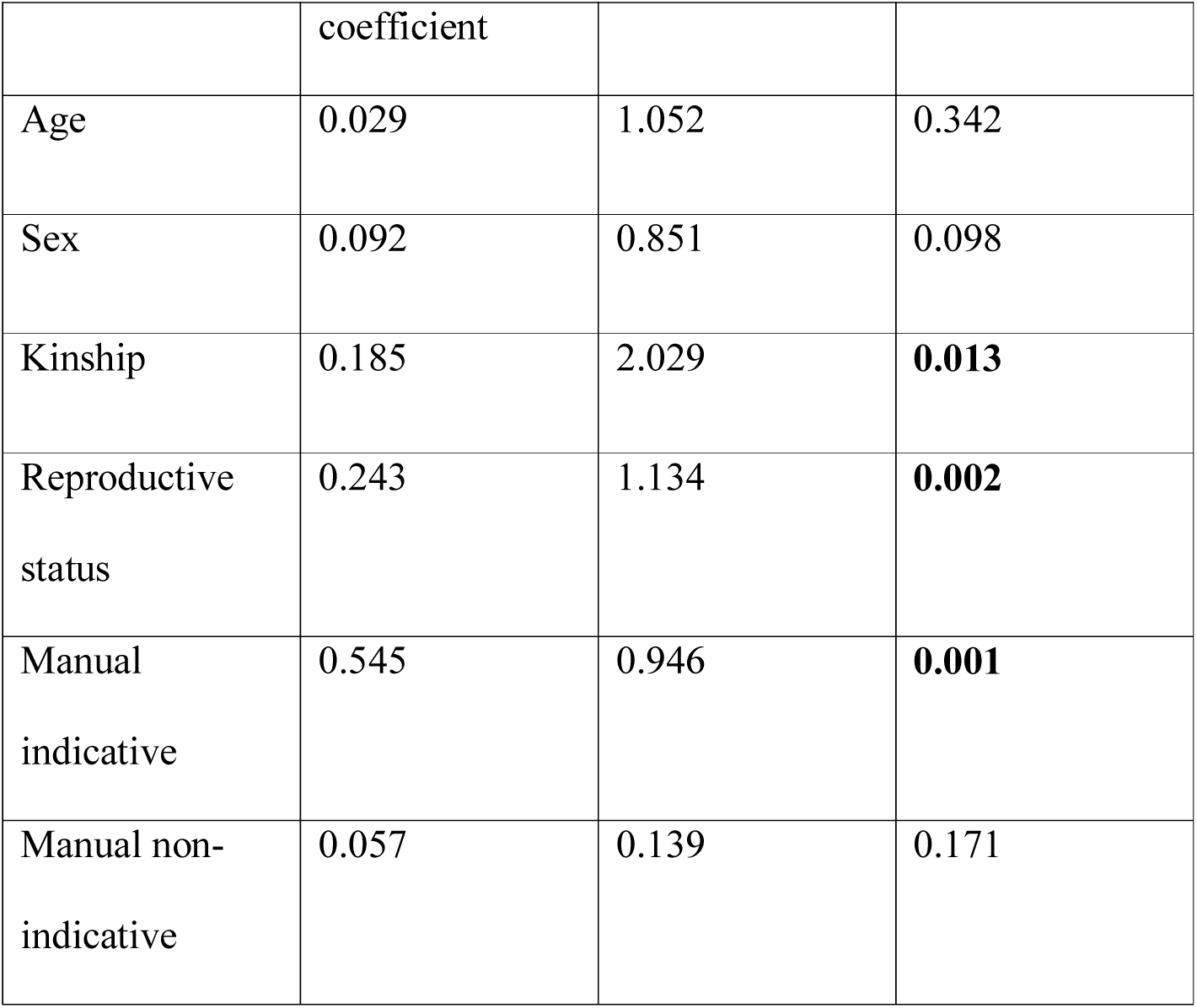
Duration of visual attention towards dyad partner (*r^2^*= 0.463)

**Table S12.8.** Duration of visual attention away from dyad partner (*r^2^* = 0.137)

|  | Standardized<br>coefficient | Standard error | *p* |
| --- | --- | --- | --- |
| Age | 0.359 | 1.719 | **0.001** |
| Sex | -0.065 | 1.277 | 0.249 |
| Kinship | 0.074 | 2.902 | 0.134 |
| Reproductive<br>status | 0.023 | 1.329 | 0.405 |
| Manual<br>indicative | 0.089 | 1.115 | 0.112 |
| Manual non-indicative | 0.071 | 0.213 | 0.120 |

**Table S12.9.** Duration of time in close proximity – within 2 m (*r^2^*= 0.312)

|  | Standardized<br>coefficient | Standard error | *p* |
| --- | --- | --- | --- |
| Age | 0.263 | 2.228 | **0.003** |
| Sex | 0.008 | 1.660 | 0.454 |
| Kinship | 0.156 | 3.667 | **0.026** |
| Reproductive<br>status | 0.154 | 1.997 | 0.062 |
| Manual<br>indicative | 0.370 | 1.763 | **0.007** |
| Manual non-indicative | 0.081 | 0.295 | 0.144 |

**Table S12.10.**
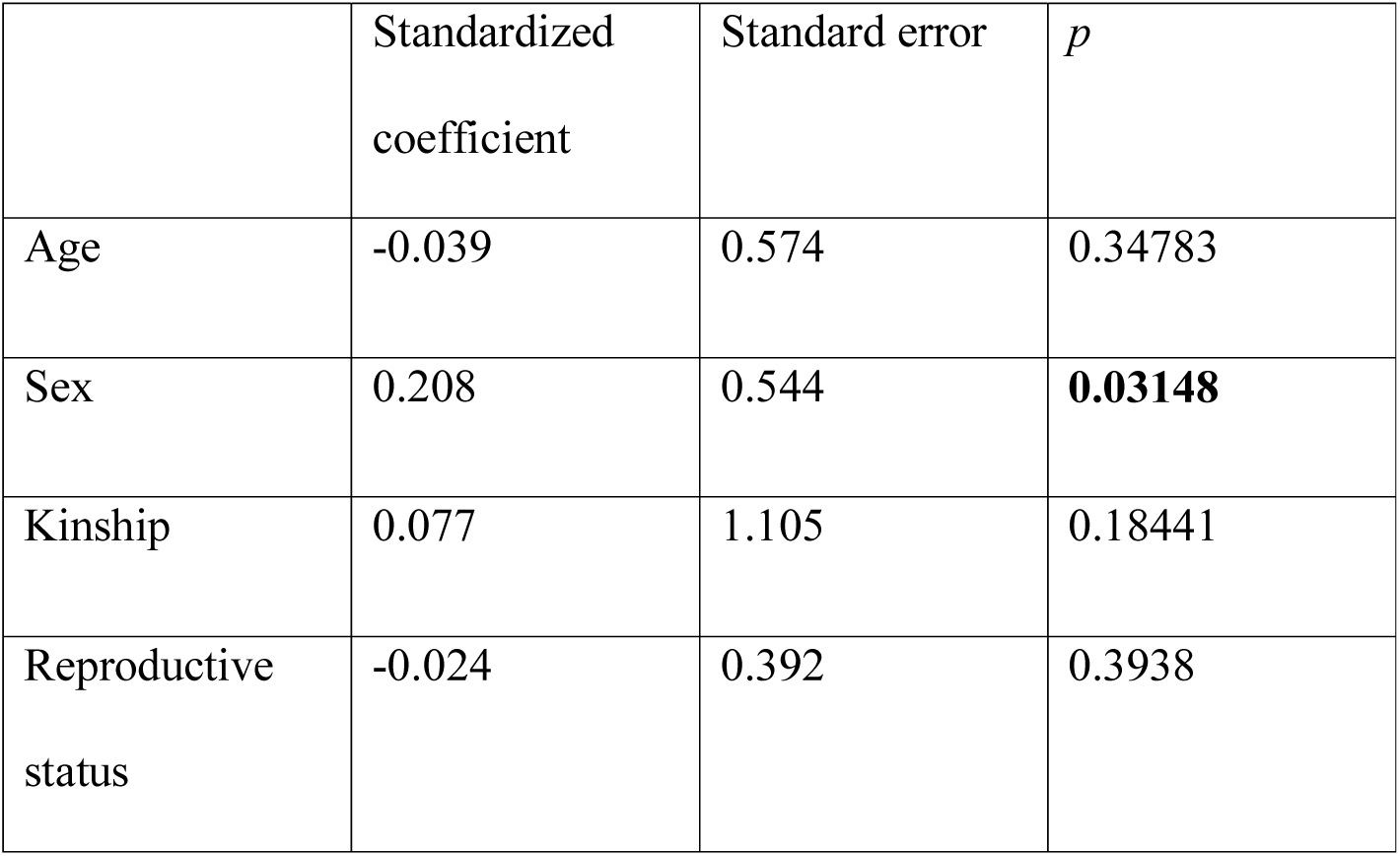
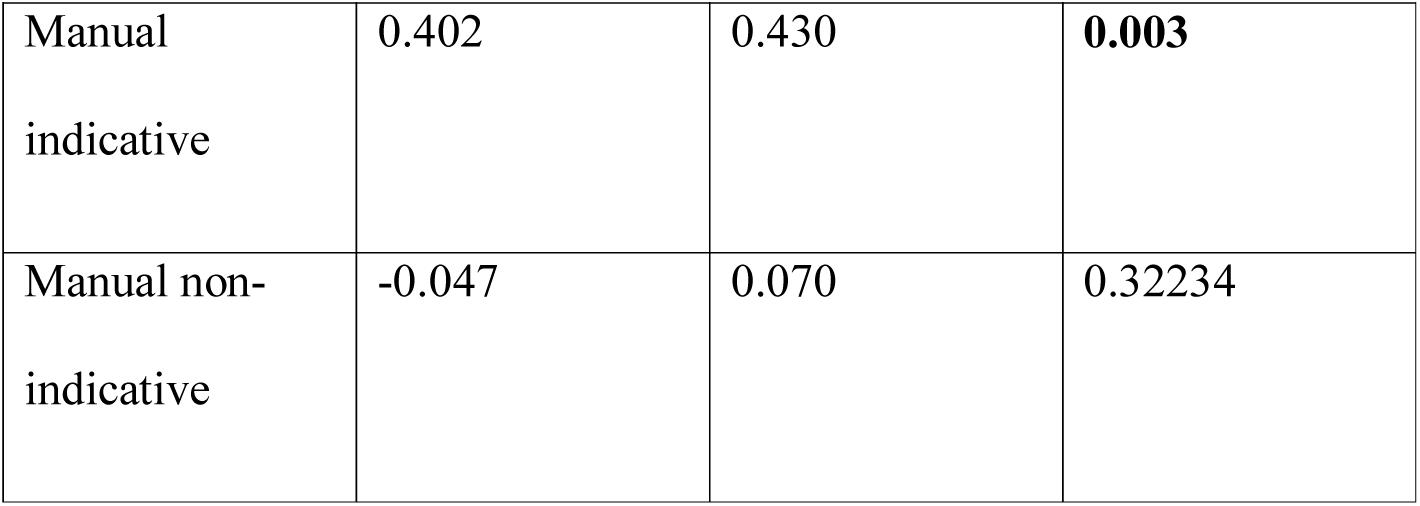
Rate of scratch produced (*r^2^*= 0.193)

**Table S12.11.**
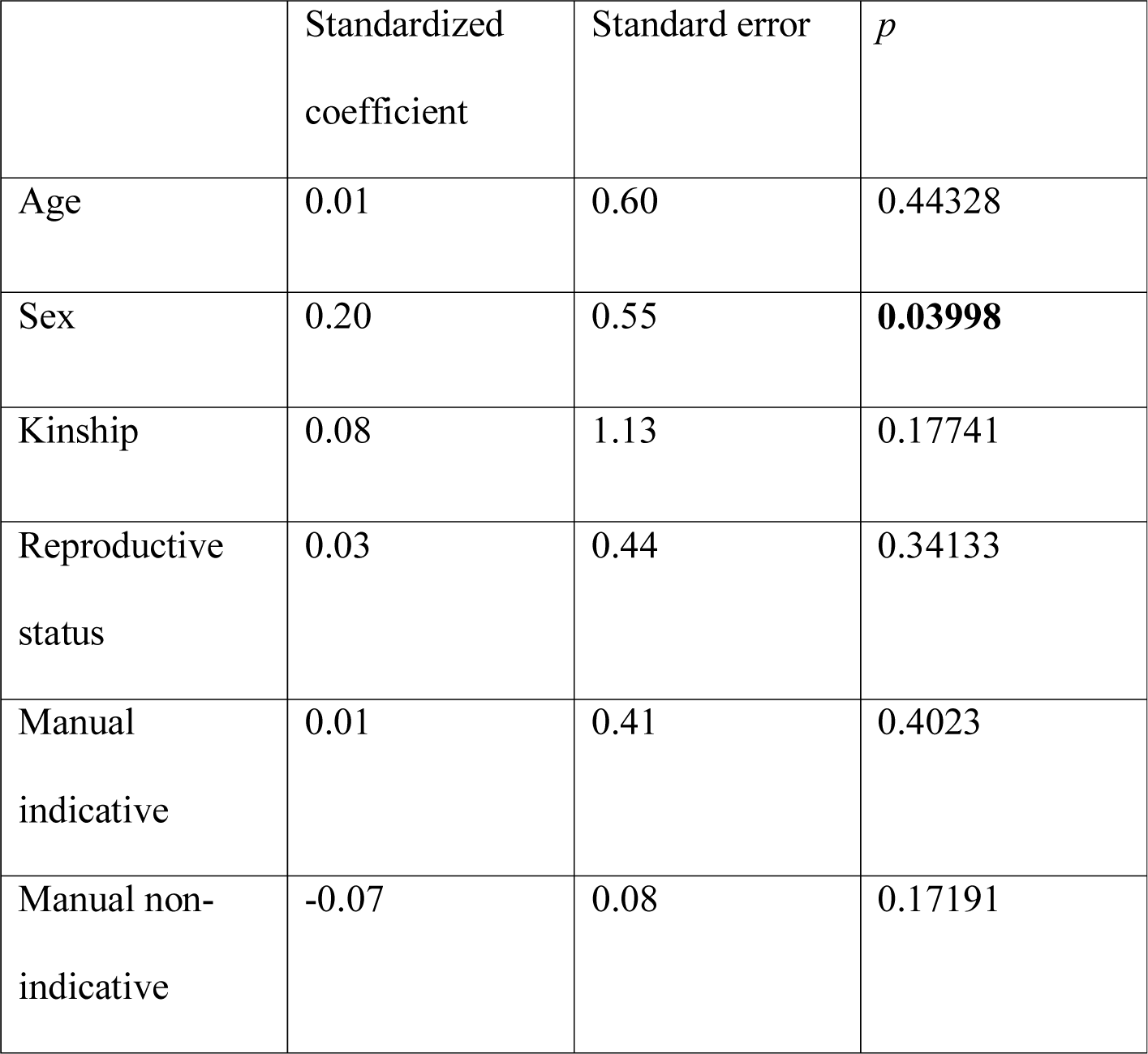
Rate of scratch received (*r^2^* = 0.051)

#### Gestures made at close and far proximity

Supplementary Table S13. MRQAP regression models predicting durations of social behavior, per hour dyad spent within 10m. Predictor variables were rates of gestures made at close and far proximity between the recipient and the signaller, per hour dyad spent within 10m and demographic variables. Based on 132 chimpanzee dyads. Significant *p* values are indicated in bold. R squared (*r^2^)* denotes amount of variance in the dependent variable explained by the regression model.

**Table S13.1.** Duration of joint feeding behaviour (*r^2^*= 0.156)

|  | Standardized<br>coefficient | Standard error | *p* |
| --- | --- | --- | --- |
| Age | -0.026 | 0.641 | 0.409 |
| Sex | 0.142 | 0.576 | 0.092 |
| Kinship | 0.282 | 1.250 | **0.006** |
| Reproductive status | 0.251 | 0.557 | **0.003** |
| Close proximity | 0.055 | 0.053 | 0.223 |
| Far proximity | -0.065 | 0.044 | 0.203 |

**Table S13.2.** Duration of joint resting behaviour (*r^2^* = 0.082)

|  | Standardized<br>coefficient | Standard error | *p* |
| --- | --- | --- | --- |
| Age | 0.284 | 1.421 | **0.005** |
| Sex | -0.131 | 1.125 | 0.055 |
| Kinship | -0.025 | 2.482 | 0.423 |
| Reproductive<br>status | -0.040 | 1.124 | 0.365 |
| Close proximity | 0.121 | 0.123 | **0.046** |
| Far proximity | -0.048 | 0.112 | 0.235 |

**Table S13.3.** Duration of joint travelling behaviour (*r^2^* = 0.241)

|  | Standardized<br>coefficient | Standard error | *p* |
| --- | --- | --- | --- |
| Age | 0.225 | 0.360 | **0.009** |
| Sex | 0.013 | 0.309 | 0.445 |
| Kinship | 0.016 | 0.660 | 0.314 |
| Reproductive<br>status | -0.059 | 0.275 | 0.248 |
| Close proximity | 0.273 | 0.030 | **0.015** |
| Far proximity | 0.223 | 0.026 | **0.013** |

**Table S13.4.** Duration of giving grooming (*r^2^* = 0.589)

|  | Standardized<br>coefficient | Standard error | *p* |
| --- | --- | --- | --- |
| Age | 0.128 | 0.346 | **0.018** |
| Sex | -0.042 | 0.282 | 0.242 |
| Kinship | 0.085 | 0.625 | 0.068 |
| Reproductive<br>status | 0.022 | 0.274 | 0.383 |
| Close proximity | 0.765 | 0.055 | **0.001** |
| Far proximity | -0.112 | 0.029 | **0.012** |

**Table S13.5.** Duration of mutual grooming (*r^2^* = 0.288)

|  | Standardized<br>coefficient | Standard error | *p* |
| --- | --- | --- | --- |
| Age | 0.125 | 0.604 | 0.101 |
| Sex | -0.136 | 0.518 | 0.073 |
| Kinship | 0.022 | 1.067 | 0.263 |
| Reproductive<br>status | 0.095 | 0.499 | 0.157 |
| Close proximity | 0.387 | 0.052 | **0.004** |
| Far proximity | 0.223 | 0.044 | **0.018** |

**Table S13.6.** Duration of receiving grooming (*r^2^* = 0.074)

|  | Standardized<br>coefficient | Standard error | *p* |
| --- | --- | --- | --- |
| Age | -0.117 | 0.495 | 0.099 |
| Sex | 0.109 | 0.400 | 0.128 |
| Kinship | 0.001 | 0.897 | 0.352 |
| Reproductive<br>status | 0.132 | 0.437 | 0.069 |
| Close proximity | 0.153 | 0.044 | 0.083 |
| Far proximity | 0.053 | 0.039 | 0.167 |

**Table S13.7.** Duration of visual attention towards dyad partner (*r^2^* = 0.439)

|  | Standardized<br>coefficient | Standard error | *p* |
| --- | --- | --- | --- |
| Age | 0.026 | 1.152 | 0.368 |
| Sex | 0.020 | 0.935 | 0.386 |
| Kinship | 0.181 | 2.133 | **0.009** |
| Reproductive<br>status | 0.242 | 1.179 | **0.010** |
| Close proximity | 0.500 | 0.119 | **0.001** |
| Far proximity | 0.131 | 0.086 | **0.045** |

**Table S13.8.** Duration of visual attention away from dyad partner (*r^2^*= 0.190)

|  | Standardized<br>coefficient | Standard error | *p* |
| --- | --- | --- | --- |
| Age | 0.341 | 1.683 | **0.001** |
| Sex | -0.102 | 1.258 | 0.105 |
| Kinship | 0.062 | 2.737 | 0.144 |
| Reproductive<br>status | -0.005 | 1.292 | 0.481 |
| Close proximity | 0.287 | 0.147 | **0.021** |
| Far proximity | -0.066 | 0.116 | 0.158 |

**Table S13.9.** Duration of time in close proximity – within 2 m (*r^2^* = 0.376)

|  | Standardized<br>coefficient | Standard error | *p* |
| --- | --- | --- | --- |
| Age | 0.249 | 2.163 | **0.004** |
| Sex | -0.058 | 1.643 | 0.226 |
| Kinship | 0.145 | 3.680 | **0.035** |
| Reproductive<br>status | 0.134 | 2.103 | 0.094 |
| Close proximity | 0.480 | 0.213 | **0.001** |
| Far proximity | 0.029 | 0.155 | 0.297 |

**Table S13.10.** Rate of scratch produced (*r^2^* = 0.174)

|  | Standardized<br>coefficient | Standard error | *p* |
| --- | --- | --- | --- |
| Age | -0.032 | 0.582 | 0.3903 |
| Sex | 0.175 | 0.539 | 0.05697 |
| Kinship | 0.089 | 1.120 | 0.15192 |
| Reproductive<br>status | -0.007 | 0.445 | 0.47126 |
| Close proximity | 0.210 | 0.051 | **0.02399** |
| Far proximity | 0.244 | 0.045 | **0.01299** |

**Table S13.11.** Rate of scratch received (*r^2^* = 0.058)

|  | Standardized<br>coefficient | Standard error | *p* |
| --- | --- | --- | --- |
| Age | 0.03 | 0.66 | 0.3908 |
| Sex | 0.22 | 0.58 | **0.03598** |
| Kinship | 0.08 | 1.19 | 0.18691 |
| Reproductive<br>status | 0.05 | 0.45 | 0.27236 |
| Close proximity | -0.11 | 0.05 | **0.04898** |
| Far proximity | 0.01 | 0.05 | 0.38131 |

#### Non-repetitive and repetitive gestures

Supplementary Table S14. MRQAP regression models predicting durations of social behavior, per hour dyad spent within 10m. Predictor variables were rates of non-repetitive and repetitive gestures between the recipient and the signaller, per hour dyad spent within 10m and demographic variables. Based on 132 chimpanzee dyads. Significant *p* values are indicated in bold. R squared (*r^2^)* denotes amount of variance in the dependent variable explained by the regression model.

**Table S14.1.**
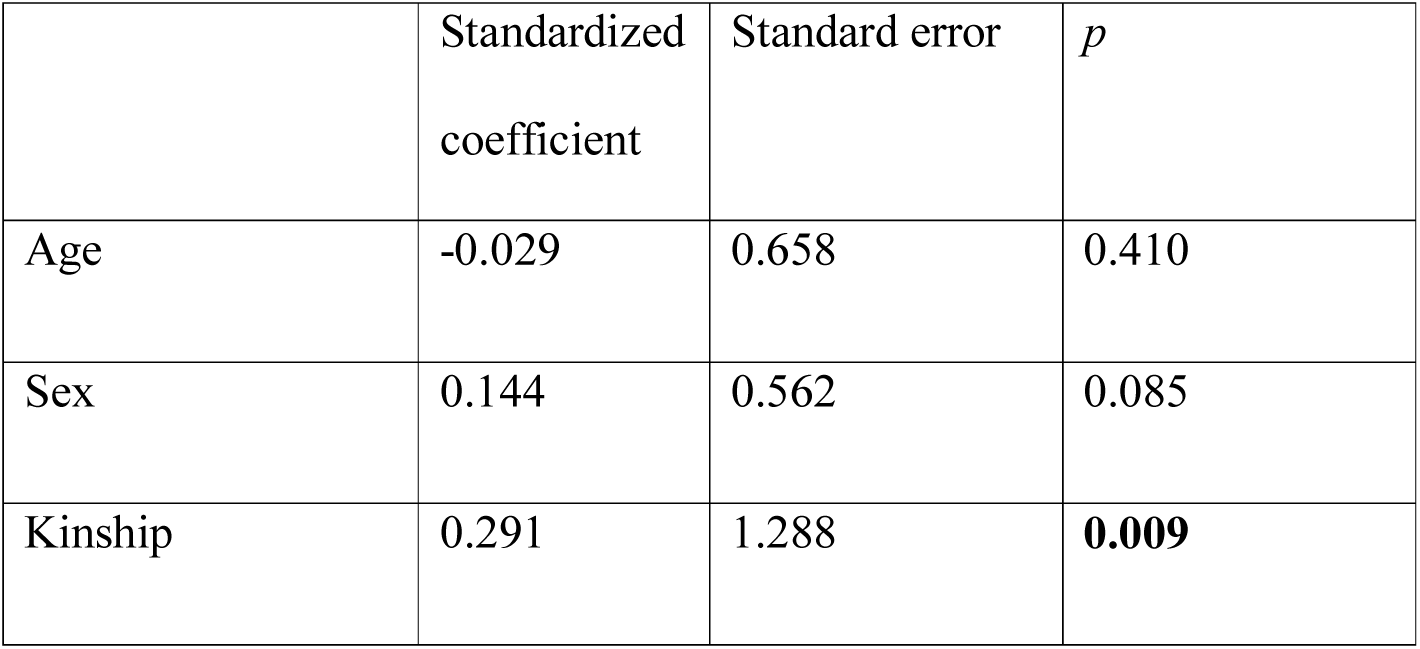
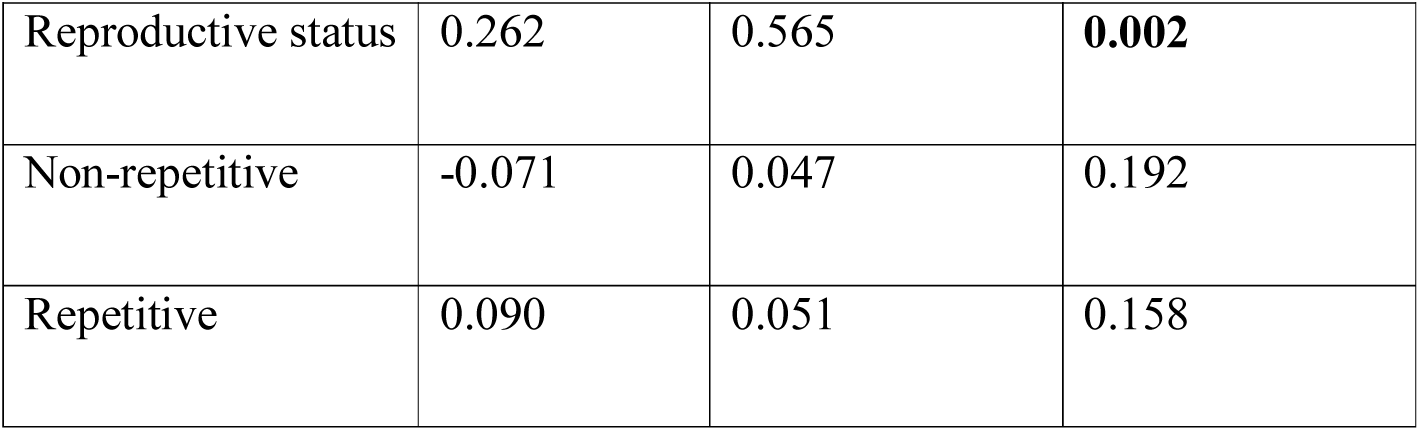
Duration of joint feeding behaviour (*r^2^* = 0.158)

**Table S14.2.**
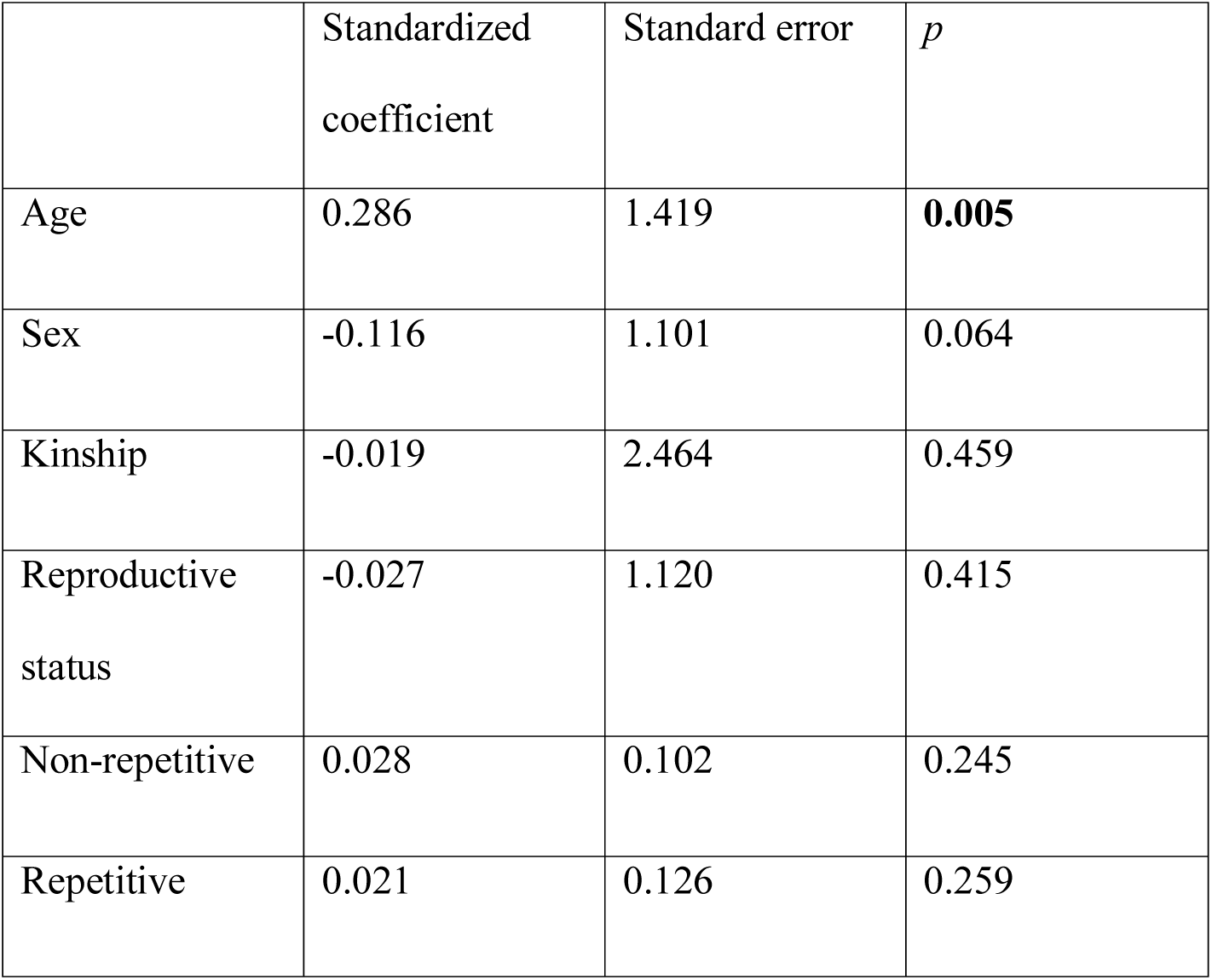
Duration of joint resting behaviour (*r^2^* = 0.072)

|  | Standardized coefficient | Standard error | *p* |
| --- | --- | --- | --- |
| Age | 0.286 | 1.419 | **0.005** |
| Sex | -0.116 | 1.101 | 0.064 |
| Kinship | -0.019 | 2.464 | 0.459 |
| Reproductive status | -0.027 | 1.120 | 0.415 |
| Non-repetitive | 0.028 | 0.102 | 0.245 |
| Repetitive | 0.021 | 0.126 | 0.259 |

**Table S14.3.** Duration of joint travelling behaviour (*r^2^* = 0.353)

|  | Standardized coefficient | Standard error | *p* |
| --- | --- | --- | --- |
| Age | 0.222 | 0.309 | **0.004** |
| Sex | 0.054 | 0.250 | 0.234 |
| Kinship | 0.002 | 0.551 | 0.391 |
| Reproductive status | -0.080 | 0.215 | 0.138 |
| Non-repetitive | 0.559 | 0.028 | **0.006** |
| Repetitive | -0.092 | 0.028 | 0.065 |

**Table S14.4.**
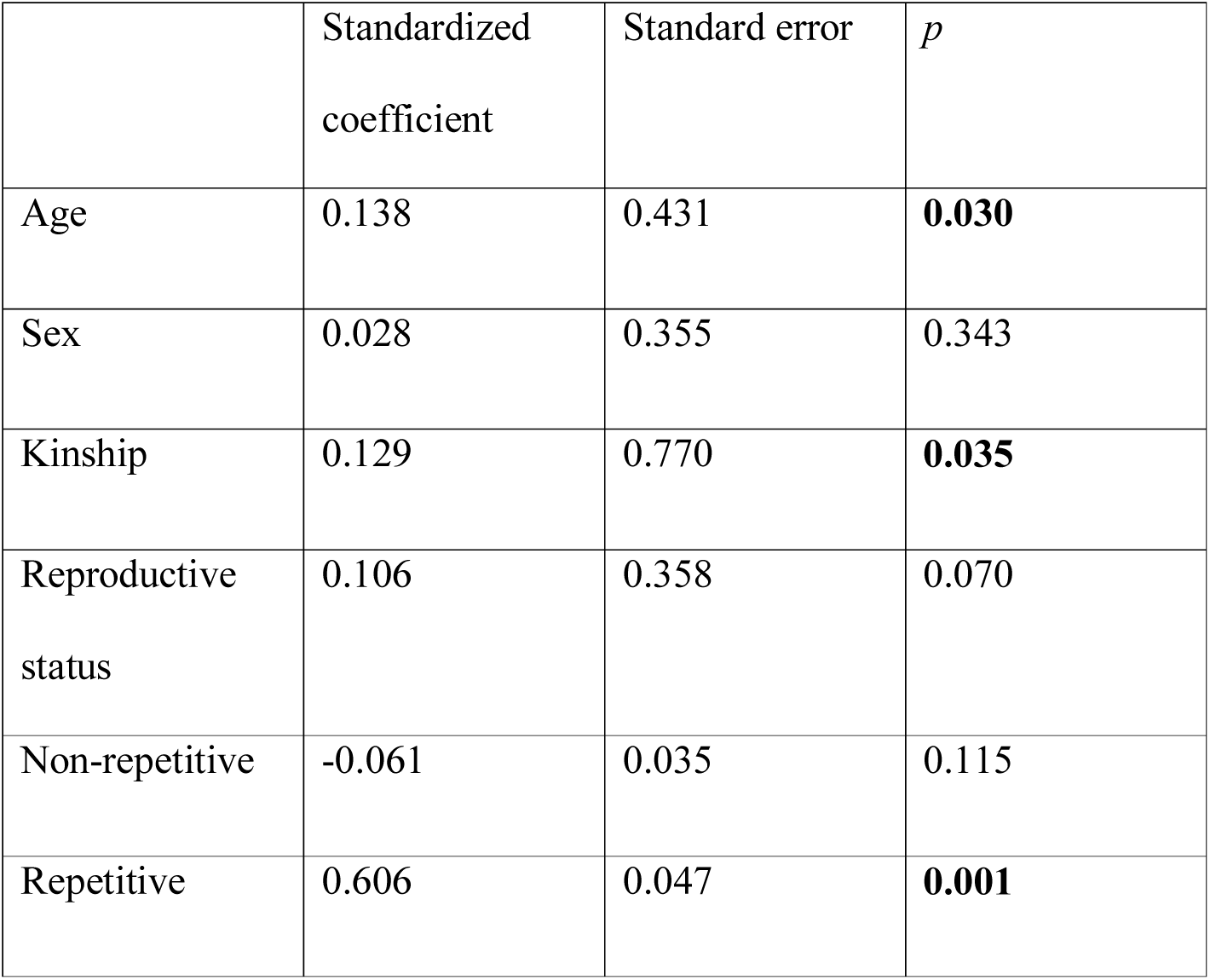
Duration of giving grooming (*r^2^* = 0.409)

**Table S14.5.** Duration of mutual grooming (*r^2^* = 0.470)

|  | Standardized coefficient | Standard error | *p* |
| --- | --- | --- | --- |
| Age | 0.126 | 0.509 | 0.059 |
| Sex | -0.074 | 0.420 | 0.186 |
| Kinship | 0.005 | 0.917 | 0.371 |
| Reproductive status | 0.077 | 0.424 | 0.181 |
| Non-repetitive | 0.726 | 0.052 | **0.001** |
| Repetitive | -0.189 | 0.045 | **0.004** |

**Table S14.6.** Duration of receiving grooming (*r^2^* = 0.080)

|  | Standardized coefficient | Standard error | *p* |
| --- | --- | --- | --- |
| Age | -0.115 | 0.503 | 0.094 |
| Sex | 0.131 | 0.424 | 0.088 |
| Kinship | -0.002 | 0.858 | 0.654 |
| Reproductive status | 0.133 | 0.436 | 0.065 |
| Non-repetitive | 0.207 | 0.039 | **0.035** |
| Repetitive | -0.040 | 0.045 | 0.424 |

**Table S14.7.** Duration of visual attention towards dyad partner (*r^2^* = 0.464)

|  | Standardized coefficient | Standard error | *p* |
| --- | --- | --- | --- |
| Age | 0.028 | 1.143 | 0.376 |
| Sex | 0.088 | 0.950 | 0.141 |
| Kinship | 0.179 | 2.173 | **0.032** |
| Reproductive status | 0.249 | 1.127 | **0.002** |
| Non-repetitive | 0.567 | 0.104 | **0.004** |
| Repetitive | -0.003 | 0.090 | 0.543 |

**Table S14.8.**
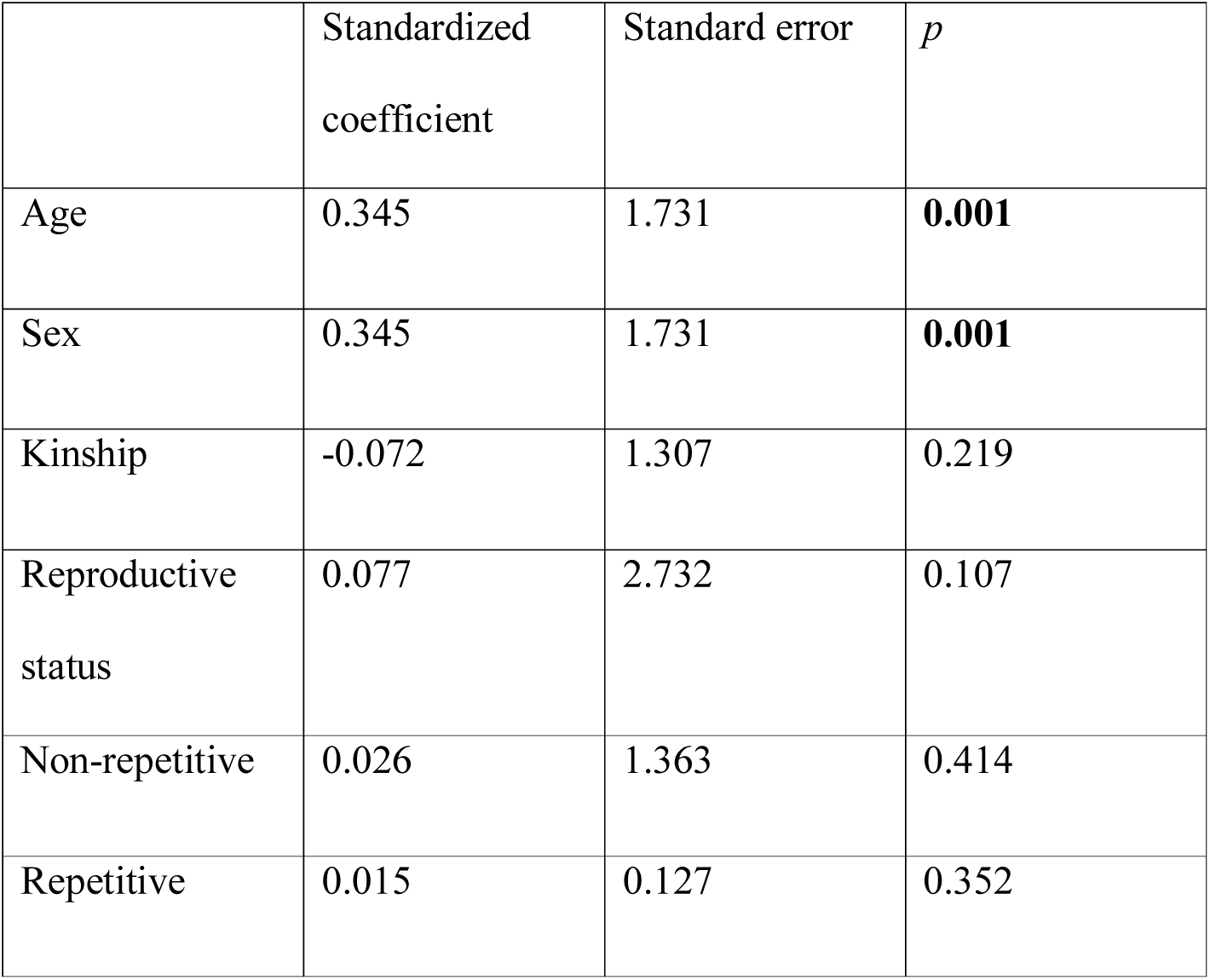
Duration of visual attention away from dyad partner (*r^2^* = 0.149)

**Table S14.9.** Duration of time in close proximity – within 2 m (*r^2^* = 0.307)

|  | Standardized coefficient | Standard error | *p* |
| --- | --- | --- | --- |
| Age | 0.253 | 2.241 | **0.004** |
| Sex | 0.000 | 1.681 | 0.483 |
| Kinship | 0.154 | 3.809 | **0.033** |
| Reproductive status | 0.159 | 2.109 | 0.067 |
| Non-repetitive | 0.331 | 0.177 | **0.012** |
| Repetitive | 0.111 | 0.193 | 0.118 |

**Table S14.10.** Rate of scratch produced (*r^2^* = 0.215)

|  | Standardized coefficient | Standard error | *p* |
| --- | --- | --- | --- |
| Age | -0.0346 | 0.5810 | 0.37881 |
| Sex | 0.2024 | 0.5279 | **0.03248** |
| Kinship | 0.0765 | 1.1127 | 0.1919 |
| Reproductive status | -0.0259 | 0.4175 | 0.38731 |
| Non-repetitive | 0.4196 | 0.0419 | **0.0045** |
| Repetitive | -0.0013 | 0.0470 | 0.55822 |

**Table S14.11.** Rate of scratch received (r2 = 0.060)

|  | Standardized coefficient | Standard error | *p* |
| --- | --- | --- | --- |
| Age | 0.03 | 0.63 | 0.38731 |
| Sex | 0.21 | 0.54 | **0.02499** |
| Kinship | 0.07 | 1.19 | 0.1959 |
| Reproductive status | 0.03 | 0.45 | 0.34633 |
| Non-repetitive | 0.05 | 0.05 | 0.22239 |
| Repetitive | -0.14 | 0.05 | **0.04498** |

#### Heterogeneous and homogeneous gestures

Supplementary Table S15. MRQAP regression models predicting durations of social behavior, per hour dyad spent within 10m. Predictor variables were rates of heterogeneous and homogeneous gestures between the recipient and the signaller, per hour dyad spent within 10m and demographic variables. Based on 132 chimpanzee dyads. Significant *p* values are indicated in bold. R squared (*r^2^)* denotes amount of variance in the dependent variable explained by the regression model.

**Table S15.1.** Duration of joint feeding behaviour (*r^2^*= 0.)

|  | Standardized coefficient | Standard error | *p* |
| --- | --- | --- | --- |
| Age | -0.02 | 0.66 | 0.41629 |
| Sex | 0.15 | 0.58 | 0.08346 |
| Kinship | 0.29 | 1.29 | **0.01199** |
| Reproductive status | 0.26 | 0.55 | **0.002** |
| Heterogeneous | 0.00 | 0.08 | 0.43328 |
| Homogeneous | -0.01 | 0.04 | 0.53173 |

**Table S15.2.**
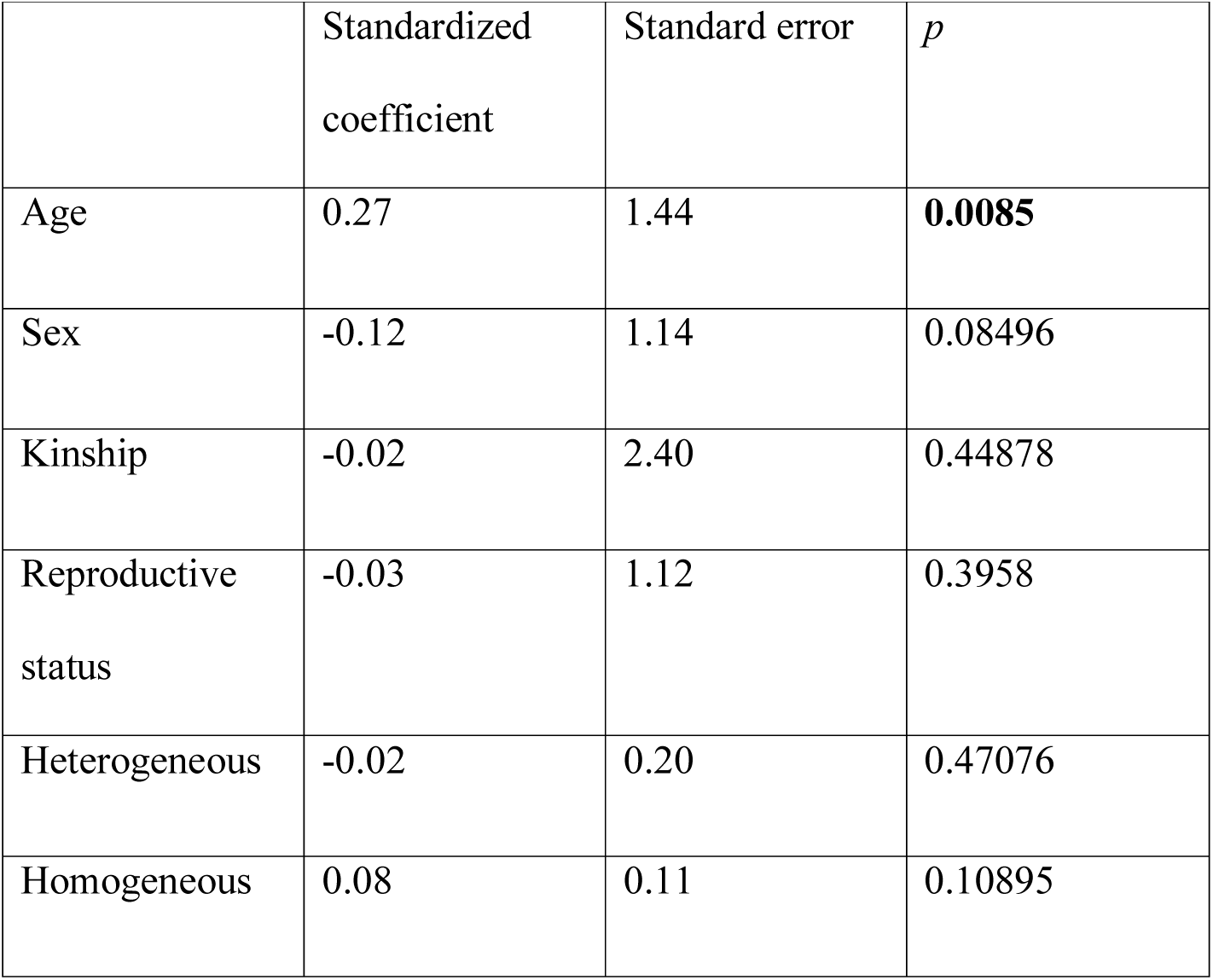
Duration of joint resting behaviour (*r^2^* = 0.075)

**Table S15.3.** Duration of joint travelling behaviour (*r^2^* = 0.251)

|  | Standardized<br>coefficient | Standard error | *p* |
| --- | --- | --- | --- |
| Age | 0.17 | 0.36 | **0.04048** |
| Sex | 0.01 | 0.29 | 0.44428 |
| Kinship | 0.01 | 0.60 | 0.33633 |
| Reproductive<br>status | -0.07 | 0.27 | 0.2019 |
| Heterogeneous | -0.03 | 0.05 | 0.38281 |
| Homogeneous | 0.43 | 0.03 | **0.01049** |

**Table S15.4.** Duration of giving grooming (*r^2^* = 0.364)

|  | Standardized<br>coefficient | Standard error | *p* |
| --- | --- | --- | --- |
| Age | 0.09 | 0.47 | 0.12944 |
| Sex | 0.02 | 0.39 | 0.3953 |
| Kinship | 0.11 | 0.84 | 0.06697 |
| Reproductive<br>status | 0.08 | 0.37 | 0.14543 |
| Heterogeneous | 0.11 | 0.07 | 0.06747 |
| Homogeneous | 0.52 | 0.04 | **0.004** |

**Table S15.5.** Duration of mutual grooming (*r^2^* = 0.358)

|  | Standardized<br>coefficient | Standard error | *p* |
| --- | --- | --- | --- |
| Age | 0.03 | 0.56 | 0.34533 |
| Sex | -0.14 | 0.49 | 0.08046 |
| Kinship | 0.01 | 1.05 | 0.33633 |
| Reproductive<br>status | 0.08 | 0.46 | 0.16392 |
| Heterogeneous | -0.15 | 0.08 | **0.01649** |
| Homogeneous | 0.60 | 0.05 | **0.0005** |

**Table S15.6.**
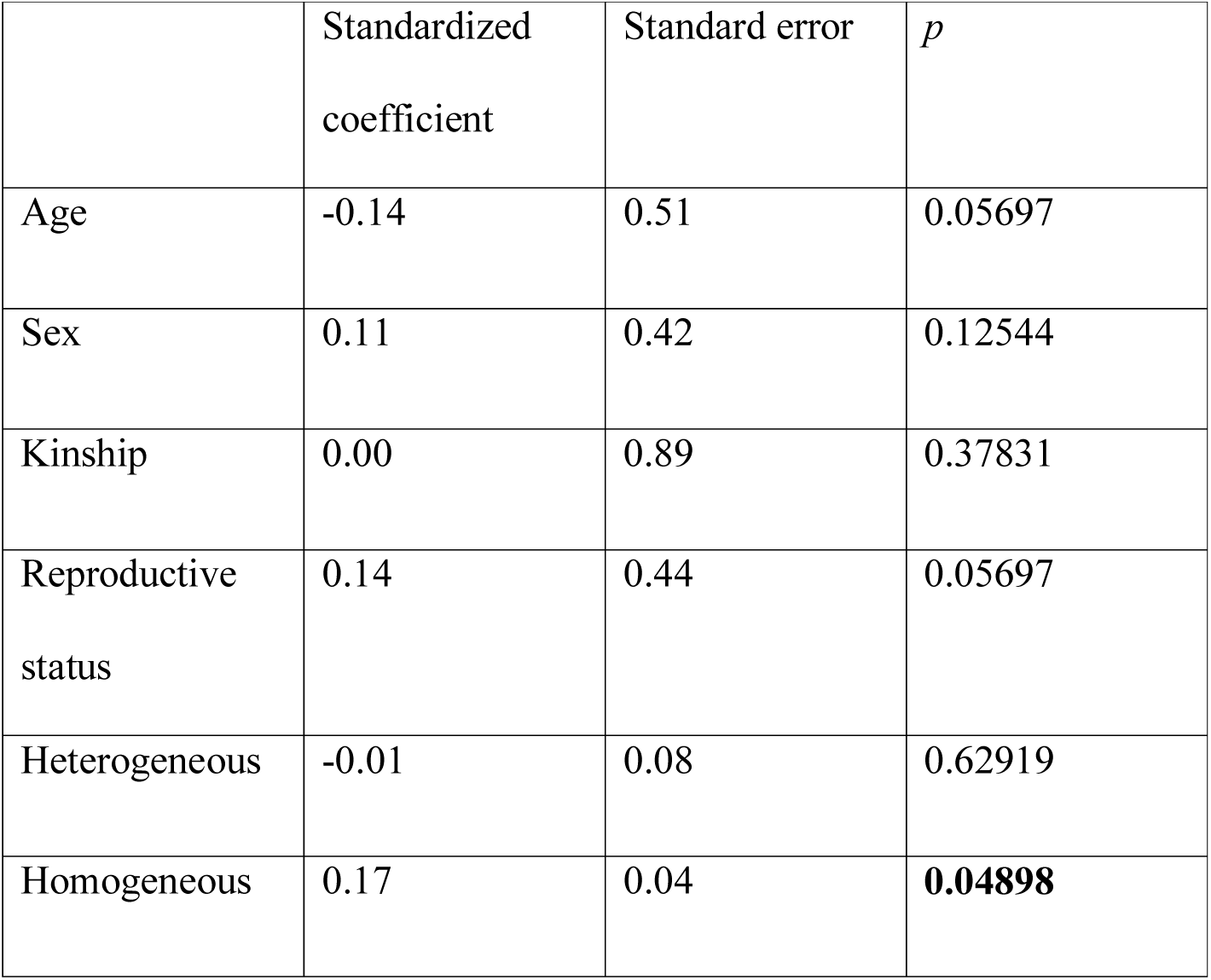
Duration of receiving grooming (*r^2^* = 0.072)

**Table S15.7.**
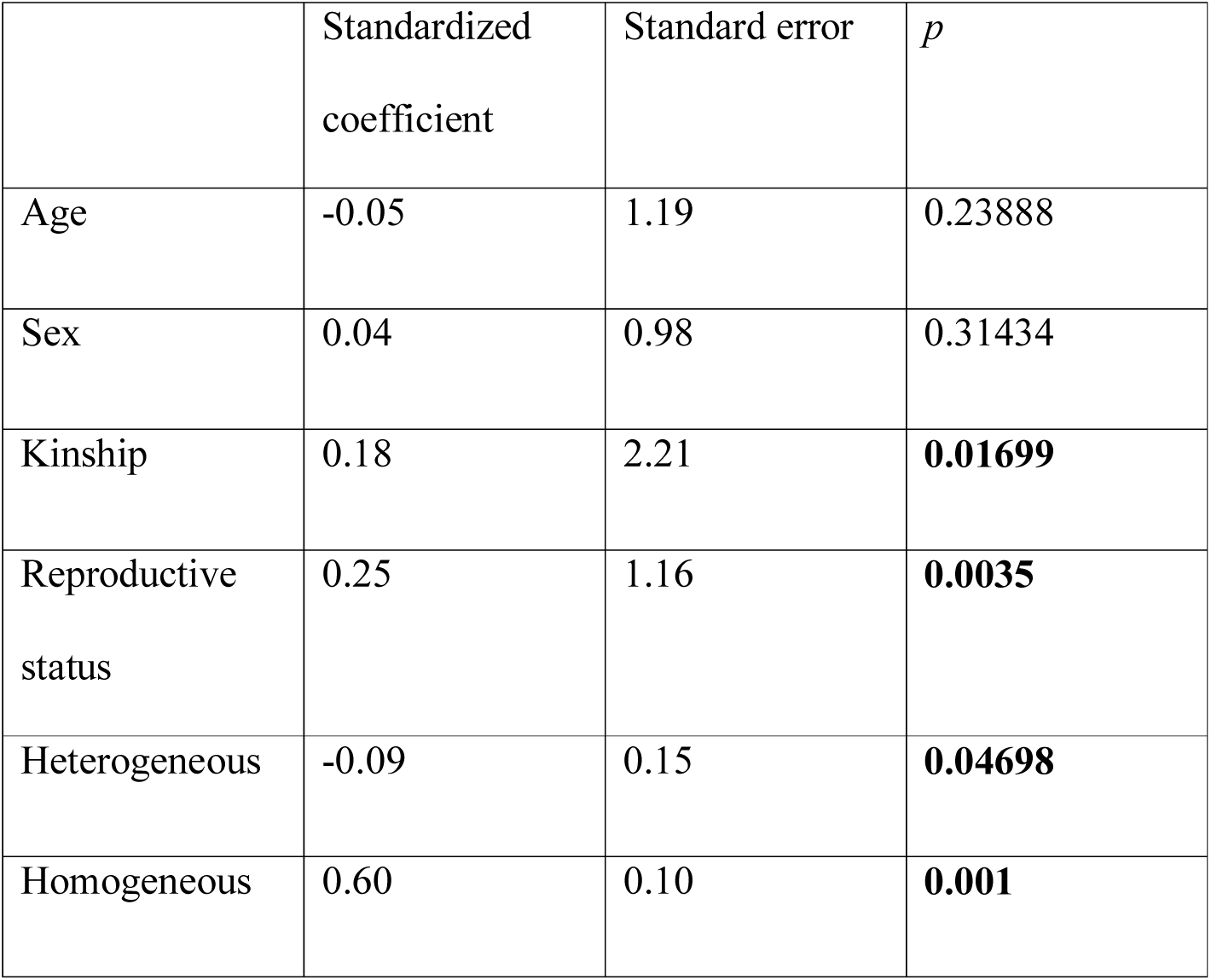
Duration of visual attention towards dyad partner (*r^2^* = 0.455)

**Table S15.8.** Duration of visual attention away from dyad partner (*r^2^* = 0.151)

|  | Standardized<br>coefficient | Standard error | *p* |
| --- | --- | --- | --- |
| Age | 0.33 | 1.77 | **0.0035** |
| Sex | -0.08 | 1.31 | 0.1924 |
| Kinship | 0.07 | 2.85 | 0.13793 |
| Reproductive<br>status | 0.02 | 1.32 | 0.44478 |
| Heterogeneous | 0.04 | 0.23 | 0.21289 |
| Homogeneous | 0.17 | 0.12 | **0.04348** |

**Table S15.9.** Duration of time in close proximity – within 2 m (*r^2^* = 0.335)

|  | Standardized<br>coefficient | Standard error | *p* |
| --- | --- | --- | --- |
| Age | 0.20 | 2.25 | **0.01899** |
| Sex | -0.03 | 1.67 | 0.35082 |
| Kinship | 0.15 | 3.93 | **0.03798** |
| Reproductive<br>status | 0.15 | 2.08 | 0.06597 |
| Heterogeneous | -0.03 | 0.30 | 0.4013 |
| Homogeneous | 0.45 | 0.18 | **0.0005** |

**Table S15.10.** Rate of scratch produced (*r^2^* = 0.186)

|  | Standardized<br>coefficient | Standard error | *p* |
| --- | --- | --- | --- |
| Age | -0.067 | 0.603 | 0.25437 |
| Sex | 0.169 | 0.540 | 0.07046 |
| Kinship | 0.083 | 1.099 | 0.17341 |
| Reproductive<br>status | -0.019 | 0.436 | 0.42579 |
| Heterogeneous | 0.053 | 0.077 | 0.21939 |
| Homogeneous | 0.378 | 0.042 | **0.0025** |

**Table S15.11.** Rate of scratch received (*r^2^* = 0.054)

|  | Standardized<br>coefficient | Standard error | *p* |
| --- | --- | --- | --- |
| Age | 0.03 | 0.65 | 0.36182 |
| Sex | 0.21 | 0.57 | **0.03748** |
| Kinship | 0.08 | 1.19 | 0.17141 |
| Reproductive<br>status | 0.04 | 0.45 | 0.29985 |
| Heterogeneous | -0.03 | 0.09 | 0.4008 |
| Homogeneous | -0.08 | 0.04 | 0.16142 |

#### Gestures accompanied by penile erection

Supplementary Table S16. MRQAP regression models predicting durations of social behavior, per hour dyad spent within 10m. Predictor variables were rates of gestures between the recipient and the signaller are accompanied by penile erection, per hour dyad spent within 10m and demographic variables. Based on 132 chimpanzee dyads. Significant *p* values are indicated in bold. R squared (*r^2^)* denotes amount of variance in the dependent variable explained by the regression model.

**Table S16.1.** Duration of joint feeding behaviour (*r^2^* = 0.153)

|  | Standardized coefficient | Standard error | *p* |
| --- | --- | --- | --- |
| Age | -0.03 | 0.64 | 0.42429 |
| Sex | 0.16 | 0.58 | 0.07646 |
| Kinship | 0.29 | 1.28 | **0.007** |
| Reproductive status | 0.26 | 0.54 | **0.0035** |
| Penile erection | 0.04 | 0.22 | 0.27036 |

**Table S16.2.** Duration of joint resting behaviour (*r^2^*= 0.070)

|  | Standardized coefficient | Standard error | *p* |
| --- | --- | --- | --- |
| Age | 0.29 | 1.40 | **0.002** |
| Sex | -0.12 | 1.12 | 0.08696 |
| Kinship | -0.02 | 2.49 | 0.45727 |
| Reproductive status | -0.02 | 1.10 | 0.4018 |
| Penile erection | -0.01 | 0.49 | 0.5947 |

**Table S16.3.** Duration of joint travelling behaviour (*r^2^* = 0.095)

|  | Standardized coefficient | Standard error | *p* |
| --- | --- | --- | --- |
| Age | 0.28 | 0.43 | **0.005** |
| Sex | 0.06 | 0.34 | 0.27886 |
| Kinship | 0.01 | 0.74 | 0.29885 |
| Reproductive status | -0.02 | 0.33 | 0.44378 |
| Penile erection | 0.07 | 0.14 | 0.13443 |

**Table S16.4.** Duration of giving grooming (*r^2^* = 0.086)

|  | Standardized coefficient | Standard error | *p* |
| --- | --- | --- | --- |
| Age | 0.21 | 0.56 | **0.01899** |
| Sex | 0.06 | 0.45 | 0.26137 |
| Kinship | 0.11 | 0.97 | 0.11444 |
| Reproductive status | 0.14 | 0.50 | 0.06997 |
| Penile erection | 0.02 | 0.20 | 0.27086 |

**Table S16.5.** Duration of mutual grooming (*r^2^* = 0.055)

|  | Standardized coefficient | Standard error | *p* |
| --- | --- | --- | --- |
| Age | 0.19 | 0.74 | 0.07746 |
| Sex | -0.09 | 0.61 | 0.1959 |
| Kinship | 0.01 | 1.22 | 0.28636 |
| Reproductive status | 0.16 | 0.65 | 0.10745 |
| Penile erection | -0.04 | 0.23 | 0.36432 |

**Table S16.6.**
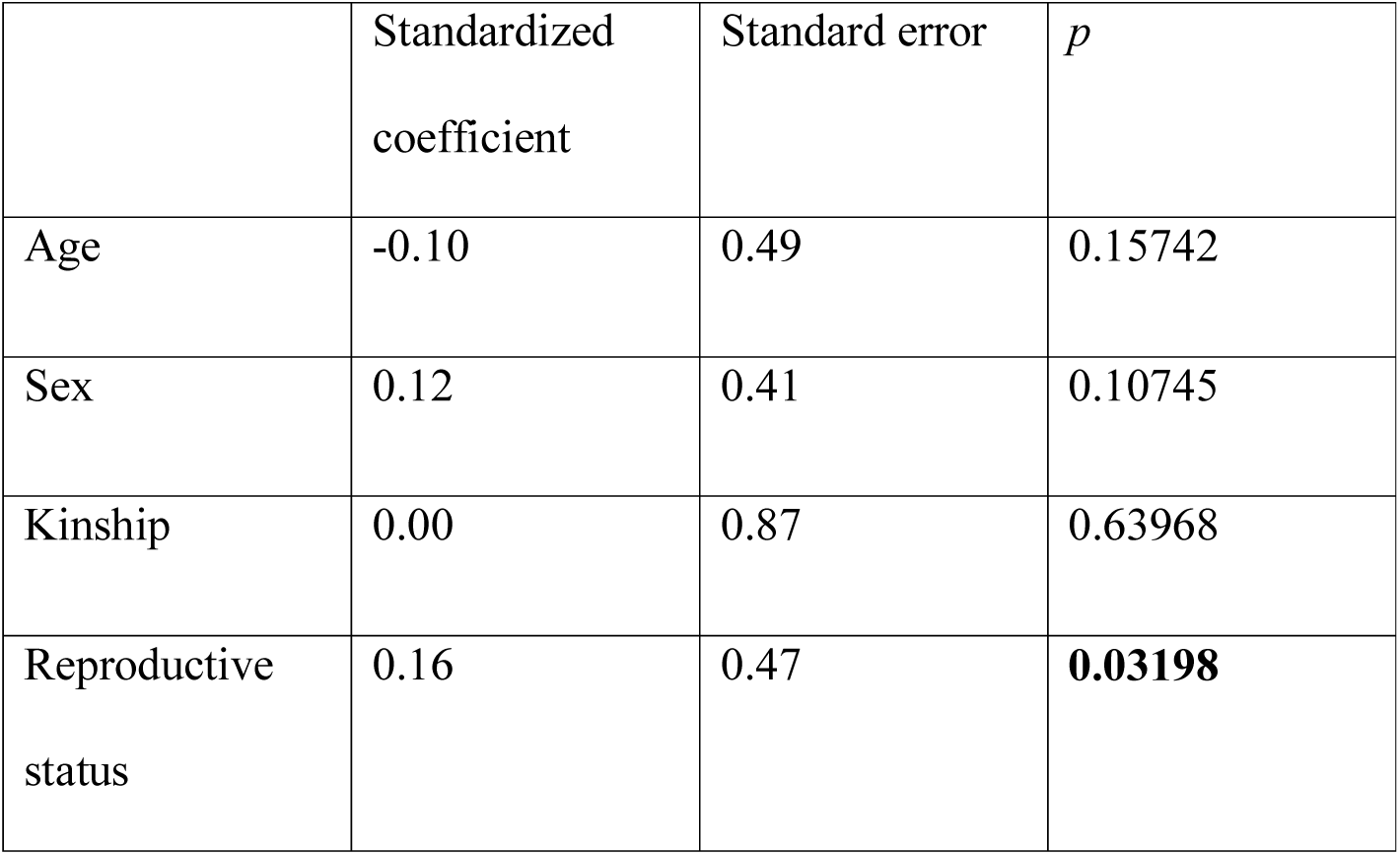
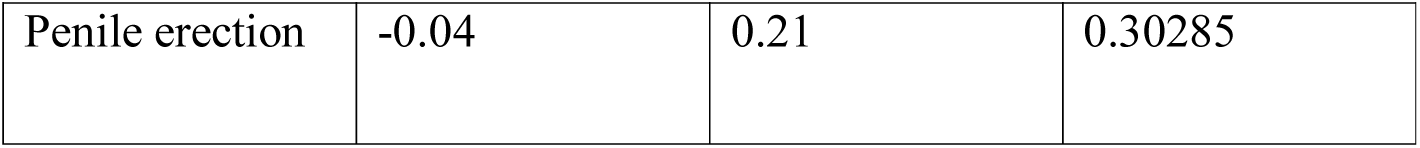
Duration of receiving grooming (*r^2^* = 0.047)

|  | Standardized coefficient | Standard error | *p* |
| --- | --- | --- | --- |
| Age | -0.10 | 0.49 | 0.15742 |
| Sex | 0.12 | 0.41 | 0.10745 |
| Kinship | 0.00 | 0.87 | 0.63968 |
| Reproductive status | 0.16 | 0.47 | **0.03198** |
| Penile erection | -0.04 | 0.21 | 0.30285 |

**Table S16.7.** Duration of visual attention towards dyad partner (*r^2^* = 0.155)

|  | Standardized<br>coefficient | Standard error | *p* |
| --- | --- | --- | --- |
| Age | 0.10 | 1.40 | 0.16492 |
| Sex | 0.08 | 1.15 | 0.18241 |
| Kinship | 0.18 | 2.68 | 0.06297 |
| Reproductive<br>status | 0.33 | 1.66 | **0.0045** |
| Penile erection | -0.01 | 0.47 | 0.50825 |

**Table S16.8.** Duration of visual attention away from dyad partner (*r^2^* = 0.121)

|  | Standardized<br>coefficient | Standard error | *p* |
| --- | --- | --- | --- |
| Age | 0.37 | 1.75 | **0.0015** |
| Sex | -0.06 | 1.29 | 0.25187 |
| Kinship | 0.07 | 2.83 | 0.13493 |
| Reproductive<br>status | 0.04 | 1.39 | 0.36932 |
| Penile erection | -0.01 | 0.62 | 0.56722 |

**Table S16.9.** Duration of time in close proximity – within 2 m (*r^2^* = 0.156)

|  | Standardized<br>coefficient | Standard error | *p* |
| --- | --- | --- | --- |
| Age | 0.31 | 2.55 | **0.001** |
| Sex | 0.00 | 1.80 | 0.45977 |
| Kinship | 0.15 | 4.22 | **0.04248** |
| Reproductive<br>status | 0.21 | 2.68 | 0.05147 |
| Penile erection | -0.01 | 0.80 | 0.52074 |

**Table S16.10.** Rate of scratch produced (*r^2^* = 0.056)

|  | Standardized<br>coefficient | Standard error | *p* |
| --- | --- | --- | --- |
| Age | 0.021 | 0.607 | 0.41429 |
| Sex | 0.218 | 0.558 | **0.02849** |
| Kinship | 0.086 | 1.124 | 0.15942 |
| Reproductive<br>status | 0.018 | 0.431 | 0.42229 |
| Penile erection | 0.102 | 0.222 | 0.11494 |

**Table S16.11.** Rate of scratch received (*r^2^* = 0.046)

|  | Standardized | Standard error | *p* |
| --- | --- | --- | --- |

|  | coefficient |  |  |
| --- | --- | --- | --- |
| Age | 0.02 | 0.61 | 0.44728 |
| Sex | 0.20 | 0.55 | **0.04148** |
| Kinship | 0.08 | 1.13 | 0.18291 |
| Reproductive status | 0.03 | 0.44 | 0.36282 |
| Penile erection | 0.00 | 0.23 | 0.5942 |

#### Gestures accompanied by piloerection

Supplementary Table S17. MRQAP regression models predicting durations of social behavior, per hour dyad spent within 10m. Predictor variables were rates of gestures between the recipient and the signaller are accompanied by piloerection, per hour dyad spent within 10m and demographic variables. Based on 132 chimpanzee dyads. Significant *p* values are indicated in bold. R squared (*r^2^)* denotes amount of variance in the dependent variable explained by the regression model.

**Table S17.1.** Duration of joint feeding behaviour (*r^2^*= 0.153)

|  | Standardized coefficient | Standard error | *p* |
| --- | --- | --- | --- |
| Age | -0.02 | 0.64 | 0.43828 |
| Sex | 0.15 | 0.57 | 0.08396 |
| Kinship | 0.29 | 1.29 | **0.0085** |
| Reproductive status | 0.26 | 0.57 | **0.002** |
| Piloerection | -0.04 | 0.04 | 0.35782 |

**Table S17.2.** Duration of joint resting behaviour (*r^2^* = 0.070)

|  | Standardized<br>coefficient | Standard error | *p* |
| --- | --- | --- | --- |
| Age | 0.30 | 1.38 | **0.004** |
| Sex | -0.12 | 1.11 | 0.08496 |
| Kinship | -0.02 | 2.39 | 0.43878 |
| Reproductive<br>status | -0.02 | 1.13 | 0.43728 |
| Piloerection | -0.02 | 0.10 | 0.49525 |

**Table S17.3.** Duration of joint travelling behaviour (*r^2^* = 0.231)

|  | Standardized<br>coefficient | Standard error | *p* |
| --- | --- | --- | --- |
| Age | 0.22 | 0.38 | **0.01749** |
| Sex | 0.06 | 0.30 | 0.24438 |
| Kinship | 0.02 | 0.65 | 0.28036 |
| Reproductive<br>status | -0.03 | 0.28 | 0.36982 |
| Piloerection | 0.38 | 0.03 | **0.006** |

**Table S17.4.** Duration of giving grooming (*r^2^* = 0.094)

| | Standardized<br>coefficient | Standard error | $p$ |
| --- | --- | --- | --- |
| Age | 0.19 | 0.56 | **0.03098** |
| Sex | 0.06 | 0.44 | 0.24588 |
| Kinship | 0.11 | 1.00 | 0.11994 |
| Reproductive<br>status | 0.14 | 0.49 | 0.06747 |
| Piloerection | 0.09 | 0.04 | 0.10845 |

**Table S17.5.** Duration of mutual grooming (*r^2^* = 0.229)

| | Standardized<br>coefficient | Standard error | $p$ |
| --- | --- | --- | --- |
| Age | 0.13 | 0.63 | 0.10495 |
| Sex | -0.07 | 0.52 | 0.23188 |
| Kinship | 0.03 | 1.10 | 0.24588 |
| Reproductive<br>status | 0.14 | 0.54 | 0.08646 |
| Piloerection | 0.42 | 0.04 | **0.004** |

**Table S17.6.**
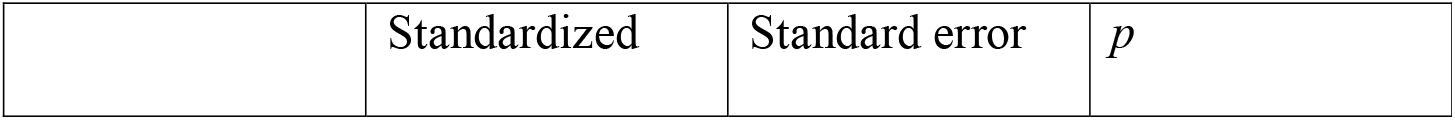
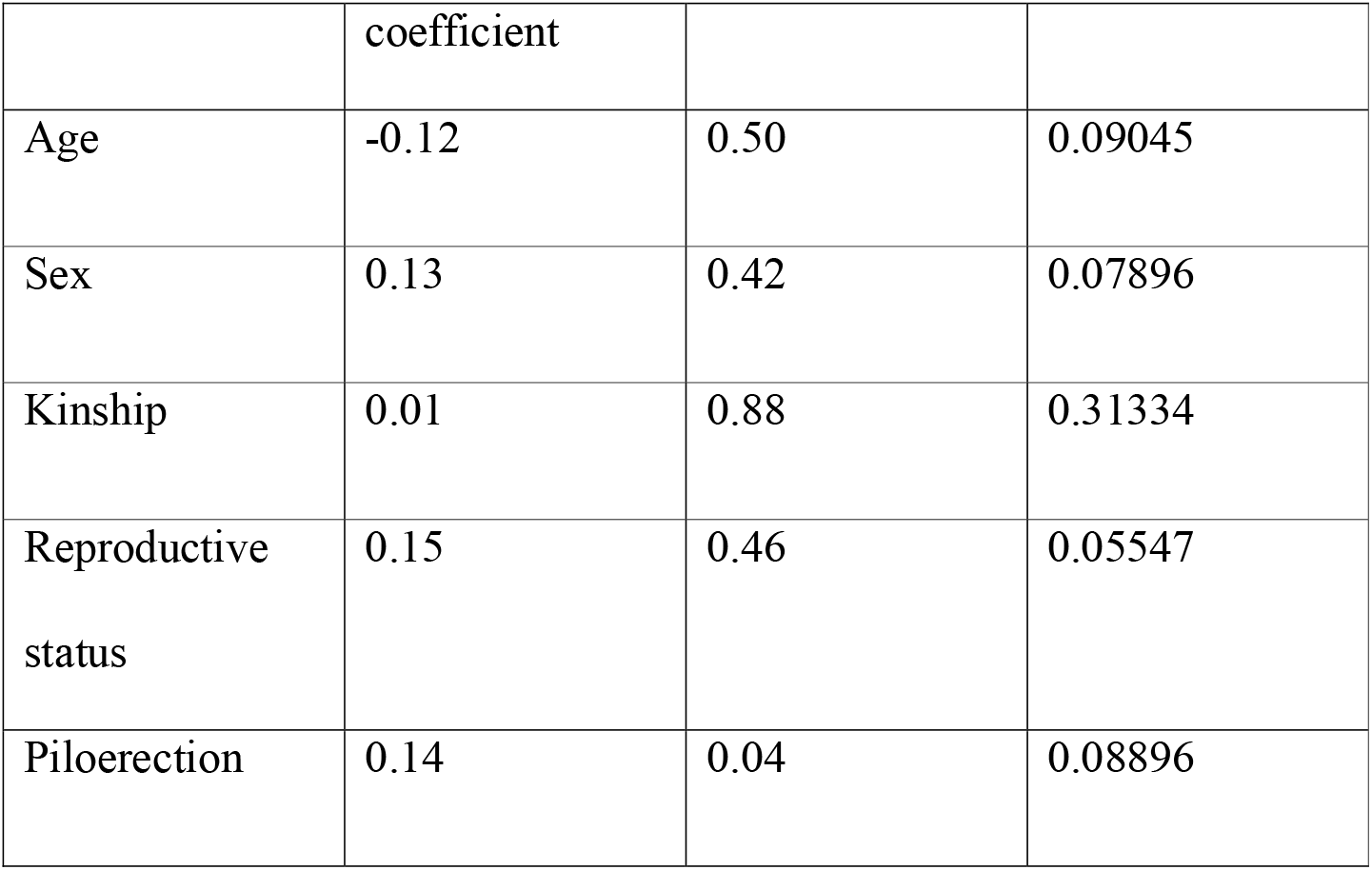
Duration of receiving grooming (*r^2^* = 0.065)

**Table S17.7.** Duration of visual attention towards dyad partner (*r^2^* = 0.278)

|  | Standardized<br>coefficient | Standard error | *p* |
| --- | --- | --- | --- |
| Age | 0.04 | 1.32 | 0.31284 |
| Sex | 0.10 | 1.08 | 0.14193 |
| Kinship | 0.20 | 2.42 | **0.02099** |
| Reproductive<br>status | 0.31 | 1.46 | **0.0015** |
| Piloerection | 0.35 | 0.10 | **0.0055** |

**Table S17.8.** Duration of visual attention away from dyad partner (*r^2^* = 0.121)

|  | Standardized<br>coefficient | Standard error | *p* |
| --- | --- | --- | --- |
| Age | 0.36 | 1.76 | **0.001** |
| Sex | -0.06 | 1.32 | 0.24188 |
| Kinship | 0.07 | 2.87 | 0.13493 |
| Reproductive status | 0.04 | 1.42 | 0.35882 |
| Piloerection | 0.02 | 0.13 | 0.29685 |

**Table S17.9.** Duration of time in close proximity – within 2 m (*r^2^* = 0.200)

|  | Standardized coefficient | Standard error | *p* |
| --- | --- | --- | --- |
| Age | 0.27 | 2.37 | **0.002** |
| Sex | 0.01 | 1.77 | 0.41279 |
| Kinship | 0.16 | 4.15 | **0.03898** |
| Reproductive status | 0.20 | 2.52 | **0.04248** |
| Piloerection | 0.21 | 0.17 | **0.02749** |

**Table S17.10.** Rate of scratch produced (*r^2^* = 0.101)

|  | Standardized coefficient | Standard error | *p* |
| --- | --- | --- | --- |
| Age | -0.020 | 0.618 | 0.43228 |
| Sex | 0.210 | 0.548 | **0.02699** |
| Kinship | 0.088 | 1.150 | 0.15892 |
| Reproductive status | 0.020 | 0.408 | 0.3973 |
| Piloerection | 0.237 | 0.043 | **0.02249** |

**Table S17.11.** Rate of scratch received (*r^2^*= 0.048)

|  | Standardized coefficient | Standard error | *p* |
| --- | --- | --- | --- |
| Age | 0.02 | 0.64 | 0.4058 |
| Sex | 0.20 | 0.56 | **0.04598** |
| Kinship | 0.08 | 1.14 | 0.18091 |
| Reproductive status | 0.03 | 0.44 | 0.36332 |
| Piloerection | -0.05 | 0.04 | 0.32784 |

#### Single gestures, rapid and persistence sequences

Supplementary Table S18. MRQAP regression models predicting durations of social behavior, per hour dyad spent within 10m. Predictor variables were rates of single gestures, rapid and persistence sequences between the signaller and the recipient, per hour dyad spent within 10m and demographic variables. Based on 132 chimpanzee dyads. Significant *p* values are indicated in bold. R squared (*r^2^)* denotes amount of variance in the dependent variable explained by the regression model.

**Table S18.1.** Duration of joint feeding behaviour (*r^2^* = 0.157)

|  | Standardized | Standard error | *p* |
| --- | --- | --- | --- |

|  | coefficient |  |  |
| --- | --- | --- | --- |
| Age | -0.03 | 0.67 | 0.3978 |
| Sex | 0.14 | 0.58 | 0.09445 |
| Kinship | 0.28 | 1.31 | **0.008** |
| Reproductive status | 0.25 | 0.56 | **0.0045** |
| Rapid sequence | -0.06 | 0.19 | 0.27286 |
| Single no sequence | 0.08 | 0.06 | 0.16042 |
| Persistence | -0.03 | 0.53 | 0.37131 |

**Table S18.2.** Duration of joint resting behaviour (*r^2^*= 0.077)

|  | Standardized<br>coefficient | Standard error | *p* |
| --- | --- | --- | --- |
| Age | 0.29 | 1.39 | **0.0015** |
| Sex | -0.13 | 1.12 | 0.05497 |
| Kinship | -0.02 | 2.41 | 0.44328 |
| Reproductive<br>status | -0.04 | 1.13 | 0.3993 |
| Rapid sequence | -0.03 | 0.47 | 0.35282 |
| Single no<br>sequence | 0.08 | 0.15 | 0.09995 |
| Persistence | 0.03 | 1.25 | 0.22889 |

**Table S18.3.** Duration of joint travelling behaviour (*r^2^* = 0.259)

|  | Standardized<br>coefficient | Standard error | *p* |
| --- | --- | --- | --- |
| Age | 0.22 | 0.37 | **0.01449** |
| Sex | 0.02 | 0.31 | 0.38581 |
| Kinship | 0.02 | 0.65 | 0.25737 |
| Reproductive<br>status | -0.07 | 0.28 | 0.21689 |
| Rapid sequence | 0.20 | 0.11 | **0.02049** |
| Single no<br>sequence | 0.22 | 0.04 | **0.02699** |
| Persistence | 0.14 | 0.29 | **0.04548** |

**Table S18.4.**
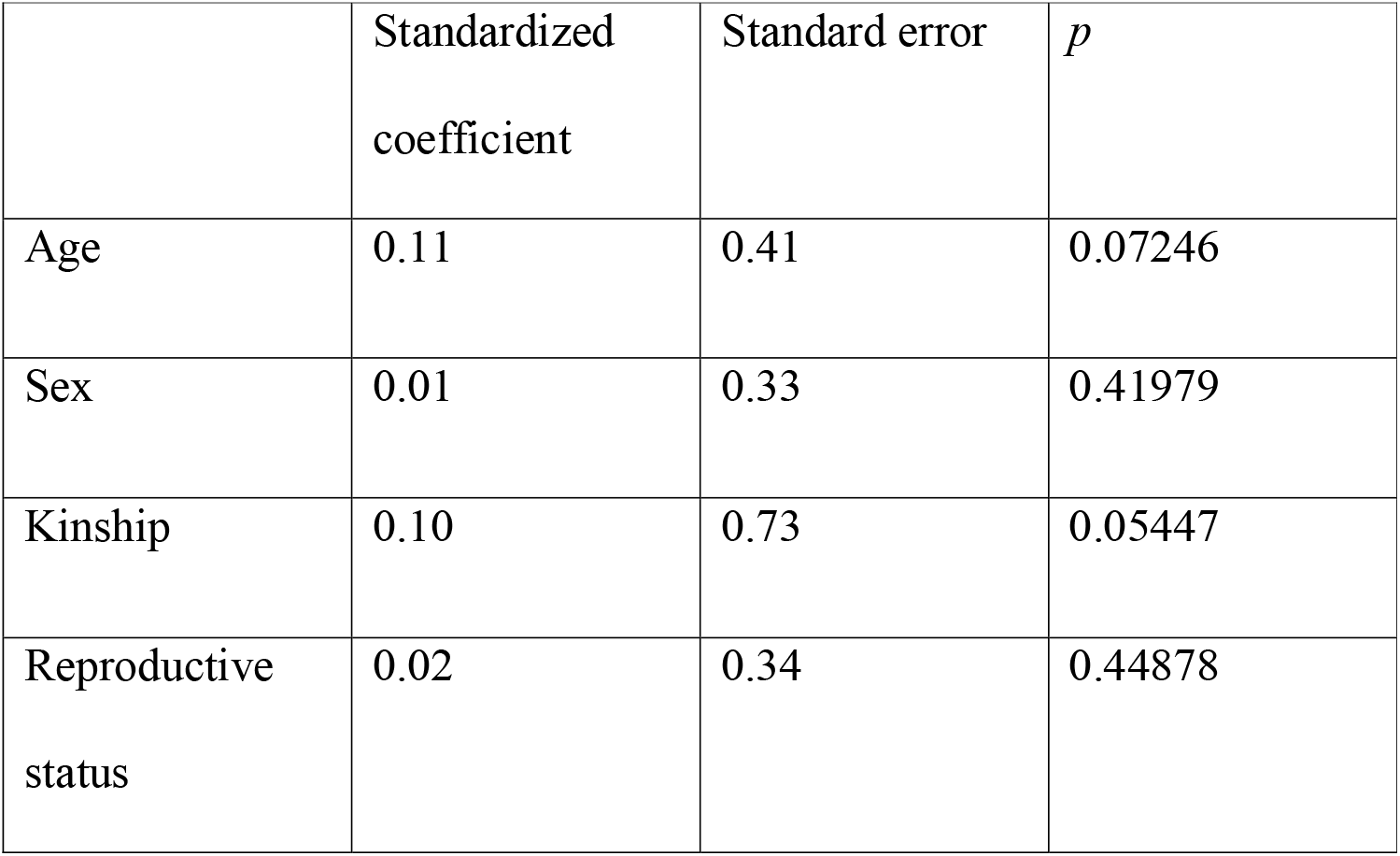
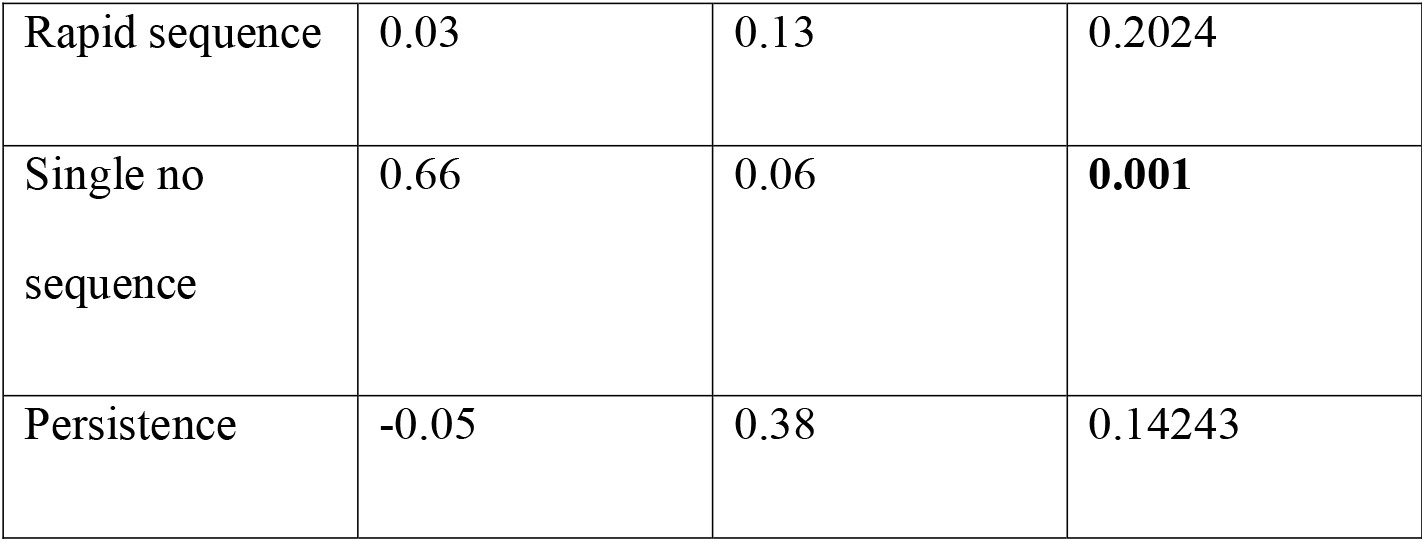
Duration of giving grooming (*r^2^*= 0.474)

**Table S18.5.**
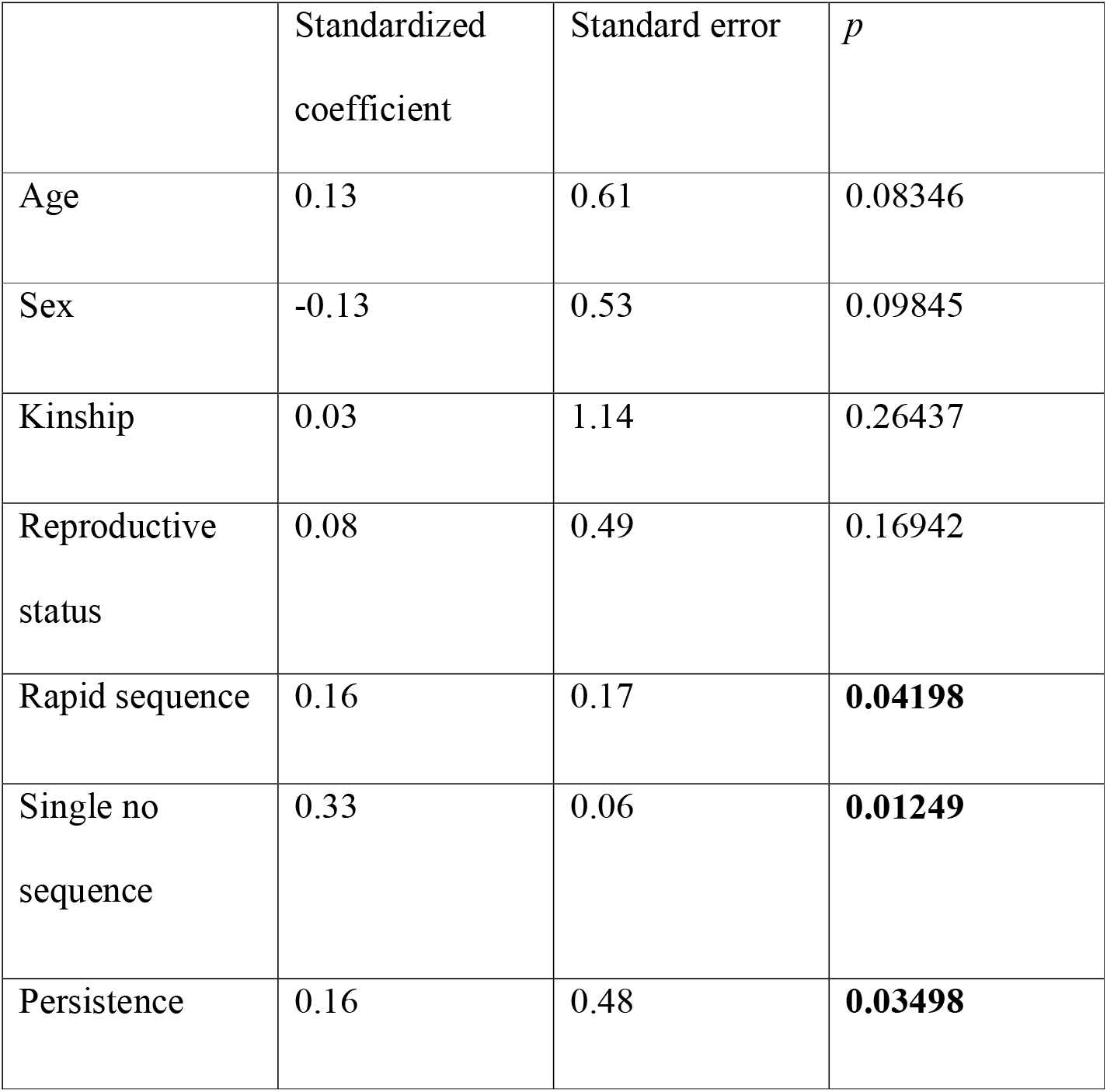
Duration of mutual grooming (*r^2^* = 0.286)

**Table S18.6.** Duration of receiving grooming (*r^2^* = 0.077)

|  | Standardized<br>coefficient | Standard error | *p* |
| --- | --- | --- | --- |
| Age | -0.11 | 0.51 | 0.12044 |
| Sex | 0.11 | 0.43 | 0.13293 |
| Kinship | 0.00 | 0.84 | 0.31384 |
| Reproductive<br>status | 0.12 | 0.46 | 0.09595 |
| Rapid sequence | 0.00 | 0.16 | 0.37331 |
| Single no<br>sequence | 0.16 | 0.05 | 0.05147 |
| Persistence | 0.05 | 0.46 | 0.14343 |

**Table S18.7.** Duration of visual attention towards dyad partner (*r^2^* = 0.433)

|  | Standardized<br>coefficient | Standard error | *p* |
| --- | --- | --- | --- |
| Age | 0.02 | 1.18 | 0.38431 |
| Sex | 0.04 | 0.97 | 0.30135 |
| Kinship | 0.19 | 2.19 | **0.01049** |
| Reproductive<br>status | 0.23 | 1.15 | **0.02099** |
| Rapid sequence | 0.09 | 0.36 | 0.12094 |
| Single no<br>sequence | 0.47 | 0.13 | **0.001** |
| Persistence | 0.09 | 1.04 | 0.12444 |

**Table S18.8.** Duration of visual attention away from dyad partner (*r^2^* = 0.168)

|  | Standardized<br>coefficient | Standard error | *p* |
| --- | --- | --- | --- |
| Age | 0.34 | 1.74 | **0.001** |
| Sex | -0.09 | 1.31 | 0.16492 |
| Kinship | 0.07 | 2.71 | 0.12994 |
| Reproductive<br>status | 0.00 | 1.33 | 0.48226 |
| Rapid sequence | -0.01 | 0.48 | 0.47926 |
| Single no<br>sequence | 0.23 | 0.17 | **0.02649** |
| Persistence | 0.02 | 1.49 | 0.33083 |

**Table S18.9.** Duration of time in close proximity – within 2 m (*r^2^* = 0.354)

|  | Standardized<br>coefficient | Standard error | *p* |
| --- | --- | --- | --- |
| Age | 0.25 | 2.19 | **0.003** |
| Sex | -0.04 | 1.66 | 0.28636 |
| Kinship | 0.15 | 3.68 | **0.02549** |
| Reproductive<br>status | 0.13 | 2.07 | 0.09995 |
| Rapid sequence | 0.04 | 0.65 | 0.27586 |
| Single no<br>sequence | 0.42 | 0.25 | **0.0005** |
| Persistence | 0.06 | 1.86 | 0.1954 |

**Table S18.10.** Rate of scratch produced (*r^2^* = 0.397)

|  | Standardized<br>coefficient | Standard error | *p* |
| --- | --- | --- | --- |
| Age | 0.04 | 0.51 | 0.29735 |
| Sex | 0.12 | 0.46 | 0.10395 |
| Kinship | 0.09 | 0.92 | 0.10645 |
| Reproductive<br>status | -0.02 | 0.34 | 0.4053 |
| Rapid sequence | -0.02 | 0.15 | 0.44528 |
| Single no<br>sequence | 0.08 | 0.05 | 0.12244 |
| Persistence | 0.57 | 0.50 | **0.0005** |

**Table S18.11.**
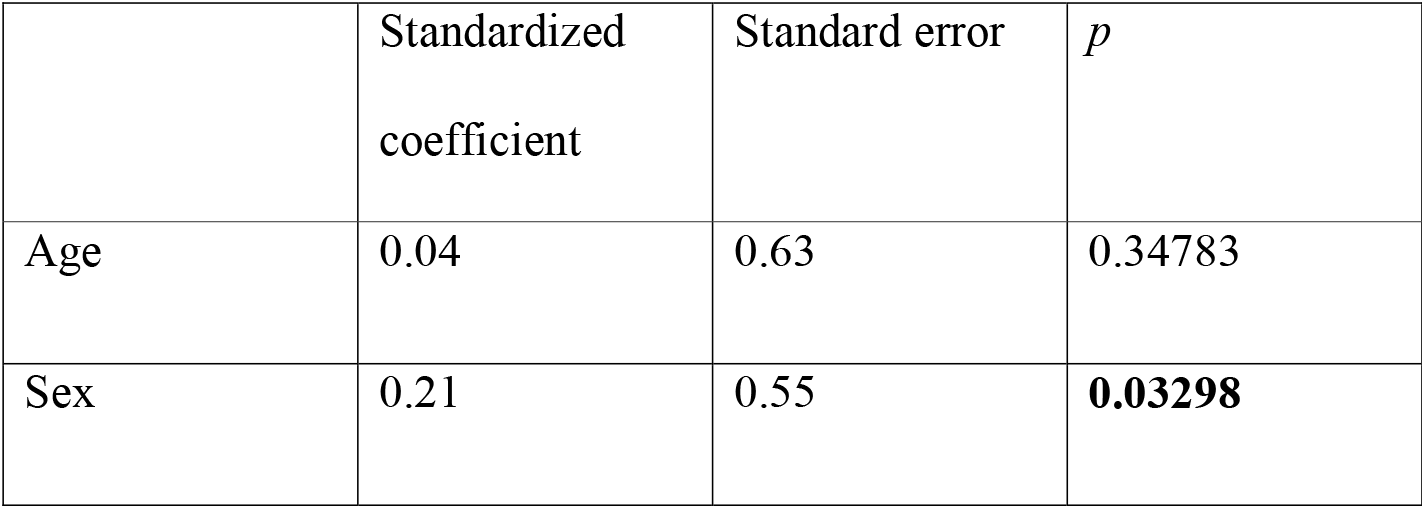
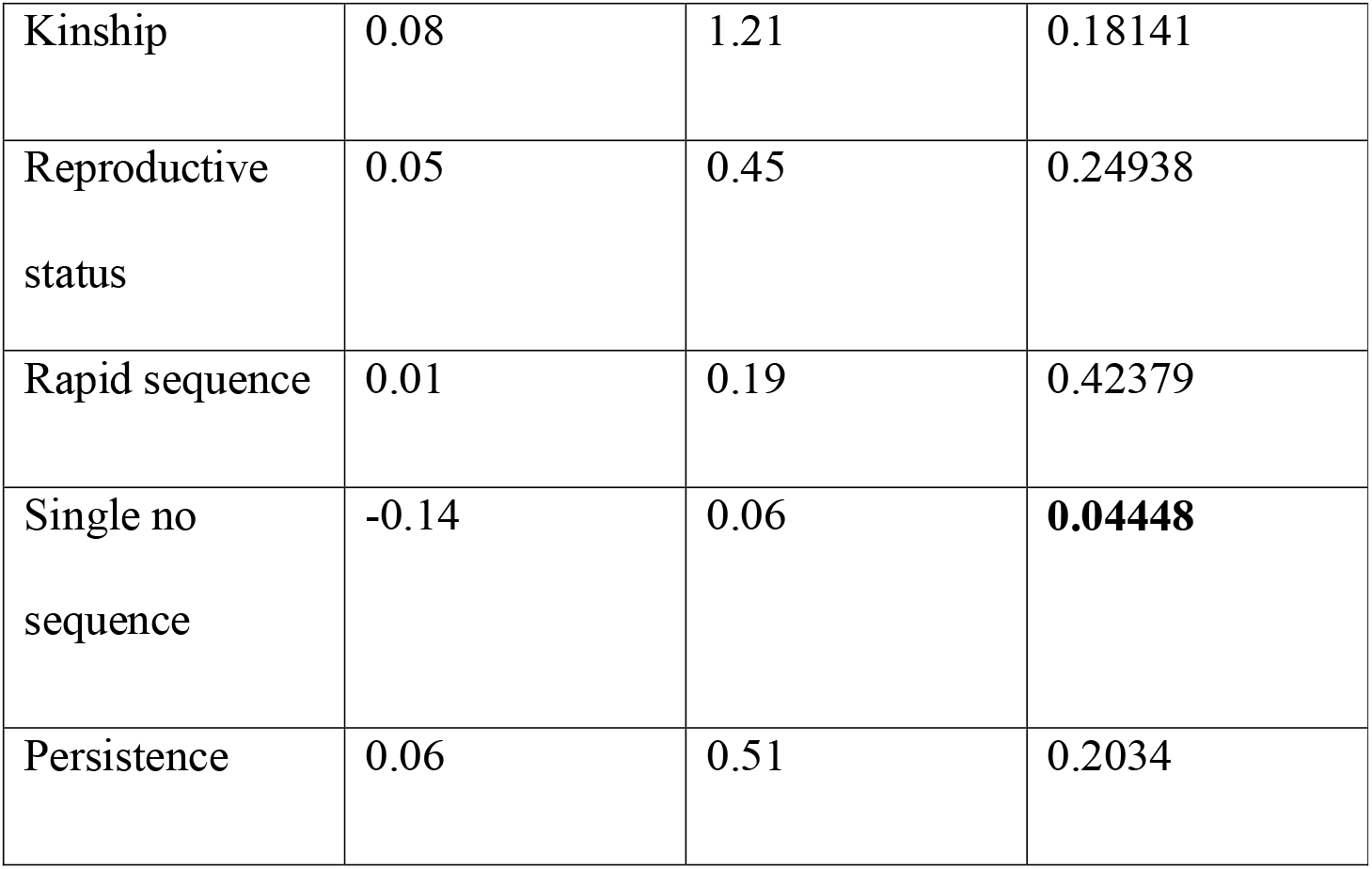
Rate of scratch received (*r^2^* = 0.061)

### Association between the duration of social behaviour and gestural communication categorized according to function

#### Gesture functions

Supplementary Table S19. MRQAP regression models predicting durations of social behavior, per hour dyad spent within 10m. Predictors were demographic variables and functions of gestures. Dyads were classified as same age or different age (within 5 years), same sex or different sex, related by maternal kinship and as the same or different reproductive status (reproductively active, not reproductively active). Based on 132 dyadic relationships of the chimpanzees. Significant *p* values are indicated in bold. R squared (*r^2^)* denotes amount of variance in the dependent variable explained by the regression model.

**Table S19.1.**
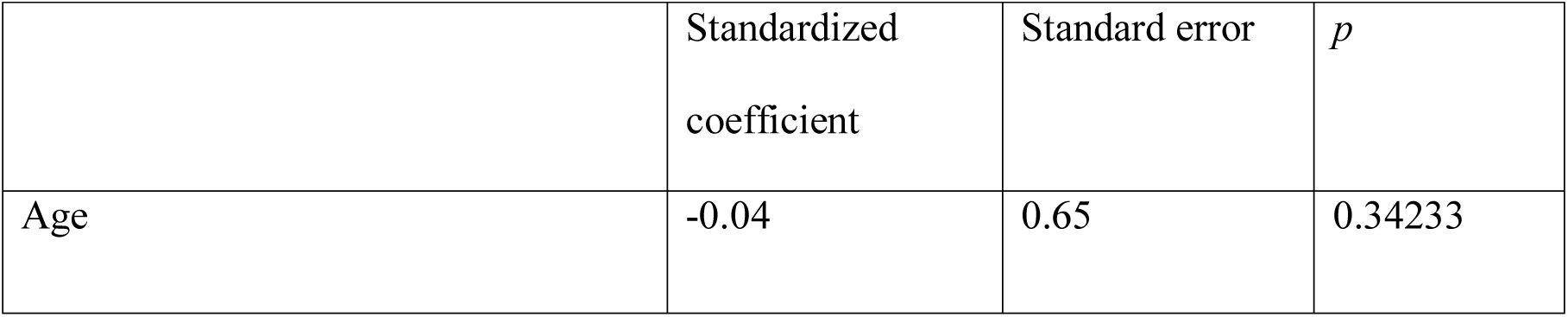
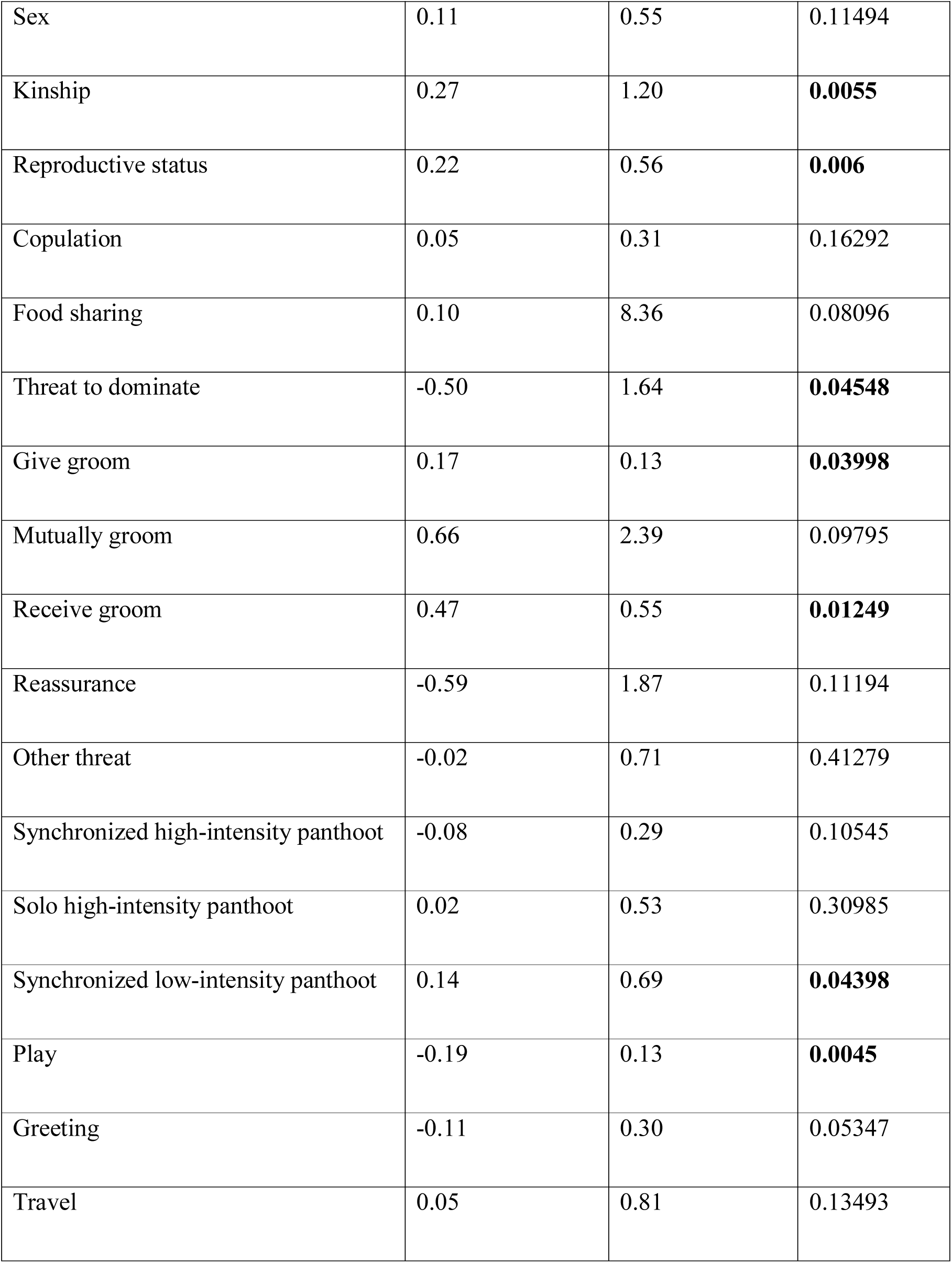
Duration of joint feeding behaviour (*r^2^*= 0.310)

| | Standardized coefficient | Standard error | $p$ |
| --- | --- | --- | --- |
| Age | -0.04 | 0.65 | 0.34233 |
| Sex | 0.11 | 0.55 | 0.11494 |
| Kinship | 0.27 | 1.20 | **0.0055** |
| Reproductive status | 0.22 | 0.56 | **0.006** |
| Copulation | 0.05 | 0.31 | 0.16292 |
| Food sharing | 0.10 | 8.36 | 0.08096 |
| Threat to dominate | -0.50 | 1.64 | **0.04548** |
| Give groom | 0.17 | 0.13 | **0.03998** |
| Mutually groom | 0.66 | 2.39 | 0.09795 |
| Receive groom | 0.47 | 0.55 | **0.01249** |
| Reassurance | -0.59 | 1.87 | 0.11194 |
| Other threat | -0.02 | 0.71 | 0.41279 |
| Synchronized high-intensity panthoot | -0.08 | 0.29 | 0.10545 |
| Solo high-intensity panthoot | 0.02 | 0.53 | 0.30985 |
| Synchronized low-intensity panthoot | 0.14 | 0.69 | **0.04398** |
| Play | -0.19 | 0.13 | **0.0045** |
| Greeting | -0.11 | 0.30 | 0.05347 |
| Travel | 0.05 | 0.81 | 0.13493 |

**Table S19.2.** Duration of joint resting behaviour (*r^2^* = 0.099)

|  | Standardized<br>coefficient | Standard error | *p* |
| --- | --- | --- | --- |
| Age | 0.27 | 1.63 | **0.01499** |
| Sex | -0.12 | 1.25 | 0.08746 |
| Kinship | -0.02 | 2.70 | 0.43428 |
| Reproductive status | -0.05 | 1.28 | 0.35432 |
| Copulation | 0.01 | 0.84 | 0.33883 |
| Food sharing | 0.01 | 23.98 | 0.32234 |
| Threat to dominate | -0.20 | 3.57 | 0.17641 |
| Give groom | 0.11 | 0.31 | 0.06097 |
| Mutually groom | 0.10 | 6.48 | 0.35732 |
| Receive groom | 0.08 | 1.44 | 0.16492 |
| Reassurance | 0.03 | 4.55 | 0.49125 |
| Other threat | -0.03 | 1.75 | 0.32534 |
| Synchronized high-intensity<br>panthoot | -0.07 | 0.66 | 0.09645 |
| Solo high-intensity panthoot | -0.01 | 1.34 | 0.49825 |
| Synchronized low-intensity panthoot | 0.05 | 2.00 | 0.09045 |
| Play | -0.04 | 0.27 | 0.18841 |
| Greeting | 0.04 | 0.75 | 0.17291 |
| Travel | 0.03 | 1.73 | 0.13493 |

**Table S19.3.** Duration of joint travelling behaviour (*r^2^* = 0.420)

|  | Standardized<br>coefficient | Standard error | *p* |
| --- | --- | --- | --- |
| Age | 0.20 | 0.35 | **0.01399** |
| Sex | 0.08 | 0.29 | 0.18191 |
| Kinship | 0.02 | 0.58 | 0.24038 |
| Reproductive status | -0.06 | 0.26 | 0.21139 |
| Copulation | 0.05 | 0.18 | 0.08496 |
| Food sharing | -0.01 | 5.05 | 0.44828 |
| Threat to dominate | 0.03 | 1.10 | 0.34733 |
| Give groom | 0.05 | 0.07 | 0.16042 |
| Mutually groom | 0.03 | 1.42 | 0.44578 |
| Receive groom | 0.08 | 0.32 | 0.17391 |
| Reassurance | 0.37 | 1.02 | 0.12394 |
| Other threat | -0.05 | 0.39 | 0.11044 |
| Synchronized high-intensity panthoot | -0.06 | 0.14 | 0.11694 |
| Solo high-intensity panthoot | 0.10 | 0.30 | 0.05547 |
| Synchronized low-intensity panthoot | 0.26 | 0.45 | **0.01249** |
| Play | -0.03 | 0.07 | 0.2099 |
| Greeting | 0.01 | 0.17 | 0.34783 |
| Travel | 0.10 | 0.44 | **0.04898** |

**Table S19.4.**
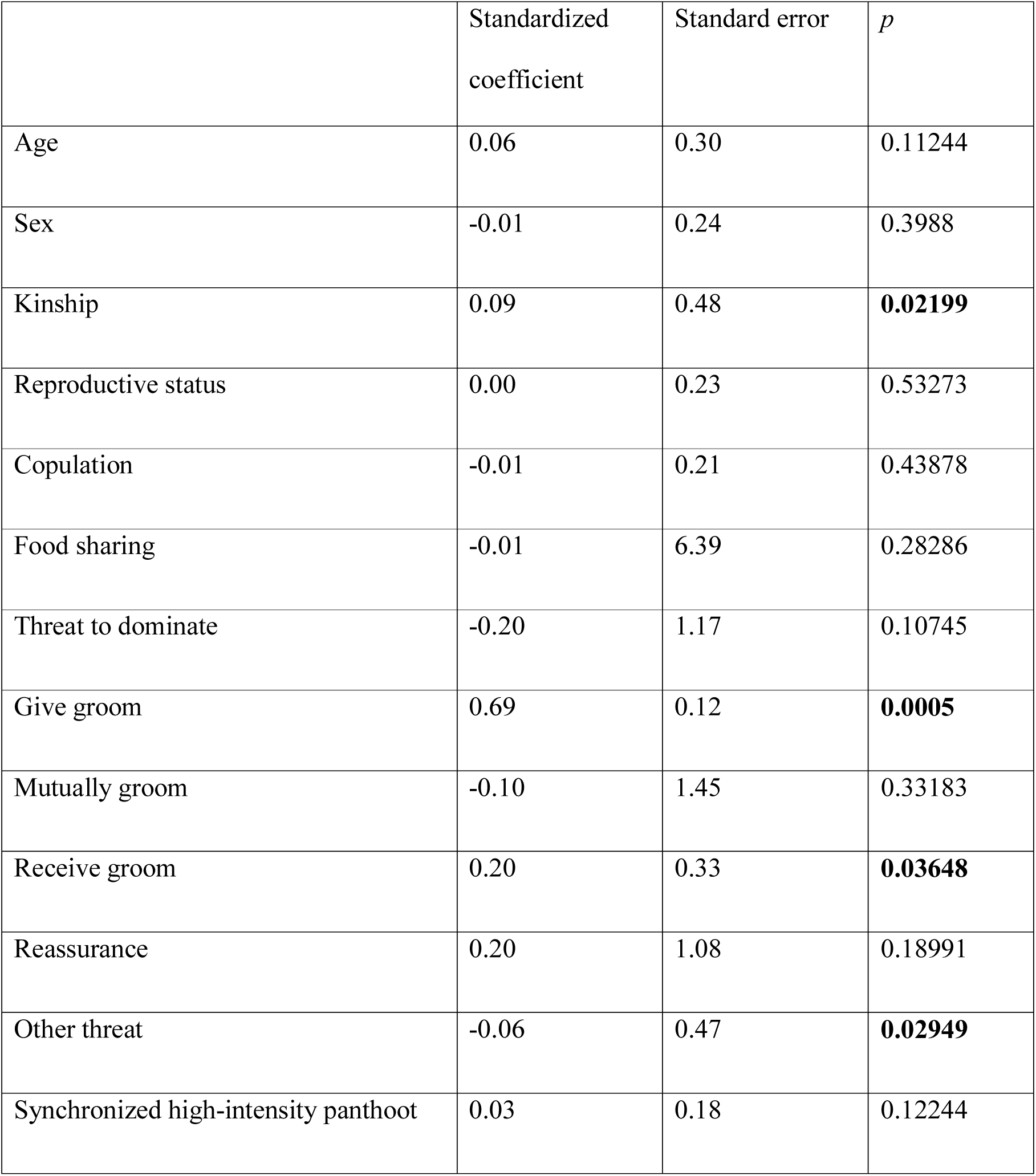
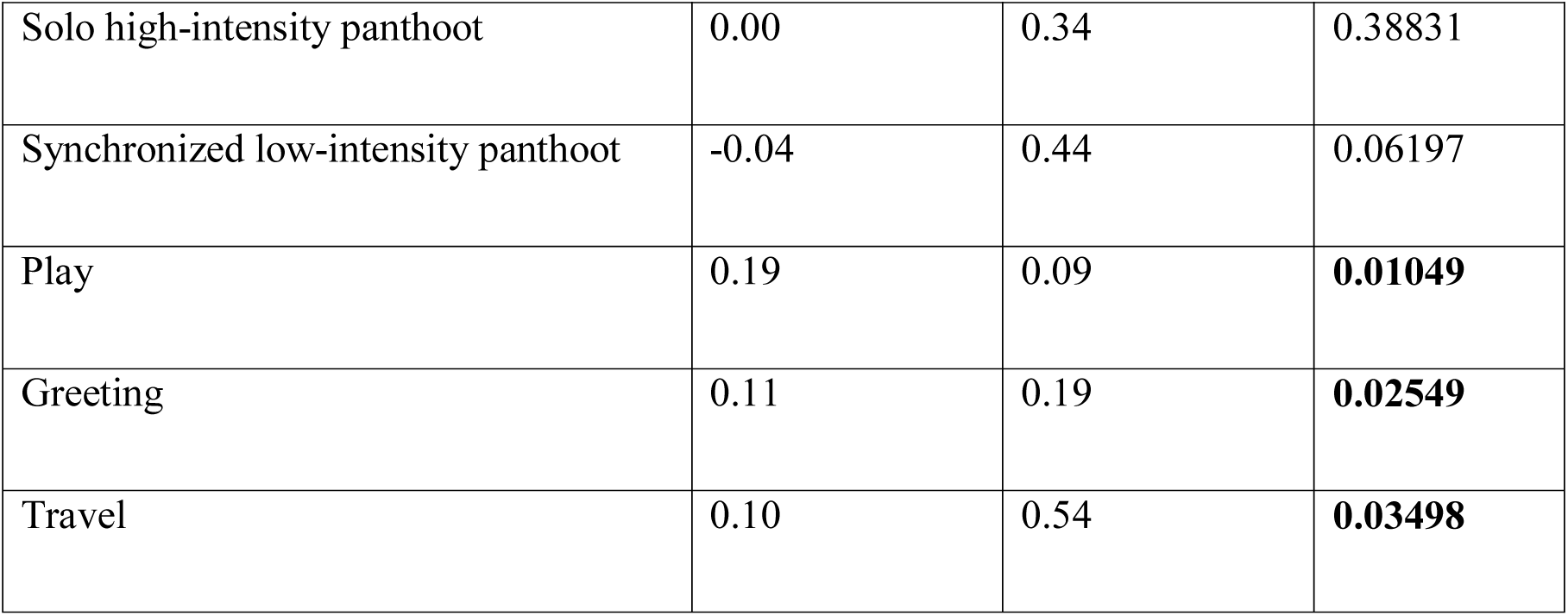
Duration of giving grooming (*r^2^* = 0.740)

**Table S19.5.** Duration of mutual grooming (*r^2^* = 0.586)

|  | Standardized<br>coefficient | Standard error | *p* |
| --- | --- | --- | --- |
| Age | 0.12 | 0.53 | 0.07246 |
| Sex | -0.16 | 0.45 | **0.007** |
| Kinship | -0.01 | 0.90 | 0.58871 |
| Reproductive status | 0.08 | 0.42 | 0.11044 |
| Copulation | 0.01 | 0.25 | 0.27386 |
| Food sharing | -0.02 | 9.54 | 0.28936 |
| Threat to dominate | -0.65 | 1.45 | **0.006** |
| Give groom | 0.03 | 0.11 | 0.21839 |
| Mutually groom | 1.14 | 1.97 | **0.01049** |
| Receive groom | 0.53 | 0.48 | **0.0085** |
| Reassurance | -0.25 | 1.47 | 0.21439 |
| Other threat | -0.03 | 0.58 | 0.31184 |
| Synchronized high-intensity panthoot | 0.01 | 0.27 | 0.5922 |
| Solo high-intensity panthoot | 0.01 | 0.40 | 0.35582 |
| Synchronized low-intensity panthoot | 0.05 | 0.58 | 0.08146 |
| Play | -0.10 | 0.10 | **0.01249** |
| Greeting | -0.05 | 0.21 | 0.11144 |
| Travel | 0.01 | 0.73 | 0.34483 |

**Table S19.6.** Duration of receiving grooming (*r^2^* = 0.338)

|  | Standardized<br>coefficient | Standard error | *p* |
| --- | --- | --- | --- |
| Age | -0.07 | 0.50 | 0.24388 |
| Sex | 0.03 | 0.41 | 0.33033 |
| Kinship | -0.03 | 0.85 | 0.30585 |
| Reproductive status | 0.10 | 0.41 | 0.11844 |
| Copulation | -0.02 | 0.24 | 0.43228 |
| Food sharing | -0.01 | 6.74 | 0.4038 |
| Threat to dominate | 0.04 | 1.29 | 0.32384 |
| Give groom | 0.01 | 0.10 | 0.52524 |
| Mutually groom | -0.52 | 2.01 | 0.07346 |
| Receive groom | 0.90 | 0.53 | **0.001** |
| Reassurance | -0.04 | 1.37 | 0.37881 |
| Other threat | 0.02 | 0.53 | 0.1919 |
| Synchronized high-intensity<br>panthoot | -0.02 | 0.20 | 0.37381 |
| Solo high-intensity panthoot | -0.01 | 0.41 | 0.48626 |
| Synchronized low-intensity panthoot | -0.01 | 0.60 | 0.56172 |
| Play | -0.20 | 0.10 | **0.002** |
| Greeting | -0.08 | 0.25 | **0.04148** |
| Travel | 0.02 | 0.55 | 0.13093 |

**Table S19.7.** Duration of visual attention towards dyad partner (*r^2^* = 0.641)

|  | Standardized<br>coefficient | Standard error | *p* |
| --- | --- | --- | --- |
| Age | 0.00 | 1.07 | 0.47326 |
| Sex | 0.01 | 0.86 | 0.43478 |
| Kinship | 0.16 | 1.96 | **0.01349** |
| Reproductive status | 0.23 | 1.06 | **0.0015** |
| Copulation | 0.02 | 0.53 | 0.26837 |
| Food sharing | 0.01 | 16.60 | 0.28686 |
| Threat to dominate | -0.64 | 2.91 | **0.0045** |
| Give groom | 0.25 | 0.26 | **0.01149** |
| Mutually groom | 0.79 | 3.97 | **0.02549** |
| Receive groom | 0.51 | 0.96 | **0.001** |
| Reassurance | -0.02 | 3.10 | 0.47926 |
| Other threat | -0.01 | 1.16 | 0.51474 |
| Synchronized high-intensity<br>panthoot | -0.07 | 0.48 | 0.07146 |
| Solo high-intensity panthoot | 0.03 | 0.89 | 0.24588 |
| Synchronized low-intensity panthoot | 0.10 | 1.25 | **0.04848** |
| Play | -0.09 | 0.24 | **0.01799** |
| Greeting | -0.02 | 0.60 | 0.41529 |
| Travel | 0.04 | 1.33 | 0.14193 |

**Table S19.8.** Duration of visual attention away dyad partner (*r^2^* = 0.276)

|  | Standardized<br>coefficient | Standard error | *p* |
| --- | --- | --- | --- |
| Age | 0.33 | 1.87 | **0.001** |
| Sex | -0.10 | 1.41 | 0.11044 |
| Kinship | 0.06 | 2.76 | 0.12744 |
| Reproductive status | -0.03 | 1.35 | 0.38731 |
| Copulation | 0.01 | 0.84 | 0.32584 |
| Food sharing | 0.09 | 26.60 | 0.05147 |
| Threat to dominate | 0.10 | 4.63 | 0.25037 |
| Give groom | 0.24 | 0.37 | **0.01999** |
| Mutually groom | -0.10 | 6.45 | 0.43478 |
| Receive groom | 0.38 | 1.55 | **0.02249** |
| Reassurance | -0.29 | 5.47 | 0.2019 |
| Other threat | -0.05 | 2.13 | 0.21489 |
| Synchronized high-intensity<br>panthoot | -0.09 | 0.72 | **0.04998** |
| Solo high-intensity panthoot | 0.01 | 1.47 | 0.33333 |
| Synchronized low-intensity<br>panthoot | 0.13 | 2.06 | **0.03598** |
| Play | -0.04 | 0.33 | 0.2044 |
| Greeting | 0.04 | 0.90 | 0.23938 |
| Travel | 0.06 | 2.13 | 0.09545 |

**Table S19.9.** Duration of time in close proximity – within 2 m (*r^2^* = 0.515)

|  | Standardized coefficient | Standard error | *p* |
| --- | --- | --- | --- |
| Age | 0.23 | 2.15 | **0.004** |
| Sex | -0.06 | 1.62 | 0.16442 |
| Kinship | 0.13 | 3.59 | **0.04448** |
| Reproductive status | 0.11 | 1.77 | 0.07546 |
| Copulation | 0.02 | 0.99 | 0.25287 |
| Food sharing | 0.07 | 29.25 | 0.10595 |
| Threat to dominate | -0.29 | 5.50 | 0.09545 |
| Give groom | 0.30 | 0.47 | **0.0075** |
| Mutually groom | 0.38 | 8.36 | 0.16192 |
| Receive groom | 0.55 | 1.95 | **0.006** |
| Reassurance | -0.21 | 6.33 | 0.27486 |
| Other threat | -0.03 | 2.35 | 0.27936 |
| Synchronized high-intensity<br>panthoot | -0.10 | 0.89 | **0.01299** |
| Solo high-intensity panthoot | 0.03 | 1.70 | 0.25737 |
| Synchronized low-intensity<br>panthoot | 0.14 | 2.59 | **0.02399** |
| Play | -0.08 | 0.40 | **0.04698** |
| Greeting | 0.01 | 1.08 | 0.4013 |
| Travel | 0.06 | 2.56 | 0.09995 |

**Table S19.10.** Rate of scratch produced (*r^2^* = 0.465)

|  | Standardized coefficient | Standard error | *p* |
| --- | --- | --- | --- |
| Age | -0.05 | 0.55 | 0.30035 |
| Sex | 0.17 | 0.49 | **0.03348** |
| Kinship | 0.10 | 0.99 | 0.09045 |
| Reproductive status | 0.01 | 0.41 | 0.46977 |
| Copulation | 0.12 | 0.27 | 0.07396 |
| Food sharing | -0.02 | 7.31 | 0.49425 |
| Threat to dominate | 0.19 | 1.42 | 0.1929 |
| Give groom | 0.10 | 0.11 | 0.10545 |
| Mutually groom | 0.48 | 2.05 | 0.15692 |
| Receive groom | -0.12 | 0.46 | 0.16542 |
| Reassurance | -0.20 | 1.54 | 0.32334 |
| Other threat | 0.48 | 0.65 | **0.002** |
| Synchronized high-intensity<br>panthoot | -0.04 | 0.23 | 0.33933 |
| Solo high-intensity panthoot | -0.04 | 0.45 | 0.29785 |
| Synchronized low-intensity<br>panthoot | 0.11 | 0.59 | 0.07496 |
| Play | -0.05 | 0.11 | 0.1954 |
| Greeting | 0.07 | 0.25 | 0.14793 |
| Travel | -0.04 | 0.60 | 0.27586 |

**Table S19.11.** Rate of scratch received (*r^2^* = 0.095)

|  | Standardized<br>coefficient | Standard error | *p* |
| --- | --- | --- | --- |
| Age | 0.09 | 0.72 | 0.1974 |
| Sex | 0.21 | 0.63 | **0.03448** |
| Kinship | 0.09 | 1.34 | 0.17041 |
| Reproductive status | 0.05 | 0.49 | 0.28336 |
| Copulation | 0.03 | 0.34 | 0.27186 |
| Food sharing | -0.07 | 11.31 | 0.12894 |
| Threat to dominate | 0.01 | 1.88 | 0.3978 |
| Give groom | -0.15 | 0.15 | **0.02099** |
| Mutually groom | 0.20 | 2.67 | 0.34183 |
| Receive groom | 0.10 | 0.60 | 0.24288 |
| Reassurance | -0.35 | 2.01 | 0.26787 |
| Other threat | 0.02 | 0.71 | 0.27536 |
| Synchronized high-intensity panthoot | -0.13 | 0.31 | **0.01949** |
| Solo high-intensity panthoot | 0.05 | 0.52 | 0.22139 |
| Synchronized low-intensity panthoot | 0.09 | 0.67 | 0.10795 |
| Play | -0.05 | 0.14 | 0.28136 |
| Greeting | 0.08 | 0.35 | 0.17341 |
| Travel | 0.01 | 0.83 | 0.35132 |

## Supplementary Information 2 Social bonding, gestural complexity and displacement behaviour of wild chimpanzee

**Table 1.**
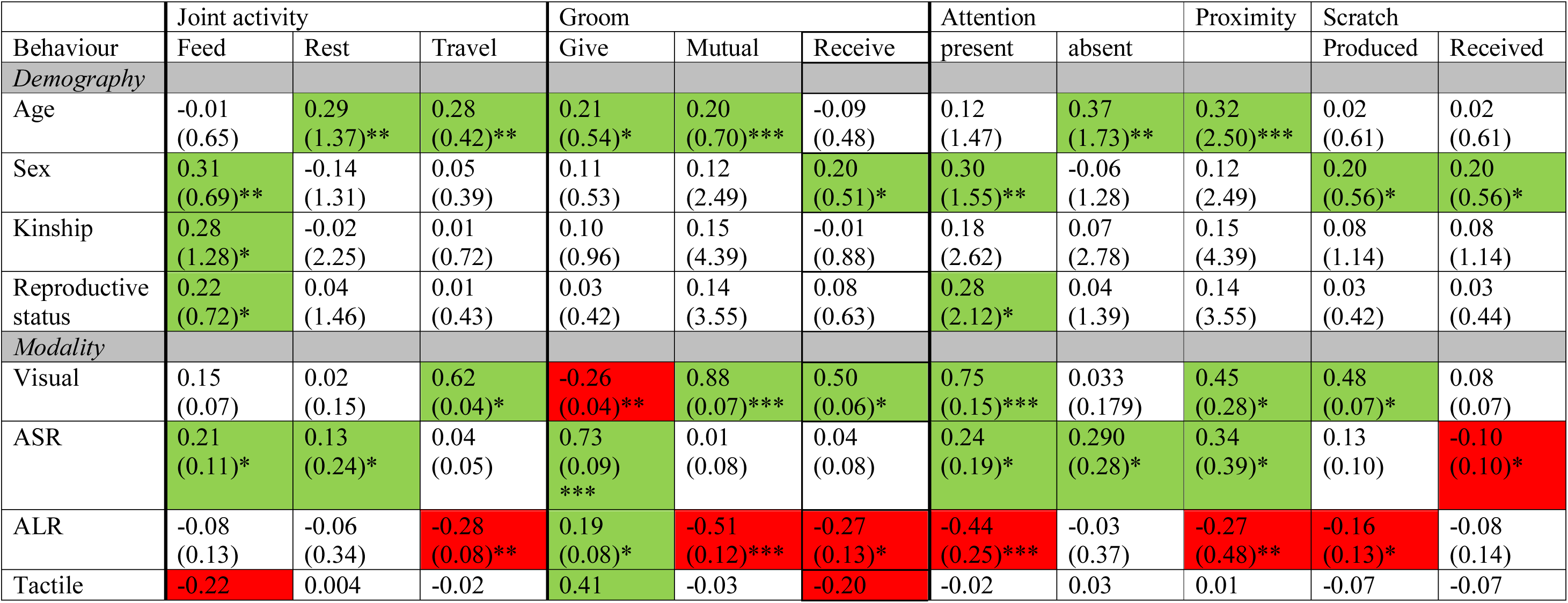
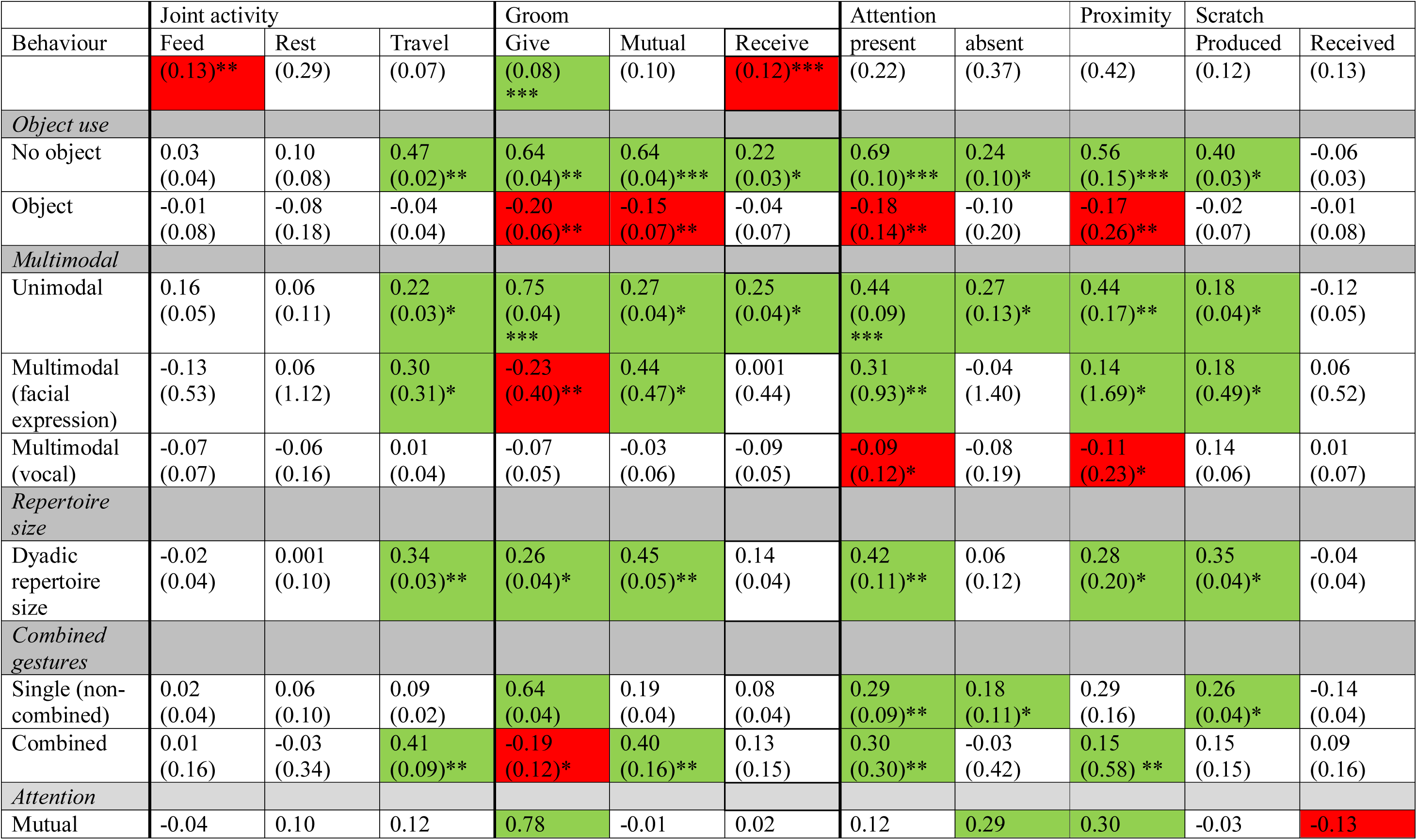
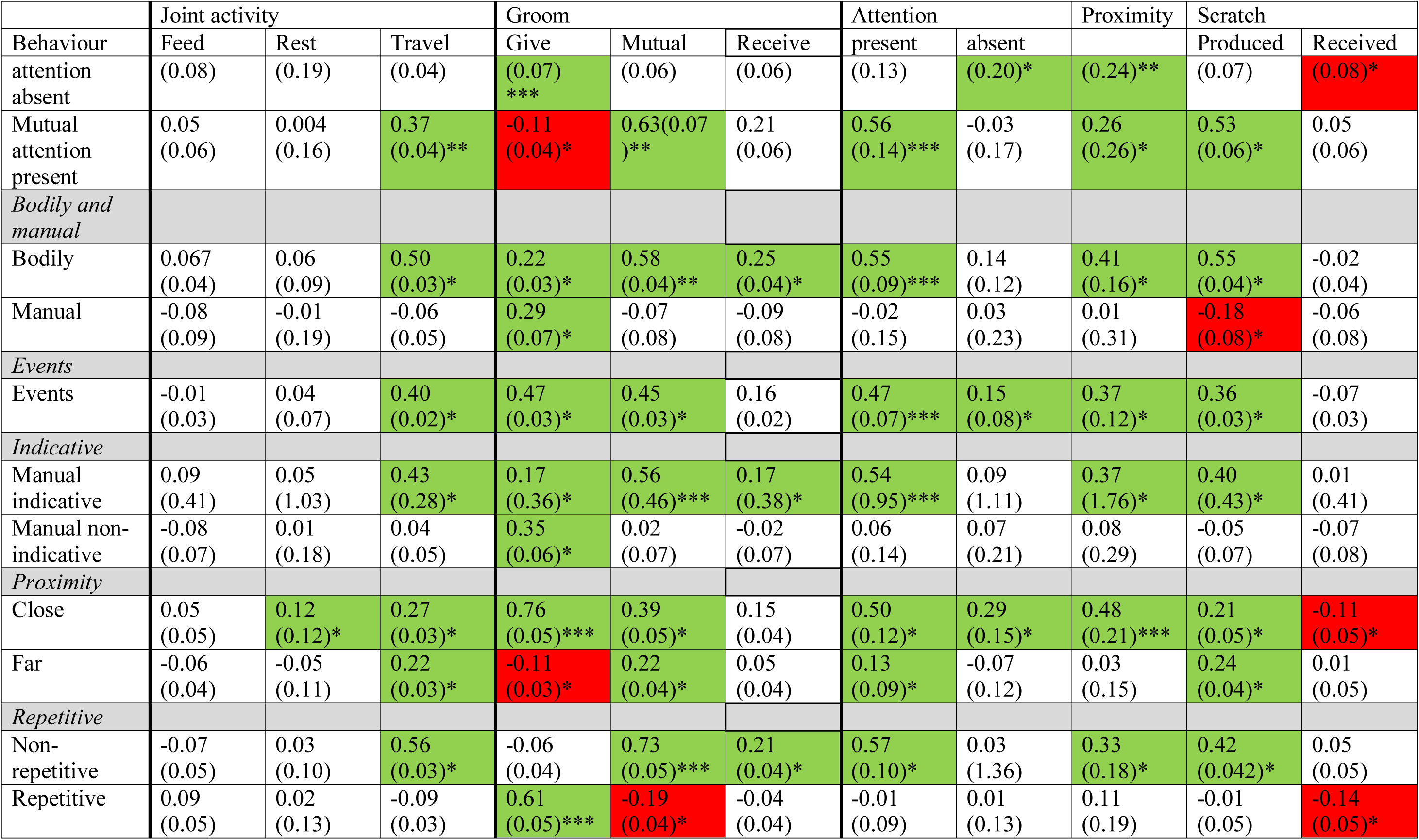
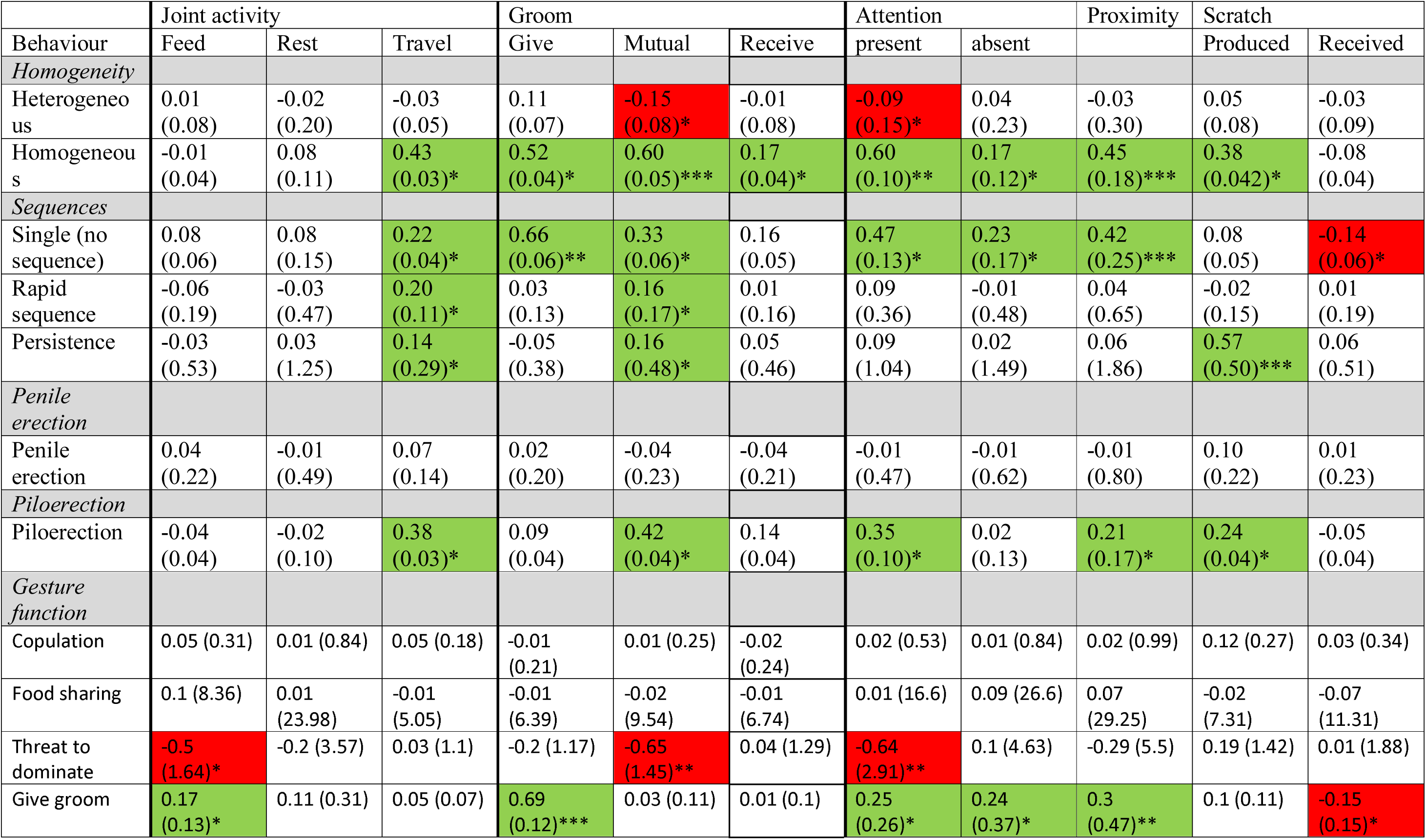
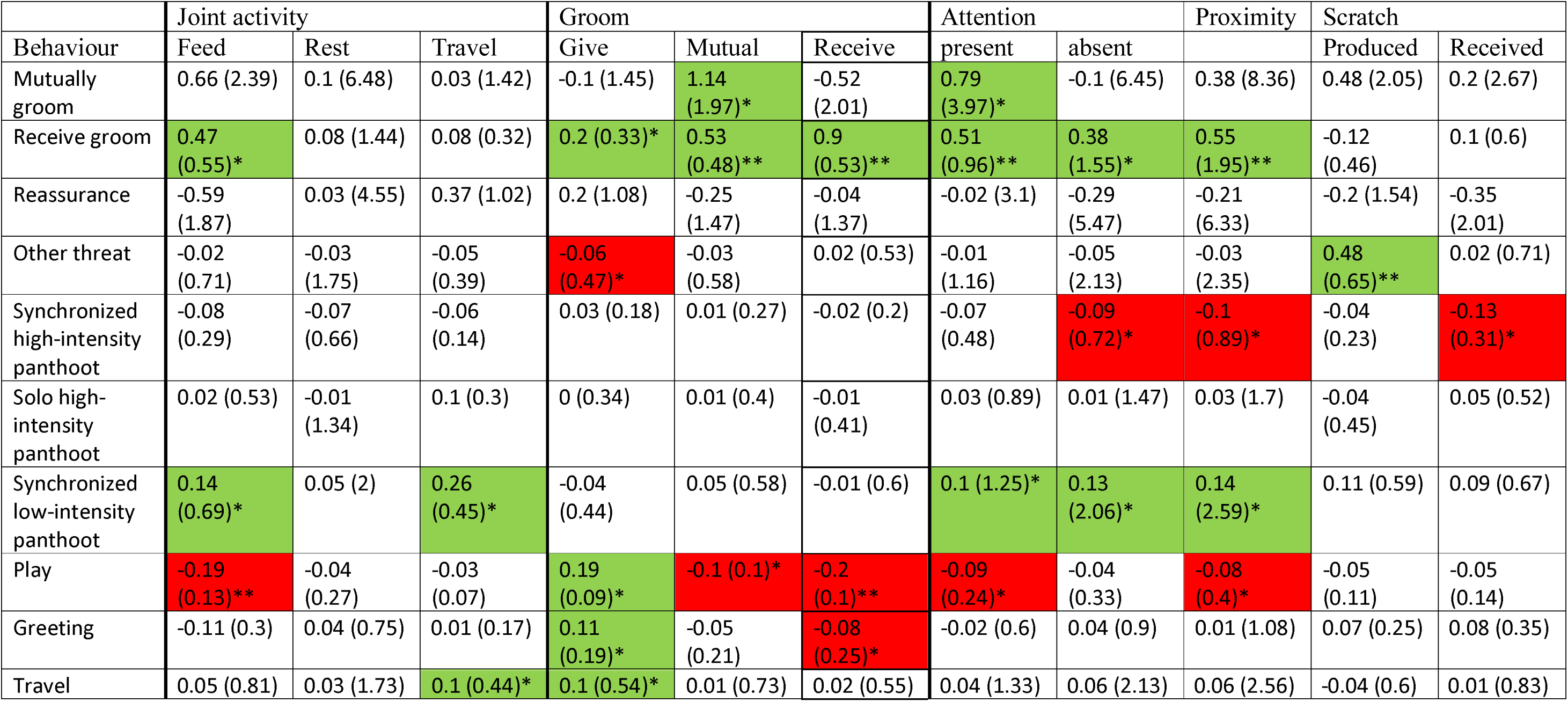
Multiple Regression Quadratic Assignment Procedure (MRQAP) regression models predicting duration of social behaviour and rates of scratch from rates of gestural communication. Summary table provides standardized coefficients (standard errors) and *p* values. In all models, the dependent variable was the duration of behaviour in mins, per hour dyad spent within 10 meters or rates of scratch per hour dyad spent within 10 meters. Shaded lines indicate different MRQAP models for demographic variables and for each type of gestural communication. All models include the control variables relating to the age, sex, kinship and reproductive status of the dyad. Green shading indicates statistically significant positive relationships, red shading indicates statistically significant negative relationships. Full results for all models are provided in Supplementary Tables. * *p* < 0.05, ** *p* < 0.01, *** *p* < 0.001

## References

Altmann, J. (1979). Age cohorts as paternal sibships. Behavioral Ecology and Sociobiology, 6(2), 161–164.

Aureli, F. (1997). Post-conflict anxiety in nonhuman primates: the mediating role of emotion in conflict resolution. Aggressive Behavior: Official Journal of the International Society for Research on Aggression, 23(5), 315–328.

Aureli, F., & Schaik, C. P. v. (1991). Post-conflict behaviour in long-tailed macaques (Macaca fascicularis) II. Coping with the uncertainty. Ethology, 89(2), 101–114.

Ay, N., Flack, J., & Krakauer, D. C. (2007). Robustness and complexity co-constructed in multimodal signalling networks. Philosophical Transactions of the Royal Society B: Biological Sciences, 362(1479), 441–447.

Baker, K. C., & Aureli, F. (1997). Behavioural indicators of anxiety: an empirical test in chimpanzees. Behaviour, 134(13), 1031–1050.

Boesch, C. (1996). Social grouping in Tai chimpanzees. In W. C. McGrew, L. Marchant, & T. Nishida (Eds.), Great Ape Societies (pp. 101–113). Cambridge: Cambridge University Press.

Borgatti, S. P., Everett, M. G., & Johnson, J. C. (2013). Analyzing Social Networks: SAGE Publications Limited.

Castles, D. L., & Whiten, A. (1998). Post-conflict behaviour of wild olive baboons. II. Stress and self-directed behaviour. Ethology, 104(2), 148–160.

Clark, A. P., & Wrangham, R. W. (1994). Chimpanzee arrival pant-hoots: Do they signify food or status? International Journal of Primatology, 15(2), 185–205.

Conty, L., Dezecache, G., Hugueville, L., & Grèzes, J. (2012). Early binding of gaze, gesture, and emotion: neural time course and correlates. The Journal of Neuroscience, 32(13), 4531–4539.

Croft, D. P., James, R., & Krause, J. (2010). Exploring Animal Social Networks. Princeton, New Yersey: Princeton University Press.

Curry, O., Roberts, S. G., & Dunbar, R. I. (2013). Altruism in social networks: Evidence for a ‘kinship premium’. British Journal of Psychology, 104(2), 283–295.

Dawkins, M. S., & Guilford, T. (1997). Conspicuousness and diversity in animal signals. Communication, 55–75.

de Waal, F. B. (1986). The integration of dominance and social bonding in primates. Quarterly Review of Biology, 61(4), 459–479.

De Waal, F. B. M. (1982). Chimpanzee politics: Power and sex among apes. New York: Harper & Row.

Dekker, D., Krackhardt, D., & Snijders, T. A. (2007). Sensitivity of MRQAP tests to collinearity and autocorrelation conditions. Psychometrika, 72(4), 563–581.

Diezinger, F. t., & Anderson, J. (1986). Starting from scratch: A first look at a “displacement activity” in group-living rhesus monkeys. American journal of primatology, 11(2), 117–124.

Dunbar, R. I. M. (1992). Neocortex size as a constraint on group size in primates. Journal of Human Evolution, 20, 469–493.

Feldman, R., Gordon, I., & Zagoory-Sharon, O. (2011). Maternal and paternal plasma, salivary, and urinary oxytocin and parent–infant synchrony: considering stress and affiliation components of human bonding. Developmental science, 14(4), 752–761.

Field, A. (2013). Discovering statistics using IBM SPSS statistics: sage.

Flack, J. C., de Waal, F. B., & Krakauer, D. C. (2005). Social structure, robustness, and policing cost in a cognitively sophisticated species. The American Naturalist, 165(5), E126–E139.

Flack, J. C., De Waal, F. B., & Waal, D. (2004). Dominance style, social power, and conflict management: a conceptual framework. In: Macaque Societies: A Model for the Study of Social Organization (Ed. by.

Flack, J. C., Girvan, M., De Waal, F. B., & Krakauer, D. C. (2006). Policing stabilizes construction of social niches in primates. Nature, 439(7075), 426–429.

Foerster, S., McLellan, K., Schroepfer-Walker, K., Murray, C. M., Krupenye, C., Gilby, I. C., & Pusey, A. E. (2015). Social bonds in the dispersing sex: partner preferences among adult female chimpanzees. Animal Behaviour, 105, 139–152.

Gerhardt, H. C., & Doherty, J. A. (1988). Acoustic communication in the gray treefrog, Hyla versicolor: evolutionary and neurobiological implications. Journal of Comparative Physiology A, 162(2), 261–278.

Goldberg, T. L., & Wrangham, R. W. (1997). Genetic correlates of social behaviour in wild chimpanzees: evidence from mitochondrial DNA. Animal Behaviour, 54(3), 559–570.

Goodall, J. (1986). The Chimpanzees of Gombe: Patterns of Behaviour. Cambridge, Massachusetts: Harward University Press.

Hamilton, W. D. (1964). The genetical evolution of social behaviour. II. Journal of theoretical biology, 7(1), 17–52.

Hewes, G. W. (1973). Primate communication and the gestural origin of language. Current Anthropology, 14, 5–24.

Hinde, R. A. (1976). Interactions, relationships and social structure. Man, 1–17.

Jackson, J. C., Jong, J., Bilkey, D., Whitehouse, H., Zollmann, S., McNaughton, C., & Halberstadt, J. (2018). Synchrony and Physiological Arousal Increase Cohesion and Cooperation in Large Naturalistic Groups. Scientific reports, 8(1), 127. doi:10.1038/s41598-017-18023-4

Kerth, G., Perony, N., & Schweitzer, F. (2011). Bats are able to maintain long-term social relationships despite the high fission–fusion dynamics of their groups. Proceedings of the Royal Society of London B: Biological Sciences, 278(1719), 2761–2767.

Krause, J., Croft, D., & James, R. (2007). Social network theory in the behavioural sciences: Potential applications. Behavioral Ecology and Sociobiology, 62(1), 15–27.

Langergraber, K., Mitani, J., & Vigilant, L. (2009). Kinship and social bonds in female chimpanzees (Pan troglodytes). American journal of primatology, 71(10), 840–851.

Langergraber, K. E., Mitani, J. C., & Vigilant, L. (2007). The limited impact of kinship on cooperation in wild chimpanzees. Proceedings of the National Academy of Sciences, 104(19), 7786–7790.

Liebal, K., Call, J., & Tomasello, M. (2004). Use of gesture sequences in chimpanzees. American journal of primatology, 64(4), 377–396.

Maestripieri, D. (1999). Primate social organization, gestural repertoire size, and communication dynamics: a comparative study of macaques. In B. J. King (Ed.), The origins of language: what nonhuman primates can tell us (pp. 55–77). Santa Fe: School of American Research Press.

Maestripieri, D. (2005). Gestural communication in three species of macaques (Macaca mulatta, M. nemestrina, M. arctoides): Use of signals in relation to dominance and social context. Gesture, 5(1), 55–71.

Massen, J. J., & Koski, S. E. (2014). Chimps of a feather sit together: chimpanzee friendships are based on homophily in personality. Evolution and Human Behavior, 35(1), 1–8.

McComb, K., & Semple, S. (2005). Coevolution of vocal communication and sociality in primates. Biology Letters, 1(4), 381–385.

Mitani, J. C., Watts, D. P., & Muller, M. N. (2002). Recent developments in the study of wild chimpanzee behavior. Evolutionary Anthropology, 11(1), 9–25.

Mitani, J. C., Watts, D. P., Pepper, J. W., & Merriwether, D. A. (2002). Demographic and social constraints on male chimpanzee behaviour. Animal Behaviour, 64(5), 727–737.

Möller, L. M., Beheregaray, L. B., Harcourt, R. G., & Krützen, M. (2001). Alliance membership and kinship in wild male bottlenose dolphins (Tursiops aduncus) of southeastern Australia. Proceedings of the Royal Society of London B: Biological Sciences, 268(1479), 1941–1947.

Muller, M. N. (2002). Agonistic relations among Kanyawara chimpanzees. Behavioural diversity in chimpanzees and bonobos, 112–124.

Murray, C. M., Lonsdorf, E. V., Stanton, M. A., Wellens, K. R., Miller, J. A., Goodall, J., & Pusey, A. E. (2014). Early social exposure in wild chimpanzees: Mothers with sons are more gregarious than mothers with daughters. Proceedings of the National Academy of Sciences, 111(51), 18189–18194.

Nakayama, K., Goto, S., Kuraoka, K., & Nakamura, K. (2005). Decrease in nasal temperature of rhesus monkeys (Macaca mulatta) in negative emotional state. Physiology & behavior, 84(5), 783–790.

Nishida, T., Zamma, K., Matsusaka, T., Inaba, A., & McGrew, W. C. (2010). Chimpanzee behavior in the wild: An audio-visual encyclopedia. Tokyo: Springer.

Noë, R. (2006). Cooperation experiments: coordination through communication versus acting apart together. Animal Behaviour, 71(1), 1–18.

Ortega-Ortiz, J. G., Engelhaupt, D., Winsor, M., Mate, B. R., & Rus Hoelzel, A. (2012). Kinship of long-term associates in the highly social sperm whale. Molecular Ecology, 21(3), 732–744.

Payne, R. J., & Pagel, M. (1997). Why do animals repeat displays? Animal Behaviour, 54(1), 109–119.

Pearce, E., Wlodarski, R., Machin, A., & Dunbar, R. I. (2017). Variation in the β-endorphin, oxytocin, and dopamine receptor genes is associated with different dimensions of human sociality. Proceedings of the National Academy of Sciences USA, 201700712.

Roberts, A. I. (2018). Influence of party size on social bonding and gestural persistence in wild chimpanzees. Advances in Biology and Earth Scienes, 3(3), 205–228.

Roberts, A. I., Chakrabarti, A., & Roberts, S. G. B. (2019). Gestural repertoire size is associated with social proximity measures in wild chimpanzees. American journal of primatology, e22954. doi:10.1002/ajp.22954

Roberts, A. I., & Roberts, S. G. B. (2015). Gestural communication and mating tactics in wild chimpanzees. PloS One, 10(11), e0139683. doi:10.1371/journal.pone.0139683

Roberts, A. I., & Roberts, S. G. B. (2016). Wild chimpanzees modify modality of gestures according to the strength of social bonds and personal network size. Scientific reports, 6(33864). doi:10.1038/srep33864

Roberts, A. I., & Roberts, S. G. B. (2017). Convergence and divergence in gestural repetoires as an adaptive mechanism for social bonding in primates. Royal Society Open Science, 4: 170181. doi:https://doi.org/10.1098/rsos.170181

Roberts, A. I., & Roberts, S. G. B. (2018). Persistence in gestural communication predicts sociality in wild chimpanzee. Animal Cognition. doi:10.1101/365858

Roberts, A. I., Roberts, S. G. B., & Vick, S.-J. (2014). The repertoire and intentionality of gestural communication in wild chimpanzees. Animal Cognition, 17(2), 317–336. doi:10.1007/s10071-013-0664-5

Roberts, A. I., Vick, S.-J., & Buchanan-Smith, H. (2012). Usage and comprehension of manual gestures in wild chimpanzees. Animal Behaviour, 84(2), 459–470. doi:10.1016/j.anbehav.2012.05.022

Roberts, A. I., Vick, S.-J., & Buchanan-Smith, H. (2013). Communicative intentions in wild chimpanzees: Persistence and elaboration in gestural signalling. Animal Cognition, 16(2), 187–196. doi:10.1007/s10071-012-0563-1

Roberts, A. I., Vick, S.-J., Roberts, S. G. B., Buchanan-Smith, H. M., & Zuberbühler, K. (2012). A structure-based repertoire of manual gestures in wild chimpanzees: Statistical analyses of a graded communication system. Evolution and Human Behavior, 33(5), 578–589. doi:10.1016/j.evolhumbehav.2012.05.006

Roberts, A. I., Vick, S.-J., Roberts, S. G. B., & Menzel, C. R. (2014). Chimpanzees modify intentional gestures to coordinate a search for hidden food. Nature Communications 5, 3088. doi:10.1038.ncomms4088

Roberts, S., & Roberts, A. I. (2018). Visual attention, indicative gestures, and calls accompanying gestural communication are associated with sociality in wild chimpanzees (Pan troglodyres schweinnfurthii). Journal of Comparative Psychology. doi:10.1037/com0000128

Roberts, S. G., & Dunbar, R. I. (2011). The costs of family and friends: an 18-month longitudinal study of relationship maintenance and decay. Evolution and Human Behavior, 32(3), 186–197.

Roberts, S. G. B., & Roberts, A. I. (2016). Social brain hypothesis, vocal and gesture networks of wild chimpanzees. Frontiers in psychology, 7(1756). doi:10.3389/fpsyg.2016.01756

Schino, G., Perretta, G., Taglioni, A. M., Monaco, V., & Troisi, A. (1996). Primate displacement activities as an ethopharmacological model of anxiety. Anxiety, 2(4), 186–191.

Schino, G., Scucchi, S., Maestripieri, D., & Turillazzi, P. G. (1988). Allogrooming as a tension reduction mechanism: A behavioral approach. American journal of primatology, 16(1), 43–50.

Silk, J. B., Seyfarth, R. M., & Cheney, D. L. (2018). Quality versus quantity: do weak bonds enhance the fitness of female baboons? Animal Behaviour, 140, 207–211.

Silk, M. J., Jackson, A. L., Croft, D. P., Colhoun, K., & Bearhop, S. (2015). The consequences of unidentifiable individuals for the analysis of an animal social network. Animal Behaviour, 104, 1–11.

Van Schaik, C. P., & Van Hooff, J. A. (1994). Male bonds: afilliative relationships among nonhuman primate males. Behaviour, 130(3-4), 309–337.

Watts, D. P. (1992). Social relationships of immigrant and resident female mountain gorillas. I. Male-female relationships. American journal of primatology, 28(3), 159–181.

Widdig, A., Nurnberg, P., Krawczak, M., Streich, W. J., & Bercovitch, F. (2002). Affiliation and aggression among adult female rhesus macaques: a genetic analysis of paternal cohorts. Behaviour, 139(2-3), 371–392.

Wrangham, R. W. (1980). An ecological model of female-bonded primate groups. Behaviour, 75(3), 262–300.

Zahavi, A., & Zahavi, A. (1997). The handicap principle: a missing piece of Darwin’s puzzle: Oxford University Press.

Zajonc, R. B., & Sales, S. M. (1966). Social facilitation of dominant and subordinate responses. Journal of Experimental Social Psychology, 2(2), 160–168.

## References

Hobaiter K, Byrne R. 2011. Serial gesturing by wild chimpanzees: Its nature and function for communication. Animal Cognition 14:827–838.

Nishida T, Zamma K, Matsusaka T, Inaba A, McGrew WC. 2010. Chimpanzee behavior in the wild: An audio-visual encyclopedia. Tokyo: Springer.

Roberts AI, Roberts SGB, Vick S-J. 2014. The repertoire and intentionality of gestural communication in wild chimpanzees. Animal Cognition 17(2):317–336.

